# Exploring genetic signatures of zoonotic influenza A virus at the swine-human interface with phylogenetic and ancestral sequence reconstruction

**DOI:** 10.1101/2024.10.26.619880

**Authors:** K.M. Anker, M.M. Ciucani, J.N. Nissen, T.K. Anderson, A.G. Pedersen, R. Trebbien

## Abstract

Influenza A viruses (IAVs) in swine have zoonotic potential and pose a continuous threat of causing human pandemics, as demonstrated by the H1N1 pandemic in 2009. Despite increased genomic surveillance, we have limited knowledge of the IAV evolutionary dynamics leading to such zoonotic events and no clear understanding of genetic markers associated with interspecies transmission of IAV between humans and swine. To explore this, we analyzed a comprehensive publicly available whole genome dataset of human and swine IAV sequences. We conducted phylogenetic analyses and inference of ancestral host and sequence states for each IAV segment and mapped inferred mutations that were associated with transmission within and between swine and human hosts. We developed a custom python library to combine information from host and ancestral sequence annotated trees and applied statistical models to identify genetic markers associated with intra- or interspecies transmissions between swine and humans. This included analyzing mutation rates and the selective pressures acting on the viral proteins following intra- and interspecies transmissions and using a scalable, gradient-boosted decision tree machine learning approach to predict key amino acid positions critical for different transmission types. Our analyses indicated complex mutational patterns within and across viral proteins, but also suggested that specific protein regions and amino acid positions of especially several of the internal gene segments were more important for interspecies transmission. Our findings identify potential genetic signatures across the IAV proteins associated with host adaptation and zoonotic potential, offering valuable markers for early-warning genomic surveillance systems to enhance animal health and minimize the potential for zoonotic transmission of IAV.

## Introduction

Influenza A virus (IAV) is a significant public health concern due to its ability to cause seasonal epidemics and occasional pandemics. Its capacity to infect a wide range of hosts, including birds, swine, human, and other mammals, is an important factor in the evolution and adaptability of the virus that impacts its zoonotic potential [1]. The IAV genome is composed of 8 segments of negative sense RNA, each encoding one or more proteins: the polymerase-encoding segments, polymerase basic (PB2 and PB1) and polymerase acidic (PA), the surface proteins hemagglutinin (HA) and neuraminidase (NA), the nucleoprotein (NP), the matrix proteins (MP segment, M1 and M2 proteins) and the nonstructural proteins (NS segment, NS1 and NEP proteins) [2]. Additionally, some lineages of IAV express accessory proteins like PB1-F2 and PA-X, encoded by alternative reading frames in the PB1 and PA segments, respectively, that may affect virus pathogenicity and host immune responses [3–5]. The segmented nature of the IAV genome, combined with an error-prone RNA-dependent RNA polymerase, contributes to its evolution and the emergence of frequent immune-escaping variants, resulting in a high degree of diversity among circulating lineages [6,7].

Swine are critical hosts associated with zoonotic transmission of IAVs, acting not only as potential intermediates in the generation of novel strains, but as important contributors to the diversification of IAVs [8]. The bidirectional transmission between humans and pigs has significantly influenced the evolutionary history of IAV in both species, with almost every human pandemic resulting in reverse zoonotic transmission of different H1 and H3 subtype IAVs into swine populations globally [9,10]. This dynamic has led to a complex array of IAV genotypes in swine, including reassortants with human-origin segments, which pose an increasing risk of introduction back into the human population [10–14]. Even though zoonotic transmission between swine and humans is sporadic [9], IAVs of both avian and swine origin have been responsible for numerous human pandemics, including the 2009 pandemic caused by a reassorted H1N1 IAV from swine (H1N1pdm09) [15–17].

Knowledge of specific genetic factors that facilitated the transmission of the H1N1pdm09 or other swine IAVs to humans is limited. Some markers of host adaptation are well-established, like amino acid positions affecting the receptor binding specificity or stability of the HA protein [18], or the efficiency of the IAV polymerase complex including the well-characterized E627K mutation [19], or analogous mutations G590S and Q591R observed in the H1N1pdm09 virus [20,21]. However, most of the characterization of IAV host adaptation markers has focused on avian IAVs and their adaptation for transmission in mammalian hosts [1,22–26], while markers for swine-to-human IAV transmissions are less well characterized or focused on specific lineages and a subset of viral proteins [20,27,28]. Recent advancements in bioinformatics and machine learning offer new avenues for exploring these complex genetic interactions. Advanced algorithms can now analyze vast genomic datasets to uncover patterns and potential biomarkers indicative of zoonotic risk which can enhance our understanding of viral adaptation and improve our ability to anticipate and respond to future pandemic strains [29–31].

In this study, we systematically explore the genetic factors implicated in the zoonotic and reverse zoonotic transmission of IAVs between swine and humans. Our approach combines comprehensive phylogenetic analysis across all IAV lineages in swine and humans with ancestral host and sequence reconstruction to trace and compare genetic changes that occur through intra- and interspecies transmission. We apply statistical methods and train a predictive machine learning model to identify recurring genetic patterns associated with intra- or interspecies transmissions and gain insights into the molecular mechanisms underlying the zoonotic potential of IAVs.

## Methods

### Data collection, quality control, and creating a representative IAV dataset

We downloaded nucleotide sequence data from all eight genomic segments of IAVs of human and swine origin from both GISAID [32] and NCBI Influenza Virus Resource [33] databases (data downloaded between 2023.01.26-2023.02.02). Only viruses with an H1 or H3 subtype HA segment combined with an N1 or N2 subtype NA segment were selected, and from the NCBI database, identical sequences were collapsed and represented by the oldest sequence for download.

The downloaded data was stored in separate files for each segment and host species. We used a modified Sequence Cleaner script from Biopython [34] (sequence_cleaner.py) to clean the fasta files by removing duplicate or poor-quality sequences, including sequences shorter than or much longer than the coding regions, with missing or more than five ambiguous bases, or labeled as “synthetic” or “environment” samples.

The human and swine datasets were then combined for each segment, and HA and NA segments were divided into datasets for each subtype, H1, H3, N1 and N2. The datasets were then down-sampled using CD-HIT v4.8.1 [35] only keeping sequences with less than 97% pairwise identity (settings: cd-hit-est -c 0.970 -n 11 -d 0 -M 16000 -T 4), while ensuring that we did not discard closely related sequences isolated from different host-species. Specifically, if a CD-HIT cluster (with more than 97% identity) contained sequences isolated from both human and swine, we re-extracted up to 5 sequences from each host. The datasets were then annotated with the NCBI Influenza Sequence Annotation Tool (https://www.ncbi.nlm.nih.gov/genomes/FLU/annotation/), and sequences with missing start or stop codons or frameshifts in the coding region of the major proteins were removed. This resulted in final datasets of ∼500-1000 sequences per segment.

### Phylogenetic analysis and reconstruction of ancestral states

Sequences in each dataset were aligned with mafft v7.487 [36] using the “--auto” option and alignments were inspected for poorly aligned sequences causing large gaps with Aliview v1.27 [37]. These sequences were removed before realignment of the remaining sequences. Finally, columns in the alignment with >40% fraction of gaps were removed from the alignment using seqconverter [38] Maximum likelihood phylogenetic trees were initially inferred with IQ-TREE v2.0.3 [39] using ModelFinder [40] to choose the best substitution model and ultrafast bootstrap approximation (UFBoot) [41] (settings: -nt auto -ntmax 16 -nm 3000 -bb 1000 -asr). The maximum likelihood trees were then used as input for inference of molecular-clock phylogenies and ancestral sequence and host state reconstruction in TreeTime v0.11.1 [42]. First, time-scaled phylogenetic trees were estimated based on the initial tree topologies and tip dates, while also inferring ancestral sequences of internal nodes in the trees (treetime --tree tree.nwk --dates dates.csv --aln aln.fasta). Most segments were fitted with a relaxed molecular clock, however for some segments, specifically the PB2 and PA segments, fitting to a strict molecular clock while accounting for covariance was more appropriate and led to better inference of the overall evolutionary rate. The resulting root-to-tip regression output was used to check for outliers, which were further manually inspected to evaluate incongruence by assessing metadata or through BLAST (e.g., to see if a sequence was derived from an avian-to-swine transmission) and problematic sequences were then removed from the dataset. If any sequences were removed in this step, the remaining data were re-aligned and new maximum likelihood and temporally scaled trees were inferred. The final molecular-clock trees were used as input trees for inference of ancestral host states using the TreeTime “mugration” subcommand and a state file with host states (either swine or human) for each tip in the tree. The flag “--confidence” was used to get an output file with the inferred host state probabilities at each internal node in the tree. Host annotated time trees were visualized using Figtree v1.4.4 (http://tree.bio.ed.ac.uk/software/figtree/).

### Summarizing sequence and mutation data across all phylogenetic branches representing inter- or intraspecies transmissions

We used the python library, anclib v0.11.0 (https://github.com/agormp/anclib), to analyze information about host states and ancestral sequences for all nodes in the trees and create the final datasets for sequence comparisons. For these analyses, we kept track of sequences and host states for all nodes in the trees as well as sequence changes along each branch. We also grouped the different host transmission types (swine-to-swine, human-to-human, human-to-swine, or swine-to-human) for additional analyses. Only branches with nodes having a minimum 60% host state probability were included in the analysis.

Processing and plotting of output files from anclib listing full sequences or amino acid changes along each branch in the phylogenetic trees were performed in R using tidyverse and ggplot2 [43–45]. Sequences from branches with a length >15 years were removed from the datasets, as long branches in the phylogenies would be expected to lead to larger uncertainties in the ancestral inference of connected nodes and often represent connections between different IAV lineages with lower rates of sequence sampling.

### Bayesian detection of sites important for species-specificity

To identify protein sites important for species-specificity, we performed an analysis of whether some sites displayed higher rates of substitution following a jump between species, compared to when the virus remained within a host-type. Assuming that an amino acid at a given site was conserved within a species, but more often changed following a jump to the other species, this could indicate selection for a specific amino acid with higher fitness in the new species.

Specifically, for each variable site in the protein alignments, we estimated separate substitution rates for the four types of viral transmission: swine-to-swine, human-to-human, human-to-swine, and swine-to-human. For this purpose, we relied on the ancestral reconstructions of host states and amino acid sequences produced by TreeTime: each internal node in the tree corresponded to a hypothetical ancestor virus, for which TreeTime inferred the most likely host state (human or swine) and amino acid sequence. Based on observed and inferred host states we classified each branch in the phylogeny as belonging to one of the four transmission types (e.g. if the host-state was human for the parent-node and swine for the child-node, then that branch corresponded to a reverse zoonotic jump from humans to swine). Similarly, for a given branch we used the observed and reconstructed ancestral amino acid sequences of the parent and child-nodes to infer amino acid substitutions (or no substitution) for each of the variable sites on that branch. Lastly, branch lengths in the phylogenetic trees had been fitted to a molecular clock-model using TreeTime and thus represented the time in years elapsed during the given transmission between the two hosts.

To estimate substitution rates for the four transmission classes for each variable site, we fitted an F81-like substitution model to the data, i.e. a model where the substitution rate is assumed to depend on the equilibrium frequency of the target amino acid [46]. We furthermore assumed that the evolution followed a strict clock-model, i.e. that the rate for a given class and site was constant across the entire phylogeny. Specifically, we assumed that the rate matrix, Q, for a given site had the form: *Q_ij_* = µ_c_π_j_. Here, *Q_ij_* is the rate (in substitutions per site per year) from amino acid i to amino acid j, µ_c_ is a scaling-factor that is specific to the transmission class, c, and π_*j*_ is the equilibrium frequency of the target amino acid. The transition probability matrix P(t), giving the *probability* that amino acid i would change to amino acid j, on a branch of length t, could be computed from the rate-matrix Q using the expression *P*(*t*) = *e^Qt^*. For the F81-like model used here that equation has an analytical solution allowing simple computation of likelihoods: if *i* ≠ *j* (the amino acid has changed) then *P_ij_*(*t*) = π_*j*_(1 − *e*^−μ*t*^). If i=j (the amino acid has not changed) then *P_ij_*(*t*) = π_*j*_ + (1 − π_*j*_)*e*^−μ*t*^. In these expressions we knew, for a given site and branch, the branch-length, t, the original amino acid, i, and the derived amino acid, j. For the equilibrium frequencies of the target amino acids, π_j_ we used the empirically observed amino acid frequencies based on the alignment of the relevant viral protein. The site-specific scaling-factors µ_c_ for each of the four transmission classes were unknown and estimated using a Bayesian approach implemented using the probabilistic programming language Stan, version 2.32.6, via the CmdStanR interface [47–49]. The prior for all µ parameters were set to normal (0.002, 0.003). This is a weakly informative prior allowing some uncertainty about the value, while regularizing the fit to avoid unrealistic areas of parameter-space and is based on the known typical mutation rate for influenza RNA sequences (on the order of 2 x10^-3^ substitutions per site per year). Branches with a length > 15 years were discarded. We ran MCMC for 2000 iterations with 1000 iterations for warm-up, using four parallel chains. Convergence was checked by ensuring that R-hat values were close to 1, and that there were no divergent transitions. After obtaining MCMC samples we computed the posterior probabilities for each site that one transmission class had a higher rate than another transmission class (all 6 pairwise comparisons were performed).

### Analysis of positive selection across branches or sites

To further analyze the selective pressures across the phylogenies and protein-coding sequences of each viral segment, we utilized tools implemented in the HyPhy (Hypothesis testing using Phylogenies) software package v2.5.4 [50]. We employed the aBSREL (adaptive Branch-Site Random Effects Likelihood) model to examine branches for positive selection and identify lineages with episodic diversification [51] (example, ns1: hyphy absrel --alignment ns1.fasta --tree ns1_annotated_tree.nexus --output ns1_absrel.json). We employed the FUBAR (Fast Unbiased Basian Approximation) model to identify sites under pervasive positive or purifying selection across the entire phylogenies [52] (example, ns1: hyphy fubar --alignment ns1.fasta --tree ns1_annotated_tree.nexus --output ns1_fubar), and we applied the MEME (Mixed Effects Model of Evolution) model to detect episodic positive or diversifying selection at specific sites on a subset of branches [53] (example, ns1, human-to-swine branches: hyphy meme --alignment ns1.fasta --tree ns1_annotated_tree.nexus --branches human-swine --output ns1_meme_hu-sw.json). Positive selection was inferred at positions or branches where the non-synonymous (beta) to synonymous (alpha) substitution rate ratio (omega) significantly exceeded 1. We conducted four separate MEME analyses, each focusing on a different set of transmission branches (swine-swine, human-human, human-swine, or swine-human) as “foreground” to identify sites with distinct positive selection pressures compared to the rest of the branches (“background”). This approach aimed to highlight differentially selected sites among the four transmission groups, potentially reflecting variations in evolutionary pressures. For the MEME analysis, we excluded the PB1-F2 protein due to its overlapping reading frame with PB1 and high variability in protein length, which could bias the statistical modelling.

### Machine learning analysis for prediction of transmission types

As an indirect way of finding IAV protein features associated with transmission between or within human and swine hosts, we employed the XGBoost v.1.7.4 [54] algorithm implemented in Python through the Scikit-learn library [55]. XGBoost (Extreme Gradient Boosting) is an ensemble machine learning method built on decision trees and tree boosting that is widely acknowledged for its performance in various machine learning tasks including classification. The strategy was to train XGBoost to predict the transmission type from sequence data, and then identify sequence features important for the predictive performance, since these might play a role in species specificity and could function as biomarkers for zoonotic transmission.

Our datasets consisted of aligned observed and ancestral amino acid sequences obtained via anclib as previously described, and we thus trained separate models for each of the viral proteins. For each branch in the phylogenies, the sequence at the child-node was labeled as resulting from either human-to-human, swine-to-swine, human-to-swine, or swine-to-human transmission (for instance: if the host-state was human for the parent-node and swine for the child-node, then the sequence at the child-node was labeled as being the result of a human-to-swine transmission). Sequences from branches with a length >15 years were removed, as these would represent some of the more uncertain branches in the phylogeny, e.g. connecting different lineages. The sequences were represented using a one-hot encoding scheme, where each column of the matrix represented the presence or absence of a specific amino acid at each position within the alignment. An additional column for each position was used to indicate whether the position had undergone a substitution at the specific branch, giving a total of 21 columns per alignment position. Columns that were conserved across the alignment (uninformative) were filtered out from the dataset. Prior to model training, we tuned the hyperparameters of each XGBoost model using a Bayesian optimization approach and the resulting best-fit parameters were subsequently applied in a 20-fold cross-validation. For each iteration of this cross-validation, the one-hot encoded data was split into training (81%), validation (9%), and test (10%) sets, ensuring representation from all classes in each subset through stratification. The models were trained on the training subset while being evaluated on the validation subset. Given the imbalanced nature of our datasets with notably more observations in the swine-to-swine category and few in the swine-to-human category (**Figure 1d**), we applied weights to the training data, assigning greater significance to classes with fewer observations to enhance their influence during model training. We set an initial limit of 200 boosting rounds (iterations) but incorporated an early stopping criterion to stop the training if the logarithm to the loss function (logloss) for the validation set did not improve over 20 consecutive rounds to avoid overfitting. The performance of each model was assessed on the test set, and following the 20 independent training iterations, we aggregated the results to evaluate the robustness and generalizability of our models. This included assessing the average accuracy of the models and the mean sensitivity and specificity derived through confusion matrices. Additionally, we generated receiver operating characteristic (ROC) curves and calculated the corresponding mean area under the curve (AUC).

**Figure 1:**
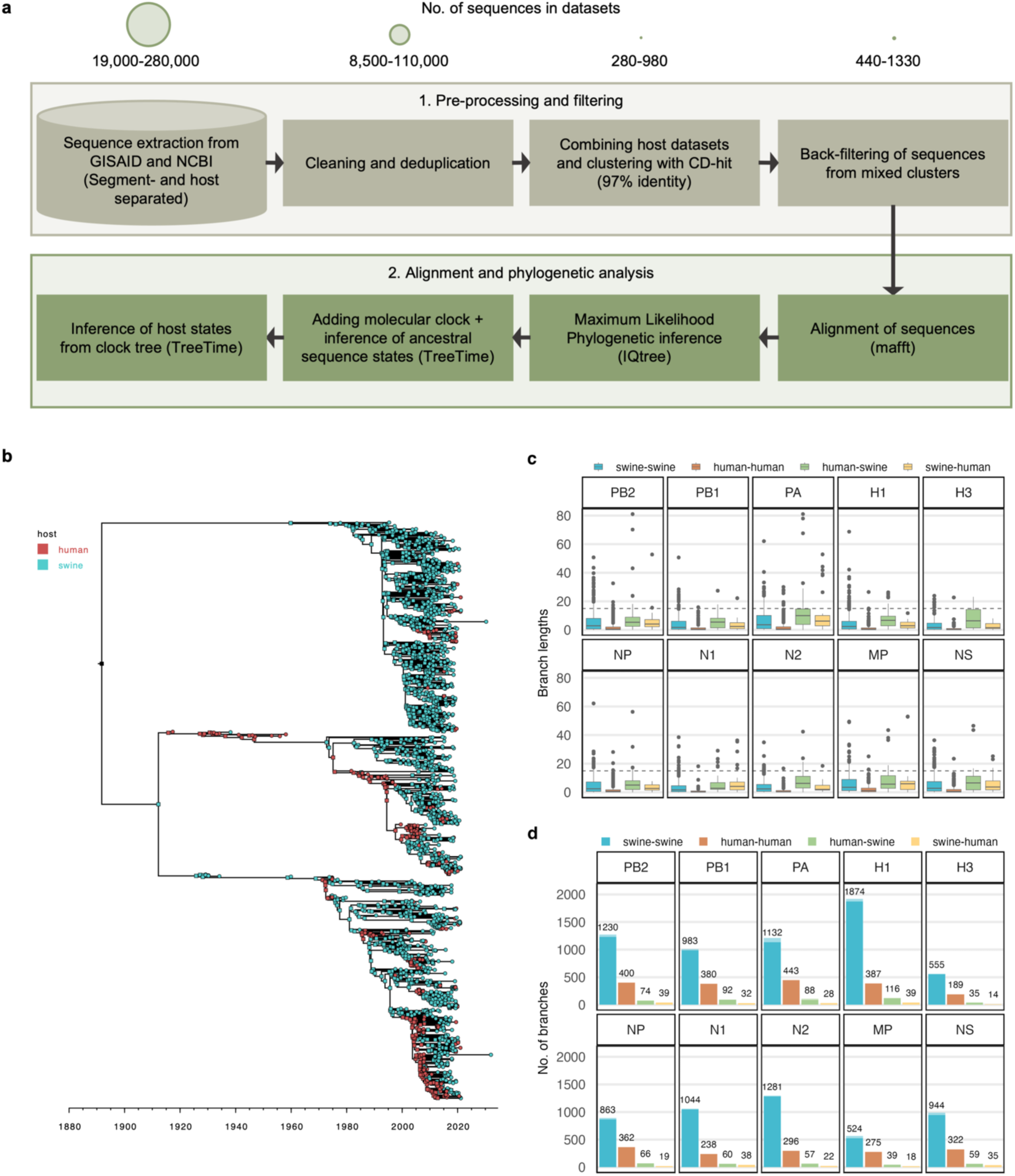
Construction of datasets of IAV transmissions between swine and human. **a.** Workflow for data preparation and phylogenetic analysis. Initial datasets encompassed all IAV sequences extracted from GISAID and NCBI Influenza databases (as of February 2023) for either human or swine IAVs and belonging to H1 or H3 and N1 or N2 subtypes. This resulted in datasets of ∼19,000 – 280,000 sequences depending on the host type and IAV segment. Deduplication and removal of poor-quality sequences filtered away approximately half of the data. Human and swine datasets were then combined for each segment and clustered based on sequence similarity using CD-HIT at a 97% similarity threshold. To avoid discarding examples of interspecies transmissions in the dataset, we retained closely related sequences isolated from different host-species (i.e., belonging to the same CD-HIT cluster). Specifically, up to five sequences from each host were added back to the dataset from such mixed clusters, giving final datasets of ∼ 440 - 1330 sequences for each segment. **b**. Time-scaled and host annotated phylogenetic tree for the H1 hemagglutinin gene. Nodes in the tree are colored based on their observed or inferred host origin as either human (red) or swine (cyan). See Supplementary **Figure S1a-j** for remaining segments. **c**. Boxplot showing the distribution of branch lengths for branches belonging to each of four transmission categories in each of the phylogenetic trees. Blue: swine-swine, orange: human-human, green: human-swine and yellow: swine-human. The dotted horizontal line indicates the cut-off at a branch-length of 15 years used for subsequent analyses. **d.** Bar plot showing the number of branches belonging to each transmission category for each segment. Semi-transparent bars indicate the total number of branches for each category while fully saturated bars show the remaining count after branches longer than 15 years have been excluded from the analysis. The number of branches used post-filtering is annotated on top of the bars. Blue: swine-swine, orange: human-human, green: human-swine and yellow: swine-human.

To uncover the significance of various features in the models which could correspond to specific amino acids or positional markers for each transmission type, we utilized Shapley values visualized using SHAP (SHapley Additive exPlanations) summary plots [56]. We represented both the overall mean Shapley values across all predictions and the direction-specific mean values (indicating positive or negative impact) for each of the four transmission categories. We also extracted the SHAP summary statistics and divided them into two categories; either representing the presence or absence of a specific amino acid at each position and summarized the top 5 most important features for each transmission type to be represented in bar-plots using R and ggplot2.

## Results

### Construction of a unique dataset of global transmission patterns between swine and human IAVs

Through the curation and clustering approach described in **Figure 1a** we have constructed datasets that encapsulate the spatial and temporal relationships and evolution of IAVs at the swine-human interface. Our phylogenetic analysis of IAV sequences from each viral segment reveals close and intertwined relationships of human and swine IAVs across all lineages (**Figure 1b**, **Figures S1A-J**), identifying both known and previously unreported zoonotic or reverse zoonotic transmissions. In our analysis of 10 separate datasets corresponding to the eight viral segments and the H1 or H3 and N1 or N2 subtypes, we have reconstructed ancestral host and sequence states while inferring the genetic changes along branches representing transmission events from human-to-human, swine-to-swine, human-to-swine, or swine-to-human (See **Table S1** and **Figure 1d**). Given the extensive variety of IAVs across time and genetic lineages captured in our phylogenetic trees, we found that the molecular clock hypothesis – assessed through root-to-tip regression of genetic distance against sampling dates – did not fit all segments uniformly. Notably, the polymerase segments PB2 and PA exhibited different evolutionary rates preceding the sampled sequences from various lineages, which impeded fitting a single regression line to the points. We therefore applied a covariation adjustment for the temporal linear regression of these segments. Despite the variation and the lower r-squared values indicating weak fits to the regression line (**Table S1**, **Figure S1A-J**), most sequences adhered to a consistent evolutionary rate across lineages and clades, supporting the use of overall root-to-tip distance and rate estimates. We hypothesized that the low regression line fit would predominantly impact branches connecting the different lineages, which we excluded from our analysis due to their disproportionately long branch lengths. By removing branches exceeding a length of 15 years, only a smaller fraction of the data was excluded, which was not expected to impact the genetic diversity in our further analysis (**Figure 1c-d**).

### Quantifying substitutions and frequencies across IAV transmissions in swine and human hosts

For each of the viral proteins (including the accessory proteins PB1-F2 and PA-X, which are functional in only some IAV strains and truncated in others) we examined general substitution trends that could be linked to specific transmission types and serve as potential host-specific markers. The average substitution counts across all branches in the phylogenies were fairly consistent (**Figure 2a**) with the highest substitution counts observed in the surface proteins (H1, H3, N1, N2), followed by the polymerase proteins (PB2, PB1, PA) and the nonstructural protein, NS1.

**Figure 2:**
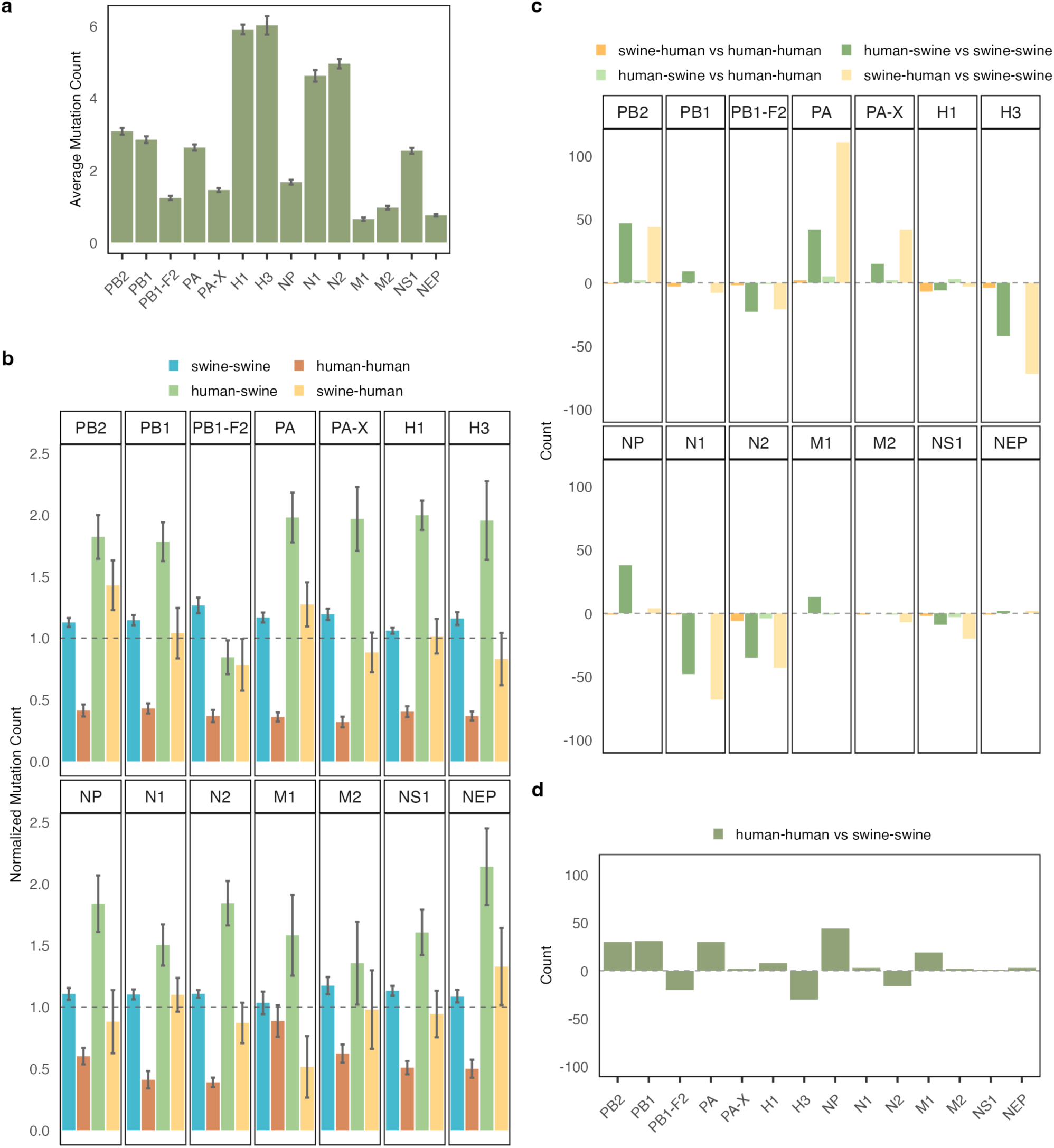
Substitution counts and rates across branches in the phylogenetic trees. **a**. Average count of substitutions observed per branch in the phylogenetic trees for each of the analyzed IAV proteins. Error bars show standard error of the mean. **b.** Average substitution count for branches of each transmission type normalized by the average substitution count across all branches in the phylogenetic trees. Error bars show standard error of the mean for each transmission type divided by the average substitution count across all branches. A dotted line at y=1 indicates the normalized substitution count at which the average substitution count in the transmission type equals the total average substitution count across all branches in the phylogenetic tree. **c**. Comparison of substitution rates across branch types. Bars represent the net amount of amino acid positions with a >95 % posterior probability of a higher substitution rate in one branch type compared to another (indicated by the colors), with the count from the opposite comparison subtracted. Positive y-values indicate more positions with higher substitution rates in the first branch type of the comparison; negative y-values indicate more positions with higher substitution rates in the second branch type. **d**. Count of positions in each protein with a higher substitution rate (>95 % posterior probability) in human-human branches compared to swine-swine branches.

Examining each transmission category individually, we observed a clear trend that branches representing human-to-human transmissions generally accumulated fewer substitutions, while human-to-swine branches displayed a higher relative substitution count in most proteins (**Figure 2b**). The human-to-swine branches were on average slightly longer for the PA, H1, H3, and N2 segments (**Figure 1c**). However, as this was not the case for all human-to-swine branches, this cannot fully account for the increased substitution frequencies observed, suggesting that at least some of the substitutions reflect adaptive evolutionary pressures following the host switch from human to swine. An additional analysis of the NS1 protein and accessory proteins, PB1-F2 and PA-X, (**Supplementary Figure S2a-c**) revealed high variability in the expressed lengths of the proteins within the human and swine IAV strains in our datasets. However, the lengths appeared to be lineage- or clade-specific rather than host-specific. This was particularly evident for the PA-X and NS1 proteins, which displayed consistent protein lengths within large clades. The PB1-F2 protein showed more variability in length; however, still with no apparent correlation to the host type.

To further explore differences in substitutions following different types of transmissions, we analyzed the substitution rates per site in each protein for the four branch types using a Bayesian framework to identify positions with significantly different substitution rates. This analysis revealed only two positions in the PA protein with a posterior probability >95% of a higher substitution rate in swine-to-human branches compared to human-to-human branches (**Figure 2c**). However, in other proteins, more positions had a higher substitution rate in human-to-swine branches (NP) or both human-to-swine and swine-to-human branches (PB2, PA, and PA-X) compared to swine-to-swine branches. Conversely, the H3, N1, N2, and to a lesser degree PB1-F2 and NS1 proteins, showed more positions with a higher substitution rate in swine-to-swine branches compared to both swine-to-human and human-to-swine branches (**Figure 2c**). This pattern aligned with the counts of positions with higher substitution rates in human-to-human versus swine-to-swine branches in most proteins (PB2, PB1-F2, PA, H3, NP, N2) (**Figure 2d**), suggesting an overall difference of substitution rates in these proteins for the swine and human host.

Hypothesizing that a higher substitution rate for zoonotic transmissions compared to intraspecies transmissions in the recipient host could be an indication of host adaptation, positions with a significantly higher substitution rate in swine-to-human versus human-to-human, and in human-to-swine versus swine-to-swine transmissions were examined in more detail. For the PA protein, there were two positions where the substitution rates for swine-to-human transmissions were higher than for human-to-human transmissions with high certainty (posterior probability >95%). These sites were located in close proximity to each other in the end of the C-terminal domain of the PA protein. For the human-to-swine transmissions, positions with significantly higher substitution rates were detected across different proteins and were especially concentrated in the Mid and Cap-binding domains of PB2, the end of the endonuclease domain of PA and PA-X, the HA2 domain of H1, and the effector domain of NS1 (**Figure 5**).

### Detection of positive selection across branches in the IAV phylogenies

We used the aBSREL method [51] to detect lineage-specific episodic diversifying selection across our phylogenies. Briefly, this approach compared the rates of synonymous (silent) and nonsynonymous (non-silent) mutations to identify codons evolving under positive selection on specific branches of a phylogeny. Our analysis identified positive selection acting on various branches for all proteins, except for the M2 protein, with the HA, NA, PB1, and PA phylogenies displaying most positively selected branches (**Table S2**).

We were especially interested in finding signs of positive selection driven by the virus adapting to a new host species after a zoonotic or reverse zoonotic jump, which could be signified by a higher incidence of positive selection on those specific branches of the phylogenetic trees. Thus, we computed the fold over-representation of branches belonging to each of the four transmission types (swine-to-swine, human-to-human, human-to-swine, or swine-to-human) among the positively selected branches for each viral protein (**Figure 3a**). While most phylogenies showed evidence for positive selection on only a smaller fraction of the total branches, we did find tendencies of such an over-representation of positively selected branches related to zoonotic and reverse-zoonotic jumps (**Figure 3a**). Swine-to-human transmissions were over-represented among positively selected branches for the PA and N1 proteins and human-to-swine transmissions were over-represented for the PB1, PA-X, H1, NP, N1, and NEP proteins. In these cases, the selected sites could potentially be related to adaptation to the new host-species after the cross-species jump. We also found signs of positive selection driven by intra-species processes. Specifically, swine-to-swine transmissions were slightly over-represented for the PB1-F2, H3, and M1 proteins (and there was only positive selection on swine-to-swine branches for the H3 and M1 proteins), while human-to-human branches were over-represented for the PA, H1, N1, N2, and NS1 proteins. In these latter cases the selection could be driven by for instance immune-related processes (which might of course also play a role for the cross-species jumps).

**Figure 3:**
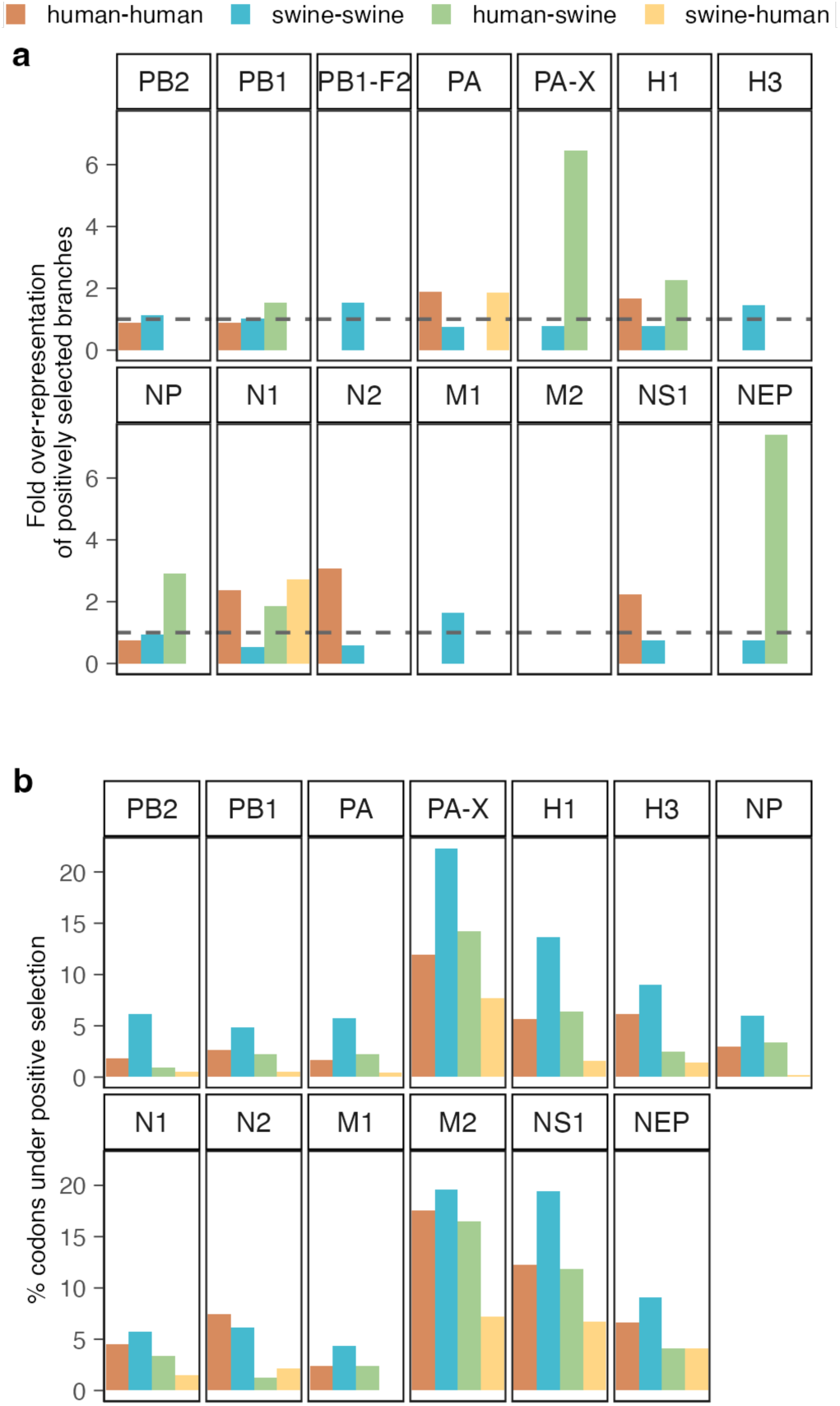
Positive selection pressures across branches or sites of each transmission type**. a.** Positive selection on branches corresponding to the four types of transmissions: human-human, swine-swine human-swine, and swine-human. Each bar represents the fold over-representation of a specific transmission type among positively selected branches compared to its overall fraction in the phylogeny. Values above 1 (indicated by the dashed line) suggest that branches of a given transmission type are over-represented among those experiencing positive selection. **b.** Proportion of codons in each protein with evidence of positive selection following the four different types of transmission. The proportions are calculated as the number of codons identified under positive selection in the MEME analysis for the respective transmission type divided by the amino acid length of the protein and shown as a percentage. The PB1-F2 protein is not full-length and functional in all lineages and the position-wise selection pressure calculated by MEME is thus not included in our analysis.

Given that this analysis is confined to individual branches and the host state at each end, it is possible that some selection pressures arising from interspecies transmissions might be overlooked. For instance, there are some positively selected branches in both the NS1 and PA phylogenies, that are classified as intraspecies (human-to-human or swine-to-swine) but where the grand-parental node in fact has a different host, forming for instance human-to-swine-to-swine chains of transmission (**Table S2**). In these cases, it is possible that adaptation to the new host is only seen after an extra transmission step within the new species.

Therefore, it appears that many internal proteins are under higher selective pressure following interspecies transmissions, indicative of adaptations to new host environments, while the surface proteins also encounter significant selective pressure during transmissions within the human or swine populations, presumably due to intense immune pressures.

### Detection of amino acid sites under positive selection following inter- or intraspecies transmissions of IAV

While the analysis of site-level selection pressures showed mostly purifying selection when constraining to a consistent pressure across the entire phylogeny (**Figure S3**), we used the MEME model [53] to test if the selective pressure on individual codon sites varied between branches representing different transmission types (**Table S3**). For all transmission types, there was a generally higher proportion of codons under positive selection in the M2, PA-X, NS1, H1, H3, and NEP proteins, which was also evident from the pervasive analysis (**Figure 3b** and **Figure S3**, **Figure S4c**). However, the swine-to-swine transmission category exhibited a higher proportion of positively selected positions across all proteins, while the swine-to-human category had the fewest (except for in the N2 and NEP proteins), possibly related to the uneven distribution of branches across the categories. Interestingly, for most proteins (PB1, PA, PA-X, H1, NP, M1, M2, and NS1) the analysis revealed similar proportions of positively selected sites for the human-to-human and human-to-swine categories of branches (**Figure 3b**), despite fewer branches in the human-to-swine category (**Figure 1d**).

Observing the positively selected sites across each protein, we noted variability in the distribution and recurrence patterns between transmission types. While there were few or no common positively selected sites in the polymerase segments, almost half of the positions were shared between transmission types for the M2 protein (**Table S3** and **Figure S4a-b**). Notably, shared positively selected positions in many proteins (PA, H1, NP, N2, M2, NS1, and NEP) were predominantly between swine-to-swine and human-to-swine transmissions, suggesting common adaptations in the swine host.

Distributions of positively selected sites, both pervasive (**Figure S3b**) and episodic (**Figure 5**), across all branch types revealed significant positive selection in regions of proteins encoded by overlapping and shifted reading frames in relation to other essential proteins. Specifically, high levels of positive selection were observed in the last ∼70 bases of PA-X and NS1, as well as in the middle of the ecto-domain of M2. For these proteins, most identified positively selected positions for all transmission categories overlapped with sites identified under perversive positive selection by the FUBAR analysis (**Figure S4c**). In other proteins, a larger proportion of the sites with evidence of positive selection in the episodic analysis, had evidence of purifying selection in the pervasive analysis, especially for the polymerase proteins (PB2, PB1 and PA). There were more unique episodic positively selected positions with neither strong evidence of pervasive purifying or positive selection for the surface and envelope proteins (H1, H3, N1, N2 and M1) and a general tendency of more unique positively selected positions found by the episodic analysis on the interspecies branches (human-to-swine and swine-to-human), suggesting a general higher selective pressure following the interspecies transmissions (**Figure S4c**).

The positively selected sites from the episodic analysis were mostly spread throughout the lengths of each segment (**Figure 5**), however some trends worth noting were a seemingly higher positive selection pressure on the mid and cap-binding domains of PB2 for the interspecies branches, a high concentration of positively selected sites in the receptor binding domain of H1 and the rest of the HA1 head domains of both H1 and H3 proteins, and an interesting concentration of positively selected sites in the middle of the HA2 domain of the H3 protein, especially for the human-to-human branches. Swine-to-human branches for the N2 protein showed notably higher rates of positively selected sites in the transmembrane and stalk domains, while most positively selected positions on the human-to-human and swine-to-swine branches were in the head domain. Specifically, the human-to-human branches had many positions under positive selection in the last part of the head domains of both N1 and N2 proteins. Positively selected positions in the M2 protein appeared to be concentrated mostly in the ectodomain and the end of the cytoplasmic tail, and, for the NS1 protein, a very high selective pressure was detected on positions in the end of the effector and C-terminal domains for all transmission types.

### Detection of species-specific signatures based on XGBoost prediction of transmission types from child-node sequences

We explored the use of the machine learning method XGBoost [54] as an indirect way of detecting residues important for species-specificity. Specifically, we trained models to predict the type of transmission (swine-to-swine, human-to-human, human-to-swine, or swine-to-human) for each branch in a phylogeny, based on (1) the amino acid sequence present at the child-node, and (2) an indication of whether each amino acid residue was different from the parental sequence at this position. Separate models were trained for each of the 14 viral proteins investigated here. The idea was to identify the sequence features that were most important for making such predictions, under the assumption that these features were likely to play a role in species-specificity. Despite the relatively small datasets, the models demonstrated a capacity to accurately predict the ancestral transmission pathways for a substantial portion of the test sequences across all proteins as evaluated by accuracy measures, ROC curves, and AUC metrics (**Table 1 and Figure S5a-n)**. For intraspecies transmissions, the models achieved high predictive accuracy with most AUC values above 0.9 (**Table 1** and **FigureS5k**) (The exception was the M1 protein which however still had AUC values of 0.88 for human-to-human and 0.85 for swine-to-swine transmissions). The models showed reduced accuracy for interspecies transmission predictions, particularly for swine-to-human transmissions, which had a median AUC of 0.56, close to the threshold of random chance (0.5). Part of the reason for this could probably be the low number of training examples belonging to this class. Performance was better for the human-to-swine category, with a median AUC of 0.80 and one AUC value at 0.9 (for predictions based on the H1 protein). Interestingly, the confusion matrices revealed a tendency for the models to misclassify interspecies transmissions as intraspecies events of the originating host. Specifically, swine-to-human transmissions were frequently predicted as swine-to-swine, while human-to-swine transmissions were classified as human-to-human, a trend consistent across all proteins except H1, H3, and N2 (**Table 1**).

**Table 1:**
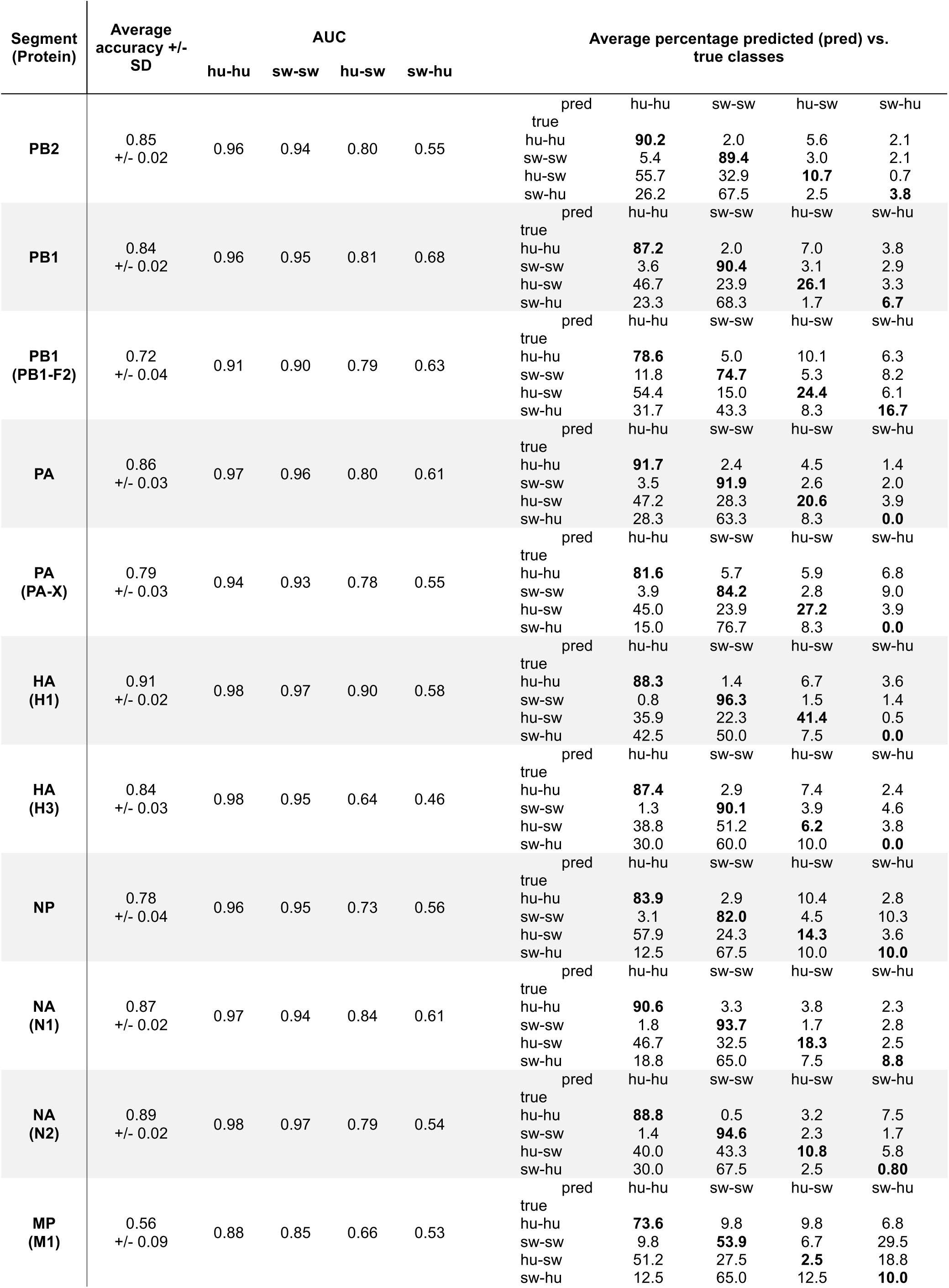
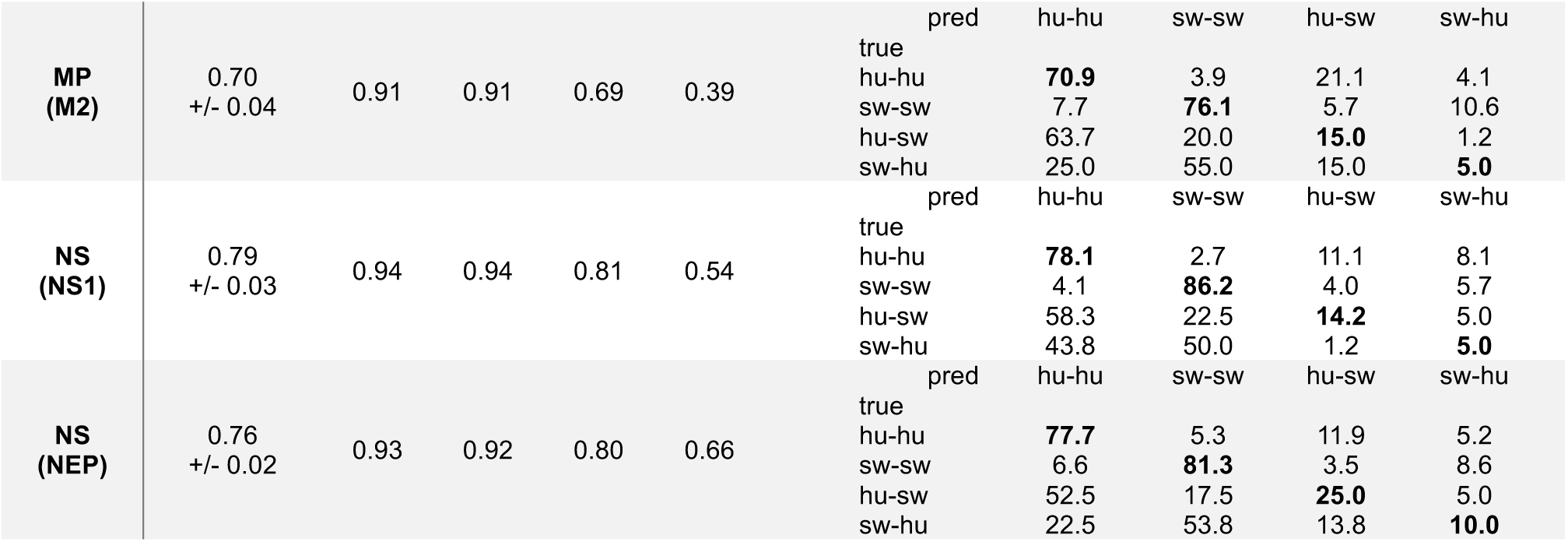
Summary of XGBoost model performances for each IAV protein. The models were trained to predict the transmission type (hu-hu, sw-sw, hu-sw, or sw-hu) based on (1) the amino acid sequence present at the child-node, and (2) an indication of whether each residue had changed from the parental sequence. All measures are averages over the 20 independent runs of model training performed for each protein and the average accuracy is specified with the margins of deviation (+/- SD). AUC values are the average area under the curves determined by the one-vs-rest ROC curves in **Figure S5**. Confusion matrices are specified for each protein with the average percentage of predicted (pred) classes vs. the true classes for all test samples. The percentage of samples where the predicted class was the true class is highlighted in bold for each of the transmission types. hu = human, sw = swine.

### Utilizing model predictions to identify host specific markers

We used SHAP summary statistics to identify the features that were most important for the predictive ability of each of the classification models. SHAP values quantify the importance of each feature by ranking their contributions to the model’s output, and thus, the most important features could be used to infer positions and amino acids with possible importance for species-specificity and zoonotic potential. The features we analyzed as predictors were: (1) presence or absence of specific amino acids at each variable position in the child-node of a branch, and (2) whether the amino acid at that position had changed compared to its corresponding residue in the parental sequence. **Figure 4** summarizes the top 5 most important features for each transmission category for each protein, showing the mean SHAP values (height of bars) while specifying if presence or absence of the specific amino acid at the position caused the impact on the models (solid or dashed outline). **Supplementary** Figure 6 shows more details about the top 20 features for each protein and transmission category, while **Supplementary Table 4** provides details about the distribution of amino acids at the positions mentioned in **Figure 4**.

**Figure 4:**
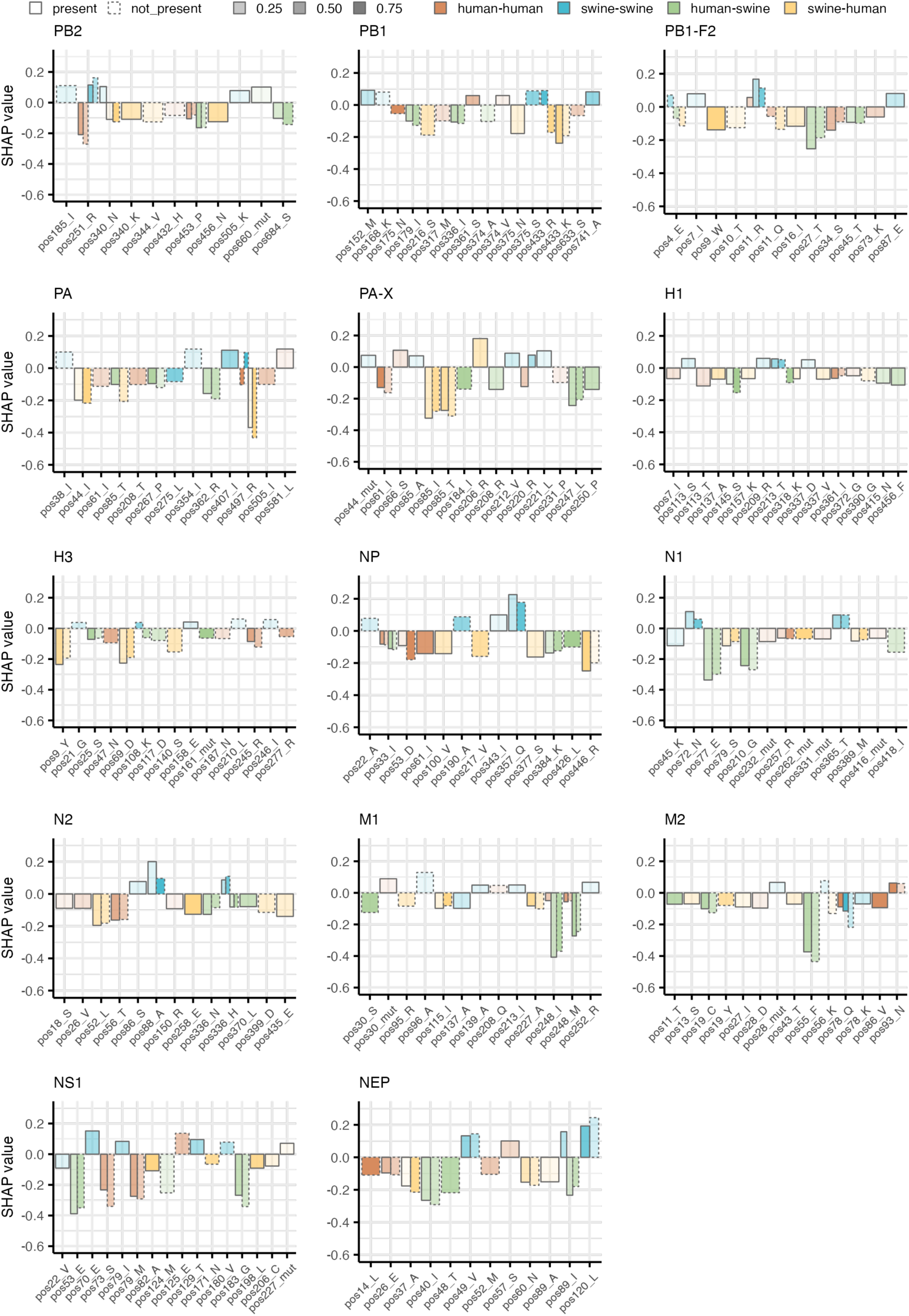
Most important amino acids and positions used for predicting IAV transmission categories. Each bar plot shows the mean SHAP values of the top 5 features used in the model predictions of each transmission category for each protein. The features are shown on the x-axis of each plot and represents the position in the protein and the specific amino acid or presence of a mutation (mut) used by the model to make predictions. The y-axis shows the mean SHAP values for the features, with positive values indicating a positive impact on model predictions (increasing the probability of this category) and negative values indicating a negative impact on model predictions. A SHAP value of 0 means that the feature has no impact on the model predictions. The bars are colored based on the transmission category (orange: human-human, blue: swine-swine, green: human-swine and yellow: swine-human) and outlined with either a solid line for presence of the feature or a dashed line for absence of the feature. The transparency of the bars shows the fraction of test samples in which the relevant feature is either present or not, with fewer samples denoted by higher transparency of the bars.

The most influential features (highest absolute SHAP values) were observed for the NS1, NEP, NP, M1, and M2 proteins, while most individual features of especially the HA proteins (H1 and H3) had relatively low impact on predictive capability (**Figure 4**). A closer examination of selected features provided further insights into the roles played by specific amino acid residues in determining species specificity. One example is PB2 position 251, where mostly the absence of arginine (R) contributed positively to predicting swine-to-swine transmissions but negatively to human-to-human transmissions. Consistent with this, we observed a clear preference for R at this position in human-derived viral sequences (93.9% R; **Supplementary Table 4**), while swine-derived sequences appeared to also tolerate lysine (K) well (61.5% R, 38% K). Focusing on the branches corresponding to swine-to-human transmissions, it could be seen that after the transmission (i.e. in the recipient human host) there was an intermediate distribution, with 77% R and 23% K, i.e. a much higher frequency of R than in the original swine host. This could indicate that IAVs that jump from swine to human hosts preferentially are the swine viruses that already have an R at this position, or that the viruses subsequently undergoing mutation to obtaining an R are selected for in the human host. Other examples include the PB1 position 375, where presence of asparagine (N) contributed negatively to predictions of swine-to-human transmissions and absence of serine (S) contributed positively to swine-to-swine transmissions (human: 79.8% S, 10.9% N; swine: 50.8% S, 17.9% N; swine-human: 75% S, 3.1% N; human-swine: 87% S, 9.8% N); PA position 497, where absence of R mostly contributed positively to predictions of swine-to-swine transmissions but negatively to human-to-human and swine-to-human transmissions (human: 89.3% K, 10.7% R; swine: 91.5% K, 8.2% R; swine-human: 82.1% K, 17.9% R; human-swine: 87.5% K, 12.5% R); or NEP position 89, where presence of isoleucine (I) contributed positively to predictions of swine-to-swine transmissions but mostly negatively to human-to-swine transmissions, and alanine (A) contributed negatively to predicting swine-to-human transmissions (human: 36% A, 17.2% I; swine: 23.8% A, 37.3% I; swine-human: 20% A, 28.6% I; human-swine: 45.8% A, 1.7% I) (**Figure 4**, **Supplementary Table 4)**.

### Integration of Analyses to Identify Potential Host-Specific Genetic Markers

The results from our three analytical approaches to identify site-specific markers of host-specificity are consolidated and compared in **Figure 5** and **Table 2**. **Figure 5** provides a schematic representation of IAV proteins, highlighting positions identified by each method. Areas or specific amino acid positions identified by multiple analyses (marked by vertical lines) may serve as potential markers of host-specificity.

**Figure 5:**
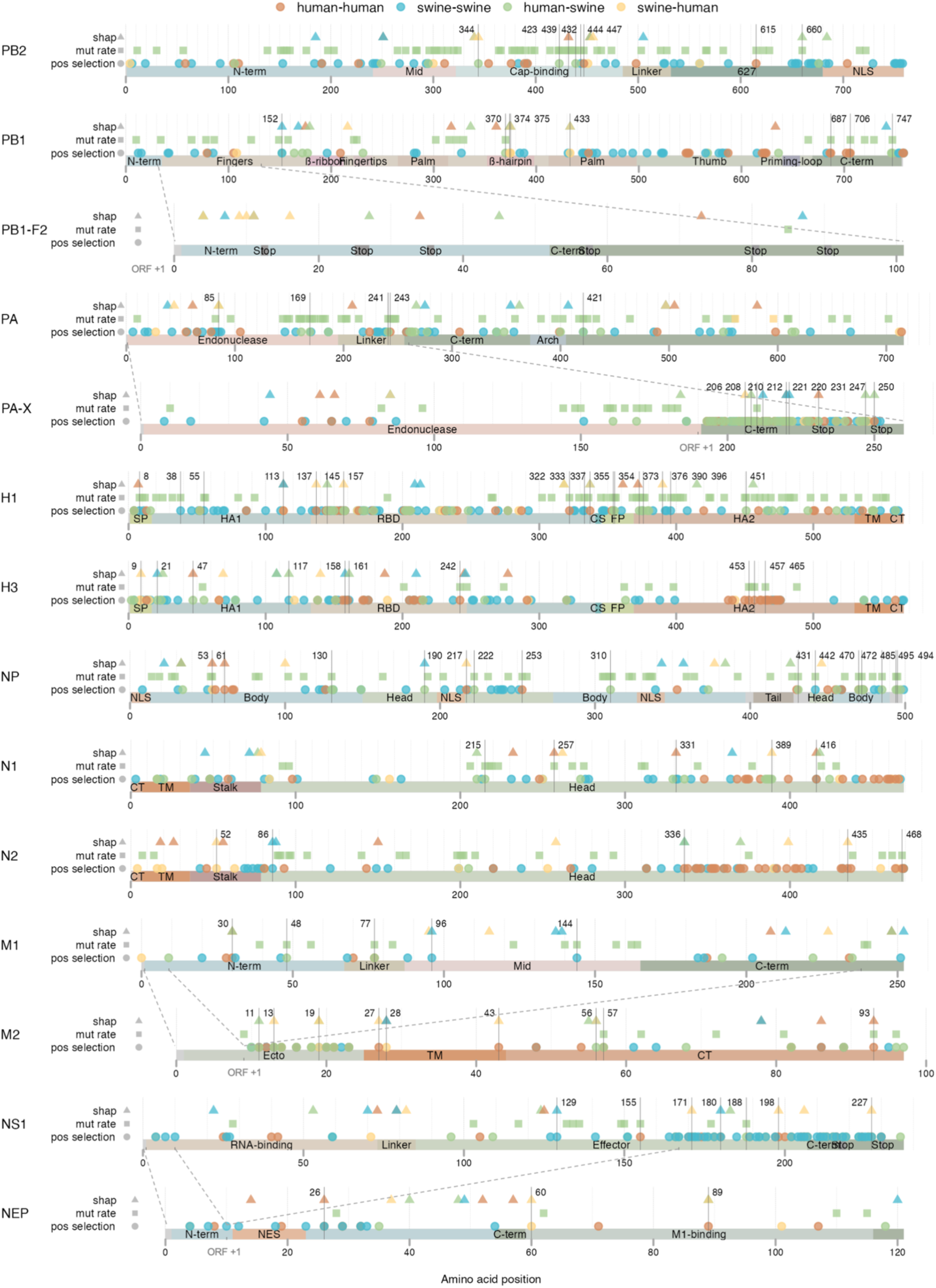
Summary of positions identified as potentially contributing to species-specificity using three different approaches. IAV proteins are represented as bars, with important protein domains highlighted by text and color. Positions identified by the three methods are indicated on three lines above each protein using different shapes: XGBoost/SHAP analysis (triangles), Bayesian mutation rate analysis (squares), and positive selection analysis (circles). Shape colors correspond to predicted transmission categories (orange: human-human, blue: swine-swine, green: human-swine and yellow: swine-human). Positions identified in at least two of the analyses are highlighted with a vertical line and the position number. Note that the lengths of the proteins in this schematic are not proportional to their actual sizes, so proteins of different lengths may appear similar in size. Overlapping reading frames between proteins from the same segment are marked with dotted lines and shifted reading frames are indicated by ORF +1. Protein domain abbreviations: N-term, N-terminal; NLS, nuclear localization signal; C-term, C-terminal; SP, signal peptide; RBD, receptor binding domain; CS, cleavage site; FP: fusion peptide; TM, transmembrane; CT, cytoplasmic tail; NES, nuclear export signal; Stop, common premature stop codons.

**Table 2:**
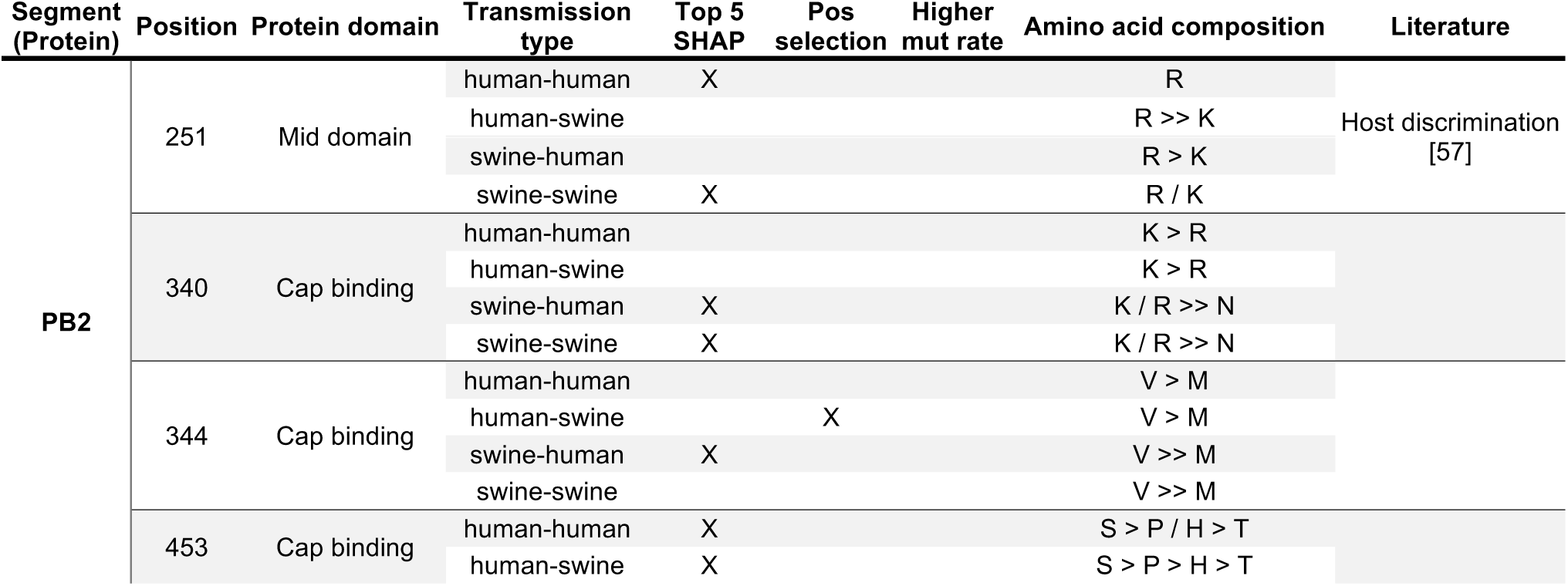
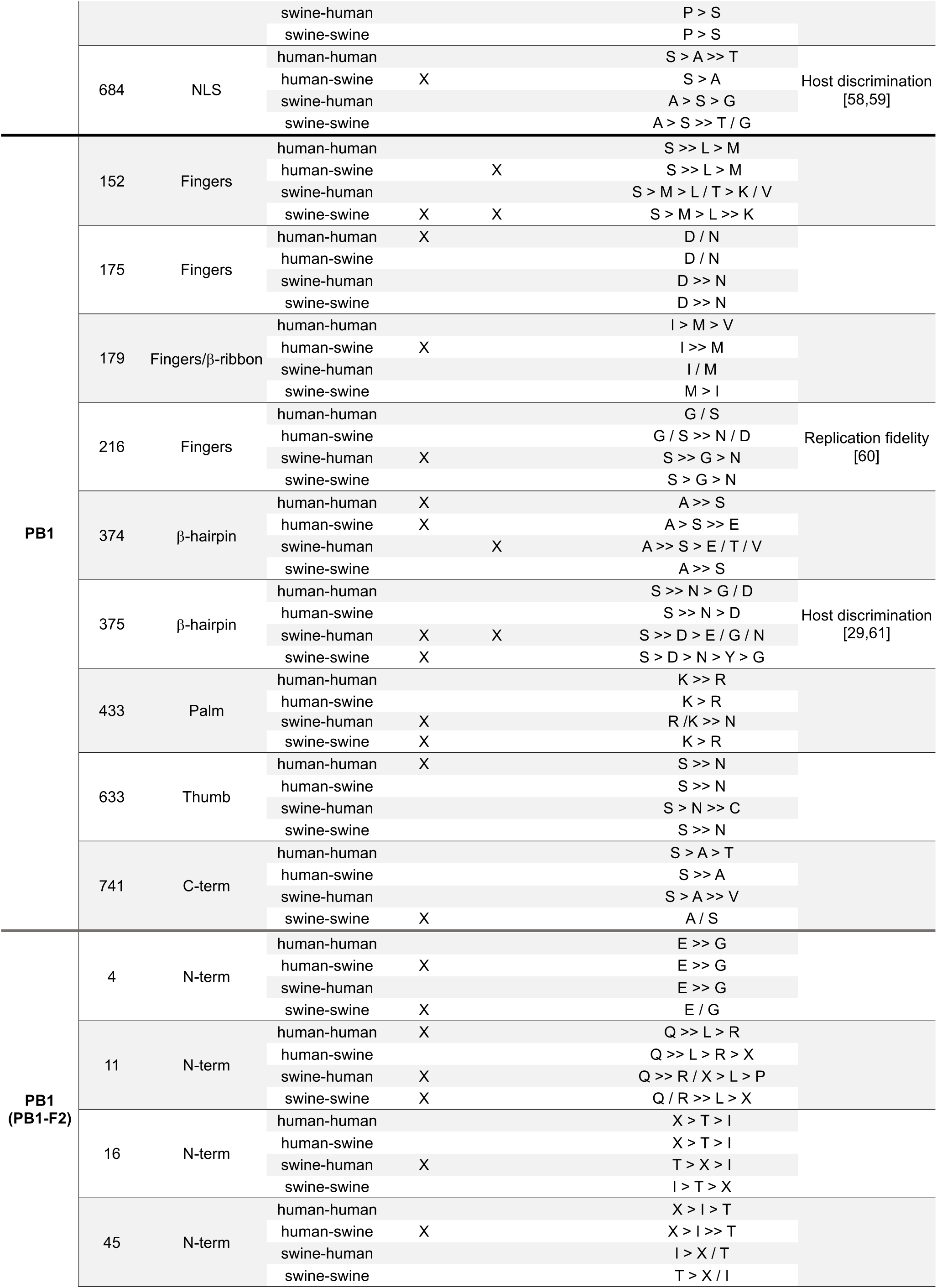
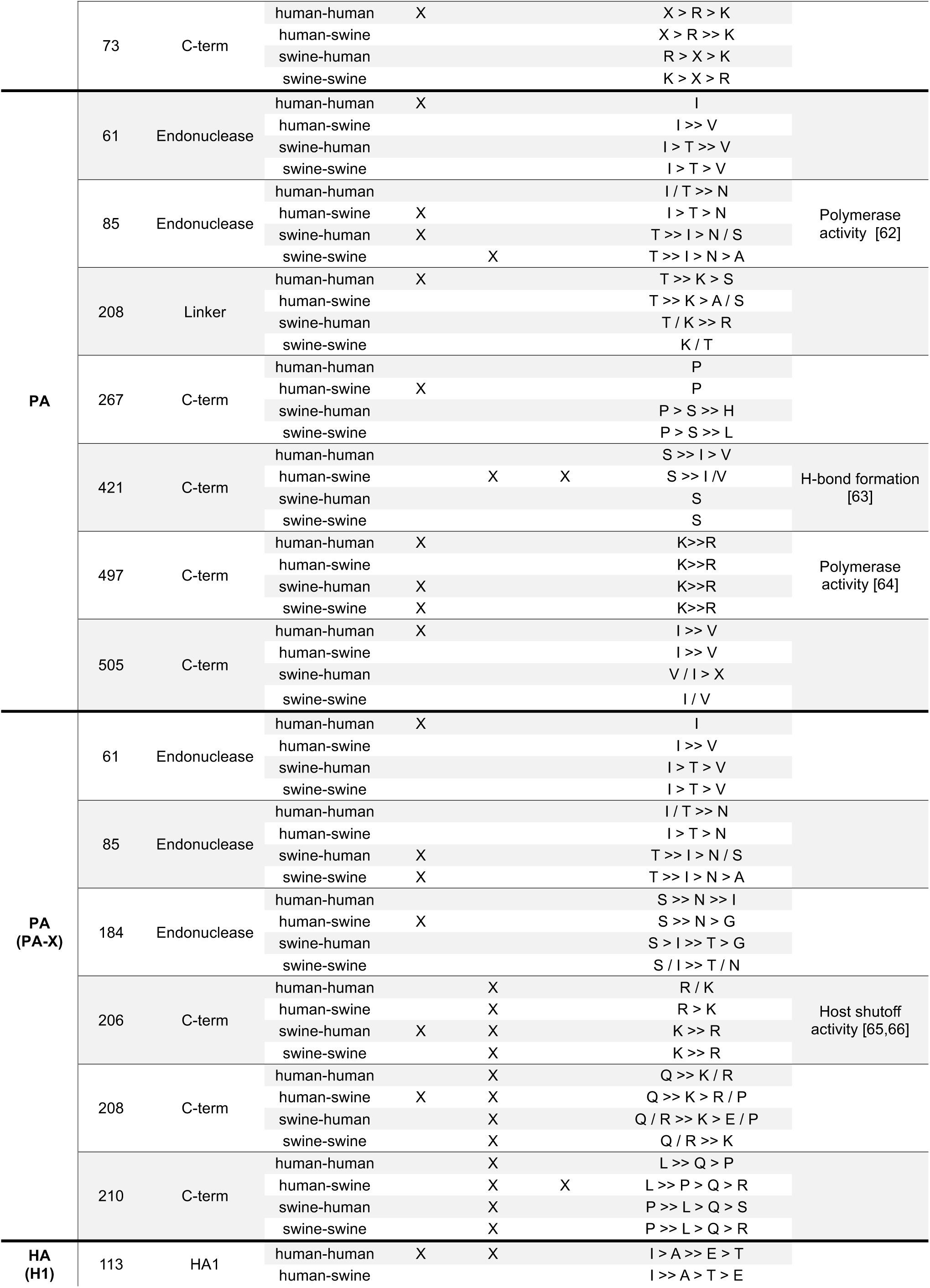
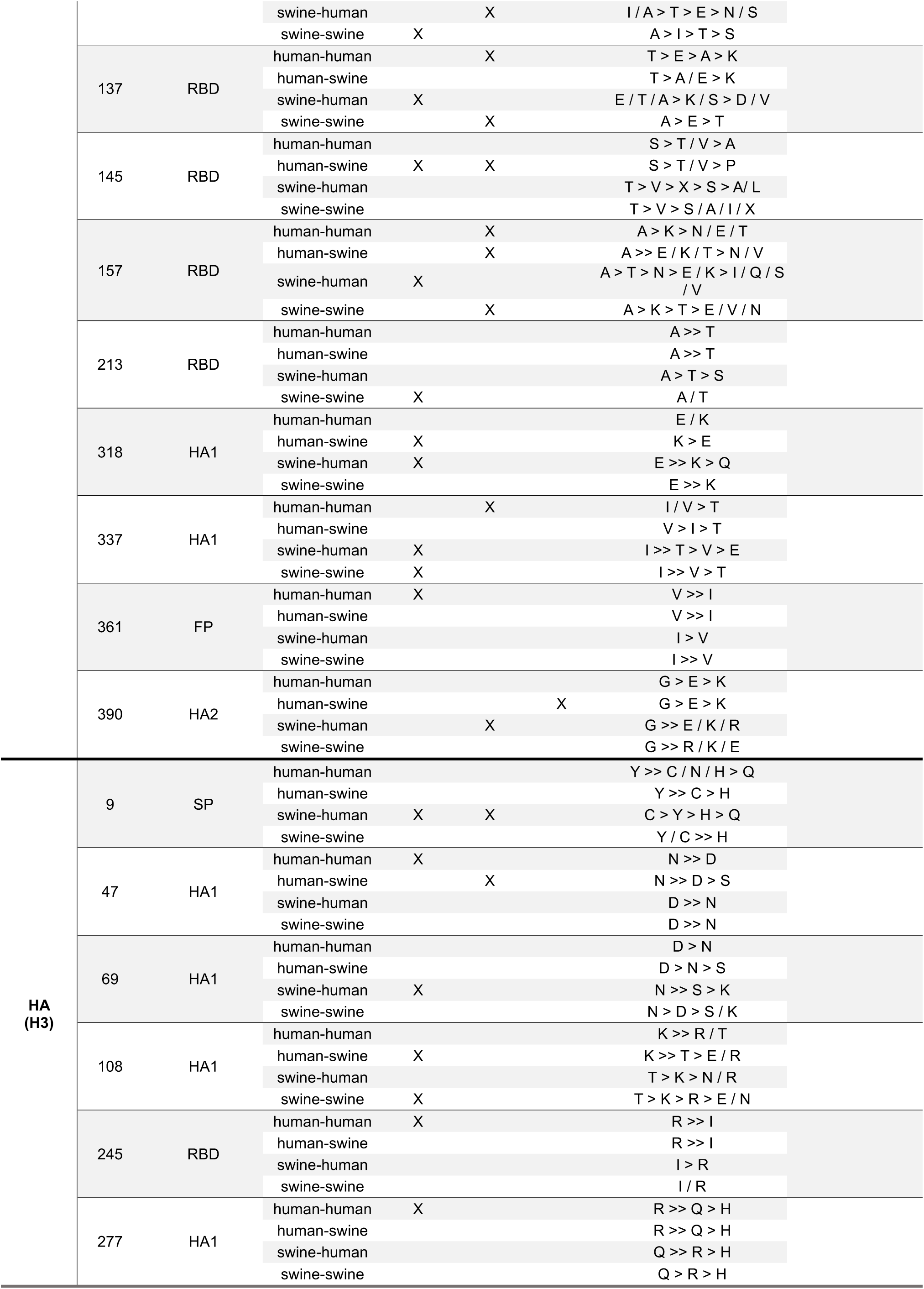
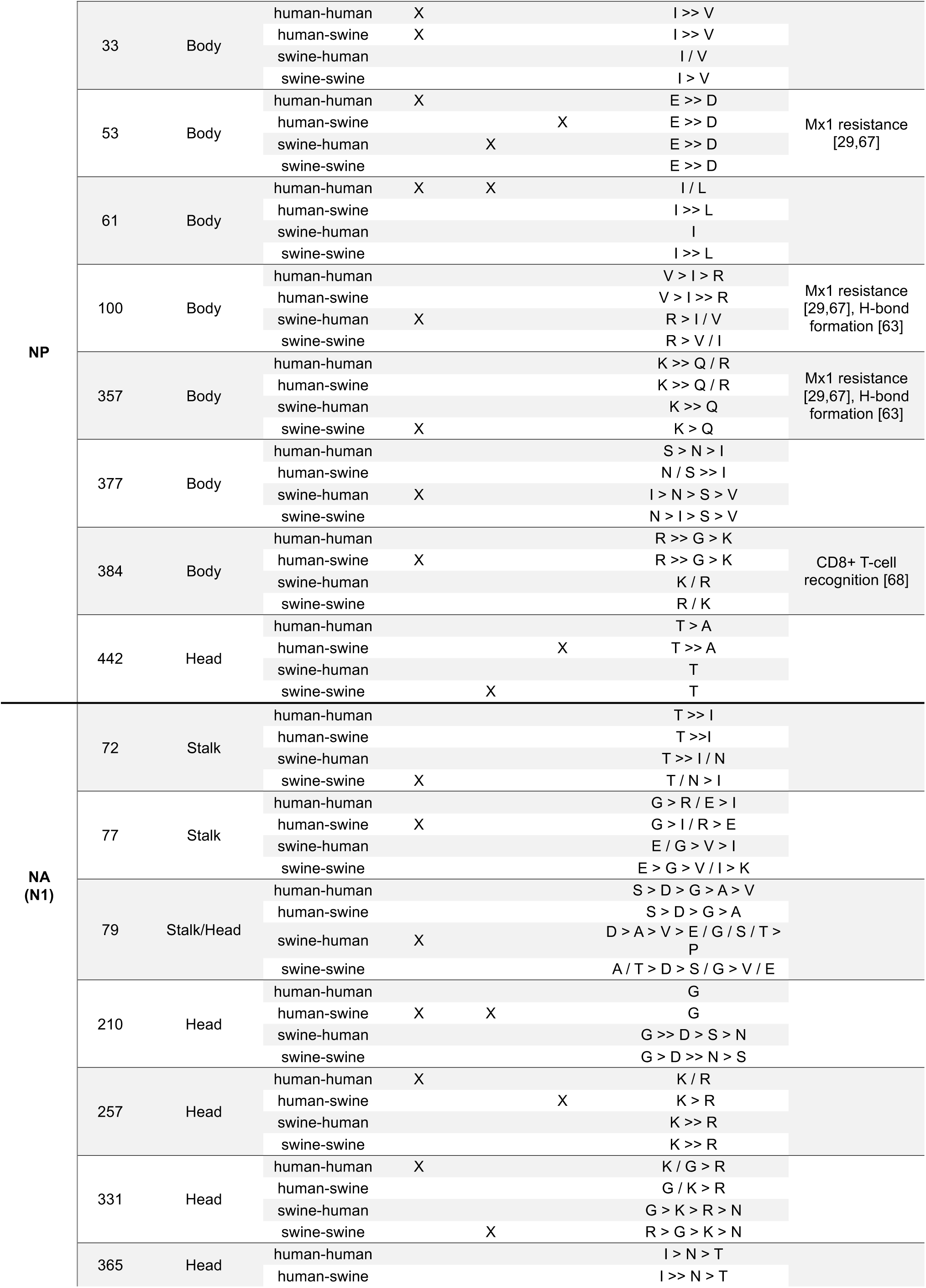
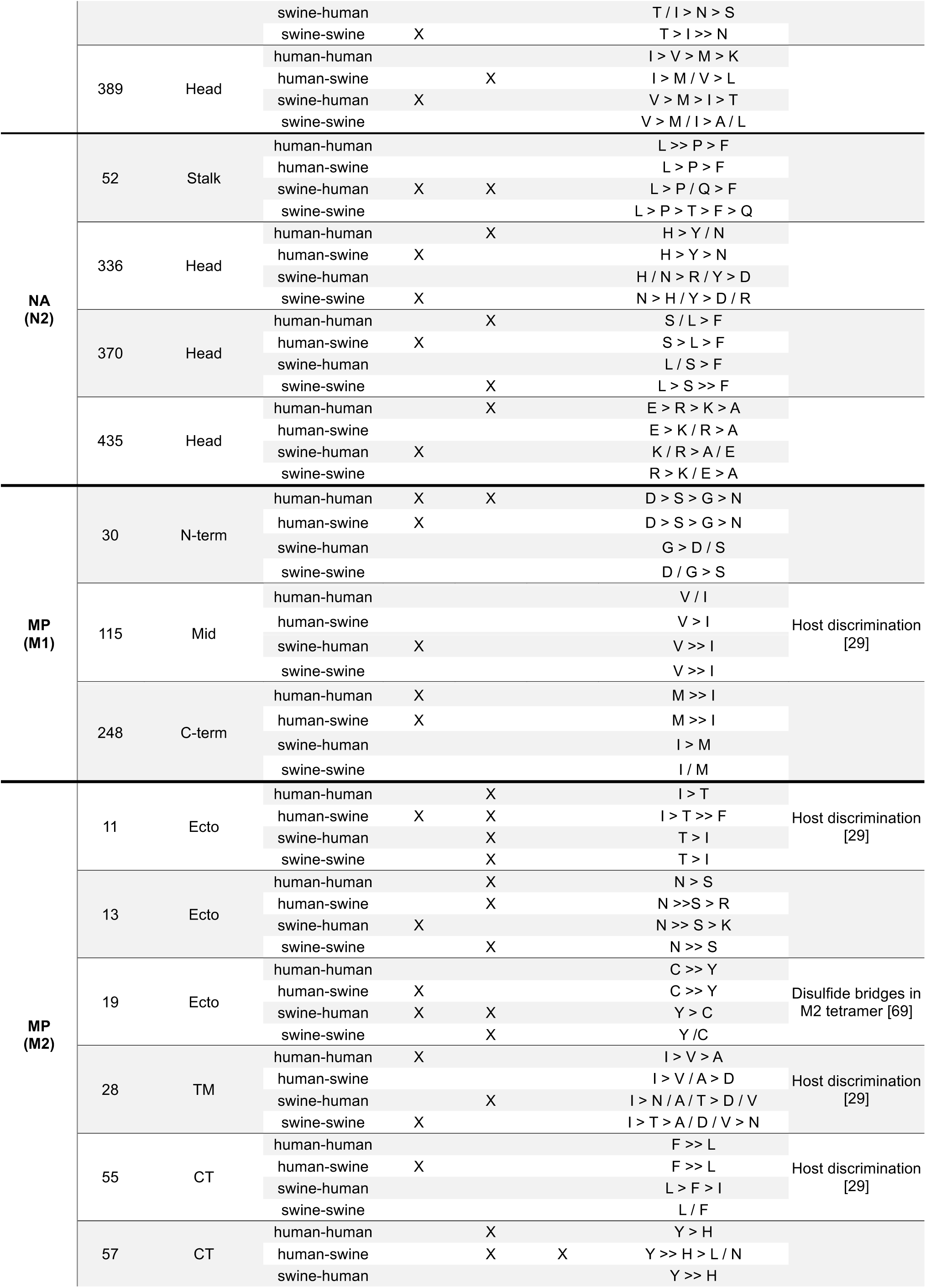
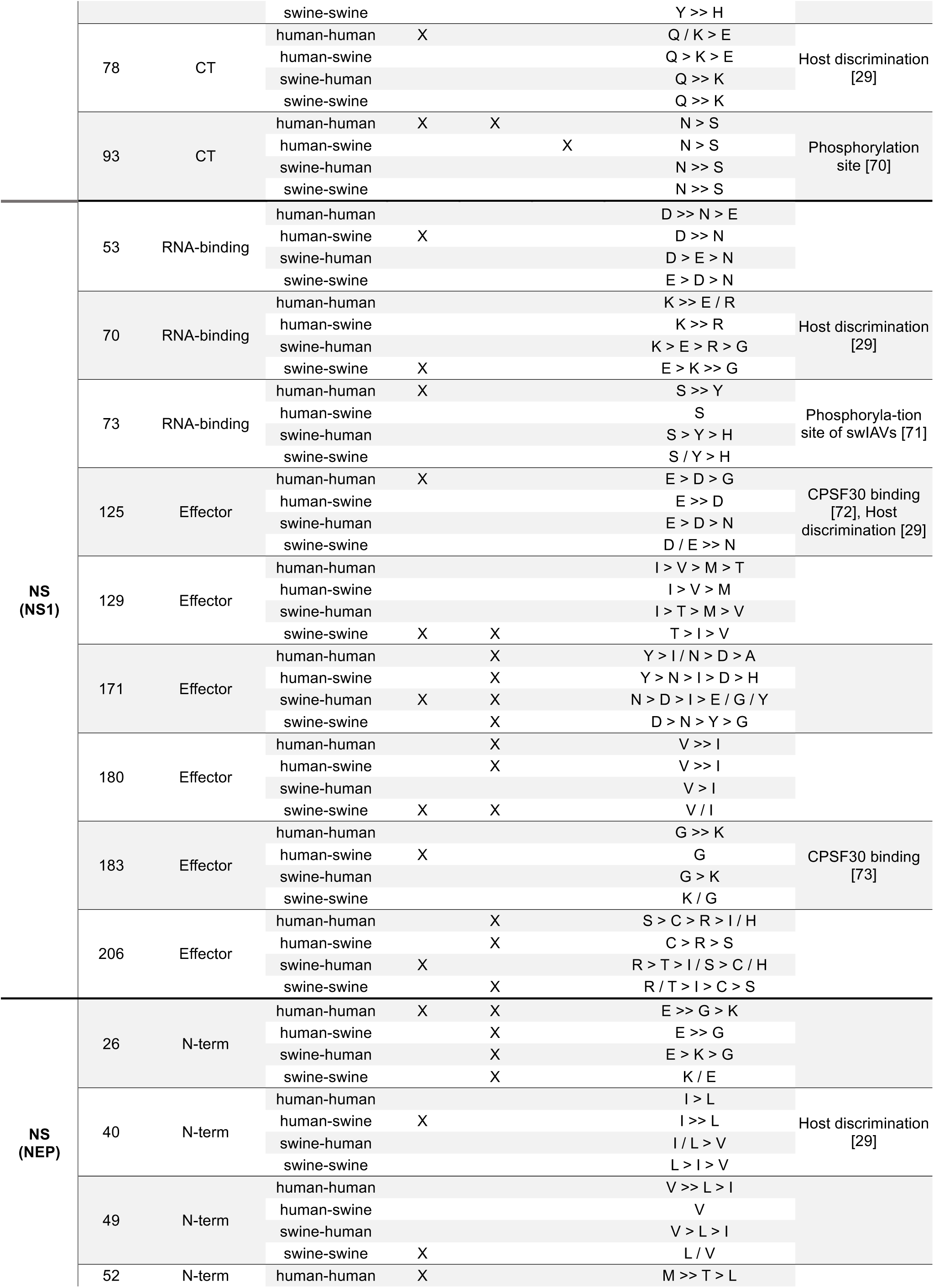
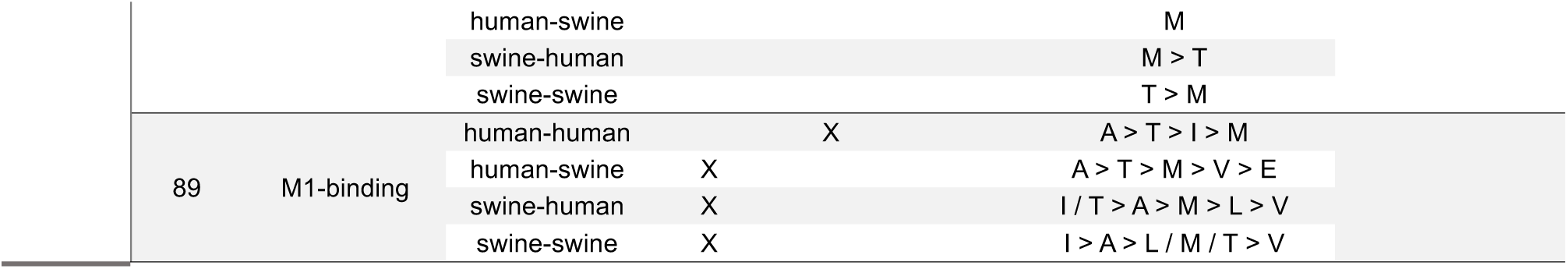
Summary of positions identified as potentially contributing to species-specificity using three different approaches. For each position the columns representing each of the analyses (Top 5 SHAP features, Positive (pos) selection, Higher mutation (mut) rate) indicate whether the position was identified for one or more transmission types in the specific analysis. The Amino acid composition column summarizes the relative distribution of amino acids present at that position for the child nodes of tree branches of each transmission type (>>: much higher proportion, >: higher proportion, /: approximately similar proportions). The Literature column provides references and brief descriptions of prior studies demonstrating a functional role for the residues.

**Table 2** offers further details on the most significant identified sites, including a summary of the amino acid composition at each position for the four types of host transitions. It also includes references to prior studies that support the role of certain residues in IAV host-specific functions. Analysis of the amino acid preferences at these positions reveals numerous instances of host-specific trends, supporting their potential as genetic markers.

## Discussion

Our study aimed to elucidate the transmission dynamics of IAVs between swine and humans with an emphasis on identifying genetic markers associated with within- and between-species transmission events. The use of comprehensive data curation and phylogenetic analysis resolved technical challenges presented by the large volumes of data associated with finding patterns across all sequenced zoonotic or reverse zoonotic transmissions of IAV between humans and swine. Furthermore, our phylogenetic approach allowed us to investigate the sequences resulting from interspecies transmissions while tracing the evolutionary paths leading to these events.

Initial findings from our phylogenetic analysis reinforced known patterns within IAV evolution in and across human and swine populations, highlighting the considerable diversity of swine influenza viruses and the more linear evolution of human IAVs, punctuated by episodic interspecies transmissions to and from swine [10,74]. This led to a much higher overall representation of swine IAVs in our datasets due to the clustering approach used for trimming down the datasets. Reverse zoonotic (human-to-swine) transmissions appeared more frequently than zoonotic transmissions (swine-to-human) [12], and despite the effort to get the widest possible representation of interspecies transmissions, these were generally rare events, impeding inference gained from statistical modeling. Nonetheless, the analysis of mutation rates and positive selection pressures indicated a notable host-specific influence, particularly following transmission of IAVs from humans to swine. Interestingly, zoonotic transmissions did not exhibit the same selective pressure, which could be caused by the low number of examples, making it difficult to identify general patterns for this type of transmission. However, it may also be associated with the sporadic nature of most of these transmissions (except for the transmission event leading to the pandemic H1N1 in 2009), impeding the accumulation of adaptive mutations, but perhaps also explaining the absence of sustained human-to-human transmissions following these events. In this context, the observations of selective pressures following reverse zoonotic transmissions from humans to swine are also highly informative, as they reveal adaptive pressures in the opposite direction and emphasize some of the differences in host specificity between human and swine hosts.

While our dataset and strategy provided advantageous conditions for broad comparisons across host species, obtaining strong site-specific signals with either of our statistical models remained challenging. Specifically, only two positions in the PA protein exhibited significantly higher substitution rates in swine-to-human transmissions compared to human-to-human transmissions. More positions with higher substitution rates were identified when comparing human-to-swine and swine-to-swine transmissions, yet these signals were difficult to interpret in isolation. A similar challenge arose in the site-specific selection analysis, where the identification of positions under positive selection provided broad trends but lacked consistent site-specific signals across transmission types. Thus, instead of revealing clear mutational patterns, findings from these analyses pointed to broader protein regions of apparent impact, rather than allowing for definitive conclusions about their specific functional roles.

Interestingly, the site-specific selection analyses using both the pervasive and episodic approaches revealed unusually high selective pressures in specific areas of both the PA-X, NS1 and to some degree the M2 protein. However, these were all concentrated in areas of overlapping reading frames with other proteins of the same segments (**Figure 5**), and thus they might reflect functional constraints in the other reading frame of the shared segment, rather than actual high positive selection pressures [75].

The difficulty in identifying strong, consistent mutational signals could be attributed to the high variability within our data, which encompasses various lineages across different time points and geographical locations, all grouped into the same four transmission categories. Within these categories, additional factors likely shaped the viral evolution, such as differing immune profiles driven by natural viral exposures and vaccination practices, which vary by time, region, and host. Therefore, while some mutational signals may have occurred following interspecies transmissions, they might also have been influenced by specific host and environmental pressures at a given time and place. Consequently, some signals of adaptive mutations could be diluted or “washed out” when viewed stochastically, as they could reflect localized host and environmental pressures rather than consistent, widespread evolutionary patterns.

Therefore, we incorporated a machine learning approach to see, if we could enhance pattern recognition and classification of the transmission categories. Interestingly, these models could accurately identify intraspecies transmissions in either swine or humans and were also mostly successful in predicting sequences from reverse zoonotic events, but did not perform as well in predicting zoonotic transmissions. This could partly be explained by our imbalanced datasets and few sequences to train the models in recognizing zoonotic transmissions. However, it could also indicate that the IAVs resemble the originating host more immediately after interspecies events and requires multiple transmissions within the new host species to accumulate adaptive mutations necessary for more efficient transmission. This was also suggested by most of our models, where confusion matrices revealed the frequent misclassification of human-to-swine transmissions as human-to-human transmissions and swine-to-human transmissions as swine-to-swine transmissions. While most previous research on IAV host specificity has focused on the adaptation of avian IAVs to mammalian hosts – a transition that likely presents a higher host-specific barrier – less pronounced adaptive signals may be expected following mammal-to-mammal transmissions, at least initially, where some of the barriers to infection might be smaller. In this analysis, only the immediate zoonotic or reverse zoonotic transmissions were classified, even though potentially important adaptive mutations could follow these events or be hidden as minority variants in the quasispecies cloud. Thus, further analyses into the phylogenetic paths might reveal more important patterns for the host adaptations.

Using machine learning algorithms to evaluate IAV host tropism based on nucleotide, codon, or protein data have recently been investigated in several studies (reviewed in [29]), however mostly focusing on certain lineages, proteins, or avian vs. human hosts. While our approach was efficient for classification and easy interpretability through Shapley values, further explorations of improvements of this or other machine learning algorithms might be able to increase model performaces on our data. Similarly, there could be benefits of training the models on nucleotide or codon rather than protein data [76], even though this may complicate interpretation and increase run time of analyses.

When analyzing the SHAP values for each of our models, we noted that there were some cases where both the presence and absence of a feature had the same predictive impact. For instance, both the absence and presence of glutamic acid (E) at position 53 in the NS1 protein had a negative impact on prediction of the human-to-swine transmission category (i.e. the presence of an E at this site made XGBoost less likely to predict the human-to-swine category, but so did the *absence* of E). There are probably two main reasons why this occurred: first, since we were trying to predict which of four transmission categories a branch belonged to, a given feature might have been mostly informative about two other categories than the one that the focus was on: perhaps the presence of E is predictive for category 1, while the absence of E is predictive for category 2, and in both of those cases category 3 will be less likely to be the correct one. This fits well with the fact that most cases where this occurred were for negative SHAP values. Secondly, there might be some cases where interactions with other sites act as confounders: the SHAP values used here assume linear additive effects of individual features and would not pick up on interactions.

Although not providing definitive answers of individual host-specific markers distinguishing human and swine IAVs, our analyses suggest that some positions within each viral protein may be more important than others for host specificity. For the surface proteins, mutations and selective pressures were mostly concentrated in the head domains for both H1, H3 and N1, N2 subtypes, supporting the expected general immune-escaping adaptive pressure. This is also where most of the XGBoost and SHAP identified positions in these proteins were located, indicating that these positions might be less related to markers specific for swine or human host adaption. Contrastingly, the internal proteins suggested more host-specific patterns with several positions indicating more distinct amino acid patterns in human and swine hosts, respectively. Several positions across the IAV proteins were identified as important for more than one type of transmission category across our analyses, and this in combination with their locations in specific protein functional domains support their potential host-specific importance. Furthermore, some of our final listed positions in **Table 2** have previously been linked to different aspects of host specificity.

While our highlighted positions deserve further investigation to fully determine their importance and potential as host-specific or zoonotic markers, it is also important to mention that our findings suggest a complex interplay within and between IAV segments in different lineages. Thus, considering cooperative effects between segments including investigations of protein or RNA-RNA interactions [77] or epistasis between genes [78] might be crucial for fully understanding the virus’s interspecies transmission and adaptation patterns.

## Acknowledgements

We gratefully acknowledge all data contributors, i.e., the authors and their originating laboratories responsible for obtaining the specimens, and their submitting laboratories for generating the genetic sequence and metadata and sharing via the GISAID initiative as well as the NCBI Influenza Virus Resource database, on which this research is based.

## Supplementary Data

Supplementary data available in separate file.

## Data availability

Data and scripts, including lists of accession numbers for used sequences, are available at https://github.com/KMAnker/IAV_ancestral_reconstruction.git

