## Supplemental tables and figures for "Exploring genetic signatures of zoonotic influenza A virus at the swine-human interface with phylogenetic and ancestral sequence reconstruction"

### Supplementary Tables and Figures

**Table S1**: Overview of the number of samples used for the final phylogenetic inferences of each viral segment. Evolutionary rates for each segment are listed as nucleotide substitutions per site per year and r-squared ($r^2$) values measuring the goodness-of-fit of the root-to-tip regression models are provided.

| Segment | Samples | Nucleotide substitutions per site per year | $r^2$ |
|---|---|---|---|
| **PB2** | 856 | $2.21 \times 10^{-3}$ +/- $2.26 \times 10^{-5}$ | 0.04 |
| **PB1** | 748 | $3.08 \times 10^{-3}$ | 0.96 |
| **PA** | 911 | $1.59 \times 10^{-3}$ +/- $2.07 \times 10^{-5}$ | 0.34 |
| **HA H1** | 1215 | $3.34 \times 10^{-3}$ | 0.84 |
| **HA H3** | 382 | $4.25 \times 10^{-3}$ | 0.96 |
| **NP** | 659 | $2.48 \times 10^{-3}$ | 0.97 |
| **NA N1** | 689 | $4.91 \times 10^{-3}$ | 0.32 |
| **NA N2** | 801 | $3.76 \times 10^{-3}$ | 0.91 |
| **MP** | 436 | $1.68 \times 10^{-3}$ | 0.74 |
| **NS** | 685 | $2.25 \times 10^{-3}$ | 0.84 |

**Figure S1:** Time-scaled and host annotated phylogenetic trees for all IAV segments. Nodes in the trees are colored based on their actual or inferred host origin as either human (red) or swine (cyan), and branches are colored based on the host state of their end node. Below each phylogenetic tree, the root-to-tip regression plot created by TreeTime is shown. **a.** PB2 segment. **b.** PB1 segment. **c.** PA segment. **d.** H1 segment rot-to-tip regression plot **e.** H3 segment. **f.** NP segment. **g.** N1 segment. **h.** N2 segment. **i.** MP segment. **j.** NS segment.

a

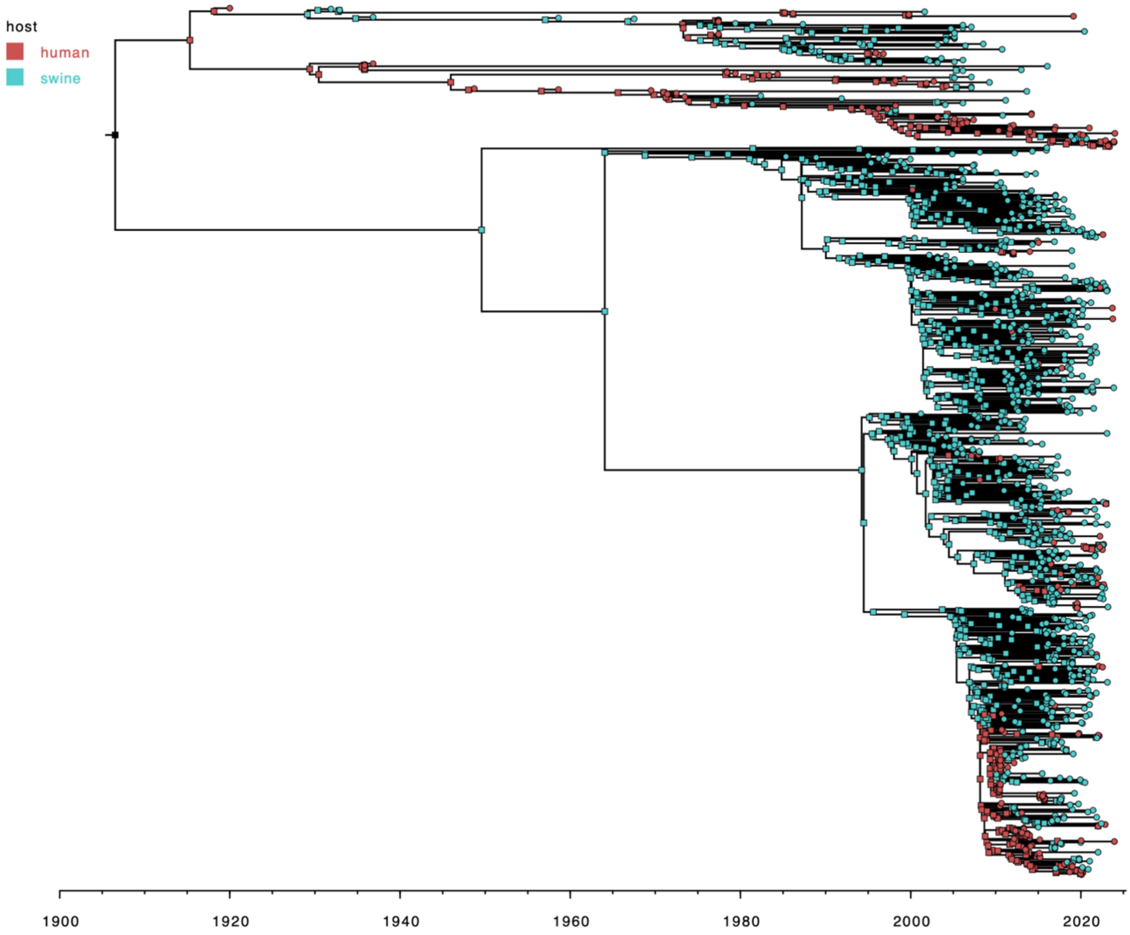
host
human
swine
1900
1920
1940
1960
1980
2000
2020

18

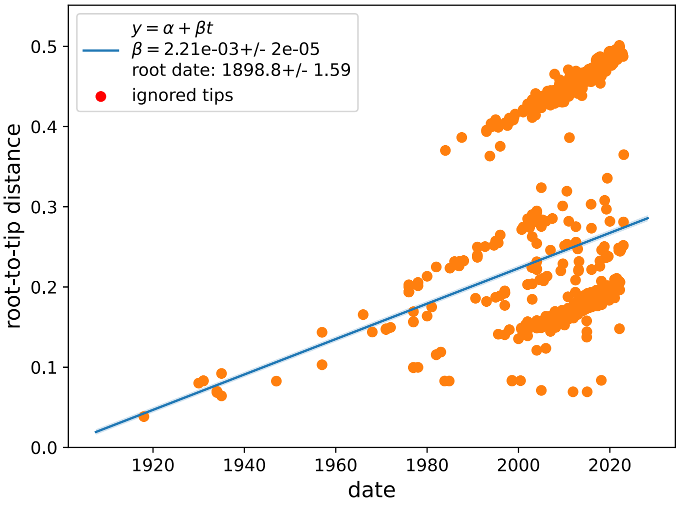
$y = \alpha + \beta t$
$\beta = 2.21e-03+/- 2e-05$
root date: 1898.8+/- 1.59
ignored tips
root-to-tip distance
0.0
0.1
0.2
0.3
0.4
0.5
1920
1940
1960
1980
2000
2020
date

19

b

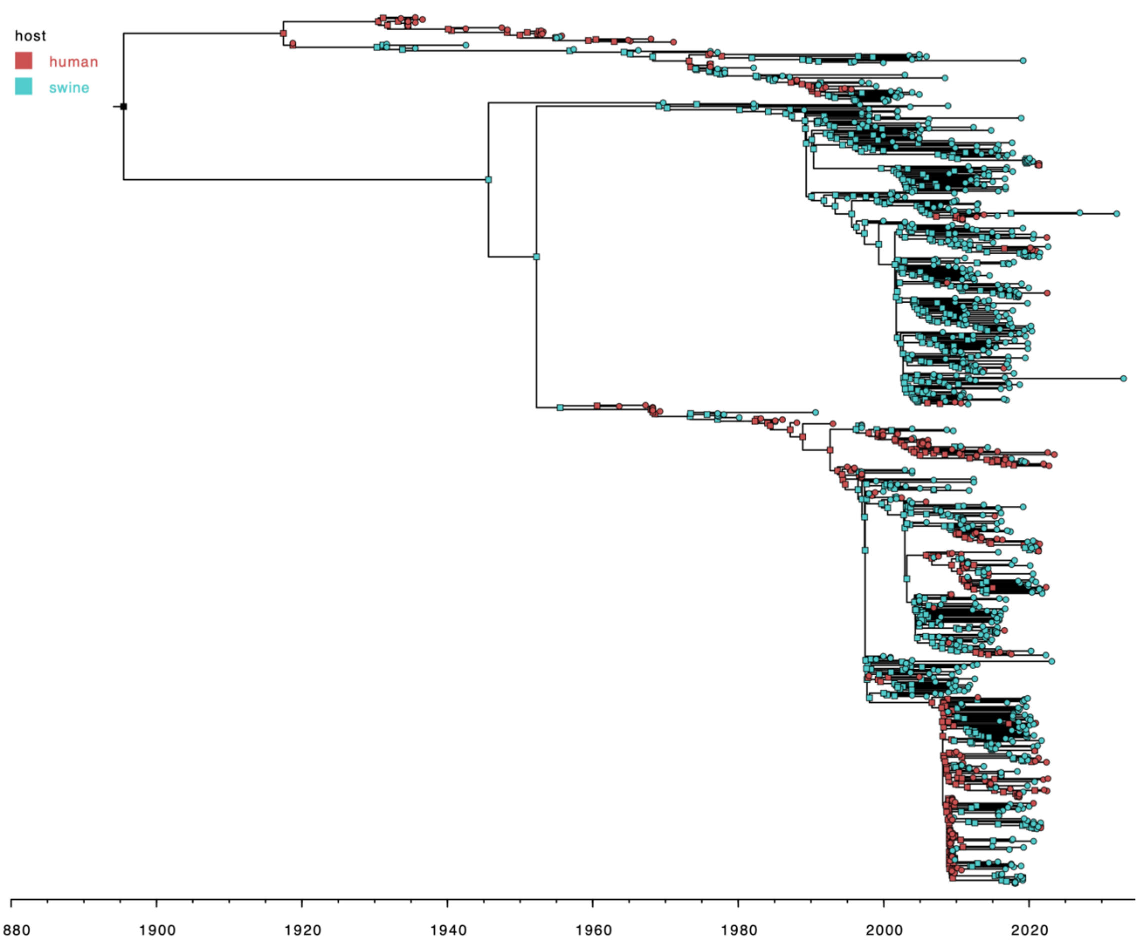
host
human
swine
1880
1900
1920
1940
1960
1980
2000
2020

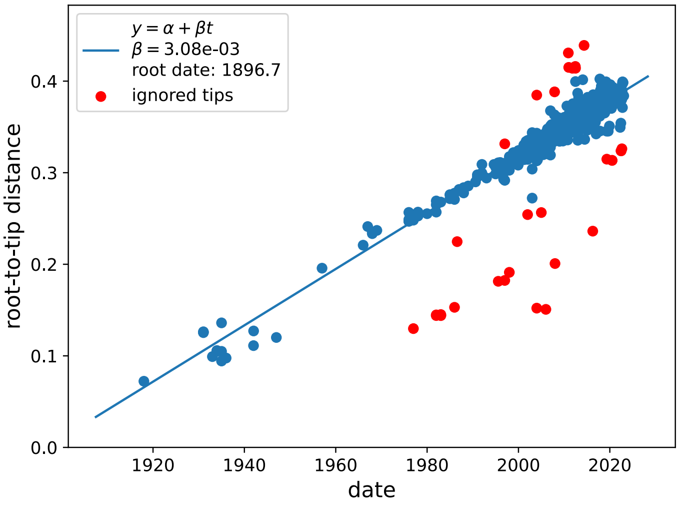
$y = \alpha + \beta t$
$\beta = 3.08e-03$
root date: 1896.7
ignored tips
root-to-tip distance
0.0
0.1
0.2
0.3
0.4
1920
1940
1960
1980
2000
2020
date

c

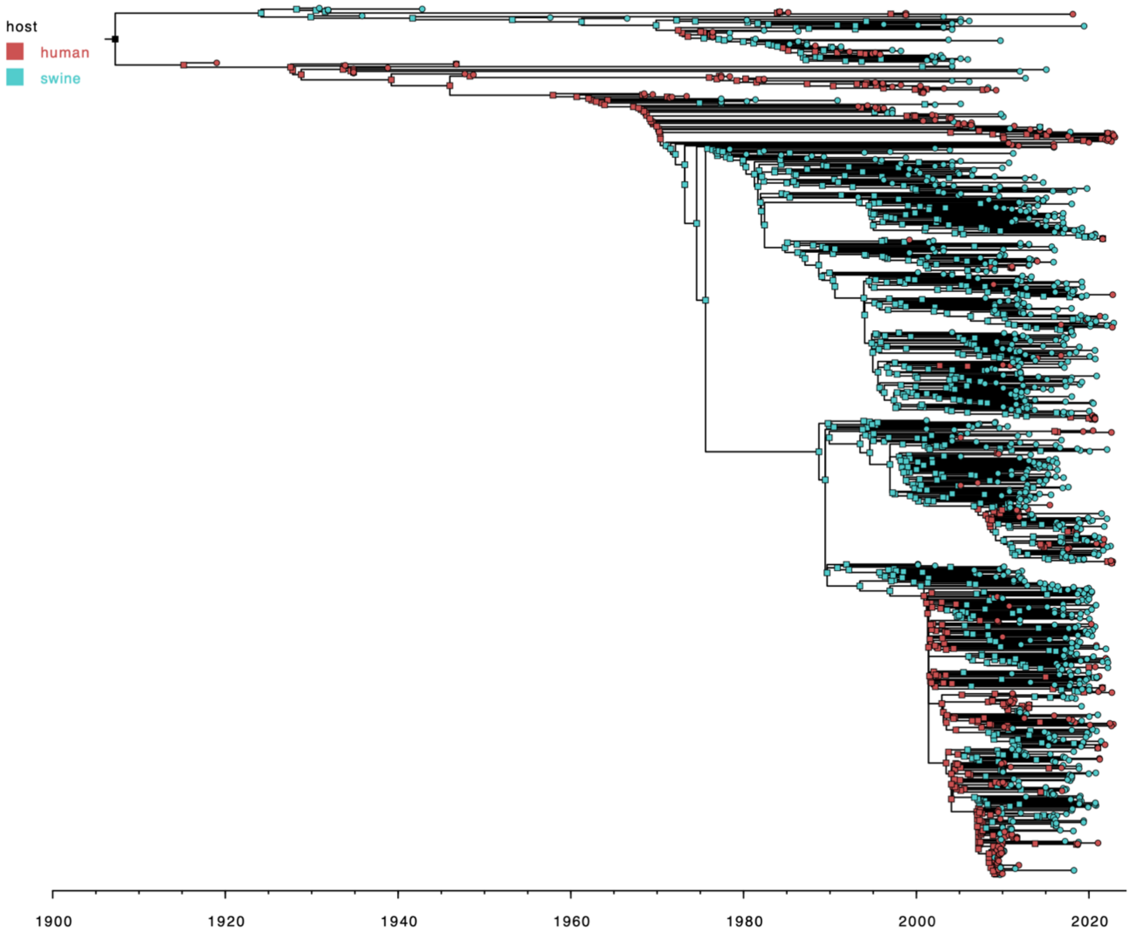
host
human
swine
1900
1920
1940
1960
1980
2000
2020

22

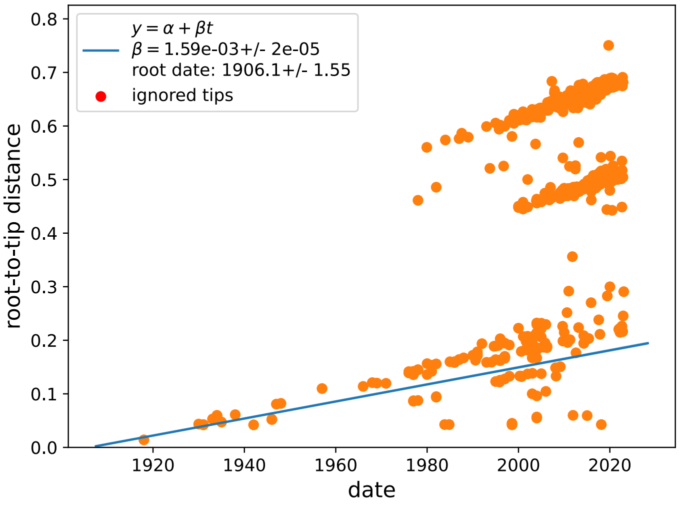
$y = \alpha + \beta t$
$\beta = 1.59e{-}03$+/- 2e-05
root date: 1906.1+/- 1.55
ignored tips
0.8
0.7
0.6
0.5
0.4
0.3
0.2
0.1
0.0
root-to-tip distance
1920
1940
1960
1980
2000
2020
date

23

d

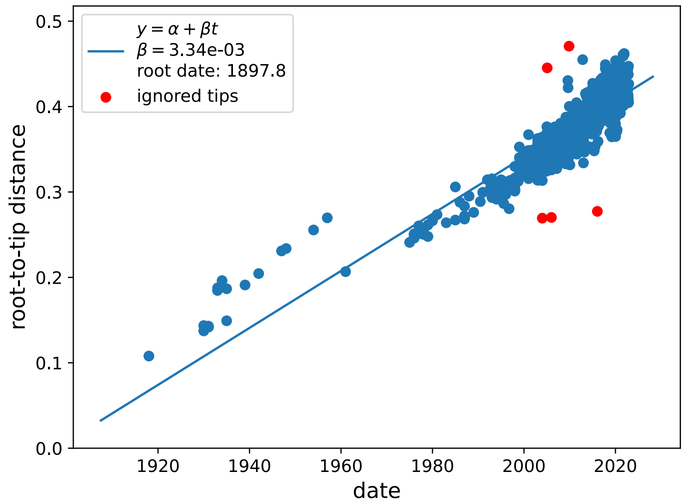
$y = \alpha + \beta t$
$\beta = 3.34e\text{-}03$
root date: 1897.8
ignored tips
root-to-tip distance
date
0.0
0.1
0.2
0.3
0.4
0.5
1920
1940
1960
1980
2000
2020

e

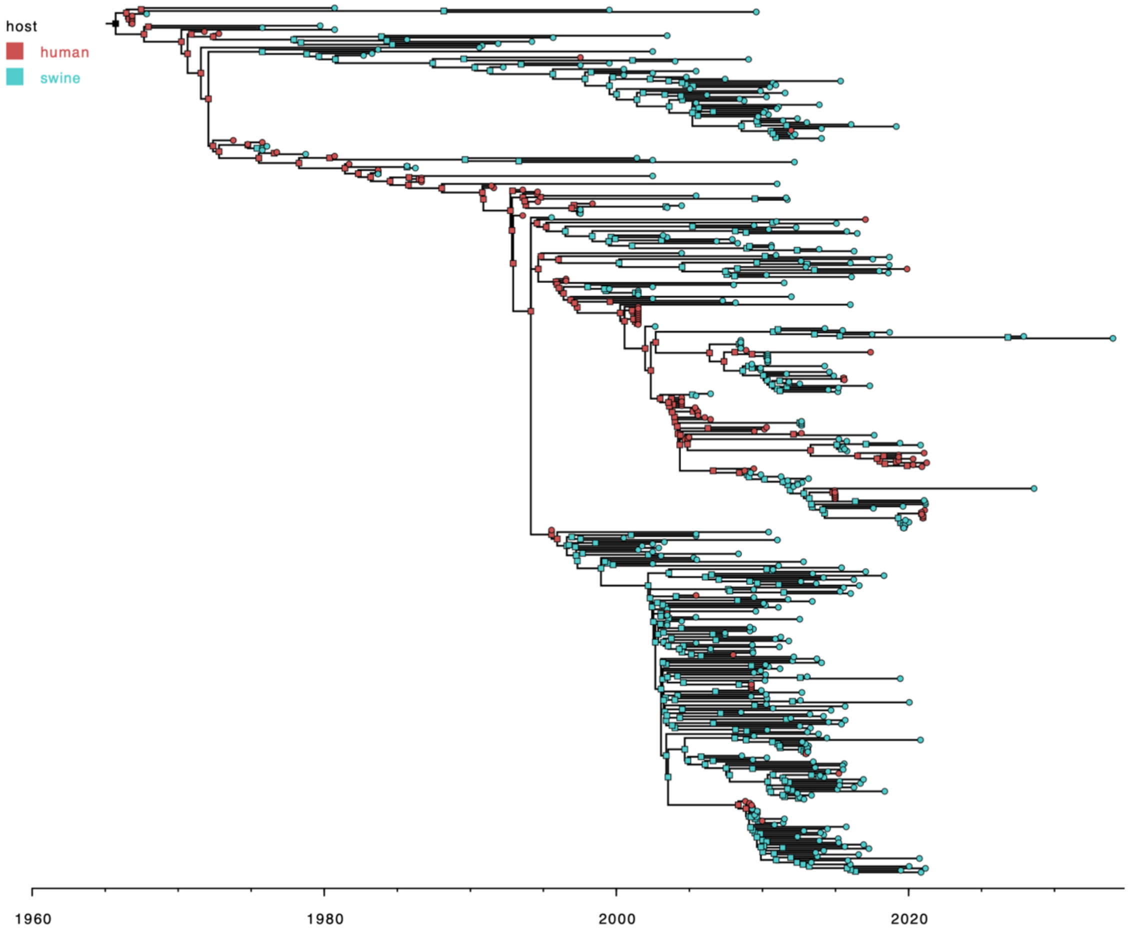
host
human
swine
1960
1980
2000
2020

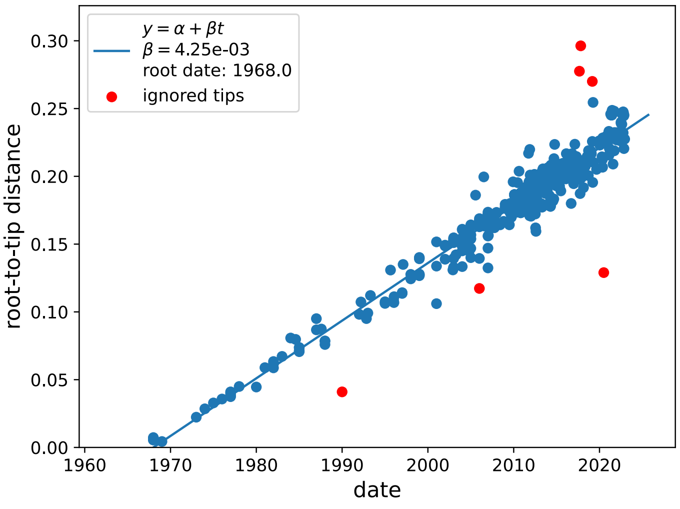
$y = \alpha + \beta t$
$\beta = 4.25e{-}03$
root date: 1968.0
ignored tips
root-to-tip distance
date
0.00
0.05
0.10
0.15
0.20
0.25
0.30
1960
1970
1980
1990
2000
2010
2020

f

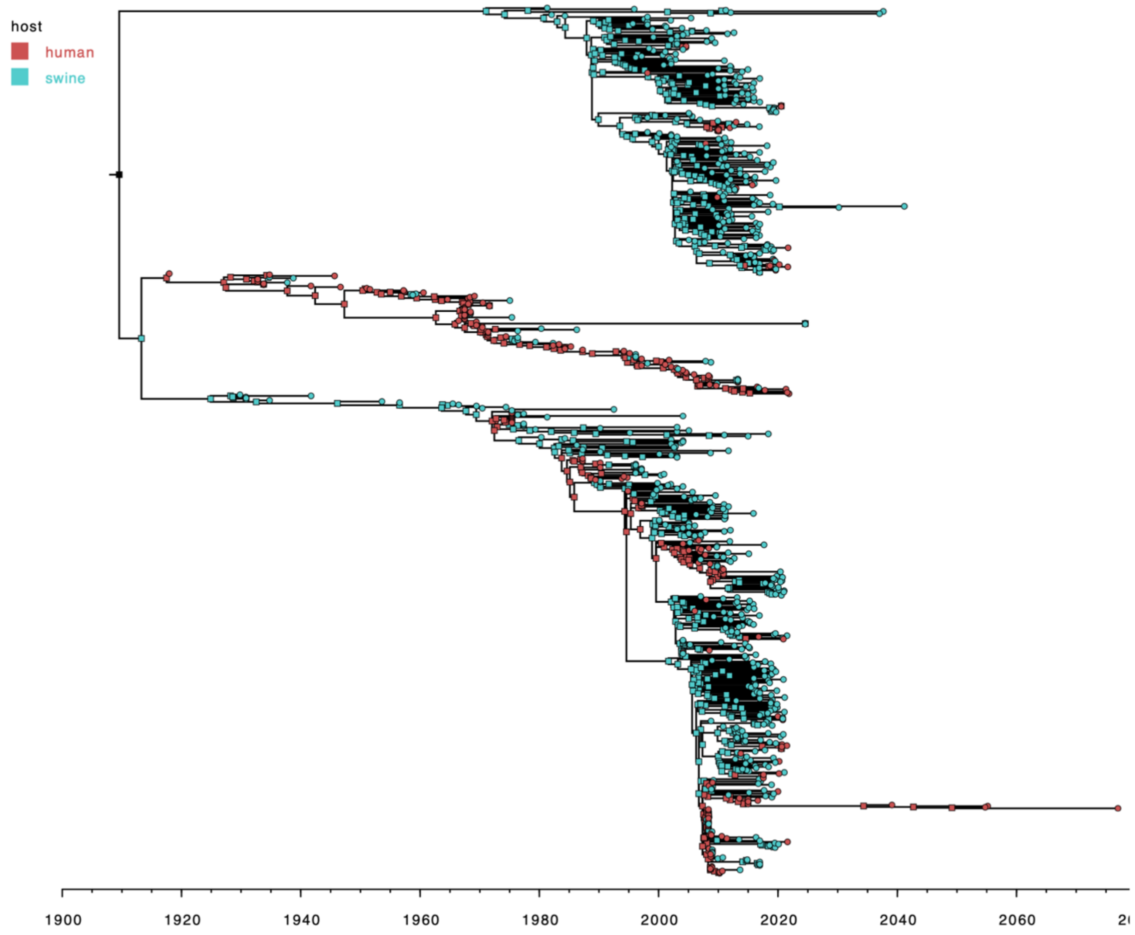

host
human
swine
1900
1920
1940
1960
1980
2000
2020
2040
2060

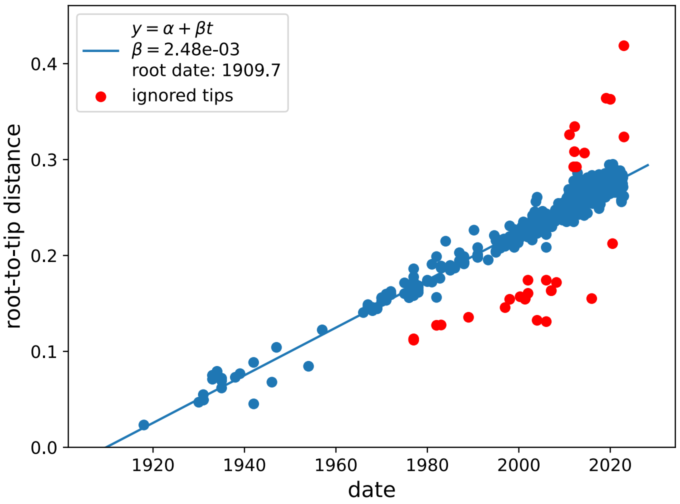

$y = \alpha + \beta t$
$\beta = 2.48e\text{-}03$
root date: 1909.7
ignored tips
root-to-tip distance
0.0
0.1
0.2
0.3
0.4
1920
1940
1960
1980
2000
2020
date

g

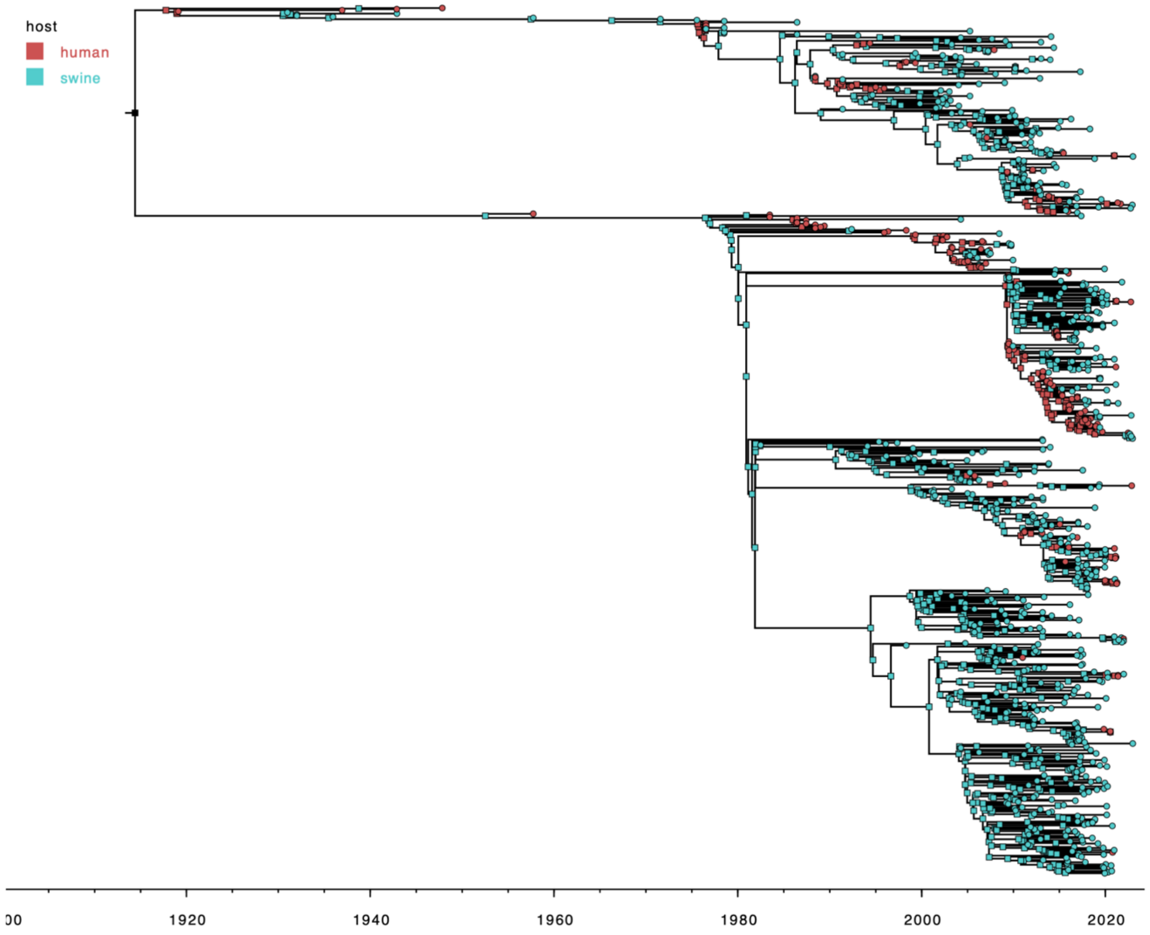
host
human
swine
00
1920
1940
1960
1980
2000
2020

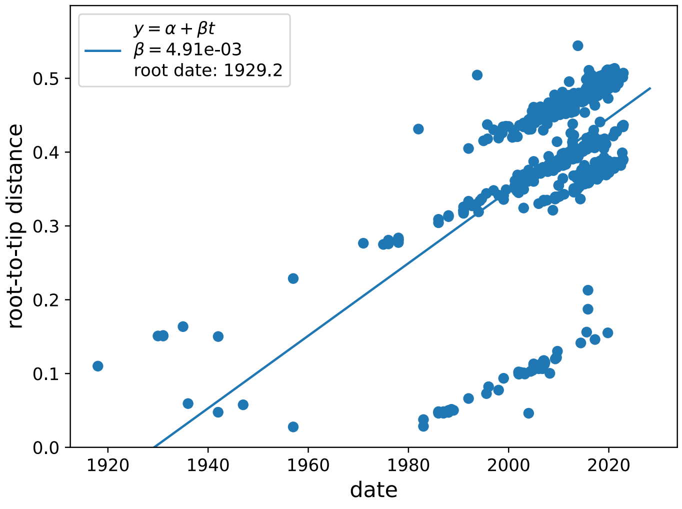
$y = \alpha + \beta t$
$\beta = 4.91e\text{-}03$
root date: 1929.2
root-to-tip distance
0.0
0.1
0.2
0.3
0.4
0.5
1920
1940
1960
1980
2000
2020
date

h

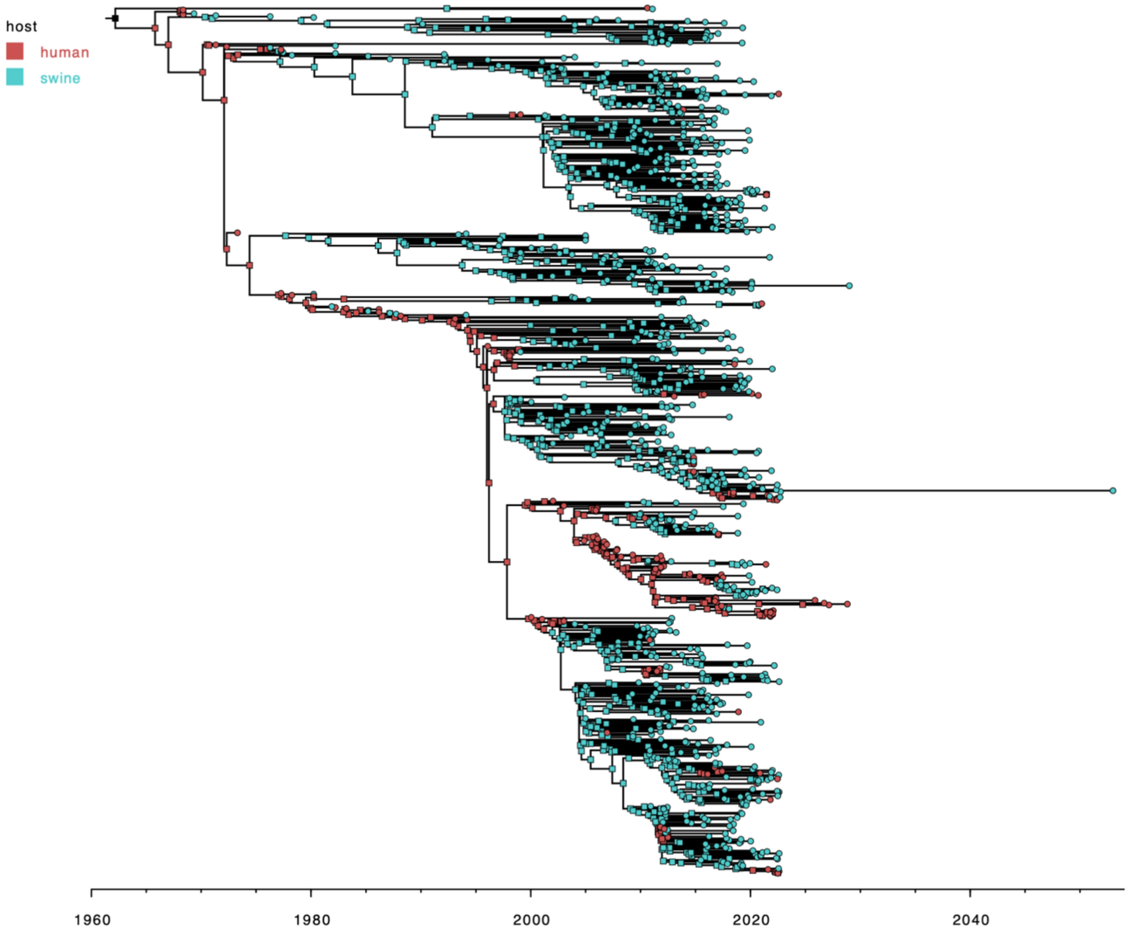
host
human
swine
1960
1980
2000
2020
2040

33

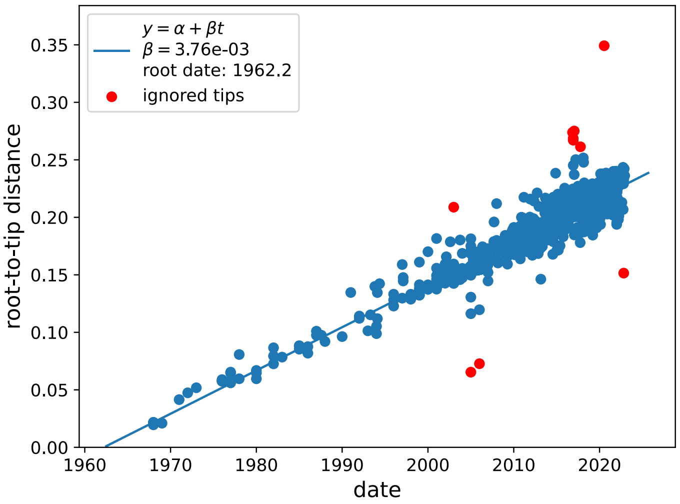
$y = \alpha + \beta t$
$\beta = 3.76e-03$
root date: 1962.2
ignored tips
root-to-tip distance
date
0.35
0.30
0.25
0.20
0.15
0.10
0.05
0.00
1960
1970
1980
1990
2000
2010
2020

34

i

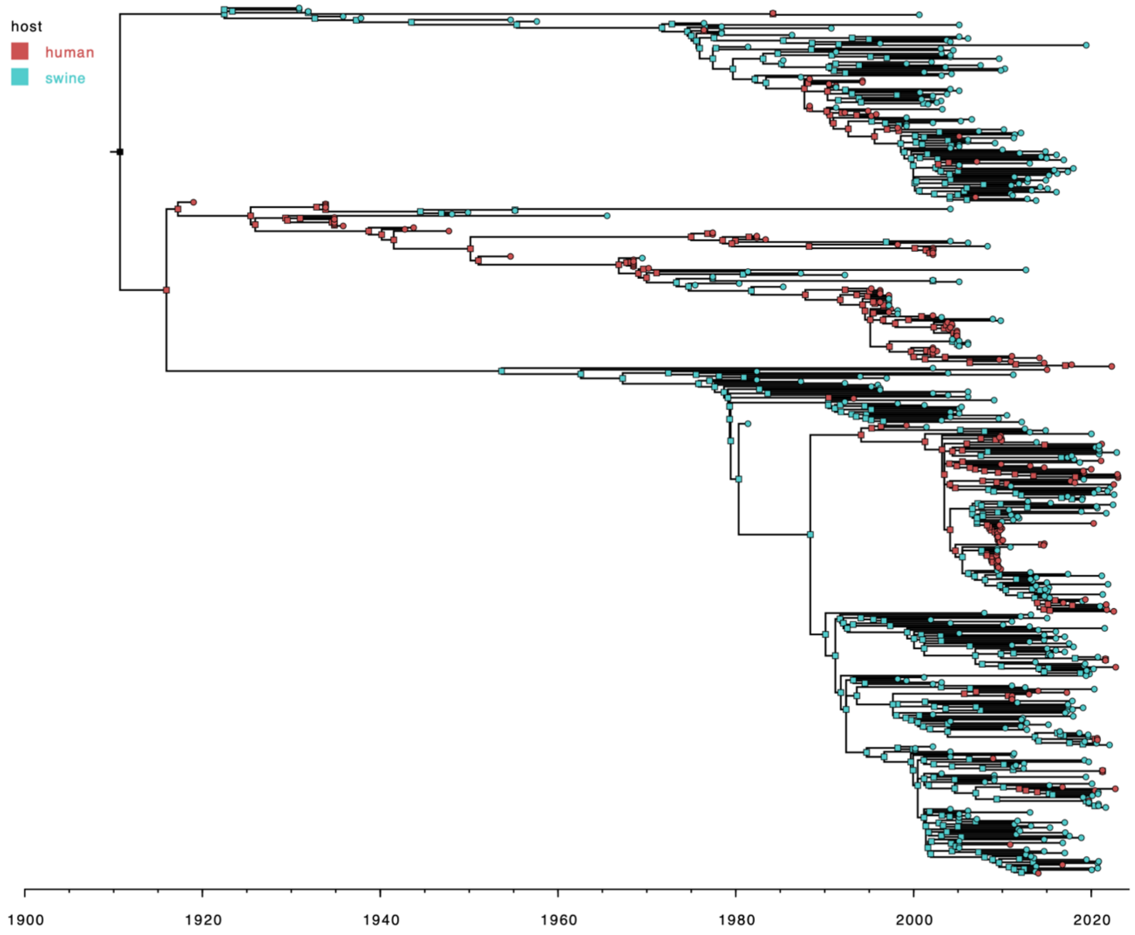
host
human
swine
1900
1920
1940
1960
1980
2000
2020

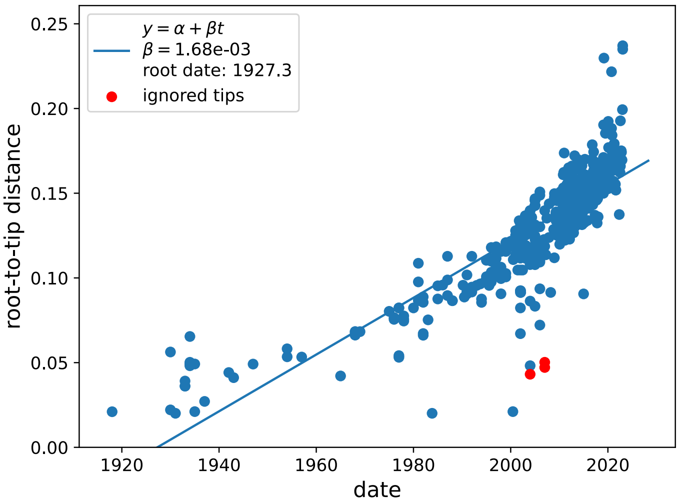
$y = \alpha + \beta t$
$\beta$ = 1.68e-03
root date: 1927.3
ignored tips
root-to-tip distance
date
0.00
0.05
0.10
0.15
0.20
0.25
1920
1940
1960
1980
2000
2020

j

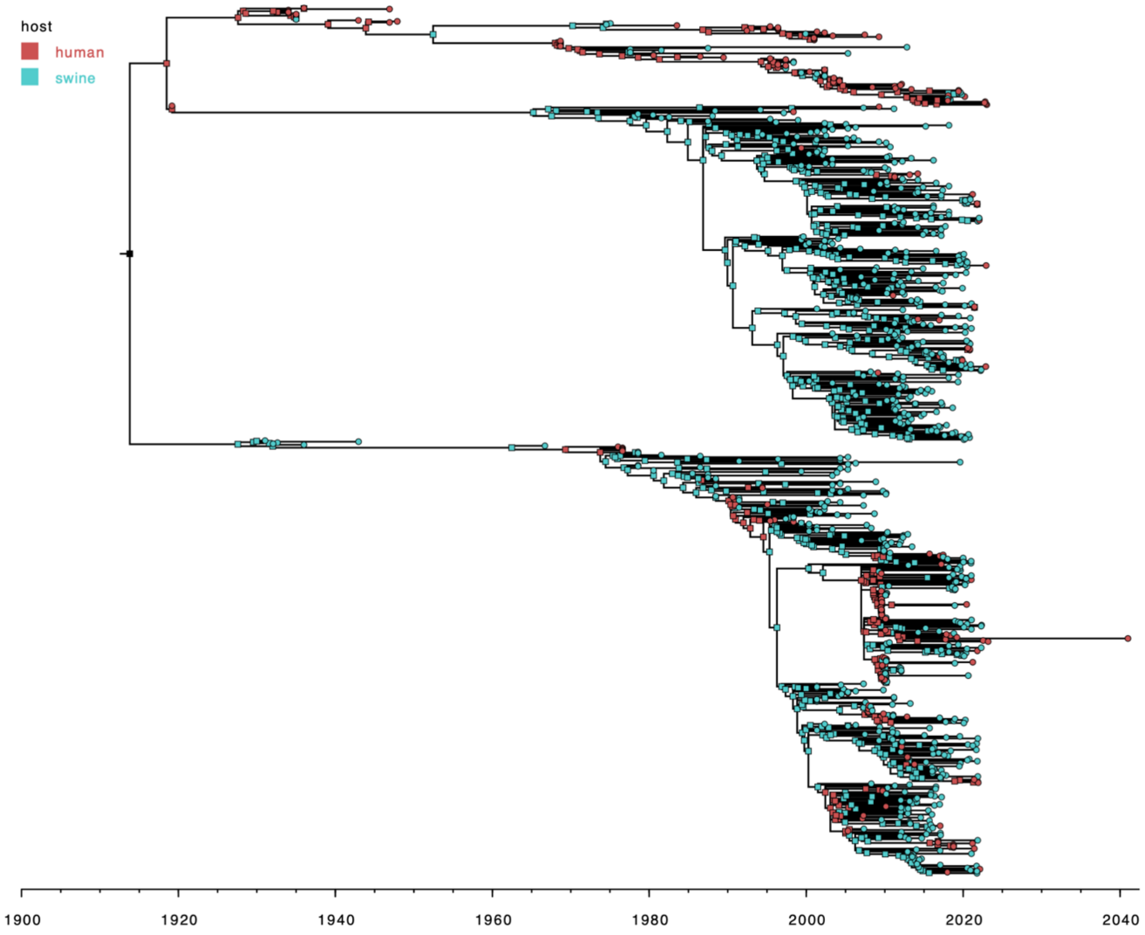
host
human
swine
1900
1920
1940
1960
1980
2000
2020
2040

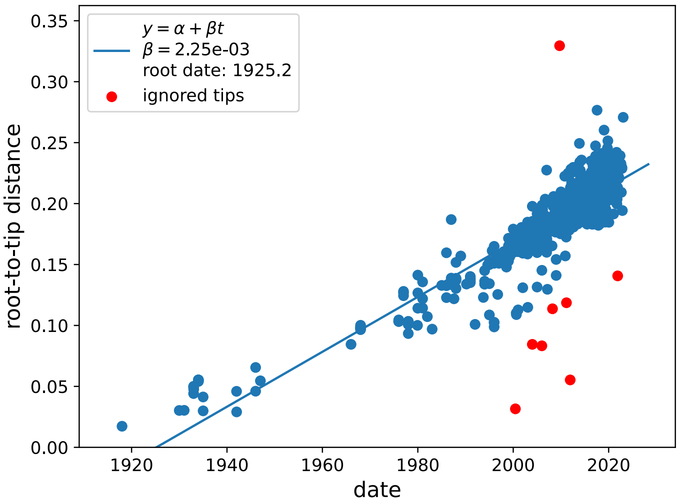
$y = \alpha + \beta t$
$\beta = 2.25e\text{-}03$
root date: 1925.2
ignored tips
root-to-tip distance
date
0.00
0.05
0.10
0.15
0.20
0.25
0.30
0.35
1920
1940
1960
1980
2000
2020

**Figure S2**: Phylogenetic trees for the PB1 (**a**), PA (**b**) and NS (**c**) segments annotated by the lengths of the PB1-F2, PA-X, and NS1 proteins respectively. The lengths are calculated as the number of amino acids from start to the first stop codon and are annotated by colored lines from each tip in the trees.

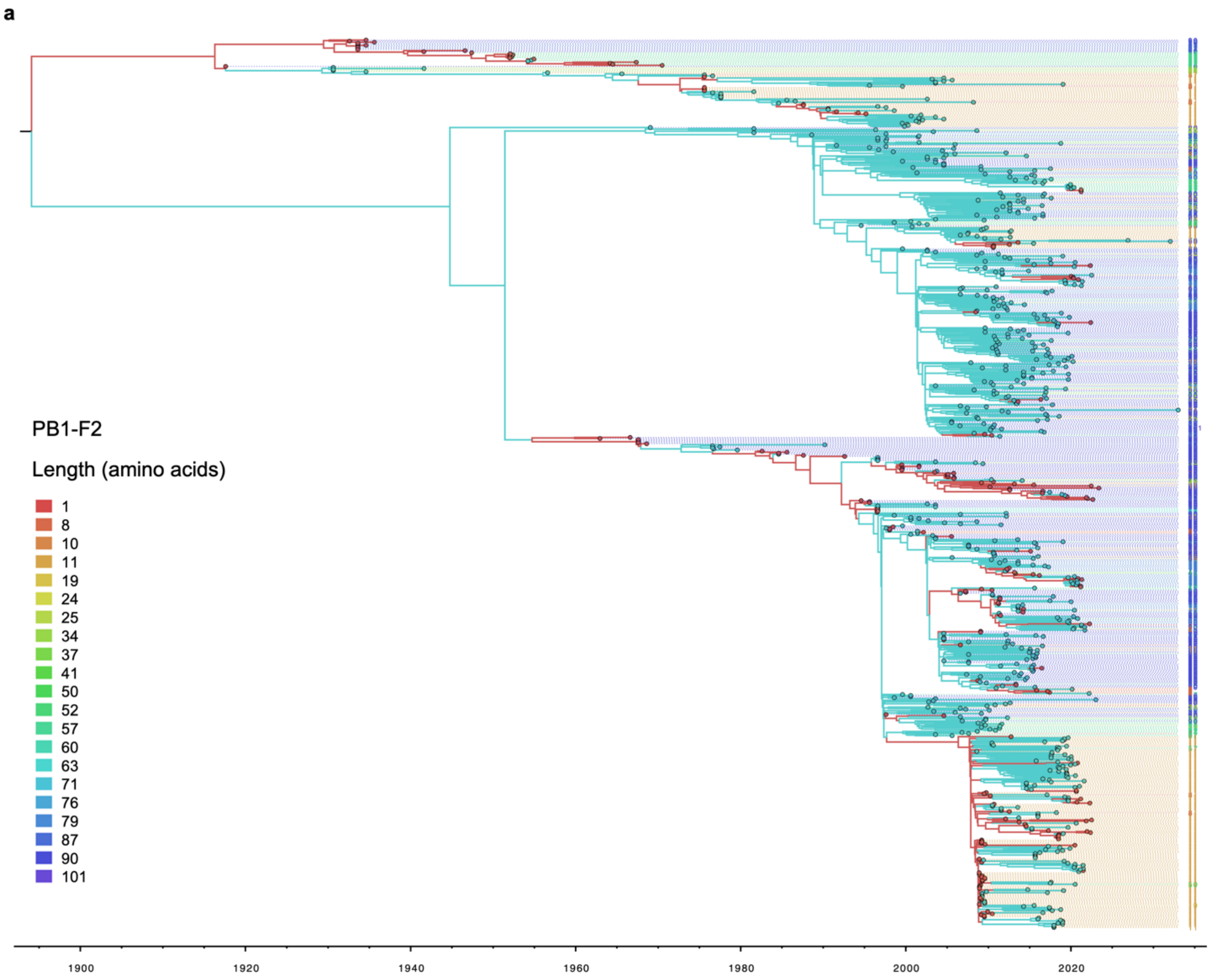

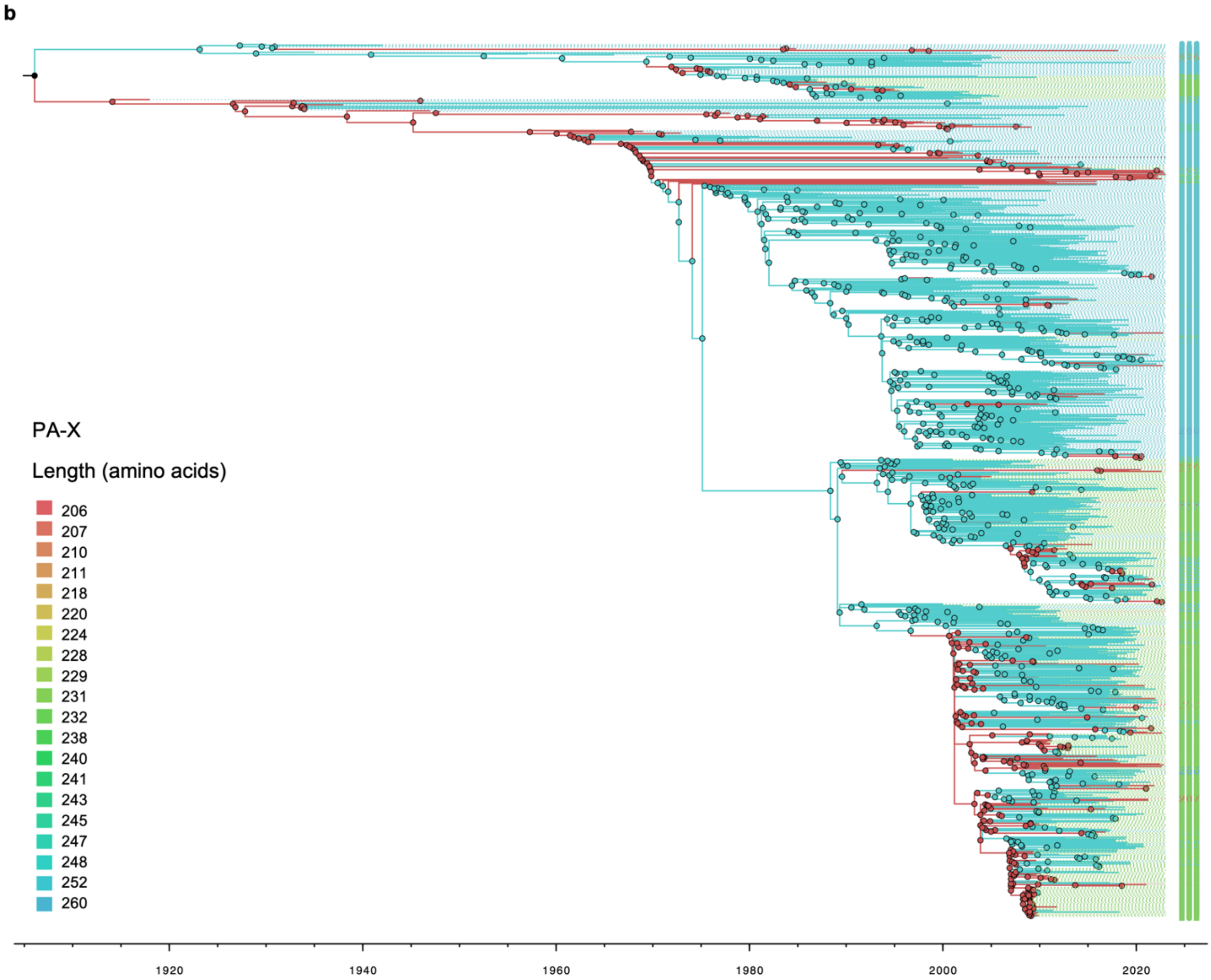
b
PA-X
Length (amino acids)
206
207
210
211
218
220
224
228
229
231
232
238
240
241
243
245
247
248
252
260
1920
1940
1960
1980
2000
2020

c

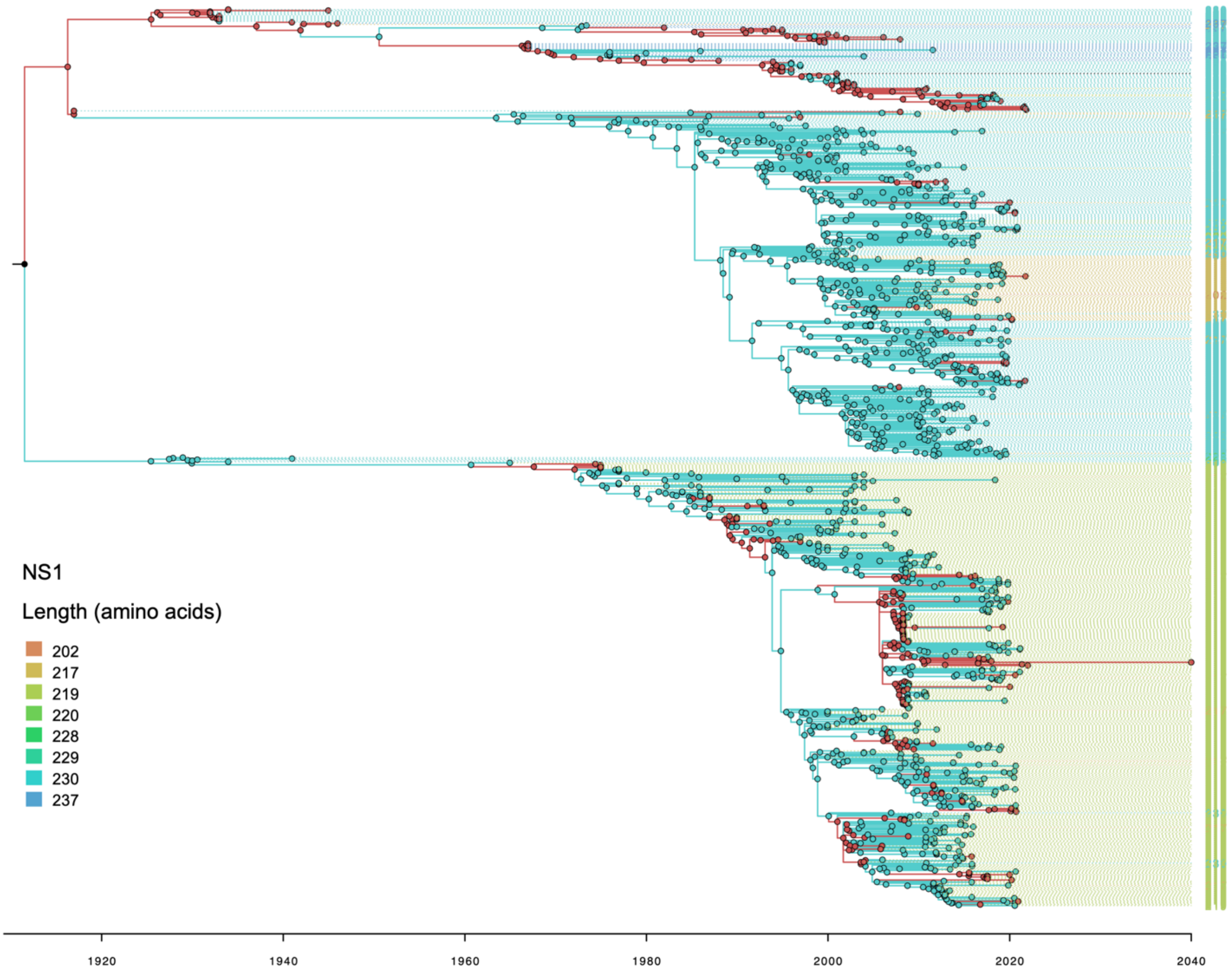
NS1
Length (amino acids)
202
217
219
220
228
229
230
237
1920
1940
1960
1980
2000
2020
2040

**Table S2**: Branches under positive selection identified by aBSREL analysis. Branches are listed with significant positive selection evidence, branch length, and host transmission type (sw = swine, hu = human), including the ancestral node in parenthesis. The likelihood ratio test supports positive selection, with a test p-value threshold of ≤ 0.05 after adjusting for multiple comparisons. The ratio of non-synonymous to synonymous substitution rates (dN/dS), denoted as ω, is divided into classes (ω1, ω2, and ω3) indicating neutral (ω = 0) or positive (ω > 0) selection, with their respective proportions over sites indicated. Amino acid changes occurring at the branches are listed, with color highlighting sites under positive selection by the MEME analysis, orange for positive selection within the same transmission category and blue for positive selection in a different transmission category. Positions in all proteins are counted from the start codon. Branches where positive selection seems to be associated with alignment gaps (many mutations to or from "X") have been shaded in dark grey and are not accounted for in further analysis.

| Segment (Protein) | Branch (length) | Transmission type | Likelihood Ratio Test | p-value | ω distribution over sites | Amino acid changes |
|---|---|---|---|---|---|---|
| PB2 | Node_548 – Seq_312 (6.08195) | (Sw)-sw-sw | 45.7504 | 0.0000 | $\omega_1$ = 0.0837 (99%) $\omega_2$ = 188 (1.0%) | H127Y, V295M, R299K, T364A, R664K, T666L, I667D, L668T, G669P, K670A, D671K |
| | Node_606 – Seq_438 (6.76317) | (Sw)-sw-sw | 74.3236 | 0.0000 | $\omega_1$ = 0.398 (96%) $\omega_2$ = 0.446 (2.2%) $\omega_3$ = 893 (1.8%) | M64N, M66I, P68L, R70K, W78L, S79I, L154I, S453F, L648V, V649D, R650D, S653E, P654I, V655L, F656N, N659S, A661E, T662N, E677D |
| | Node_667 – Seq_666 (6.11243) | (Sw)-sw-sw | 46.2322 | 0.0000 | $\omega_1$ = 0.167 (99%) $\omega_2$ = 646 (1.1%) | A44S, Y205H, I398V, I463M, P515L, Q522H, G523R, E525V, K526F, T530S, N540T, G541P, N556I |
| | Node_108 – Seq_017 (0.56291) | (Sw)-sw-sw | 20.8747 | 0.0172 | $\omega_1$ = 0.00 (100%) $\omega_2$ = 100000 (0.27%) | E6R, W78G |
| | Node_10 – Seq_804 (0.39905) | (Hu)-hu-hu | 19.4865 | 0.0345 | $\omega_1$ = 10000000000 (100%) | E391D, I554T |
| PB1 | Node_384 – Seq_071 (3.67124) | (Hu)-hu-hu | 44.5574 | 0.0000 | $\omega_1$ = 0.611 (99%) $\omega_2$ = 7570 (1.3%) | D27Y, Y30D, S31R, G33V, G35R, T42I, N44H, T57K, N77K, E78G, P79A, T243P, Y324H, T326P, T385P, R468G, T493A, D658E, T663P, W666G, E686V, Q687P, M688H, Y689P, Q690P, C693R |
| | Node_361 – Seq_219 (1.4607) | (Sw)-sw-sw | 33.9238 | 0.0000 | $\omega_1$ = 0.00 (99%) $\omega_2$ = 4980 (0.72%) | V12E, T85Q, Q127K, D565H, L590F |
| | Node_556 – Seq_403 (1.04465) | (Sw)-sw-sw | 32.4759 | 0.0000 | $\omega_1$ = 0.793 (99%) $\omega_2$ = 100000 (0.60%) | K168E, I475N, S578K, F700S, P701L |
| | Node_486 – Seq_380 (0.62759) | (Hu)-hu-sw | 30.0587 | 0.0002 | $\omega_1$ = 0.00 (100%) $\omega_2$ = 100000 (0.27%) | I746D, C747L |
| | Node_493 – Seq_513 (0.3577) | (Sw)-sw-sw | 28.7716 | 0.0003 | $\omega_1$ = 0.00 (100%) $\omega_2$ = 100000 (0.36%) | D581L, E582D, S585A |

| | | | | | | |
|---|---|---|---|---|---|---|
| | Node_492 – node_493 (Node_1351) (0.30539) | (Sw)-sw-sw | 23.3773 | 0.0043 | $\omega_1$ = 0.00 (99%) $\omega_2$ = 1390 (0.92%) | N455M, H456D, E457K, I459A, Q460K, Q584P |
| | Node_675 – Seq_188 (1.28625) | (Hu)-hu-hu | 23.2869 | 0.0045 | $\omega_1$ = 0.00 (99%) $\omega_2$ = 1030 (1.2%) | N597K, L598X, Y599X, N600X, I601X, R602X, N603X, L604X, H605X, I606X, P607X, E608X, V609X, C610X, L611X, K612X, W613X, E614X, L615X, D617E, D618V, D619V, Y620Q, R621Q, G622T, L624X |
| | Node_625 – node_626 (Node_1132) (0.54818) | (Sw)-sw-sw | 27.7361 | 0.0097 | $\omega_1$ = 0.881 (100%) $\omega_2$ = 10000 (0.17%) | Q202H, S494A |
| | Node_28 – Seq_323 (0.04549) | (Sw)-sw-sw | 21.1929 | 0.0128 | $\omega_1$ = 10000000000 (100%) | K736E |
| | Node_404 – Seq_154 (1.05703) | (Hu)-hu-hu | 18.6383 | 0.0461 | $\omega_1$ = 0.00526 (99%) $\omega_2$ = 780 (0.75%) | S702N, S703I, S704P, Y705W, R706N |
| **PB1 (PB1-F2)** | Node_0 – node_79 (Node_162) (50.6796) | (?)-sw-sw | 20.1990 | 0.0152 | $\omega_1$ = 10000000000 (100%) | G14E, T18I, E22G, R29K, H33P, R37Q, V50G, P52H, I55T, Y57S, R65K, V70G, H86Q |
| **PA** | Node591 – node_592 (Node1102) (0.49406) | (Sw)-hu-hu | 33.6726 | 0.0000 | $\omega_1$ = 10000000000 (100%) | G388S |
| | Node_452 – Seq_0258 (6.67697) | (Sw)-sw-sw | 36.1129 | 0.0000 | $\omega_1$ = 0.235 (99%) $\omega_2$ = 100000 (0.56%) | C95S, K158R, T210A, A215V, L219F, P220L, R230T, P238E, N239W, G240R, M595I |
| | Node_031 – node_032 (Node100) (0.24643) | (Hu)-hu-hu | 30.4727 | 0.0001 | $\omega_1$ = 10000000000 (100%) | L268I |
| | Node_146 – Seq_0221 (3.27501) | (Hu)-hu-hu | 31.1042 | 0.0001 | $\omega_1$ = 0.214 (99%) $\omega_2$ = 66000 (0.77%) | D50X, F51X, H52X, F53X, I54X, N55V, Q57L, G58D, E59D, S60P, I61X, V62X, V63X, E64X, L65X, D66X, D67X, P68X, N69X, A70X, L71X, L72X, K73X, H74X, R75X, F76X, E77X, I78A |
| | Node_425 – Seq_0517 (8.21139) | (Sw)-sw-sw | 31.7168 | 0.0001 | $\omega_1$ = 0.355 (99%) $\omega_2$ = 1060 (0.61%) | H212R, E319D, V438I, M441I, K615N, L666H, L667G, I668N, V669L, Q670L, A671V |
| | Node_034 – Seq_0283 (1.30148) | (Sw)-sw-sw | 29.1081 | 0.0003 | $\omega_1$ = 10000000000 (100%) | R508K, F710V |
| | Node009 – Seq_0177 (19.55068) | (Hu)-hu-hu | 28.3201 | 0.0004 | $\omega_1$ = 10000000000 (100%) | I486K |
| | Node_044 – Seq_0308 (0.15041) | (Hu)-sw-sw | 27.5229 | 0.0006 | $\omega_1$ = 10000000000 (100%) | A369T |
| | Node_775 – 5eq_0836 (7.48185) | (Sw)-sw-sw | 27.7965 | 0.0006 | $\omega_1$ = 0.0415 (98%) $\omega_2$ = 9090 (1.7%) | E2Q, I85T, S186A, T208K, L212H, T263E, L275P, H277S, P355S, R356K, I459V, T534K, L549T, V602A |
| | Node_814 – Seq_0131 (0.84707) | (Hu)-hu-hu | 25.8306 | 0.0015 | $\omega_1$ = 10000000000 (100%) | S224P |
| | Node_343 – Seq_0935 (0.29208) | (Sw)-sw-sw | 24.9349 | 0.0023 | $\omega_1$ = 10000000000 (100%) | H535Y |
| | Node_884 – Seq_0086 (0.289) | (Hu)-hu-hu | 24.4733 | 0.0030 | $\omega_1$ = 10000000000 (100%) | F105L |
| | Node_899 – Seq_0229 (0.28892) | (Hu)-hu-hu | 24.3569 | 0.0031 | $\omega_1$ = 10000000000 (100%) | E114K |

| | | | | | | |
|---|---|---|---|---|---|---|
| | Node_893 – node_894 (Node533) (0.17194) | (Hu)-hu-hu | 24.1119 | 0.0035 | ω₁ = 10000000000 (100%) | M581L |
| | Node_881 – Seq_0174 (0.14767) | (Hu)-hu-hu | 24.0455 | 0.0037 | ω₁ = 10000000000 (100%) | C489S |
| | Node_863 – Seq_0966 (0.1267) | (Sw)-sw-sw | 23.9587 | 0.0038 | ω₁ = 10000000000 (100%) | E258K |
| | Node_288 – Seq_0361 (2.18904) | (Sw)-sw-sw | 22.9342 | 0.0064 | ω₁ = 0.00 (98%) ω₂ = 100000 (1.8%) | A17G, W88S, A183P, V253L, E300Q, D347H, L417V, R559T, M607I, E677Q |
| | Node_560 – Seq_0616 (0.28954) | (Sw)-sw-sw | 22.4550 | 0.0081 | ω₁ = 10000000000 (100%) | K328R |
| | Node_564 – node_565 (Node1020) (0.83882) | (Sw)-hu-hu | 22.2314 | 0.0091 | ω₁ = 10000000000 (100%) | V100I |
| | Node_898 – Seq_0077 (0.28893) | (Hu)-hu-hu | 21.9917 | 0.0102 | ω₁ = 10000000000 (100%) | V379I |
| | Node_548 – Seq_0456 (0.14956) | (Hu)-sw-sw | 21.9648 | 0.0104 | ω₁ = 10000000000 (100%) | G329C |
| | Node_334 – Seq_0521 (2.31991) | (Sw)-sw-sw | 20.9866 | 0.0169 | ω₁ = 10000000000 (100%) | E351D |
| | Node_341 – Seq_0179 (0.38831) | (Sw)-sw-hu | 20.8081 | 0.0185 | ω₁ = 10000000000 (100%) | D682N |
| | Node_447 – Seq_0186 (0.10502) | (Hu)-hu-hu | 20.6961 | 0.0195 | ω₁ = 10000000000 (100%) | A70T |
| **PA (PA-X)** | Node_146 – Seq_0221 (3.27501) | (Hu)-hu-hu | 27.5073 | 0.0006 | ω₁ = 0.427 (98%) ω₂ = 63700 (2.3%) | D50X, F51X, H52X, F53X, I54X, N55V, Q57L, G58D, E59D, S60P, I61X, V62X, V63X, E64X, L65X, D66X, D67X, P68X, N69X, A70X, L71X, L72X, K73X, H74X, R75X, F76X, E77X, I78A |
| | Node_775 – Seq_0836 (7.48185) | (Sw)-sw-sw | 25.6317 | 0.0015 | ω₁ = 0.00 (97%) ω₂ = 143 (2.9%) | E2Q, I85T, S186A, L205S, Q208R, S212I |
| | Node_712 – Seq_0492 (10.17196) | (Hu)-hu-sw | 25.1383 | 0.0019 | ω₁ = 0.505 (92%) ω₂ = 79.3 (7.6%) | H144C, F148L, F150S, D160E, Y161L, E165A, A169Y, I171F, T173M, F176C, R179M, Q180H, E181Q, R185M, L187R, D189N, K198M, R199K, K202E, K203E, L205S, L207I, C211S |
| **HA (H1)** | Node_688 – Seq_0362 (1.89771) | (Sw)-sw-sw | 61.4624 | 0.0000 | ω₁ = 0.505 (97%) ω₂ = 227 (3.0%) | V33D, T35K, V36I, E38P, T42K, S46R, V47E, E51L, H54Q, C59S, K62R, I64R, A65S, L67Q, L69V, G70A, N71S, N73I, A75V, L79S, I88F, V97A, S325T, T332P, N336H, H369Y |
| | Node_595 – Seq_0499 (5.64687) | (Sw)-sw-sw | 50.4977 | 0.0000 | ω₁ = 0.541 (98%) ω₂ = 310 (1.7%) | H45P, H54Q, N55F, K57M, L58R, K62S, G63V, A65C, Q68R, G70K, N71T, S73T, V74N, W77R, E83K, D114N, V169A, N171D, L173S, G224E, T283I, N393T, V519L |
| | Node_933 – Seq_0500 (0.0929) | (Sw)-sw-sw | 198.1019 | 0.0000 | ω₁ = 0.354 (38%) ω₂ = 1.00 (56%) ω₃ = 100000 (5.9%) | V5I, I133R, F134T, P135R, K136S, A137K, W140X, P141X, N142X, H143X, D144X, T145X, N146Q, G147N, V148M, T149I, A150T, A151X, C152X, P153C, Y154F, A155T, G156R, A157S, S159R, R162K, N163Y, L164P, I165A, W166L, L167N, K169D, |

| | | | | | |
|---|---|---|---|---|---|
| | | | | | E171Q, N172M, Y174N, P175L, K176T, L177N, S178C, K179T, Y181G, I182R, N183V, N184H, K185H, L190P, I192L |
| Node_1007 – Seq_0530 (2.54773) | (Sw)-sw-sw | 58.9458 | 0.0000 | $\omega_1$ = 0.608 (0.080%) $\omega_2$ = 0.650 (99%) $\omega_3$ = 59500 (1.3%) | L5I, I532K, T535P, L540S, F551L, W552S, C554R, G557V, L559T, Q560A, C561R, R562K |
| Node_707 – Seq_0597 (1.46323) | (Hu)-hu-sw | 87.9093 | 0.0000 | $\omega_1$ = 0.00000000658 (97%) $\omega_2$ = 10400 (3.4%) | L69S, N396H, I399L, N403D, N424Y, L432Q, L442F, L451Y, V458E, K459Y, R466S, K470L, I476N, G477R, K496S, D517L |
| Node_726 – Seq_1062 (6.44001) | (Sw)-sw-sw | 45.5379 | 0.0000 | $\omega_1$ = 0.302 (95%) $\omega_2$ = 102 (5.4%) | K136E, H196P, H209D, E211D, N212H, V218L, H221P, R225P, K235W, Q239H, N244I, E251G, T255I, E272Q, S274T, S279P, G280R, S284A, N285H, M288L, N292C, Q299P, I302M, G304V, S305T, Q309H, N310T, H312P, V314L, T315S, P320A, K321T, Y322F, S325F, K327E, M330R, G333M, K466R, I476L, K486G, S497G |
| Node_1027 – Seq_1108 (1.97404) | (Sw)-sw-sw | 40.3591 | 0.0000 | $\omega_1$ = 1.00 (99%) $\omega_2$ = 629 (1.5%) | N142S, H154Y, S201T, N210Y, R221T, R224S, K235N, V250C, P252T, G253V, D254N, I256Q, T261S, A272T, V288I, I316M, G317E |
| Node_604 – Seq_1259 (5.17704) | (Sw)-sw-sw | 40.1559 | 0.0000 | $\omega_1$ = 0.259 (98%) $\omega_2$ = 115 (2.3%) | S53R, S127L, T144N, G148A, A150S, E182K, D238G, F346R, G347Y, A348I, I349F, G359D, M360E, G363V, W364R, N393T, K470Q, M524V |
| Node_901 – Seq_0203 (4.94775) | (Sw)-sw-sw | 28.9494 | 0.0004 | $\omega_1$ = 0.330 (99%) $\omega_2$ = 9090 (1.3%) | A3T, G23D, N27H, V33D, S86L, T99P, K227I, P234W, N244S, I250V, T255K, L264V, P267S, G276N, P287S, T294S, T332R, I337V, Q381L, E400D, Q415H, V519I, V541I |
| Node_232 – Seq_0950 (9.35363) | (Hu)-?-sw | 29.2350 | 0.0004 | $\omega_1$ = 0.0 (92%) $\omega_2$ = 48.2 (7.6%) | D18E, N27Y, V33D, E51Q, C59S, L87V, T89R, W93R, N104D, Y108F, D114H, V125L, P135A, N142I, W166C, P175R, V191E, S202T, P228R, Q239E, T248S, D254E, S279C, S284P, D362H, A387G, V398L, T407S, L442H, E448Q, L456V, C487S, C491W, G525R, S558T |
| Node_652 – Seq_0187 (1.99507) | (Hu)-hu-hu | 26.7378 | 0.0013 | $\omega_1$ = 0.169 (98%) $\omega_2$ = 200 (1.7%) | I201V, D289G, A326S, Y500K, D501K, Y505N, E508K, S509T, L511K, E514D, I516S, G518A, V519G |
| Node_27 – Seq_0309 (6.81738) | (Sw)-sw-sw | 22.6044 | 0.0102 | $\omega_1$ = 0.184 (98%) $\omega_2$ = 38.5 (2.0%) | L86F, A90T, A113T, Q204H, T206P, Y208S, Y212L, P234S, T257I, M282I, I314M, V337I, G390K, I391G, S392E, N393F, I443V |
| Node_1214 – Seq_0141 (3.55733) | (Hu)-hu-hu | 20.9847 | 0.0229 | $\omega_1$ = 0.00 (93%) $\omega_2$ = 72.8 (6.6%) | T14P, L167V, V168G, L192V, A202G, A211G, E251D, D254A, A270P, A301G, S305I, C319W, Y322G, K327R, L328W, L330V, A331G, A348G, A350G, M360V, Y365D, A387G, N438S, T450S, Y502S, Y527S, T535P, A537P, S555C |
| Node_238 – Seq_0097 (0.17597) | (Hu)-hu-hu | 20.7112 | 0.0263 | $\omega_1$ = 10000000000 (100%) | F226L |
| Node_1214 – Seq_0140 (3.56555) | (Hu)-hu-hu | 19.8620 | 0.0403 | $\omega_1$ = 1.00 (97%) $\omega_2$ = 147 (3.1%) | N27T, D31A, D34A, T35P, T42P, D52A, K57T, C59S, L67F, A90P, Y95S, C152M, Y174S, T180P, Y181S, V189A, D203A, A211P, T219P, S220P, T326P, T332P, |

| | | | | | | |
|---|---|---|---|---|---|---|
| | | | | | | A350P, H368P, E372D, A378P, A379P, T384P, A387P, T407P, A408P, F413S, N471T, A473P, Y484S, C491S |
| HA (H3) | Node_190 – node_191 (Node_684) (1.9383) | (Hu)-hu-hu | 37.0272 | 0.0000 | ω1 = 1.00 (98%) ω2 = 100000 (2.0%) | N440S, A441F, L443F, D454G, L455Q, D457A, E459A, M460I, N461I, K462X, L463X, F464X, E465X, K466X, T467X, K468X, K469X, Q470X, L471X, R472X, E473X, N474X, A475X, E476X, D477X, M478X, G479X, N480X, G481X, C482X, F483X, K484X, I485X, Y486X, H487X, K488X, C489X, D490X, N491X, A492X, C493X, I494X, G495X, S496X, I497X, R498X, N499X, G500X, T501X, Y502X, D503X, H504X, D505X, V506X, Y507X, R508X, D509X, E510X, A511X, L512X, N513X, N514X, R515X, F516X, Q517X, I518X, K519X, G520X, V521X, E522X, L523X, K524X, S525X, G526X, Y527X, K528X, D529X, W530X, I531X, L532X, W533X, I534X, S535X, F536X, A537X, I538X, S539X, C540X, F541X, L542X, L543X, C544X, V545X, A546X, L547X, L548X, G549X, F550X, I551X, M552X, W553X, A554X, C555X, Q556X, K557X, G558X, N559X, I560X, R561X, C562X, N563X, I564X, C565X, I566X |
| | Node_191 – Seq_070 (4.05328) | (Hu)-hu-hu | 44.0971 | 0.0000 | ω1 = 0.385 (97%) ω2 = 458 (2.7%) | S61N, N137Y, T144A, R158K, N160T, S440C, F443S, A446S, N449D, I453A, A459V, I461F, X462C, X463F, X464L, X465S, X466D, X467L, X468R, X469S, X470Q, X471L, X472A, X473V, X474G, X475Y, X476M, X477D, X478M, X479G, X480N, X481G, X482C, X483F, X484K, X485I, X486Y, X487H, X488K, X489C, X490D, X491N, X492A, X493C, X494I, X495G, X496S, X497I, X498R, X499N, X500G, X501T, X502Y, X503D, X504H, X505D, X506V, X507Y, X508R, X509D, X510E, X511A, X512L, X513N, X514N, X515R, X516F, X517Q, X518I, X519K, X520G, X521V, X522E, X523L, X524K, X525S, X526G, X527Y, X528K, X529D, X530W, X531I, X532L, X533W, X534I, X535S, X536F, X537A, X538I, X539S, X540C, X541F, X542L, X543L, X544C, X545V, X546A, X547L, X548L, X549G, X550F, X551I, X552M, X553W, X554A, X555C, X556Q, X557K, X558G, X559N, X560I, X561R, X562C, X563N, X564I, X565C, X566 |
| | Node_191 – Seq_071 (3.91082) | (Hu)-hu-hu | 31.5950 | 0.0000 | ω1 = 1.00 (98%) ω2 = 1740 (2.2%) | N161S, A214S, V239I, N328S, S438G, Y439H, S440R, F441C, L444P, V445D, A446E, E448K, Q450Y, T452S, I453W, G454S, Q455E, S458L, I460V, I461V, X462L, X463H, X464S, X465G, X466H, X467T, X468R, X469S, X470Q, X471L, X472R, X473E, X474N, X475A, X476E, X477D, |

| | | | | | | |
|---|---|---|---|---|---|---|
| | | | | | | X478M, X479G, X480N, X481G, X482C, X483F, X484K, X485I, X486Y, X487H, X488K, X489C, X490D, X491N, X492A, X493C, X494I, X495G, X496S, X497I, X498R, X499N, X500G, X501T, X502Y, X503D, X504H, X505D, X506V, X507Y, X508R, X509D, X510E, X511A, X512L, X513N, X514N, X515R, X516F, X517Q, X518I, X519K, X520G, X521V, X522E, X523L, X524K, X525S, X526G, X527Y, X528K, X529D, X530W, X531I, X532L, X533W, X534I, X535S, X536F, X537A, X538I, X539S, X540C, X541F, X542L, X543L, X544C, X545V, X546A, X547L, X548L, X549G, X550F, X551I, X552M, X553W, X554A, X555C, X556Q, X557K, X558G, X559N, X560I, X561R, X562C, X563N, X564I, X565C, X566I |
| | Node_110 – Seq_218 (1.51154) | (Sw)-sw-sw | 33.7977 | 0.0000 | $\omega_1$ = 0.185 (99%) $\omega_2$ = 479 (1.3%) | V4I, F7L, T251D, I252T, T264Q, Y273S, P300S, I304F, K308Y |
| | Node_330 – Seq_413 (1.95566) | (Sw)-sw-sw | 60.6726 | 0.0000 | $\omega_1$ = 0.00 (96%) $\omega_2$ = 106 (4.4%) | G216W, I494L, T501I, D509V, N514H, F516L, Q517L, G520H, V521F, L523M, L525V, W530S, W533G, I534N, A537R, S539A, F541C, L542Q, A545H, L547T, A554P |
| | Node_152 – node_153 (Node_562) (11.63431) | (Sw)-sw-sw | 29.3909 | 0.0001 | $\omega_1$ = 0.163 (97%) $\omega_2$ = 267 (2.7%) | F15L, A16R, V50I, A85S, R108K, V128I, D149N, T171Y, F175I, A179M, V183T, D189A, V208I, R217K, D241G, V258I, L260I, T264N, R277H, I284M, R285K, G291D, K292T, D320A, K342R, I494M, G495E |
| | Node_363 – Seq_192 (1.65352) | (Sw)-sw-sw | 27.2187 | 0.0003 | $\omega_1$ = 0.315 (98%) $\omega_2$ = 98.0 (2.4%) | V146I, A154T, N172K, Q372K, Y439D, S461N, L463K, F464S, R466K, T467K, R468E, Q470N, E476A, D477V, M478T, C482R, Y486H, D490V, G495Q |
| | Node_217 – Seq_350 (15.93445) | (Sw)-sw-sw | 25.9471 | 0.0006 | $\omega_1$ = 0.0962 (95%) $\omega_2$ = 22.7 (5.3%) | N25S, M26T, I41L, E66R, N69D, Q73R, E78K, Q91H, K99E, V104I, H110Y, A122S, N137T, S140D, T147A, A151G, S153Y, A154S, R158G, S160V, T171H, H172K, N174E, F175Y, E181N, E188D, Q189K, T264P, N312K, T329A, L330V, I339R, D364H, Y367D, R370T, N373P, E375G, R377T |
| | Node_263 – Seq_157 (1.64658) | (Sw)-sw-sw | 24.0637 | 0.0016 | $\omega_1$ = 0.838 (100%) $\omega_2$ = 100000 (0.42%) | A179T, R236K, P289S, I560S, C565H, I566G |
| | Node_328 – Seq_249 (1.69911) | (Sw)-sw-sw | 23.1471 | 0.0025 | $\omega_1$ = 0.638 (99%) $\omega_2$ = 1700 (0.57%) | T3A, I304L, L471R, R515V, K557T |
| **NP** | Node_094 – Seq_174 (5.24698) | (Sw)-?-sw | 54.4039 | 0.0000 | $\omega_1$ = 0.681 (95%) $\omega_2$ = 291 (4.7%) | Q42L, G54R, E107K, D114N, Q149L, R150I, R152I, A153P, V155L, G158E, R162T, Q168H, R175T, A178S, A182P, V183L, K184R, R195Q, I197M, N202Y, K227I, G228W, F230L, A233P, A234P, Q235L, A237L, Q241A, V242L, |
| | Node_6 – Seq_385 (28.57858) | (Sw)-sw-sw | 39.5814 | 0.0000 | $\omega_1$ = 0.439 (97%) $\omega_2$ = 286 (3.5%) | A2V, S3L, W207R, P277L, A278D, V280G, A284V, H289Y, D290N, N309S, R317S, P322S, A323V, S326N, Q327H, L328I, |

| | | | | | | |
|---|---|---|---|---|---|---|
| | | | | | | V329I, P334H, F335S, A336S, D340G, V343G, A366E, T378N, S392N, Q409H, P410R, T411S, T423K, T424P, M426R, T442R, M447K, M448K, S450R, R452I, S457A, F458E, D468E, T472Q, V476M, P477Q, D480Q, S486Y, F489Y |
| | Node_503 – Seq_549 (8.70997) | (Sw)-sw-sw | 52.5326 | 0.0000 | ω₁ = 0.0953 (97%) ω₂ = 98.7 (3.2%) | G34A, I109T, L133V, T171S, L172V, P173S, A178R, M196K, K198E, R199S, D203S, R216W, A218S, C223R, I225R, F230L, M239E, V242I, R246P, N247P, V313F |
| | Node_639 – Seq_726 (3.69539) | (Hu)-hu-hu | 61.1303 | 0.0000 | ω₁ = 0.424 (98%) ω₂ = 435 (2.1%) | R26Q, F39L, R55V, L56I, I61T, T62S, R65M, M66P, V67N, S69F, F71S, D72V, E81A, P89L, K91R, I96T, V119E, W120S, G126P, P248Q |
| | Node_433 – Seq_145 (2.03844) | (Hu)-?_sw | 27.3928 | 0.0005 | ω₁ = 0.200 (99%) ω₂ = 100000 (0.79%) | Q149Y, E454K, D455N, D497R, S498R, X499L |
| **NA (N1)** | Node_321 – seq_119 (2.91135) | (Hu)-hu-hu | 42.9902 | 0.0000 | ω₁ = 0.377 (91%) ω₂ = 53.6 (8.9%) | R77K, V80G, K84N, A86G, N88I, P93L, A98F, S101K, D103N, S105R, I108F, K111R, E119N, E128K, A138V, N146E, Y170F, T188I, D330E, S340F, N344K, Y353F, G354N, T362S, F371Y, Q377R, A381G, D384G, N392K, I393K, E398K, F406C, L415P, D416G, C417G, I418V, N434K, T438I, S442G, S444R, C446S |
| | Node_331 – seq_126 (2.8777) | (Hu)-hu-hu | 65.3259 | 0.0000 | ω₁ = 0.328 (94%) ω₂ = 166 (6.1%) | K6R, I8L, I10L, G11C, S12W, C14W, Y402D, E411Q, V424I, I427K, R428K, R430Q, E432N, E433R, N434K, T435I, S439C, G440A, F445I, C446F, G447V, D449N, T452I, V453I, G454F, W455S, S456T, W457C, D459R, G460L, A461W, E462A |
| | Node_335 – seq_127 (2.02968) | (Hu)-hu-hu | 49.6517 | 0.0000 | ω₁ = 0.231 (97%) ω₂ = 359 (2.5%) | E228K, S366I, R368A, F371L, M373S, I374K, W375M, D376N, G379R, W380K, T381I, G382M, D384I, N385H, K386N, F387L |
| | Node_321 – seq_153 (2.72916) | (Hu)-hu-hu | 36.2948 | 0.0000 | ω₁ = 0.543 (94%) ω₂ = 87.6 (6.0%) | A81V, I264V, G333V, C335G, V338A, A343V, G348L, K352N, G354V, N355D, V357G, W358R, I359M, K363S, K369R, I374S, D376P, Q377K, N378K, W380S, T383A, N385K, F387L, S388T, I389K, Q391E, V394G, G395R, K397E, A400G, Y402N, V407D, T413L, L415V, P420S, L426Q, W437C, S441G, I443R, V448D, D449N, D451A, G454C, W455C, G460A, L463S, F465L |
| | Node_168 – Seq_475 (2.03597) | (Hu)-hu-sw | 48.6841 | 0.0000 | ω₁ = 0.111 (99%) ω₂ = 522 (1.2%) | S21N, R64Q, T81A, S385W, S386I, F387R, S388Q, M389Q, D398G |
| | Node_249 – Seq_677 (6.09972) | (Sw)-sw-sw | 48.7125 | 0.0000 | ω₁ = 0.427 (92%) ω₂ = 40.9 (8.1%) | I29T, I34T, Q64P, Y66P, I69L, S70G, A75P, Q78L, S79M, V81I, P82S, V83M, L85I, L91R, C92S, A98D, N104R, V106I, G109S, K111I, G112W, I122K, F133L, L134S, T135H, G147T, T148P, D151I, R152W, S153T, R156Q, C161N, P162H, G164E, E165A, Y170H, F174L, A181P, W190C, T215I, I216R, K217T, R220K, L224F, E228K, E230K, C231I, S237T, C238Y, F239I, A251V, |

| | | | | | | |
|---|---|---|---|---|---|---|
| | | | | | | Y253F, I255V, K265Q, P272R, E278G, C281F, S286N |
| | Node_155 – Seq_350 (3.23388) | (Sw)-sw-sw | 27.8493 | 0.0004 | ω₁ = 1.00 (97%) ω₂ = 99.6 (3.3%) | T9N, M15K, V17I, S21G, L24F, I26M, G27E, N28K, I29L, V30A, W33X, S35Y, H36P, S37F, Q39F, T40P, G41S, H45Y, Q51P, V53G, V57G, N58K, R64P, I69L, N71H, A75G, T76N, D79G, T81A, L85P, S90L, V99G, S101R, N104H, I117L, C129W, R130T, T131N, H144P, M159V, S176L, M389L, D462E |
| | Node_179 – Seq_025 (10.19964) | (Sw)-sw-hu | 27.6185 | 0.0005 | ω₁ = 0.0803 (99%) ω₂ = 720 (0.74%) | A157R, V177I, P250Q, R327P |
| | Node_441 – Seq_622 (3.09441) | (Sw)-sw-sw | 25.1369 | 0.0016 | ω₁ = 0.120 (99%) ω₂ = 100000 (0.95%) | V264I, Y276P, E278S, C279Y, Y282F, E287V, I289T, R430Q |
| | Node_117 – Seq_251 (1.38611) | (Sw)-sw-sw | 24.3434 | 0.0024 | ω₁ = 0.385 (99%) ω₂ = 1120 (0.78%) | I16V, I38V, K43R, I117M, E259K, N329I, K332R, I467Q, D468E, K469Q, X470K |
| **NA (N2)** | Node_531 – seq_092 (3.12813) | (Hu)-hu-hu | 120.9998 | 0.0000 | ω₁ = 0.00 (94%) ω₂ = 422 (6.1%) | N2H, I73X, T138K, K328I, N329I, D330G, S331I, S333T, S334K, H336R, C337S, L338K, P340R, N341I, N342Q, E343I, G346Y, H347Q, K350N, W352F, A353L, F354S, D355P, G357A, M362S, N367Y, T369K |
| | Node_539 – seq_095 (4.29804) | (Hu)-hu-hu | 79.1552 | 0.0000 | ω₁ = 0.551 (97%) ω₂ = 100000 (3.0%) | V396I, I397K, V398R, D399Y, R400N, G401I, D402H, R403A, S404N, G405C, Y406H, S407L, G408D, D463A, L464M, N465S, M467F, I469X |
| | Node_600 – seq_244 (3.58434) | (Sw)-sw-sw | 35.5473 | 0.0000 | ω₁ = 0.535 (97%) ω₂ = 138 (3.3%) | P3Q, Q5K, K6R, S12C, I77L, L81I, Y84N, T153S, A188S, S204R, Y207F, D213N, W218G, I222N, L223F, C230S, C232F, I233T, N234K, C237A, T238P, A246G, D251V, E259K, V275I, C278G, C280V, P285L, C289G, C291F, D330A, S331R, V360M |
| | Node_440 – seq_433 (1.15397) | (Sw)-sw-sw | 31.1080 | 0.0001 | ω₁ = 0.901 (99%) ω₂ = 3280 (1.2%) | L338V, N339Y, P340S, E343S, K350N, V360S, W361X, I366L, G382X |
| | Node_184 – node_185 (node292) (1.77149) | (Sw)-sw-sw | 30.0848 | 0.0002 | ω₁ = 0.303 (98%) ω₂ = 100000 (1.5%) | I47N, T69S, T71I, H184Y, N368E, E369D, F390L, N401R, D402N, E432S |
| | Node_555 – node_557 (node_637) (0.98069) | (Hu)-hu-hu | 30.1084 | 0.0002 | ω₁ = 0.273 (99%) ω₂ = 1050 (0.85%) | G93Q, K220Q, D221N, I307M, K308E, K344R, G345X, G346X, T369D |
| | Node_520 – seq_065 (0.77052) | (Hu)-hu-hu | 24.2632 | 0.0030 | ω₁ = 10000000000 (100%) | R60K, T148I, K369R |
| | Node_437 – seq_434 (1.84403) | (Sw)-sw-sw | 24.2529 | 0.0030 | ω₁ = 1.00 (99%) ω₂ = 986 (1.5%) | D304E, E346K, H347Q, V349M, K350I, G351C, W352E, A353S, D356G, D359A, V360L, W361S, M362L, T365A, I366L, N367T, S416I |
| | Node_4 – seq_024 (0.18805) | (Hu)-hu-hu | 21.7724 | 0.0105 | ω₁ = 10000000000 (100%) | P468H |
| | Node_5 – seq_003 (0.20471) | (Hu)-hu-hu | 21.0719 | 0.0149 | ω₁ = 10000000000 (100%) | E74D, I77K |
| | Node_219 – node_220 (node_029) (3.53197) | (Sw)-sw-sw | 19.1275 | 0.0395 | ω₁ = 0.00 (95%) ω₂ = 29.7 (4.7%) | I17F, F37S, E41W, C42Y, S44P, A46T, N48S, I50M, M51X, P52X, I58V, T69A, K75R, E83N, K378R |

| | | | | | | |
|---|---|---|---|---|---|---|
| **MP (M1)** | Node_185 – Seq_234 (10.0392) | (Sw)-sw-sw | 21.0835 | 0.0077 | $\omega_1$ = 0.00 (99%) $\omega_2$ = 85.3 (0.86%) | T184H, A202R |
| **MP (M2)** | - | - | - | - | - | - |
| **NS (NS1)** | Node_396 – Seq_045 (1.82011) | (Sw)-hu-hu | 31.7863 | 0.0001 | $\omega_1$ = 0.259 (99%) $\omega_2$ = 100000 (1.5%) | Q40L, L43P, R206H, P212S |
| | Node_280 – Seq_446 (2.25894) | (Sw)-sw-sw | 19.2982 | 0.0305 | $\omega_1$ = 0.204 (99%) $\omega_2$ = 5170 (1.0%) | S7I, T80N |
| **NS (NEP)** | Node_280 – Seq_446 (2.25894) | (Sw)-sw-sw | 20.8302 | 0.0124 | $\omega_1$ = 0.00 (99%) $\omega_2$ = 100000 (0.86%) | S7I |
| | Node_049 – Seq_238 (2.62767) | (Hu)-?-sw | 19.4866 | 0.0244 | $\omega_1$ = 10000000000 (100%) | D27G, G53R |

**Table S3**: Positions under positive selection identified by MEME analysis. For each protein, positions with evidence of positive selection at a test p-value threshold of ≤ 0.1 are listed together with the transmission type of branches tested in the corresponding analysis (sw = swine, hu = human). Specific mutations observed at the identified positions for the corresponding branch type are listed with number of observations in parentheses. Positions in all proteins are counted from the start codon.

| Segment (Protein) | Position | Transmission type | Mutations at position | Position | Transmission type | Mutations at position |
|---|---|---|---|---|---|---|
| **PB2** | 4 | sw-hu | I-V(1) | 423 | hu-sw | R-T(1) |
| | 6 | sw-sw | E-A(1), E-R(1) | 439 | hu-sw | Q-L(1) |
| | 19 | sw-sw | I-D(1) | 444 | sw-sw | V-I(4), I-V(1) |
| | 52 | sw-sw | A-Q(1) | 447 | hu-hu | Q-L(1) |
| | 58 | hu-hu | T-A(1) | 451 | sw-hu | I-V(2), I-L(1), V-I(1) |
| | 64 | sw-sw | M-I(22), M-V(6), M-T(3), M-L(2), M-R(2), I-M(1), I-V(1), M-K(1), M-N(1) | 468 | sw-sw | T-A(3), T-I(1) |
| | 92 | sw-sw | S-Q(1) | 478 | sw-sw | I-V(9), V-I(3) |
| | 105 | hu-hu | M-I(1), M-V(1), T-A(1), T-M(1), V-T(1) | 498 | hu-hu | I-D(1), I-N(1) |
| | 106 | hu-sw | T-A(3), A-S(1), T-S(1) | 525 | sw-sw | E-D(1), E-V(1) |
| | 110 | hu-hu | H-T(1) | 526 | sw-sw | K-R(10), K-F(1), R-K(1) |
| | 153 | sw-sw | D-G(2), D-H(2), D-N(1) | 535 | sw-sw | M-L(10) |
| | 184 | sw-sw | T-A(25), A-T(6), T-K(5), T-S(5), A-S(2), M-V(1), T-I(1), T-P(1) | 541 | sw-sw | G-P(1) |
| | 191 | hu-hu | E-K(2) | 560 | sw-hu | L-V(2) |
| | 191 | sw-sw | E-K(17), E-D(4), E-G(2), E-R(2), E-S(1) | 560 | sw-sw | V-I(9), V-L(5), L-V(2), L-M(1), V-M(1) |
| | 207 | sw-sw | L-M(1) | 588 | sw-sw | T-I(20), A-T(15), A-V(8), T-A(8), A-S(6), T-N(5), A-F(2), A-I(2), T-S(2), A-N(1), I-V(1), S-F(1), T-F(1), V-F(1) |
| | 227 | hu-hu | I-V(1), V-I(1) | 615 | hu-hu | I-V(2) |
| | 229 | sw-sw | I-V(5), I-N(1), I-Y(1), N-Y(1) | 650 | sw-sw | R-D(1) |
| | 241 | sw-sw | E-G(1) | 653 | sw-sw | S-E(1) |
| | 243 | sw-sw | M-L(6), M-I(3), M-K(3), I-V(1), K-M(1), M-R(1), M-T(1), M-V(1) | 654 | sw-sw | P-I(1), P-Q(1), P-S(1) |
| | 245 | sw-sw | T-I(1), T-N(1) | 656 | sw-sw | F-L(1), F-N(1) |

| | | | | | | |
|---|---|---|---|---|---|---|
| | 249 | hu-sw | E-G(2), E-N(1) | 666 | sw-sw | T-L(1), T-S(1) |
| | 267 | sw-sw | V-I(2), V-D(1) | 668 | sw-sw | L-T(1) |
| | 281 | sw-sw | L-I(1) | 669 | sw-sw | G-P(1), G-R(1) |
| | 293 | sw-sw | R-K(18), R-G(4), R-Q(2), G-E(1), R-M(1), R-N(1), R-W(1) | 670 | sw-sw | K-A(1) |
| | 295 | hu-sw | V-I(1), V-T(1) | 671 | sw-sw | D-K(1), D-V(1) |
| | 295 | sw-sw | V-I(22), V-M(13), V-T(3), I-T(1), I-V(1), T-V(1) | 676 | hu-sw | T-I(2), T-A(1), T-D(1), T-K(1), T-S(1) |
| | 300 | sw-hu | Q-L(1) | 676 | sw-sw | T-A(15), T-I(12), T-N(7), N-S(2), M-T(1), N-D(1), S-I(1), T-K(1), T-L(1), T-M(1), T-R(1), T-S(1), T-V(1) |
| | 311 | hu-hu | C-S(1) | 683 | sw-sw | T-K(9), T-R(2), T-A(1), T-M(1), T-N(1), T-Q(1), T-S(1) |
| | 344 | hu-sw | V-M(2) | 686 | sw-sw | V-I(5), I-V(1), M-I(1), V-M(1) |
| | 354 | hu-hu | I-L(5), I-V(1), L-I(1), L-R(1) | 704 | hu-hu | Y-H(1) |
| | 376 | hu-hu | K-G(1) | 729 | sw-sw | G-C(1) |
| | 383 | sw-sw | Q-L(2) | 750 | sw-sw | A-V(1) |
| | 389 | hu-hu | R-K(3) | 755 | sw-sw | R-Q(1) |
| | 391 | sw-sw | E-D(7), D-E(2), D-N(1), E-G(1), E-N(1) | 756 | sw-sw | L-M(2), M-L(2), M-V(2), L-T(1), M-T(1) |
| | 392 | hu-hu | Q-E(1) | 758 | sw-sw | I-T(6), I-N(1), I-R(1) |
| PB1 | 12 | sw-sw | V-I(7), V-E(1), V-L(1) | 475 | sw-sw | I-N(1), I-V(1) |
| | 15 | sw-sw | Q-G(1) | 484 | sw-sw | I-L(1), I-V(1) |
| | 41 | sw-sw | D-F(1) | 494 | sw-sw | S-A(1) |
| | 42 | sw-sw | T-L(1) | 508 | sw-sw | E-D(1), E-G(1) |
| | 67 | sw-sw | N-K(1) | 522 | sw-sw | D-G(1), D-N(1) |
| | 79 | hu-hu | P-A(1) | 523 | sw-sw | M-Q(1) |
| | 80 | sw-sw | N-S(2), S-C(1), S-I(1), S-T(1), T-N(1) | 524 | sw-sw | S-I(1), S-K(1) |
| | 85 | sw-sw | T-Q(1) | 546 | hu-hu | M-L(1) |
| | 105 | hu-hu | N-T(2) | 565 | sw-sw | D-G(1), D-H(1) |
| | 108 | sw-hu | L-I(1) | 567 | hu-sw | Q-L(1) |
| | 152 | hu-sw | S-L(2), S-M(1) | 578 | sw-sw | K-R(9), R-K(2), S-K(2), S-N(2), K-Q(1), R-G(1), R-S(1) |
| | 152 | sw-sw | L-S(4), S-L(4), M-T(3), L-M(2), K-M(1), L-F(1), M-V(1), S-A(1), S-T(1), T-E(1) | 581 | sw-sw | E-D(5), D-N(4), D-E(1), D-G(1), D-L(1) |
| | 164 | hu-sw | I-V(1) | 582 | sw-sw | E-D(1), Q-E(1), Q-H(1), Q-K(1) |
| | 172 | hu-sw | E-R(1) | 585 | sw-sw | S-A(3) |
| | 177 | hu-sw | E-K(1), E-R(1), E-V(1) | 587 | hu-hu | V-A(5), A-T(4), A-V(1), T-V(1) |
| | 196 | sw-sw | T-S(3) | 615 | sw-sw | L-M(1), L-V(1) |
| | 209 | hu-hu | K-R(2), K-N(1) | 620 | hu-hu | Y-Q(1) |
| | 281 | sw-sw | K-T(1) | 622 | hu-hu | G-T(1) |
| | 283 | sw-sw | A-P(1), A-V(1) | 623 | hu-hu | R-E(1) |
| | 327 | sw-sw | R-K(6), R-E(3), K-R(1) | 624 | hu-hu | L-V(1) |
| | 370 | hu-sw | A-G(1), A-T(1) | 636 | hu-sw | E-R(1) |
| | 374 | sw-hu | A-E(1), A-T(1), A-V(1) | 683 | sw-sw | L-I(2), L-S(1) |
| | 375 | sw-hu | D-E(1), D-S(1), G-S(1) | 686 | hu-hu | E-V(1) |
| | 384 | sw-sw | S-P(7), S-L(3), S-T(2), P-Q(1), P-T(1), S-A(1), S-I(1), S-Q(1) | 687 | hu-hu | Q-H(1), Q-P(1) |
| | 388 | sw-sw | K-R(5) | 702 | hu-hu | S-N(1) |
| | 431 | sw-sw | Y-H(2), Y-P(1) | 705 | hu-hu | Y-W(1) |
| | 433 | sw-hu | K-R(1) | 706 | hu-hu | R-N(1) |
| | 445 | sw-sw | D-P(1), D-S(1) | 746 | hu-sw | I-D(1), I-T(1) |
| | 446 | sw-sw | D-S(1) | 747 | hu-sw | C-L(1) |
| | 448 | sw-sw | A-V(5), A-C(1), A-T(1) | 753 | sw-sw | L-F(5), F-L(4), F-S(4), L-I(4), F-I(2), S-F(2), I-V(1) |
| | 451 | hu-hu | V-I(1) | 754 | sw-sw | R-G(6), R-K(5) |
| | 455 | sw-sw | N-D(10), N-S(3), D-N(2), D-G(1), M-L(1), N-M(1), N-Q(1) | 757 | hu-hu | R-K(1) |
| | 459 | sw-sw | I-V(3), I-A(1) | | | |

| | | | | | | |
|---|---|---|---|---|---|---|
| PA | 6 | sw-sw | R-E(1) | 261 | hu-sw | L-R(1), L-S(1), L-T(1) |
| | 17 | sw-sw | A-G(1) | 261 | sw-sw | L-F(2), L-S(2), L-T(1), L-V(1), L-W(1), S-T(1) |
| | 20 | sw-sw | A-T(16), A-S(4), T-A(3), A-V(2), A-E(1), T-K(1) | 263 | hu-sw | T-E(2), T-A(1) |
| | 42 | sw-sw | L-M(1), L-S(1), M-A(1) | 263 | sw-sw | T-K(7), E-K(6), T-E(5), T-A(4), E-G(2), K-R(2), T-N(2), E-D(1), N-D(1), N-K(1), N-S(1), T-S(1) |
| | 55 | hu-hu | D-N(2), N-D(2), N-V(1) | 272 | sw-sw | D-N(10), D-G(8), D-E(7), D-V(5), N-D(3), C-S(1), D-A(1), D-K(1), D-S(1), G-C(1), S-G(1), S-K(1) |
| | 56 | sw-sw | E-R(1) | 277 | hu-sw | H-S(2), F-S(1), H-P(1), S-Y(1) |
| | 65 | hu-hu | S-L(3), L-P(1) | 278 | sw-sw | Q-N(1) |
| | 65 | sw-sw | S-P(2), L-S(1), S-T(1), S-Y(1) | 279 | sw-sw | R-K(4) |
| | 70 | sw-sw | A-V(9), A-I(2), A-T(2), A-S(1) | 284 | sw-sw | L-M(4), L-R(1) |
| | 78 | hu-hu | I-A(1) | 329 | sw-sw | G-C(1), G-E(1) |
| | 79 | sw-sw | I-L(2), I-V(1) | 341 | hu-sw | I-V(1) |
| | 81 | sw-sw | G-A(1), G-V(1) | 341 | sw-sw | V-I(2), V-L(1) |
| | 83 | sw-sw | D-C(1), D-H(1) | 348 | hu-sw | I-L(1), I-T(1) |
| | 85 | sw-sw | T-A(15), T-N(9), T-I(4), I-T(3), I-V(3), N-S(2), I-A(1), T-G(1), T-S(1) | 357 | sw-sw | T-I(9), T-A(2), T-N(2), I-V(1), T-L(1), T-V(1), V-I(1) |
| | 87 | sw-sw | A-Y(1) | 399 | hu-hu | E-K(2) |
| | 105 | hu-hu | F-L(2) | 400 | hu-sw | P-F(1), P-L(1), P-S(1) |
| | 146 | sw-sw | H-D(1) | 421 | hu-sw | S-I(2), S-G(1) |
| | 156 | sw-sw | A-P(1) | 421 | sw-sw | S-I(3) |
| | 161 | hu-sw | Y-L(1) | 437 | hu-sw | H-I(1), H-Y(1) |
| | 169 | hu-sw | A-Y(1) | 450 | sw-sw | V-I(1) |
| | 170 | sw-sw | R-T(1) | 486 | sw-sw | I-L(4), I-M(1) |
| | 186 | hu-sw | S-N(3), S-A(2), S-G(1) | 489 | hu-hu | C-S(1) |
| | 186 | sw-sw | G-A(4), G-S(4), S-N(3), A-G(2), A-N(2), S-G(2), S-A(1) | 528 | sw-sw | V-I(3), T-I(1), T-N(1), T-V(1) |
| | 218 | sw-sw | S-A(1) | 538 | sw-sw | E-K(5), E-G(2), E-R(1), G-V(1) |
| | 224 | hu-hu | S-P(2) | 550 | hu-sw | L-I(1), L-R(1) |
| | 230 | sw-sw | R-T(2), R-K(1) | 555 | sw-sw | G-R(1) |
| | 231 | sw-sw | A-S(3), A-T(2), A-V(2), S-F(1) | 558 | sw-sw | S-L(2), A-T(1), S-A(1), S-E(1) |
| | 233 | sw-sw | V-I(2) | 598 | hu-hu | A-G(1) |
| | 238 | sw-sw | P-A(1), P-E(1) | 613 | hu-sw | E-D(1) |
| | 239 | sw-sw | N-T(1), N-W(1) | 616 | hu-sw | S-Q(1) |
| | 241 | hu-sw | C-Y(11) | 630 | sw-sw | E-D(1), E-H(1), E-K(1) |
| | 243 | hu-hu | E-D(1), E-P(1) | 667 | sw-sw | L-G(1) |
| | 258 | hu-hu | E-H(1) | 712 | sw-hu | V-T(1) |
| | 258 | sw-sw | D-E(7), E-D(3), D-Y(2), D-Q(1), D-S(1), E-A(1), E-K(1), E-Q(1), Y-F(1), Y-H(1) | 714 | hu-hu | A-T(1) |
| PA (PA-X) | 8 | sw-sw | C-H(1), C-R(1), C-S(1), C-Y(1) | 212 | sw-sw | A-V(9), I-T(7), V-I(7), A-E(4), V-A(4), I-V(3), A-T(2), I-A(1), I-L(1), I-R(1), S-I(1), V-G(1), V-L(1), V-T(1) |
| | 17 | sw-sw | A-G(1) | 213 | hu-hu | S-G(4), G-S(2), S-N(1) |
| | 42 | sw-sw | L-M(1), L-S(1), M-A(1) | 213 | hu-sw | S-G(7), G-D(1) |
| | 55 | hu-hu | D-N(2), N-D(2), N-V(1) | 213 | sw-hu | G-D(1) |
| | 56 | sw-sw | E-R(1) | 213 | sw-sw | G-D(11), G-S(11), S-G(8), S-N(3), G-V(2), D-S(1), G-C(1), G-N(1), S-D(1) |
| | 65 | hu-hu | S-L(3), L-P(1) | 214 | hu-hu | L-S(1), S-L(1) |
| | 65 | sw-sw | S-P(2), L-S(1), S-T(1), S-Y(1) | 214 | hu-sw | L-S(2) |
| | 70 | sw-sw | A-V(9), A-I(2), A-T(2), A-S(1) | 214 | sw-sw | L-S(5), L-W(1), S-L(1) |
| | 78 | hu-hu | I-A(1) | 216 | hu-hu | T-I(3), I-T(1) |

| | | | | | |
|---|---|---|---|---|---|
| 79 | sw-sw | I-L(2), I-V(1) | 216 | hu-sw | T-I(2) |
| 87 | sw-sw | A-Y(1) | 216 | sw-hu | I-T(1) |
| 151 | sw-sw | T-N(5), T-A(2), T-I(2), T-E(1) | 216 | sw-sw | T-I(30), I-T(3), T-A(1) |
| 161 | hu-sw | Y-L(1) | 217 | hu-sw | K-R(2) |
| 169 | hu-sw | A-Y(1) | 217 | sw-sw | K-R(13), K-T(1) |
| 186 | hu-sw | S-N(3), S-A(2), S-G(1) | 218 | hu-sw | V-A(3) |
| 186 | sw-sw | G-A(4), G-S(4), S-N(3), A-G(2), A-N(2), S-G(2), S-A(1) | 218 | sw-sw | V-A(29), A-P(1), A-V(1) |
| 193 | hu-hu | S-N(2) | 219 | hu-hu | S-F(1) |
| 193 | hu-sw | S-N(3) | 219 | hu-sw | S-F(4), S-Y(1) |
| 193 | sw-hu | S-N(1) | 219 | sw-hu | S-F(1) |
| 193 | sw-sw | S-N(17), S-C(1) | 219 | sw-sw | S-F(13), S-Y(11), S-C(3), F-C(1) |
| 194 | sw-sw | P-L(6), P-Q(6), Q-P(1) | 220 | hu-hu | R-H(2), H-P(1), H-R(1) |
| 195 | hu-hu | R-K(2), K-R(1) | 220 | hu-sw | H-L(3), H-R(3), R-H(2) |
| 195 | hu-sw | K-R(1) | 220 | sw-hu | R-H(1) |
| 195 | sw-sw | R-K(15), K-R(9), K-N(1) | 220 | sw-sw | R-H(34), R-L(9), H-R(4), H-P(1), H-Y(1), R-P(1) |
| 196 | hu-hu | E-K(2), E-G(1) | 221 | hu-hu | R-Q(5), Q-R(2) |
| 196 | hu-sw | E-K(2), E-G(1) | 221 | hu-sw | R-Q(2) |
| 196 | sw-hu | E-K(1) | 221 | sw-sw | R-Q(42), R-L(12), R-P(2), L-P(1), L-R(1), Q-P(1) |
| 196 | sw-sw | E-K(12), E-G(8), E-R(1) | 222 | hu-sw | T-I(1) |
| 197 | hu-hu | A-E(2), A-V(2) | 222 | sw-hu | T-I(1) |
| 197 | sw-sw | A-V(15), A-E(2), A-T(1) | 222 | sw-sw | T-I(24), T-A(3), T-N(1) |
| 198 | hu-sw | K-M(1), K-R(1) | 224 | sw-sw | P-L(11), P-Q(7), P-S(4) |
| 198 | sw-sw | K-R(13) | 226 | sw-hu | L-S(1) |
| 199 | hu-hu | R-K(5) | 226 | sw-sw | L-S(9) |
| 199 | hu-sw | R-K(6) | 227 | hu-hu | K-R(3), R-K(1) |
| 199 | sw-hu | I-T(1) | 227 | hu-sw | K-R(2), I-T(1) |
| 199 | sw-sw | R-K(28), R-I(5), I-T(2), I-R(1), K-R(1) | 227 | sw-sw | K-R(12), K-T(3), K-I(2), I-T(1), K-E(1), K-N(1), R-I(1), R-K(1), T-I(1), T-K(1) |
| 201 | sw-sw | L-S(6), L-P(1), L-W(1) | 228 | hu-hu | T-I(3), I-F(1), I-T(1) |
| 202 | hu-sw | K-E(1), K-R(1) | 228 | hu-sw | T-I(2), T-A(1) |
| 202 | sw-sw | K-R(7) | 228 | sw-sw | T-I(10), T-A(6), T-P(5), T-N(2), I-T(1) |
| 203 | hu-sw | K-R(2), K-E(1) | 229 | hu-hu | L-S(2), S-L(2) |
| 203 | sw-sw | K-R(11), K-N(1) | 229 | sw-sw | L-S(15), L-F(1) |
| 204 | hu-hu | D-N(4), N-S(2), D-G(1), N-D(1) | 230 | sw-hu | E-G(1) |
| 204 | hu-sw | N-S(3), D-N(2), N-D(1) | 230 | sw-sw | E-G(10), E-D(2), E-Q(2), E-K(1) |
| 204 | sw-hu | D-N(2) | 231 | hu-sw | P-H(2) |
| 204 | sw-sw | D-N(25), D-G(16), G-D(16), G-S(10), N-S(3), G-N(2), N-D(2), S-G(2), S-N(2), N-T(1) | 231 | sw-sw | P-H(8), P-L(5), P-S(3) |
| 205 | hu-sw | L-S(3) | 235 | sw-sw | D-G(1), G-D(1) |
| 205 | sw-sw | L-S(11) | 236 | sw-sw | S-L(12) |
| 206 | hu-hu | R-K(3) | 237 | sw-sw | N-S(7), N-K(1) |
| 206 | hu-sw | R-K(6) | 238 | sw-sw | R-Q(8), R-L(1) |
| 206 | sw-hu | K-R(1), R-K(1) | 240 | sw-sw | A-D(7), A-V(6) |
| 206 | sw-sw | K-R(14), R-K(12), K-N(2), K-Q(1), K-T(1) | 241 | hu-hu | A-V(1), T-A(1) |
| 207 | hu-hu | S-L(2), L-S(1) | 241 | sw-sw | A-T(6), A-V(3) |
| 207 | hu-sw | L-S(4), L-I(1), S-F(1) | 244 | sw-sw | A-E(3) |
| 207 | sw-sw | L-S(15), S-L(5) | 245 | sw-hu | S-N(1) |
| 208 | hu-hu | Q-K(1) | 245 | sw-sw | S-N(7) |
| 208 | hu-sw | Q-K(4), K-R(1), Q-R(1) | 246 | hu-hu | F-C(1) |
| 208 | sw-hu | R-E(1) | 246 | hu-sw | F-S(1) |
| 208 | sw-sw | R-K(20), Q-R(6), R-G(6), Q-K(4), R-Q(3), R-M(2), G-R(1), Q-L(1), Q-P(1) | 246 | sw-sw | F-S(3) |

| | | | | | | |
|---|---|---|---|---|---|---|
| | 209 | hu-hu | E-G(5), E-K(1) | 247 | hu-sw | L-F(1) |
| | 209 | hu-sw | E-G(4), E-V(1) | 247 | sw-sw | L-P(7), L-H(4) |
| | 209 | sw-sw | E-G(31), G-E(3), E-A(2), E-K(1) | 248 | sw-hu | K-R(1) |
| | 210 | hu-hu | L-P(2), P-Q(1), Q-L(1) | 248 | sw-sw | K-R(3) |
| | 210 | hu-sw | L-P(6), L-R(2), L-Q(1), P-L(1), Q-R(1) | 250 | hu-hu | P-Q(1) |
| | 210 | sw-hu | L-P(1), P-S(1) | 250 | sw-sw | P-Q(5), P-L(4), Q-P(2) |
| | 210 | sw-sw | P-Q(8), P-L(7), L-P(4), L-Q(3), R-Q(3), L-I(1), P-S(1) | 251 | sw-sw | K-R(6), E-K(1), K-E(1) |
| | 212 | hu-hu | A-V(2), A-E(1) | 252 | sw-sw | K-R(9), K-E(2), K-M(1), K-N(1) |
| | 212 | hu-sw | A-E(2), A-I(2), A-S(1), A-V(1) | | | |
| | 4 | hu-hu | K-R(2), R-K(2), I-T(1), K-N(1) | 211 | sw-sw | A-E(12), A-V(7), A-T(6), N-S(6), N-D(5), E-D(4), A-I(2), A-N(2), E-K(2), N-K(2), N-T(2), A-D(1), A-L(1), A-P(1), E-G(1), N-I(1), T-P(1) |
| | 4 | hu-sw | I-T(6), I-V(2), I-M(1), K-E(1), R-V(1) | 217 | sw-sw | V-A(1), V-I(1) |
| | 4 | sw-sw | K-R(11), I-T(6), I-V(6), K-E(6), R-K(5), I-M(4), K-I(2), K-N(2), K-T(2), V-A(2), A-I(1), A-T(1), I-L(1), K-Q(1), R-I(1), V-I(1) | 227 | sw-hu | K-V(1), T-A(1) |
| | 8 | sw-sw | L-I(3), L-M(3), L-Q(1), L-S(1) | 227 | sw-sw | T-I(18), T-K(8), K-E(5), K-T(5), T-M(4), T-N(3), I-T(2), K-N(2), K-R(2), E-A(1), E-K(1), I-V(1), K-I(1), K-M(1), M-V(1), T-A(1) |
| | 11 | sw-sw | A-T(26), A-V(21), T-A(16), T-I(8), A-S(4), A-I(2), T-S(2), V-A(2), I-M(1), T-K(1), V-I(1) | 228 | sw-sw | P-I(1), P-S(1), P-T(1) |
| | 13 | hu-hu | A-T(2), A-S(1), S-T(1), T-A(1), T-V(1) | 229 | sw-hu | E-I(1) |
| | 16 | hu-sw | N-D(4), N-S(2), Y-H(2), K-N(1), N-T(1), S-G(1) | 231 | sw-sw | V-A(12), A-T(11), V-I(10), A-V(9), T-I(7), A-S(3), I-V(3), V-G(3), I-M(2), T-A(2), A-G(1), A-I(1), A-R(1), N-Y(1), T-S(1), V-L(1), V-N(1) |
| **HA (H1)** | 38 | sw-sw | E-P(1) | 232 | hu-hu | I-T(4), A-E(1), E-K(1), I-K(1), I-R(1), K-A(1), K-R(1), T-I(1), T-P(1) |
| | 46 | sw-sw | S-R(1) | 232 | hu-sw | I-V(5), I-R(4), I-K(3), I-T(2), T-A(2), I-M(1), K-R(1), K-T(1), T-E(1), T-K(1), T-R(1), T-S(1) |
| | 51 | sw-sw | E-L(1), E-V(1) | 232 | sw-sw | A-T(35), K-R(13), T-A(13), A-V(10), A-D(6), T-K(6), A-S(5), T-I(4), A-E(3), I-K(3), A-N(2), A-P(2), E-K(2), I-R(2), T-N(2), A-I(1), E-G(1), I-T(1), I-V(1), K-N(1), K-Q(1), Q-R(1), R-I(1), R-K(1), S-P(1), T-R(1), T-V(1), V-I(1) |
| | 52 | hu-hu | D-N(2), N-S(2), D-A(1), N-T(1) | 234 | sw-sw | P-Q(6), P-S(6), P-T(5), P-L(2), P-W(1) |
| | 52 | hu-sw | D-N(3), N-T(3), D-G(2), D-E(1), D-S(1), N-D(1) | 236 | hu-sw | V-I(1), V-L(1), V-M(1), V-T(1) |
| | 52 | sw-hu | D-E(1), D-N(1), N-S(1) | 238 | sw-hu | D-G(1), E-K(1), G-D(1), G-N(1) |
| | 52 | sw-sw | N-T(27), D-N(21), N-S(18), N-D(6), D-E(2), D-G(2), D-S(1), D-Y(1), N-H(1), T-I(1) | 238 | sw-sw | D-N(29), D-G(20), E-G(17), E-K(11), G-D(9), K-N(6), N-D(5), E-D(4), G-E(3), G-N(3), E-Q(2), K-E(2), D-E(1), K-T(1), N-G(1) |
| | 55 | sw-sw | N-S(2), D-N(1), N-D(1), N-F(1) | 240 | hu-hu | A-S(1), A-T(1), E-A(1), E-H(1), S-P(1), T-P(1) |
| | 62 | sw-sw | R-K(11), K-R(7), G-R(6), N-K(4), K-N(3), N-D(3), N-S(3), G-E(2), K-G(1), K-S(1), R- | 248 | hu-sw | T-V(1) |

| | | | | | |
|---|---|---|---|---|---|
| | | E(1), R-G(1), R-I(1), R-M(1), R-Q(1), R-S(1) | | | |
| 65 | sw-sw | A-T(16), A-S(7), A-P(6), A-V(3), I-T(2), A-C(1), A-D(1), I-N(1), I-S(1), P-T(1), S-P(1), T-A(1), T-P(1), V-I(1) | 250 | sw-sw | V-I(8), I-V(6), V-L(2), L-F(1), L-M(1), L-V(1), V-C(1), V-M(1) |
| 69 | sw-sw | L-M(2), L-S(1), L-V(1) | 256 | sw-sw | I-L(1), I-Q(1) |
| 70 | sw-sw | G-R(11), G-E(5), G-A(1), G-K(1) | 268 | hu-hu | W-R(2), W-M(1) |
| 86 | hu-hu | S-P(4), L-S(3), S-L(1), S-T(1) | 270 | sw-sw | A-R(1) |
| 86 | sw-sw | L-S(27), S-P(10), S-L(8), L-F(7), L-M(7), L-P(5), F-S(4), L-Q(3), L-V(2), I-T(1), L-I(1), L-R(1), M-I(1), P-T(1), S-T(1) | 277 | hu-hu | A-V(1), F-P(1), F-V(1), S-F(1), S-P(1) |
| 100 | sw-sw | S-P(30), P-S(7), S-T(4), P-T(2), S-A(2), P-Q(1), S-L(1), S-R(1), S-Y(1) | 277 | hu-sw | A-T(6), A-V(3), A-D(2), A-S(2), F-L(1), S-Y(1) |
| 111 | sw-sw | Y-H(17), E-K(15), E-D(8), D-N(4), D-E(3), D-S(1), E-Q(1), E-Y(1), H-Y(1), S-I(1), Y-L(1) | 285 | sw-hu | D-S(1) |
| 113 | hu-hu | A-S(1), I-A(1) | 287 | hu-hu | P-S(3), S-P(2), P-G(1), Q-K(1) |
| 113 | sw-hu | T-E(1), T-N(1) | 292 | sw-sw | D-N(11), N-D(9), N-S(3), N-T(3), T-S(3), D-E(2), S-N(2), D-T(1), D-Y(1), N-C(1), N-Y(1), T-A(1), T-N(1) |
| 126 | sw-sw | S-T(5), S-A(3), S-L(2), S-F(1), S-V(1), S-Y(1), T-N(1) | 322 | hu-hu | Y-G(1) |
| 134 | sw-sw | F-T(1) | 325 | sw-sw | S-F(1), S-T(1) |
| 136 | sw-sw | K-N(21), K-E(10), K-R(9), E-G(2), I-M(2), K-M(2), I-R(1), K-G(1), K-I(1), K-S(1), K-T(1), R-I(1), R-K(1) | 326 | sw-sw | A-T(17), T-K(14), T-A(9), T-N(5), T-S(3), A-E(2), A-S(2), T-R(2), A-V(1), K-N(1), T-E(1), T-F(1), T-I(1), T-P(1) |
| 137 | hu-hu | A-T(1), T-A(1), T-E(1) | 332 | sw-sw | T-R(2), T-P(1) |
| 137 | sw-sw | A-T(32), A-E(22), T-A(11), A-S(7), A-V(5), E-G(4), T-I(4), E-K(3), K-E(3), E-R(2), G-E(2), A-D(1), A-K(1), A-L(1), E-D(1), R-G(1), T-K(1), T-V(1) | 333 | sw-sw | G-M(1) |
| 141 | sw-sw | P-S(7), P-L(4), P-T(3), K-E(1), P-A(1), P-I(1), P-K(1), P-Q(1), S-F(1) | 337 | hu-hu | I-T(1), V-T(1) |
| 144 | hu-hu | D-E(2), E-N(1), N-T(1), T-N(1) | 346 | sw-sw | F-L(2), F-R(1) |
| 144 | sw-sw | E-D(30), T-N(12), D-E(9), E-K(9), S-N(6), T-S(5), E-T(3), N-S(3), D-N(2), E-V(2), D-K(1), E-Q(1), N-T(1), S-G(1), S-T(1), T-E(1) | 347 | sw-sw | G-Y(1) |
| 145 | hu-sw | S-P(3), S-V(1), S-W(1), T-A(1), T-P(1), V-I(1) | 348 | sw-sw | A-G(1), A-I(1) |
| 146 | hu-hu | N-D(1), N-T(1), T-L(1), T-S(1) | 353 | hu-hu | I-L(1), I-V(1) |
| 147 | sw-sw | G-N(1) | 354 | hu-sw | E-H(1) |
| 148 | hu-sw | V-I(2), V-A(1), V-K(1), V-L(1), V-T(1) | 355 | hu-sw | G-R(1) |
| 148 | sw-sw | V-A(10), A-V(4), V-G(4), V-T(4), A-T(3), G-K(3), T-S(3), V-I(3), V-L(3), M-K(2), V-K(2), V-M(2), V-S(2), A-S(1), G-A(1), I-R(1), R-K(1), T-A(1), T-K(1), T-N(1), V-E(1) | 360 | sw-sw | M-L(5), M-V(5), M-E(3), M-K(1) |
| 150 | hu-sw | A-K(2), A-S(2), A-T(1) | 373 | hu-sw | Q-H(1), Q-S(1) |
| 150 | sw-sw | V-I(20), A-S(10), A-V(9), V-A(9), A-T(6), A-I(4), A-K(4), V-G(3), I-V(2), V-E(2), A-E(1), I-M(1), I-T(1), T-A(1), V-S(1) | 375 | hu-sw | S-G(1) |
| 152 | hu-hu | C-M(1) | 375 | sw-sw | S-T(3), S-A(1), S-H(1), S-Y(1), T-N(1) |
| 154 | sw-sw | Y-H(13), H-Y(11), H-R(10), H-N(6), H-Q(5), N-K(4), H-D(1), | 376 | hu-sw | G-K(1) |

| | | | | | |
|---|---|---|---|---|---|
| | | H-K(1), H-L(1), H-S(1), Q-K(1), Y-D(1), Y-F(1), Y-S(1) | | | |
| 155 | hu-hu | A-D(1), A-E(1), A-G(1), A-K(1), K-N(1) | 383 | sw-sw | S-T(2) |
| 155 | sw-sw | A-D(6), S-L(6), N-K(4), A-T(3), K-N(3), S-F(3), A-S(2), K-Q(2), N-D(2), S-A(2), S-Y(2), A-E(1), A-G(1), A-V(1), D-G(1), D-N(1), G-S(1), K-E(1), K-R(1), S-G(1), S-N(1), S-P(1), S-T(1), T-I(1), T-N(1) | 392 | sw-sw | S-N(22), S-R(13), N-K(2), S-T(2), S-E(1), T-I(1), T-S(1) |
| 157 | hu-hu | K-E(7), A-K(1), A-T(1), K-N(1), K-R(1), N-S(1) | 393 | sw-sw | N-T(7), N-F(1), N-S(1) |
| 157 | hu-sw | A-T(5), A-E(4), A-V(4), N-K(2), A-G(1), A-I(1), A-K(1), A-S(1), K-E(1), K-R(1) | 396 | sw-sw | N-D(1), N-T(1) |
| 157 | sw-sw | A-T(42), A-V(32), K-E(20), A-S(17), E-K(15), K-N(8), A-E(5), T-A(5), A-N(4), K-R(4), A-D(3), A-K(2), K-G(2), N-K(2), T-K(2), V-I(2), D-N(1), E-A(1), E-N(1), E-Q(1), I-V(1), K-Q(1), M-K(1), N-D(1), T-E(1), T-I(1), T-S(1), V-M(1) | 408 | sw-sw | A-S(6), S-A(4), A-T(3), A-V(3), A-F(1) |
| 158 | hu-hu | K-R(2), K-N(1), N-K(1), R-K(1), S-C(1), S-R(1) | 420 | hu-hu | I-L(2), I-M(2), I-V(1), M-L(1) |
| 158 | hu-sw | K-R(6), K-N(4), K-S(2), K-P(1), N-S(1), R-G(1), S-G(1), S-R(1) | 433 | sw-sw | D-N(1), D-S(1) |
| 158 | sw-sw | N-K(73), S-R(16), R-S(15), N-S(13), K-N(10), R-G(10), K-R(9), S-N(5), R-K(4), L-R(3), N-R(3), N-Q(2), R-N(2), G-R(1), K-G(1), K-P(1), N-G(1), N-P(1), N-T(1), R-Q(1), S-L(1), T-S(1) | 451 | hu-sw | L-Y(1) |
| 167 | sw-hu | L-I(1) | 460 | hu-hu | N-D(1), N-K(1), N-S(1), N-T(1) |
| 167 | sw-sw | I-V(5), I-L(4), L-I(4), L-P(2), I-T(1), L-N(1) | 466 | hu-sw | R-K(2), R-S(1) |
| 168 | hu-hu | V-I(2), I-T(1), V-G(1), V-T(1) | 467 | sw-sw | S-N(15), S-G(14), S-L(11), S-A(5), S-I(5), S-R(5), G-S(3), N-S(3), S-T(3), G-A(1), G-D(1), G-V(1), N-H(1), N-T(1) |
| 169 | sw-sw | G-V(7), R-K(7), R-G(5), V-E(5), G-E(4), G-R(4), K-Q(4), V-A(4), E-G(3), K-R(3), R-S(3), V-M(3), K-E(2), E-Q(1), K-D(1), K-S(1), M-T(1), V-K(1), V-T(1) | 470 | hu-sw | K-R(4), K-L(1), R-K(1) |
| 171 | hu-sw | G-E(12), E-N(1), G-I(1), G-K(1), G-R(1) | 476 | hu-sw | I-N(1), I-T(1) |
| 171 | sw-sw | G-E(76), G-R(19), N-D(11), G-K(10), E-G(5), E-D(3), A-E(2), D-G(2), E-K(2), G-A(2), G-D(2), G-V(2), N-G(2), D-A(1), D-K(1), D-S(1), E-A(1), E-Q(1), E-T(1), G-N(1), G-T(1), K-E(1), K-N(1), N-H(1), R-N(1) | 479 | sw-sw | G-R(1) |
| 172 | sw-sw | N-D(26), N-S(16), G-N(7), N-T(6), N-G(4), S-G(4), S-N(4), G-D(3), D-G(2), N-E(2), N-K(2), D-N(1), D-Y(1), G-E(1), G-S(1), G-Y(1), N-H(1), N-M(1), N-Q(1) | 496 | hu-sw | K-R(3), K-S(1), K-T(1) |
| 173 | hu-sw | S-L(4), S-V(1) | 500 | hu-hu | Y-K(1) |
| 177 | sw-sw | L-I(48), L-F(4), I-L(2), L-V(2), F-L(1), I-V(1), L-N(1) | 501 | hu-hu | D-K(1) |
| 178 | hu-hu | S-N(5), N-S(2), N-K(1), S-I(1), S-T(1) | 511 | hu-hu | L-I(1), L-K(1) |
| 179 | hu-sw | K-I(2), K-A(1), K-G(1), K-R(1), Q-K(1) | 517 | hu-sw | D-E(2), D-N(2), D-L(1) |

| | | | | | | |
|---|---|---|---|---|---|---|
| | 179 | sw-sw | K-N(25), K-I(7), K-Q(7), K-R(7), K-T(7), K-M(6), K-E(3), N-K(2), T-A(2), T-M(2), I-A(1), I-K(1), K-G(1), N-E(1), Q-K(1), T-K(1), T-V(1) | 523 | sw-sw | S-T(10), S-P(8), S-A(3), S-L(2), P-Q(1), P-S(1), S-F(1), S-V(1) |
| | 181 | sw-sw | Y-G(1), Y-H(1) | 525 | hu-sw | R-K(2), R-M(1), R-Q(1), R-T(1) |
| | 183 | sw-sw | N-T(1), N-V(1) | 528 | sw-sw | Q-R(5), R-Q(3), N-D(1), Q-H(1), Q-K(1), Q-L(1), Q-N(1), Q-P(1), R-N(1), R-W(1) |
| | 200 | sw-sw | T-N(18), N-D(2), N-K(2), N-R(2), T-S(2), T-K(1) | 534 | sw-sw | S-W(1) |
| | 201 | hu-hu | D-E(1), I-M(1), I-V(1), S-G(1), S-I(1), S-M(1), S-N(1), S-T(1), T-I(1) | 536 | hu-hu | V-A(3) |
| | 202 | hu-sw | A-T(4), A-D(3), A-E(2), A-N(2), T-N(2), A-G(1), E-G(1), G-V(1), T-A(1) | 543 | hu-sw | V-I(3), V-T(2), V-M(1) |
| | 202 | sw-sw | S-N(19), G-R(13), G-E(12), A-T(9), S-R(8), E-K(6), N-S(6), T-A(6), T-N(6), A-D(5), E-G(4), S-G(4), D-N(3), N-T(3), A-S(2), R-G(2), S-D(2), T-I(2), T-S(2), A-E(1), A-G(1), A-V(1), D-A(1), E-Q(1), E-R(1), N-D(1), R-Q(1), S-I(1), S-K(1), S-T(1), T-Q(1) | 544 | hu-sw | V-I(1) |
| | 203 | hu-hu | D-E(3), D-A(2), D-N(1), D-S(1), D-V(1) | 544 | sw-hu | V-I(1) |
| | 203 | hu-sw | D-E(2), D-N(2), D-S(2), D-V(2), S-I(1), S-T(1) | 544 | sw-sw | V-I(21), V-A(2) |
| | 205 | hu-hu | R-K(4), K-R(1), Q-E(1), Q-K(1), R-M(1), R-Q(1) | 549 | sw-sw | I-V(32), T-I(10), I-L(9), I-T(9), I-F(2), T-A(1), V-A(1) |
| | 205 | hu-sw | Q-R(8), Q-K(2), M-S(1) | 559 | sw-sw | L-T(1) |
| | 205 | sw-hu | Q-K(2), H-Y(1), R-K(1), R-Q(1) | 560 | hu-sw | Q-D(1) |
| | 205 | sw-sw | R-K(15), Q-R(13), Q-K(5), Q-L(4), R-M(4), Q-H(2), Q-Y(2), R-I(2), H-Y(1), K-R(1), L-I(1), Q-E(1), R-L(1) | 560 | sw-sw | Q-A(1), Q-H(1), Q-R(1) |
| | 206 | hu-sw | S-T(5), S-R(4), S-N(3), A-T(2), T-A(2), S-A(1), S-D(1), S-E(1), S-K(1) | 562 | sw-sw | R-K(15), R-Q(1), R-S(1), R-T(1), R-V(1) |
| | 206 | sw-sw | T-A(30), A-T(18), S-T(10), S-A(5), T-S(5), R-W(4), A-V(3), S-N(2), S-R(2), S-W(2), R-Q(1), S-G(1), S-I(1), S-K(1), S-V(1), S-Y(1), T-D(1), T-I(1), T-P(1), V-A(1), W-Q(1) | | | |
| **HA (H3)** | 2 | hu-sw | K-R(1), K-S(1) | 205 | sw-sw | R-K(13), K-N(8), K-R(6), K-M(3), R-G(3), S-N(3), S-R(3), K-A(2), K-E(2), K-T(2), R-S(2), K-G(1), K-I(1), K-S(1), N-S(1), Q-K(1), R-M(1), R-N(1), S-G(1) |
| | 4 | hu-sw | I-V(1) | 208 | hu-hu | I-F(1), I-V(1), T-I(1) |
| | 4 | sw-sw | I-V(3), I-T(2), V-I(2), I-A(1), V-L(1) | 208 | sw-sw | T-I(17), I-T(4), I-V(2), T-A(2), T-N(2), V-I(1) |
| | 8 | sw-hu | S-G(1) | 209 | hu-hu | F-S(1), N-D(1), N-S(1), S-F(1), S-N(1) |
| | 9 | sw-hu | Y-H(2), C-Y(1) | 209 | sw-sw | N-S(16), S-N(7), N-Y(6), Y-S(3), F-S(2), S-A(2), S-Y(2), D-N(1), N-D(1), N-K(1), Y-N(1) |
| | 10 | sw-sw | I-T(2), I-A(1), I-F(1), I-G(1), I-V(1), T-I(1), V-I(1), V-L(1) | 213 | hu-hu | Q-R(2), R-Q(1) |
| | 13 | hu-hu | L-Q(3) | 215 | hu-hu | S-I(1) |
| | 13 | sw-sw | L-M(5), L-Q(4), L-I(1), L-P(1), L-S(1), M-V(1), Q-H(1) | 215 | hu-sw | S-A(1), S-L(1) |
| | 14 | sw-sw | V-G(8), V-A(6), V-I(6), I-V(5), I-T(4), I-N(3), I-S(1), T-A(1), V-F(1), V-T(1) | 215 | sw-sw | S-P(12), P-Q(2), S-T(2), A-T(1), P-S(1) |

| | | | | | |
|---|---|---|---|---|---|
| 15 | sw-hu | F-L(1), F-Y(1) | 219 | sw-sw | T-I(12), I-T(3), T-A(2), T-V(2), I-L(1), I-M(1), I-V(1), T-M(1) |
| 16 | sw-sw | G-S(4), A-T(3), A-D(1), A-G(1), A-P(1), A-R(1), A-S(1) | 237 | sw-sw | P-I(1), P-L(1) |
| 21 | hu-sw | G-E(2), G-R(2), G-K(1) | 242 | hu-hu | I-V(2), L-Q(2), V-I(2), L-I(1) |
| 23 | sw-sw | D-E(7), D-G(7), D-N(3), G-D(3), D-S(2), D-T(1), G-S(1), G-Y(1) | 251 | sw-sw | T-D(1) |
| 24 | hu-sw | N-D(3), N-K(2) | 252 | sw-sw | I-V(3), I-T(2), I-L(1), I-M(1) |
| 26 | sw-sw | M-R(2), M-T(2), T-K(2), T-M(2), M-L(1), T-S(1) | 260 | hu-sw | L-M(1), L-N(1), V-I(1), V-M(1) |
| 28 | sw-sw | T-M(4), T-G(1), T-I(1), T-K(1) | 262 | sw-sw | N-K(6), N-S(3), N-T(2), N-E(1), N-V(1) |
| 36 | sw-sw | V-W(1) | 264 | sw-sw | N-T(1), T-N(1), T-Q(1) |
| 47 | hu-sw | N-D(4), N-S(2) | 269 | sw-sw | A-P(1) |
| 55 | hu-sw | A-G(1) | 278 | sw-sw | S-N(14), S-G(8), N-D(2), S-I(1), S-R(1), S-T(1), T-A(1), T-N(1), T-S(1) |
| 66 | hu-hu | E-K(2), G-E(2), K-R(1), R-G(1) | 291 | hu-hu | D-G(3), G-D(3) |
| 66 | sw-hu | R-G(1) | 292 | sw-hu | N-E(1) |
| 71 | sw-sw | P-I(1) | 294 | sw-sw | N-H(3), N-S(3), S-N(3), N-D(2), N-Y(2), N-E(1), N-I(1), N-K(1), N-T(1) |
| 78 | hu-sw | K-E(2), E-K(1), I-M(1), I-T(1), I-V(1), K-G(1), K-R(1) | 320 | hu-sw | A-D(3), A-T(1) |
| 78 | sw-sw | K-E(20), E-G(4), K-R(3), E-K(1), G-E(1), K-G(1), K-N(1), T-A(1) | 320 | sw-sw | A-P(6), A-T(4), A-V(3), A-E(1), A-N(1), D-A(1), D-N(1), P-A(1), T-N(1) |
| 94 | sw-sw | G-D(10), D-E(8), D-G(6), G-S(4), D-N(3), D-S(1), G-A(1), G-E(1), G-M(1) | 323 | sw-sw | R-K(4), K-H(1) |
| 117 | sw-sw | Y-F(4), D-E(3), D-Y(3), Y-H(2) | 363 | hu-sw | V-M(1) |
| 121 | sw-hu | Y-H(1) | 370 | sw-sw | K-R(1) |
| 128 | sw-sw | V-I(13), I-V(4), V-A(1) | 388 | sw-sw | A-T(8), A-S(6), A-E(2), T-S(1) |
| 138 | hu-hu | N-K(2), T-N(1) | 399 | sw-sw | R-K(4), R-W(1) |
| 138 | hu-sw | N-S(2), N-D(1), N-T(1), T-I(1) | 438 | hu-hu | S-G(1) |
| 146 | sw-sw | V-I(3), V-L(1) | 441 | hu-hu | A-F(1), F-C(1) |
| 147 | sw-sw | A-T(10), A-S(4), A-N(3), K-E(2), T-N(2), A-D(1), A-K(1), T-A(1), T-K(1), T-S(1) | 443 | sw-hu | L-I(1) |
| 151 | hu-hu | G-K(1), K-G(1), K-T(1), T-K(1) | 450 | hu-hu | Q-Y(1) |
| 151 | hu-sw | K-S(1), K-T(1), T-A(1), T-K(1) | 453 | hu-hu | I-A(1), I-W(1) |
| 151 | sw-sw | T-A(4), S-A(3), T-S(2), A-T(1), K-E(1), K-R(1), S-T(1), T-E(1), T-K(1) | 457 | hu-hu | D-A(1) |
| 153 | sw-sw | Y-F(9), Y-N(7), N-S(3), Y-R(3), Y-S(3), N-I(2), S-F(2), S-Y(2), Y-Q(2), F-S(1), N-D(1), N-R(1), R-F(1), S-P(1), Y-I(1), Y-T(1) | 461 | hu-hu | I-F(1), I-V(1), N-D(1), N-I(1) |
| 156 | hu-hu | K-I(3), I-K(1), K-R(1) | 466 | hu-hu | R-K(2), K-R(1) |
| 158 | hu-hu | R-G(2), R-K(2), G-R(1) | 469 | hu-hu | R-K(1) |
| 158 | sw-sw | E-G(12), E-K(7), G-E(5), K-N(5), G-K(3), G-R(3), R-G(3), K-E(2), K-G(2), E-A(1), E-N(1), G-N(1), K-Q(1), K-R(1), R-K(1) | 470 | sw-sw | Q-L(7), Q-N(1) |
| 160 | hu-hu | D-G(3), N-D(2), G-S(1), G-V(1), I-N(1), N-S(1), N-T(1), V-G(1), V-I(1) | 471 | sw-sw | L-R(1) |
| 160 | sw-sw | V-D(11), V-A(9), D-N(6), D-G(4), D-E(3), D-V(3), N-K(3), N-S(2), V-I(2), V-N(2), V-T(2), A-D(1), D-A(1), G-D(1), G-V(1), I-T(1), N-D(1), N-Q(1), S-G(1), S-T(1), V-G(1), V-K(1) | 476 | sw-sw | E-A(1), E-V(1) |
| 161 | sw-sw | N-K(28), K-N(11), N-S(3), S-N(2), K-I(1), K-M(1), K-S(1), N-Y(1), S-K(1) | 515 | sw-sw | R-V(1) |
| 171 | hu-hu | Y-H(2), H-T(1), T-Y(1) | 537 | hu-sw | A-S(2) |

| | | | | | | |
|---|---|---|---|---|---|---|
| | 172 | hu-hu | E-K(2), H-S(1), K-E(1), K-Q(1), Q-H(1) | 537 | sw-sw | A-R(1), A-S(1), A-T(1) |
| | 172 | sw-sw | N-K(12), N-H(10), N-S(9), K-Q(6), K-E(3), E-A(2), E-K(2), K-N(2), A-G(1), E-G(1), E-Q(1), H-Q(1), H-R(1), H-S(1), N-A(1), Q-H(1), S-G(1) | 545 | sw-sw | V-A(12), A-T(3), V-G(3), V-I(2), A-H(1), A-V(1), V-F(1), V-T(1) |
| | 175 | hu-hu | F-Y(2), F-S(1), S-N(1), S-Y(1), Y-F(1), Y-N(1) | 554 | sw-sw | A-T(2), A-G(1), A-P(1) |
| | 175 | sw-sw | Y-H(17), Y-N(7), Y-D(3), F-S(2), H-N(2), N-H(2), N-Y(2), F-H(1), F-I(1), F-Y(1), N-D(1), N-G(1), S-N(1), Y-F(1), Y-G(1), Y-S(1), Y-T(1) | 560 | hu-hu | I-K(1), I-L(1) |
| | 189 | sw-hu | E-G(1), K-Q(1) | 565 | sw-sw | C-H(3) |
| | 205 | hu-hu | S-N(2), K-R(1), N-K(1), Q-K(1), R-S(1) | 566 | sw-sw | V-I(6), I-G(2), I-V(1) |
| **NP** | 8 | sw-sw | R-S(1) | 251 | sw-sw | A-V(1) |
| | 31 | sw-sw | K-R(8), R-K(4), G-R(1), K-P(1), R-G(1) | 253 | hu-hu | F-I(1), F-L(1), I-D(1), I-F(1) |
| | 34 | hu-sw | G-S(2), G-D(1) | 310 | hu-sw | S-T(1) |
| | 50 | sw-sw | S-N(6), S-D(1), S-G(1), S-R(1) | 339 | sw-sw | E-L(1) |
| | 53 | sw-hu | E-D(1) | 353 | hu-hu | I-L(2), V-I(2), I-V(1), S-I(1), V-L(1) |
| | 55 | hu-hu | R-V(1) | 370 | sw-sw | N-S(6), N-T(2), N-A(1), S-N(1) |
| | 61 | hu-hu | I-L(2), I-T(1), I-V(1), L-I(1) | 373 | hu-sw | A-T(4), A-V(2), T-A(1) |
| | 66 | hu-hu | M-P(1) | 430 | hu-hu | S-A(1), T-N(1) |
| | 67 | hu-hu | V-N(1) | 431 | hu-sw | G-M(1) |
| | 113 | sw-sw | K-L(1) | 450 | hu-hu | G-S(7), S-N(3), S-G(1) |
| | 122 | sw-sw | Q-L(2), Q-H(1) | 455 | hu-hu | D-E(2), D-T(1) |
| | 124 | sw-sw | N-Y(1) | 456 | hu-sw | V-M(1) |
| | 126 | hu-hu | G-P(1) | 459 | hu-hu | Q-R(2), R-Q(1) |
| | 126 | sw-sw | G-S(2), G-N(1) | 459 | sw-sw | Q-H(2), Q-N(1) |
| | 130 | hu-sw | T-I(1) | 470 | hu-hu | K-R(2) |
| | 131 | hu-sw | A-R(1) | 470 | hu-sw | K-R(2) |
| | 131 | sw-sw | A-G(1), A-P(1), A-R(1) | 472 | hu-hu | A-T(5), T-A(4) |
| | 178 | sw-sw | A-R(1) | 472 | hu-sw | T-A(3), A-T(1) |
| | 190 | hu-sw | A-N(1), A-T(1), A-V(1) | 472 | sw-sw | T-A(3) |
| | 203 | sw-sw | D-G(1), D-S(1) | 473 | hu-hu | N-S(1), S-I(1), S-N(1) |
| | 213 | sw-sw | R-P(1) | 473 | hu-sw | I-T(1), S-N(1) |
| | 217 | hu-hu | I-V(2), S-G(2), G-S(1), I-S(1), V-A(1), V-I(1) | 473 | sw-sw | S-N(10), N-T(4), N-S(2), N-K(1), S-T(1) |
| | 218 | sw-sw | A-S(1), A-V(1) | 480 | sw-sw | D-E(1), D-N(1) |
| | 222 | hu-sw | M-I(1), M-L(1), M-T(1) | 483 | sw-sw | N-Y(1) |
| | 239 | sw-sw | M-V(6), M-E(1) | 485 | hu-sw | G-C(1) |
| | 240 | sw-sw | D-S(1) | 494 | hu-sw | E-V(1) |
| | 241 | sw-sw | Q-P(1) | 495 | hu-sw | E-V(1) |
| | 246 | sw-sw | R-P(1) | | | |
| **NA (N1)** | 3 | sw-sw | P-S(6), T-A(4), T-P(2), A-T(1), P-D(1), P-Q(1) | 331 | sw-sw | R-G(26), K-G(8), K-N(8), K-R(7), R-K(6), K-E(3), K-M(3), E-G(1), G-K(1), G-R(1), K-Q(1), N-K(1), R-V(1) |
| | 16 | hu-sw | T-A(4), I-A(1), I-T(1), V-I(1) | 332 | sw-sw | T-I(7), T-K(7), K-R(4), T-A(3), T-M(3), T-P(2), E-T(1), K-E(1), T-V(1) |
| | 16 | sw-hu | A-T(1), T-A(1), T-I(1) | 336 | hu-sw | N-G(1) |
| | 16 | sw-sw | T-I(20), T-A(17), I-T(5), I-V(4), A-T(2), I-L(2), I-M(2), A-V(1) | 340 | sw-sw | S-F(11), S-T(7), S-L(6), S-P(4), S-Y(4), P-Q(2), P-S(2), S-A(2), F-L(1), L-A(1), L-R(1), P-H(1), P-L(1), S-V(1), T-I(1), T-S(1), V-I(1), V-S(1) |
| | 18 | sw-sw | G-A(1), G-W(1) | 348 | hu-hu | G-L(1) |
| | 20 | hu-sw | I-T(2), I-V(1), V-I(1) | 364 | sw-sw | S-N(6), S-D(3), S-G(2), S-K(1) |

| | | | | | | |
|---|---|---|---|---|---|---|
| | 39 | sw-sw | Q-R(2), Q-F(1), Q-H(1), Q-P(1) | 368 | hu-hu | R-A(1) |
| | 48 | hu-hu | I-V(2), I-T(1), M-V(1), V-M(1) | 373 | hu-hu | M-S(1) |
| | 48 | hu-sw | T-K(3), I-T(1), I-V(1), M-I(1), T-S(1) | 374 | hu-hu | I-K(1), I-S(1), I-V(1) |
| | 48 | sw-sw | T-A(12), I-T(8), T-I(8), T-K(5), T-P(5), I-M(3), V-I(3), I-L(2), I-V(2), M-I(2), I-K(1), I-S(1), M-L(1), M-R(1), M-V(1), P-S(1), T-R(1), V-A(1), V-M(1) | 375 | hu-hu | W-M(1) |
| | 53 | sw-sw | V-I(35), V-F(8), V-A(5), I-V(3), I-T(2), V-G(2), V-T(2), F-L(1), F-S(1), I-M(1), T-I(1) | 382 | hu-hu | D-N(1), G-M(1), K-R(1), R-G(1) |
| | 59 | hu-hu | K-R(1), N-K(1), S-N(1) | 385 | hu-sw | N-S(1), S-W(1) |
| | 61 | sw-sw | W-C(1), W-L(1), W-Y(1) | 386 | hu-sw | D-N(2), K-E(1), N-D(1), N-S(1), S-I(1) |
| | 81 | sw-sw | V-I(25), V-A(24), V-T(5), T-A(4), I-S(3), I-T(3), I-V(3), T-I(3), A-S(2), A-V(2), V-F(2), I-N(1), N-D(1), N-Y(1), T-N(1), T-P(1), T-V(1), V-D(1), V-G(1), V-N(1), Y-H(1) | 387 | hu-sw | F-R(1), F-Y(1) |
| | 84 | sw-hu | I-V(2), N-S(1), T-A(1), T-V(1), V-I(1) | 388 | hu-sw | S-Q(1) |
| | 84 | sw-sw | I-T(4), I-V(4), K-E(4), K-R(3), I-K(2), K-N(2), E-V(1), I-F(1), I-M(1), K-S(1), K-T(1), M-I(1), R-K(1), T-K(1) | 389 | hu-sw | M-L(2), I-T(1), I-V(1), M-Q(1), M-V(1), V-I(1) |
| | 98 | hu-hu | A-F(1), A-P(1) | 399 | hu-hu | W-Y(1) |
| | 101 | sw-sw | S-R(2), S-N(1), S-T(1), T-S(1) | 416 | hu-hu | D-N(2), D-G(1) |
| | 147 | sw-sw | G-T(1) | 416 | sw-sw | D-N(19), D-S(9), N-D(9), N-S(3), S-N(3), D-G(2), G-D(1), N-K(1), N-T(1), S-K(1) |
| | 149 | hu-sw | I-V(2), I-F(1), I-Y(1), V-I(1) | 424 | hu-sw | V-I(1) |
| | 157 | sw-hu | A-R(1) | 430 | sw-hu | R-E(1) |
| | 164 | sw-sw | G-E(1), G-R(1) | 432 | hu-hu | E-K(1), E-N(1), K-E(1), R-K(1) |
| | 213 | sw-sw | I-M(1), T-I(1), T-V(1) | 440 | sw-sw | G-A(3), G-S(2) |
| | 215 | hu-sw | T-I(1), T-S(1) | 444 | hu-hu | S-R(1) |
| | 231 | sw-sw | C-I(1) | 451 | hu-hu | D-G(2), D-A(1), D-N(1), E-D(1), G-E(1) |
| | 240 | sw-sw | T-A(3), T-I(1), T-V(1), V-S(1) | 454 | hu-hu | G-C(1), G-F(1), G-R(1), G-S(1), N-D(1) |
| | 248 | hu-hu | D-N(1), N-D(1) | 459 | hu-hu | D-R(1) |
| | 250 | sw-hu | P-Q(1) | 460 | hu-hu | G-A(1), G-L(1) |
| | 250 | sw-sw | A-P(1), A-Q(1), P-A(1), Q-P(1), Q-R(1) | 461 | hu-hu | A-W(1) |
| | 273 | hu-sw | N-G(1) | 462 | sw-hu | D-N(1) |
| | 276 | sw-sw | Y-P(1) | 464 | hu-hu | P-L(1) |
| | 278 | sw-sw | E-G(1), E-S(1) | 466 | hu-hu | T-A(1), T-I(1), T-N(1), T-S(1), T-V(1) |
| | 314 | sw-sw | I-M(8), I-L(5), I-K(1), I-V(1) | 466 | sw-hu | T-D(1) |
| | 317 | sw-sw | I-V(5) | 467 | sw-sw | I-V(10), I-D(1), I-M(1), I-Q(1), I-T(1), I-Y(1), V-I(1) |
| | 329 | hu-sw | K-R(2), N-D(2), N-S(2), K-E(1), N-I(1) | | | |
| **NA (N2)** | 4 | sw-hu | S-I(1) | 343 | sw-sw | E-K(1), E-Q(1), E-S(1) |
| | 16 | sw-hu | I-A(1) | 344 | hu-hu | E-K(3), K-E(1), K-R(1), R-K(1) |
| | 19 | sw-hu | T-I(1), T-L(1) | 346 | hu-hu | G-D(1), G-S(1), G-Y(1), N-S(1), N-T(1), S-G(1) |
| | 42 | sw-sw | C-Y(21), C-R(7), C-H(2), C-S(2), Y-H(2), C-F(1), F-L(1), S-G(1), S-H(1), S-N(1), Y-C(1) | 346 | sw-sw | G-S(11), G-D(8), N-G(7), N-S(7), S-N(4), G-A(2), G-E(2), N-D(2), S-G(2), E-G(1), E-K(1), G-N(1), G-V(1), N-I(1), N-T(1) |
| | 46 | hu-hu | A-P(2), P-A(2) | 350 | sw-sw | K-I(1), K-N(1) |
| | 52 | sw-hu | L-Q(1), S-L(1) | 352 | hu-hu | W-F(1) |

| | | | | | |
|---|---|---|---|---|---|
| 63 | sw-hu | T-V(1) | 353 | hu-hu | A-L(1) |
| 69 | sw-sw | T-N(14), T-A(10), A-T(6), T-I(4), A-V(3), S-F(3), T-S(3), A-F(1), A-I(1), A-S(1), S-A(1), T-V(1) | 354 | hu-hu | F-S(1) |
| 71 | sw-sw | T-I(24), T-A(4), T-N(4), T-S(2), A-T(1), I-T(1), I-V(1), T-K(1), T-L(1), T-P(1), T-V(1) | 355 | hu-hu | D-P(1) |
| 74 | sw-sw | E-K(10), E-V(4), E-D(2), E-G(1), E-Y(1) | 360 | hu-hu | V-L(1) |
| 77 | hu-sw | I-K(4), I-T(4), I-V(2) | 360 | sw-sw | V-L(9), I-V(4), V-M(4), I-L(2), I-T(2), T-I(2), V-A(2), V-I(2), I-M(1), M-I(1), V-S(1) |
| 77 | sw-sw | I-V(22), I-T(16), I-M(14), I-K(10), V-I(8), V-A(5), I-L(2), T-I(2), V-L(2), I-A(1), K-I(1), K-N(1), K-R(1), T-A(1), V-M(1), V-T(1) | 362 | hu-hu | M-S(1) |
| 79 | sw-sw | P-S(13), P-H(3), P-L(2), P-D(1), P-T(1), S-C(1), S-P(1) | 368 | sw-sw | E-D(5), K-N(4), E-K(2), K-E(2), N-E(2), N-K(2), D-N(1), E-N(1), E-T(1), K-G(1), N-S(1) |
| 81 | hu-hu | P-V(2), L-P(1), V-L(1) | 369 | hu-hu | D-E(1), E-K(1), K-E(1), K-R(1), K-T(1), T-D(1), T-K(1) |
| 81 | hu-sw | L-P(2), L-I(1), L-Q(1), L-S(1), P-S(1), P-T(1) | 370 | hu-hu | L-S(4), S-L(2), L-F(1) |
| 86 | sw-sw | N-S(25), N-D(19), N-T(6), S-N(5), N-K(4), N-H(2), N-I(2), D-E(1), D-N(1), K-R(1), S-R(1) | 370 | sw-sw | L-S(36), L-F(9), S-L(6), L-V(5), F-L(2), L-I(2), S-A(2), S-T(2), L-Q(1), S-P(1), S-V(1), V-A(1), V-I(1) |
| 93 | hu-hu | D-N(2), D-G(1), G-D(1), G-Q(1), G-S(1), K-N(1), N-D(1), Q-D(1), Q-K(1) | 372 | hu-hu | S-L(3), L-F(2), L-S(1) |
| 121 | sw-sw | Y-L(1) | 381 | hu-hu | E-G(3), G-E(1) |
| 143 | hu-hu | G-V(1), K-R(1), R-G(1), V-I(1), V-K(1), V-M(1) | 385 | hu-hu | N-T(2), K-N(1), T-K(1) |
| 149 | sw-sw | V-I(10), I-V(4), I-M(2), V-A(2), A-T(1), F-V(1), I-F(1), I-T(1), V-F(1), V-L(1) | 390 | sw-hu | L-V(1), S-T(1) |
| 156 | sw-sw | R-I(1) | 391 | sw-sw | Q-W(1) |
| 199 | hu-sw | E-K(4), K-N(3), K-E(2), E-A(1), E-T(1), K-Q(1), K-R(1), N-K(1) | 392 | hu-hu | I-T(3), I-M(2) |
| 199 | sw-hu | D-N(1), E-G(1), K-N(1), K-T(1) | 398 | hu-hu | V-R(1) |
| 199 | sw-sw | K-N(31), E-K(21), E-G(10), K-E(10), K-R(9), K-T(7), K-G(5), E-A(3), E-V(3), N-D(3), E-N(2), K-A(2), E-D(1), E-Q(1), E-R(1), K-D(1), K-I(1), K-Q(1), N-E(1), N-T(1), R-G(1), T-K(1), V-A(1) | 403 | hu-hu | R-A(1), R-K(1) |
| 205 | hu-sw | F-L(1), I-D(1) | 404 | hu-hu | S-N(1) |
| 220 | hu-hu | Q-K(2), K-N(1), K-P(1), K-Q(1) | 407 | hu-hu | S-F(1), S-L(1) |
| 220 | sw-sw | Q-K(8), Q-H(7), K-N(4), K-Q(4), Q-R(4), K-H(2), K-R(2), K-G(1), Q-L(1), Q-M(1), R-M(1) | 411 | sw-sw | S-T(5), S-C(1) |
| 237 | sw-sw | C-A(1) | 414 | sw-sw | G-S(12), G-D(9), G-N(7), S-N(7), D-N(2), D-G(1), G-R(1), G-V(1), S-G(1), V-L(1) |
| 253 | sw-hu | K-R(3) | 431 | hu-hu | E-K(2), K-N(2), E-R(1) |
| 267 | hu-hu | K-P(2), P-Q(2), P-T(2), K-T(1), P-L(1), P-S(1), Q-P(1), T-A(1), T-K(1), T-P(1) | 432 | hu-hu | E-Q(2), Q-E(2) |
| 269 | sw-sw | L-S(8), S-L(8), S-T(4), A-S(2), A-T(2), I-M(2), L-F(2), L-V(2), S-A(2), A-L(1), A-V(1), I-L(1), I-T(1), I-V(1), L-I(1), L-K(1), L-M(1), S-M(1), S-P(1), T-A(1) | 432 | sw-sw | Q-E(12), Q-K(4), E-K(3), Q-R(3), E-S(2), P-S(2), S-T(2), E-D(1), E-G(1), P-Q(1), Q-H(1), Q-L(1), Q-P(1), Q-S(1), Q-W(1), S-E(1), S-P(1), S-R(1) |
| 280 | sw-sw | C-V(1) | 435 | hu-hu | E-R(2), E-K(1), K-E(1), R-K(1) |

| | | | | | | |
|---|---|---|---|---|---|---|
| | 313 | hu-hu | A-V(1), D-A(1), V-D(1), V-T(1) | 437 | hu-hu | L-W(2), W-L(1) |
| | 313 | hu-sw | V-A(4), D-N(2), V-D(1) | 464 | hu-hu | I-L(2), L-I(1), L-M(1) |
| | 313 | sw-sw | V-A(27), D-N(20), V-I(11), D-G(5), G-D(5), G-S(5), V-F(4), A-T(3), D-E(2), G-N(2), I-T(2), S-G(2), V-G(2), A-N(1), A-V(1), D-K(1), D-S(1), D-T(1), D-Y(1), G-V(1), I-M(1), S-N(1), V-D(1), V-N(1) | 465 | sw-hu | N-D(1) |
| | 322 | sw-sw | V-T(2) | 468 | hu-hu | H-L(3), P-H(2), H-P(1), H-R(1), P-L(1) |
| | 332 | sw-sw | F-S(20), S-F(17), F-L(13), S-Y(11), S-T(7), F-V(2), S-L(2), H-Q(1), L-F(1), S-A(1), Y-H(1) | 469 | hu-hu | I-V(2) |
| | 336 | hu-hu | H-N(2), Y-H(2), H-P(1), H-R(1), N-Y(1) | 469 | hu-sw | I-M(1), V-I(1) |
| | 338 | hu-hu | R-Q(3), L-F(2), L-I(1), L-K(1), L-R(1), R-L(1) | 469 | sw-sw | I-V(26), V-I(8), I-M(6), I-L(1), I-T(1), V-A(1) |
| | 343 | hu-hu | E-D(1), E-I(1) | | | |
| **MP (M1)** | 9 | hu-sw | T-M(1) | 88 | sw-sw | G-A(1) |
| | 20 | sw-sw | F-V(1), L-F(1) | 96 | sw-sw | A-G(2), A-S(1) |
| | 28 | hu-hu | L-Q(1) | 144 | sw-sw | F-P(1) |
| | 30 | hu-hu | D-N(4), G-S(2), D-S(1), D-Y(1), S-D(1), S-N(1) | 184 | sw-sw | T-H(1) |
| | 31 | sw-sw | V-C(1) | 186 | sw-sw | A-P(1) |
| | 46 | sw-sw | L-V(2) | 187 | hu-hu | K-Y(1) |
| | 48 | hu-sw | T-K(1) | 192 | hu-sw | M-I(1) |
| | 68 | sw-sw | V-C(1) | 202 | sw-sw | A-R(1) |
| | 70 | hu-hu | S-T(1) | 222 | hu-sw | H-P(1) |
| | 77 | hu-hu | R-I(2), R-K(2), R-T(2), K-T(1) | 234 | hu-hu | L-I(2), L-F(1) |
| | 77 | hu-sw | R-I(1), R-T(1) | 235 | hu-sw | E-D(1) |
| **MP (M2)** | 10 | hu-sw | P-H(1) | 22 | sw-sw | S-L(2) |
| | 10 | sw-sw | P-H(4), P-L(2) | 23 | hu-hu | S-N(3) |
| | 11 | hu-hu | T-I(5), I-T(1) | 23 | hu-sw | S-D(1), S-N(1) |
| | 11 | hu-sw | I-F(1), I-T(1) | 23 | sw-sw | S-N(19), S-G(3), S-D(1) |
| | 11 | sw-hu | T-I(1) | 27 | hu-hu | V-I(4), V-A(1), V-T(1) |
| | 11 | sw-sw | T-I(10), I-T(4) | 28 | sw-hu | T-I(1), T-N(1) |
| | 12 | hu-hu | R-K(3) | 43 | hu-hu | L-I(2), P-L(1), T-A(1), T-I(1), T-P(1), V-T(1) |
| | 12 | sw-sw | R-K(10), K-R(2) | 48 | hu-hu | F-L(2), F-S(2) |
| | 13 | hu-hu | S-N(6), N-S(3), N-K(1) | 48 | sw-sw | F-S(3), F-L(1) |
| | 13 | hu-sw | S-N(1), S-R(1) | 54 | hu-hu | R-L(4), L-F(1), L-I(1), L-V(1), R-H(1) |
| | 13 | sw-sw | N-S(12), S-N(8), N-K(4), N-D(1), N-T(1), S-R(1) | 56 | hu-sw | K-E(1), K-N(1) |
| | 14 | hu-hu | E-G(3), E-A(1), G-E(1) | 57 | hu-hu | Y-H(3), H-Q(1) |
| | 14 | hu-sw | E-G(1) | 57 | hu-sw | H-L(1), H-N(1), H-Y(1), Y-H(1) |
| | 14 | sw-hu | G-E(2) | 61 | sw-sw | R-G(8), R-S(3), R-K(2), R-I(1) |
| | 14 | sw-sw | G-E(20), E-G(6) | 64 | sw-sw | S-L(1), S-T(1) |
| | 16 | hu-hu | G-E(6), E-G(2) | 68 | hu-sw | V-I(1) |
| | 16 | hu-sw | E-G(1) | 82 | hu-hu | S-N(4), N-S(3) |
| | 16 | sw-sw | E-G(4), G-E(3), E-V(1) | 82 | hu-sw | S-N(4) |
| | 18 | hu-sw | R-K(2) | 82 | sw-hu | S-N(1) |
| | 18 | sw-hu | R-K(1) | 82 | sw-sw | S-N(29), S-R(2), S-I(1), S-T(1) |
| | 18 | sw-sw | R-K(13), K-R(1) | 86 | hu-hu | V-A(2), A-T(1), A-V(1) |
| | 19 | sw-hu | Y-C(1) | 86 | sw-sw | V-A(2), V-S(1) |
| | 19 | sw-sw | Y-C(5), C-Y(3), Y-S(2), C-R(1) | 89 | hu-hu | S-G(4), G-S(3), S-N(1) |
| | 20 | hu-hu | S-N(3), N-S(1), S-R(1) | 89 | hu-sw | G-S(1), S-N(1) |
| | 20 | hu-sw | N-K(1), N-S(1) | 89 | sw-sw | G-S(6), G-D(4), G-C(1) |

| | | | | | | |
|---|---|---|---|---|---|---|
| | 20 | sw-sw | S-N(6), N-K(2), N-D(1), S-G(1), S-I(1), S-R(1) | 93 | hu-hu | N-S(4), S-N(4) |
| | 21 | hu-hu | D-G(5), G-D(2) | 95 | sw-sw | V-E(5), V-A(2), E-K(1), E-Q(1), V-M(1) |
| | 21 | hu-sw | D-G(2) | 96 | hu-sw | L-H(1), L-Q(1), L-S(1) |
| | 21 | sw-sw | G-D(11), D-G(9), G-V(2), D-A(1) | 97 | hu-sw | E-T(1) |
| **NS (NS1)** | 4 | sw-hu | N-T(1) | 192 | hu-sw | V-A(1), V-I(1) |
| | 4 | sw-sw | N-K(2), N-G(1), N-I(1), N-T(1) | 192 | sw-sw | V-I(7) |
| | 7 | sw-sw | S-L(6), S-I(1), S-T(1) | 194 | hu-sw | G-S(1) |
| | 10 | sw-sw | Q-E(1), Q-K(1) | 194 | sw-hu | V-I(1) |
| | 25 | hu-hu | N-S(2), N-Q(1), Q-N(1) | 194 | sw-sw | V-I(11), G-S(5), G-V(2), V-G(1) |
| | 26 | hu-hu | D-G(3), E-K(2), D-N(1), E-D(1), E-G(1), G-K(1), K-N(1) | 197 | hu-hu | S-N(1), T-A(1), T-N(1) |
| | 26 | sw-sw | G-R(5), G-E(2), G-K(2), E-G(1), R-K(1) | 197 | sw-sw | A-T(16), A-D(3), N-S(2), T-A(2), T-N(2), A-N(1), N-T(1), S-I(1), S-N(1) |
| | 27 | sw-sw | L-F(11), L-I(2), L-R(2), L-H(1), L-M(1) | 198 | hu-hu | L-I(4), I-L(1) |
| | 40 | hu-hu | Q-L(1) | 200 | hu-hu | R-K(1) |
| | 42 | hu-hu | S-A(1) | 202 | hu-hu | A-T(2) |
| | 59 | hu-sw | H-F(1), L-I(1) | 202 | hu-sw | A-S(1), A-T(1) |
| | 59 | sw-sw | R-H(22), R-C(7), L-I(5), L-F(4), R-L(3), R-S(3), L-R(1), R-Y(1) | 202 | sw-sw | A-T(16), A-S(3), T-A(1), T-N(1) |
| | 71 | sw-hu | D-G(1) | 204 | sw-sw | R-G(14), R-K(1) |
| | 96 | hu-sw | E-M(1) | 205 | hu-hu | S-N(2), N-K(1), N-S(1), S-G(1), S-R(1) |
| | 105 | hu-hu | L-V(2) | 205 | sw-sw | N-S(9), S-N(7), S-G(5), N-K(4), S-I(4), N-D(2), I-V(1), N-I(1), N-T(1), S-C(1), S-D(1), S-R(1) |
| | 109 | hu-sw | Q-G(1) | 206 | hu-hu | S-C(3), C-S(1), R-H(1), R-L(1) |
| | 127 | sw-sw | N-T(37), N-D(9), N-S(8), N-K(3), T-N(3), K-R(2), N-I(2), T-I(2), K-E(1), K-I(1), N-H(1) | 206 | hu-sw | C-S(2), R-C(2), C-R(1), S-C(1), S-N(1) |
| | 129 | sw-sw | T-I(10), I-T(8), I-M(2), I-V(2), T-A(2), V-I(2), I-L(1), T-R(1) | 206 | sw-sw | T-I(13), R-C(10), I-T(5), R-H(4), R-S(4), T-A(4), R-L(3), I-S(2), T-S(2), C-S(1), H-Y(1), I-K(1), L-I(1), S-I(1), T-P(1) |
| | 141 | sw-sw | L-M(6), L-I(4), L-V(2), L-G(1) | 207 | hu-hu | N-D(5), D-N(1) |
| | 151 | hu-sw | T-S(2) | 207 | hu-sw | D-N(2) |
| | 151 | sw-sw | T-N(2), T-D(1), T-M(1) | 207 | sw-sw | N-D(7), N-Y(2), D-N(1), N-T(1) |
| | 155 | hu-hu | A-T(4), A-R(1), A-S(1), S-A(1) | 210 | sw-sw | G-R(16), G-E(3), G-W(2), R-G(2), R-W(1) |
| | 164 | sw-sw | P-L(4), P-S(4), L-I(1), L-P(1), P-H(1), P-Y(1) | 211 | hu-hu | R-G(4), G-R(2), R-K(1) |
| | 166 | hu-hu | L-F(3), F-L(1) | 211 | sw-sw | R-G(13), R-K(1), R-N(1) |
| | 166 | sw-hu | L-F(1) | 212 | hu-hu | P-S(2) |
| | 166 | sw-sw | L-F(10), L-I(4), L-V(1) | 212 | sw-hu | P-S(1) |
| | 167 | hu-sw | P-S(1) | 212 | sw-sw | P-S(19), P-L(4), P-H(1) |
| | 167 | sw-sw | P-L(1), P-S(1) | 213 | hu-hu | P-S(3), S-P(1) |
| | 170 | hu-hu | T-I(1) | 213 | hu-sw | S-P(2) |
| | 170 | sw-sw | T-A(2), T-N(2), T-I(1) | 213 | sw-hu | S-P(2) |
| | 171 | hu-hu | I-Y(2), D-A(1), D-G(1), D-N(1), I-F(1), Y-H(1), Y-I(1) | 213 | sw-sw | P-S(12), S-P(12), P-L(2), P-T(1) |
| | 171 | hu-sw | Y-H(3), D-E(1), Y-C(1) | 215 | hu-hu | P-S(1), P-T(1), T-P(1) |
| | 171 | sw-hu | D-N(1) | 215 | hu-sw | P-S(5) |
| | 171 | sw-sw | D-N(18), D-G(9), D-Y(3), Y-H(3), D-E(2), N-T(2), N-G(1), N-I(1), N-K(1), Y-C(1) | 215 | sw-hu | T-P(1) |
| | 172 | hu-sw | E-K(1) | 215 | sw-sw | P-S(21), P-F(2), P-T(2), S-P(2), P-H(1), P-L(1), T-N(1) |
| | 172 | sw-sw | E-K(4), E-G(1) | 216 | hu-hu | P-S(2), P-T(1) |

| | | | | | | |
|---|---|---|---|---|---|---|
| | 173 | sw-sw | D-N(4) | 216 | hu-sw | P-S(3) |
| | 174 | hu-hu | V-I(2) | 216 | sw-sw | P-S(24), P-T(9), T-A(1), T-P(1) |
| | 175 | sw-sw | K-E(4), K-N(1), K-V(1) | 218 | hu-sw | Q-K(1) |
| | 176 | hu-sw | N-I(1) | 218 | sw-sw | Q-P(2), Q-H(1), Q-K(1) |
| | 176 | sw-hu | N-D(1) | 219 | hu-sw | K-E(1), K-N(1) |
| | 176 | sw-sw | N-D(7), N-I(3), N-H(1), N-S(1), N-T(1), N-Y(1), T-I(1) | 219 | sw-hu | E-K(1) |
| | 178 | hu-hu | V-I(7), I-V(2), V-A(1) | 219 | sw-sw | K-E(8), E-K(1) |
| | 178 | hu-sw | V-I(4) | 223 | sw-sw | A-E(4) |
| | 178 | sw-hu | V-I(1) | 224 | hu-hu | G-R(3), R-G(1) |
| | 178 | sw-sw | V-I(19), I-V(6), V-T(1) | 224 | sw-sw | R-G(5), G-R(2), R-K(2) |
| | 180 | hu-hu | V-I(2), V-F(1) | 225 | sw-sw | T-A(8), A-T(1) |
| | 180 | hu-sw | V-I(2) | 226 | hu-hu | I-A(1), V-A(1) |
| | 180 | sw-sw | V-I(22), I-T(7), I-V(5), V-A(3), I-F(2), I-L(1), V-F(1), V-L(1) | 226 | sw-sw | I-T(8), T-I(6), I-V(5), T-A(4) |
| | 185 | sw-sw | L-F(9), L-P(1) | 227 | hu-hu | R-G(1), R-K(1) |
| | 188 | hu-sw | N-S(2) | 227 | hu-sw | R-G(1) |
| | 189 | hu-hu | D-G(3), D-N(1), G-D(1) | 227 | sw-hu | G-R(1), R-E(1) |
| | 189 | hu-sw | G-S(2), G-D(1) | 227 | sw-sw | G-R(8), E-G(1), R-G(1) |
| | 189 | sw-hu | G-S(2) | 230 | hu-hu | V-I(2) |
| | 189 | sw-sw | D-N(7), G-D(7), G-S(5), D-G(3), G-N(2), G-C(1) | 230 | hu-sw | V-I(1) |
| | 191 | sw-sw | A-T(5), T-K(4), T-A(3), T-R(1) | 230 | sw-sw | V-I(14) |
| **NS (NEP)** | 4 | sw-hu | N-T(1) | 29 | sw-sw | N-S(23), S-N(3) |
| | 4 | sw-sw | N-K(2), N-G(1), N-I(1), N-T(1) | 32 | hu-hu | I-V(3), V-I(1) |
| | 7 | sw-sw | S-L(6), S-I(1), S-T(1) | 32 | hu-sw | V-A(1), V-I(1) |
| | 8 | hu-hu | S-T(1) | 32 | sw-sw | V-I(9), I-V(3), I-M(1), I-T(1) |
| | 10 | sw-sw | Q-E(1), Q-K(1) | 33 | sw-sw | T-M(5), M-T(3), T-I(2) |
| | 12 | sw-sw | I-L(1), I-M(1), I-R(1), I-T(1) | 35 | hu-sw | F-L(1) |
| | 18 | sw-sw | K-I(1), K-N(1), K-R(1), K-Y(1) | 54 | sw-sw | D-N(1), D-T(1) |
| | 19 | hu-hu | M-I(1), M-L(1) | 60 | sw-hu | N-H(1), N-S(1), N-T(1), S-N(1) |
| | 23 | sw-sw | S-P(10), S-Y(2) | 71 | hu-hu | Q-K(1) |
| | 26 | hu-hu | E-G(2), G-E(1) | 89 | hu-hu | T-A(3), A-V(2), A-E(1), A-T(1), I-M(1), M-S(1), T-I(1) |
| | 26 | hu-sw | E-G(2) | 101 | sw-hu | Q-E(1) |
| | 26 | sw-hu | E-G(1) | 107 | hu-hu | F-L(3), F-S(1), L-F(1) |
| | 29 | hu-hu | N-S(3), S-N(1) | 118 | hu-sw | F-C(1), F-V(1) |
| | 29 | hu-sw | N-S(3) | | | |

**Figure S3:** Pervasive positive and purifying selection across sites of each protein, measured by FUBAR analysis. **a.** Proportions of codons with evidence of pervasive positive (blue) or purifying (red) selection under a posterior probability threshold > 0.9. **b.** Distribution of non-synonymous to synonymous substitution rates, ω (dN/dS), across sites of each protein. Positions with evidence of pervasive positive (blue) or purifying (red) selection under a posterior probability threshold > 0.9 are marked with color.

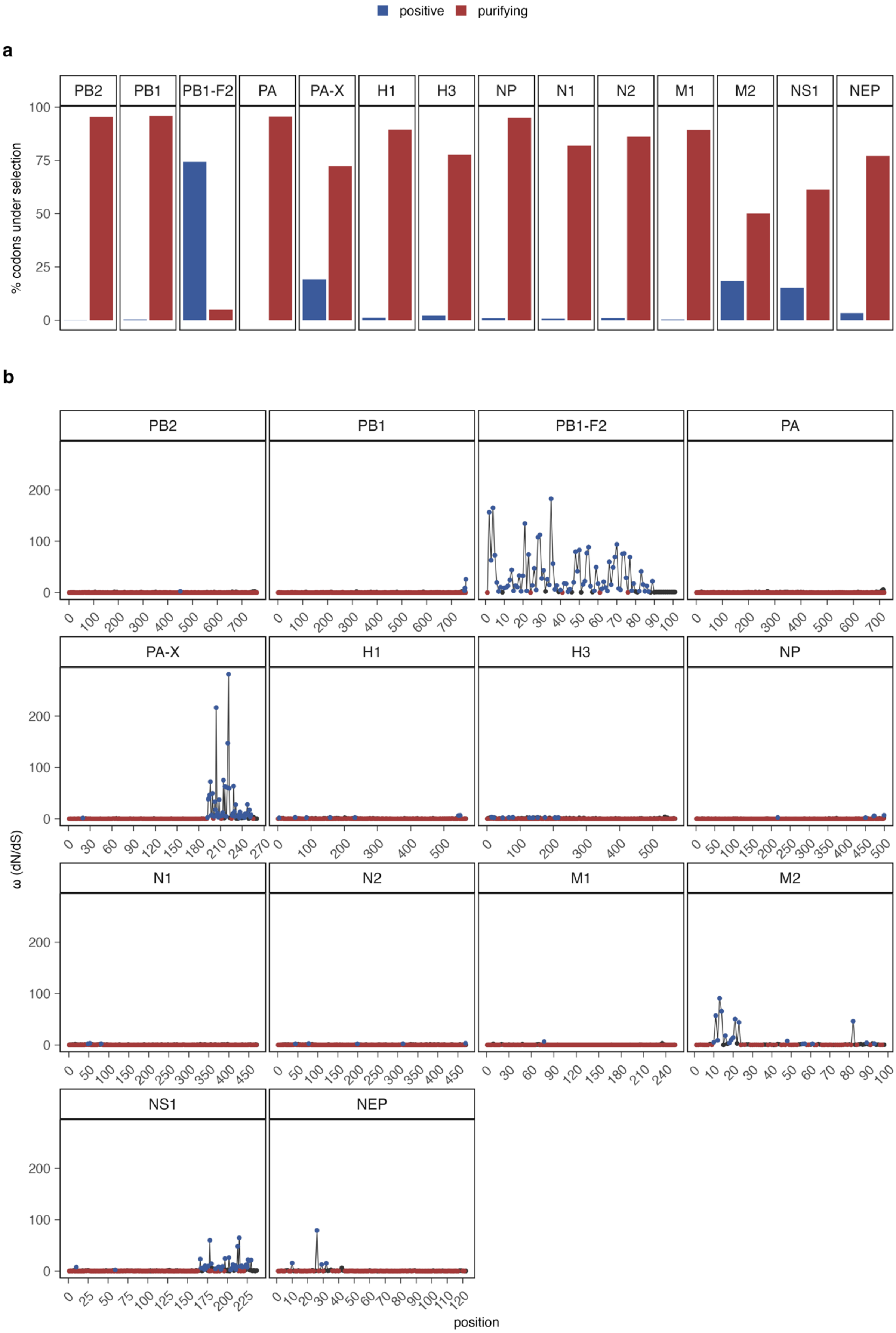
positive
purifying
a
PB2
PB1
PB1-F2
PA
PA-X
H1
H3
NP
N1
N2
M1
M2
NS1
NEP
% codons under selection
b
ω (dN/dS)
position

**Figure S4:** Proportions of positively selected sites across transmission types measured by MEME and FUBAR analysis. **a**. Proportion of unique vs. shared positively selected positions for each transmission category and protein. The proportions are calculated as the number of positively selected sites in a transmission category that are either unique or also present in one or more other transmission categories (shared) divided by the total number of positively selected sites in the transmission category. **b**. Bar plots showing the count of selected positions unique for each transmission type or shared between transmission types for each protein. Transmission categories are hu-hu: human-human, sw-sw: swine-swine, hu-sw: human-swine and sw-hu: swine-human. **c**. Percentage of positively selected sites identified by MEME analysis that overlaps with sites with either positive or purifying pervasive selection identified by FUBAR analysis. Proportions of sites that overlap with positively selected sites identified by FUBAR are shown with fully saturated color, while proportions of sites that overlap with sites under purifying selection identified by FUBAR are shown in semi-transparent color.

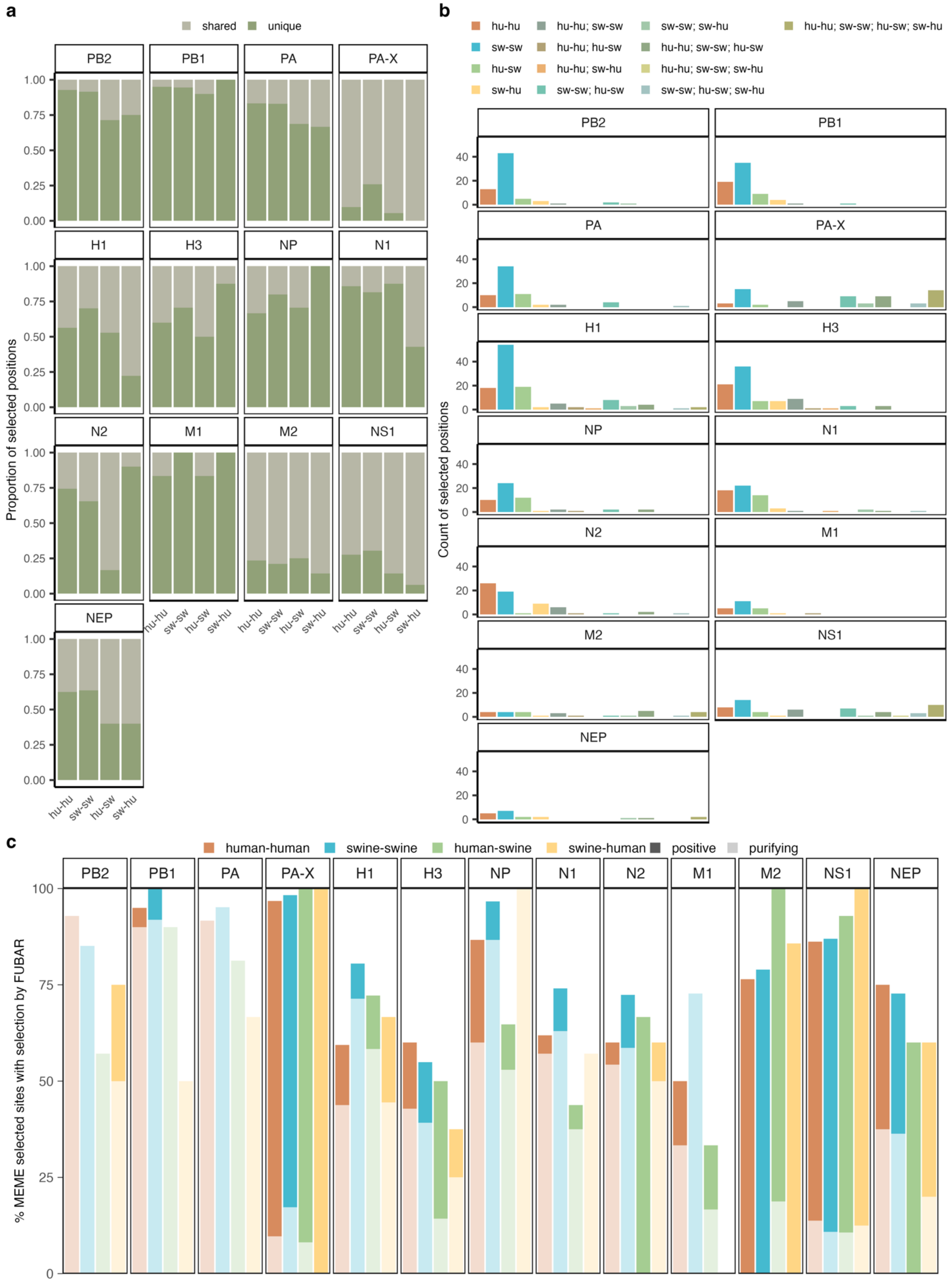

a
shared
unique
PB2
PB1
PA
PA-X
H1
H3
NP
N1
N2
M1
M2
NS1
NEP
Proportion of selected positions
1.00
0.75
0.50
0.25
0.00
hu-hu
sw-sw
hu-sw
sw-hu
b
hu-hu
sw-sw
hu-sw
sw-hu
hu-hu; sw-sw
hu-hu; hu-sw
hu-hu; sw-hu
sw-sw; hu-sw
sw-sw; sw-hu
hu-hu; sw-sw; hu-sw
hu-hu; sw-sw; sw-hu
sw-sw; hu-sw; sw-hu
hu-hu; sw-sw; hu-sw; sw-hu
Count of selected positions
40
20
0
c
human-human
swine-swine
human-swine
swine-human
positive
purifying
% MEME selected sites with selection by FUBAR
100
75
50
25
0

**Figure S5:** ROC (Receiver Operating Characteristics) curves for each of the protein XGBoost models as indicated in the headings of each subfigure **a-n**. Each plot shows the true positive rate (y-axis) vs. the false positive rate (x-axis), and lines represent rates for each of the four transmission classes in a one-vs-rest scheme, meaning that each class is compared against all other classes (assumed as one class). Rates are shown for each of the 20 independent model runs, color-coded by the transmission type (orange = human-human, blue = swine-swine, green = human-swine, yellow = swine-human), and the average area under the curve (AUC) for each of the transmission types are stated in the legend. A red dotted line indicates the rate of random guessing, where the true positive rate = the false positive rate and the AUC = 0.5.

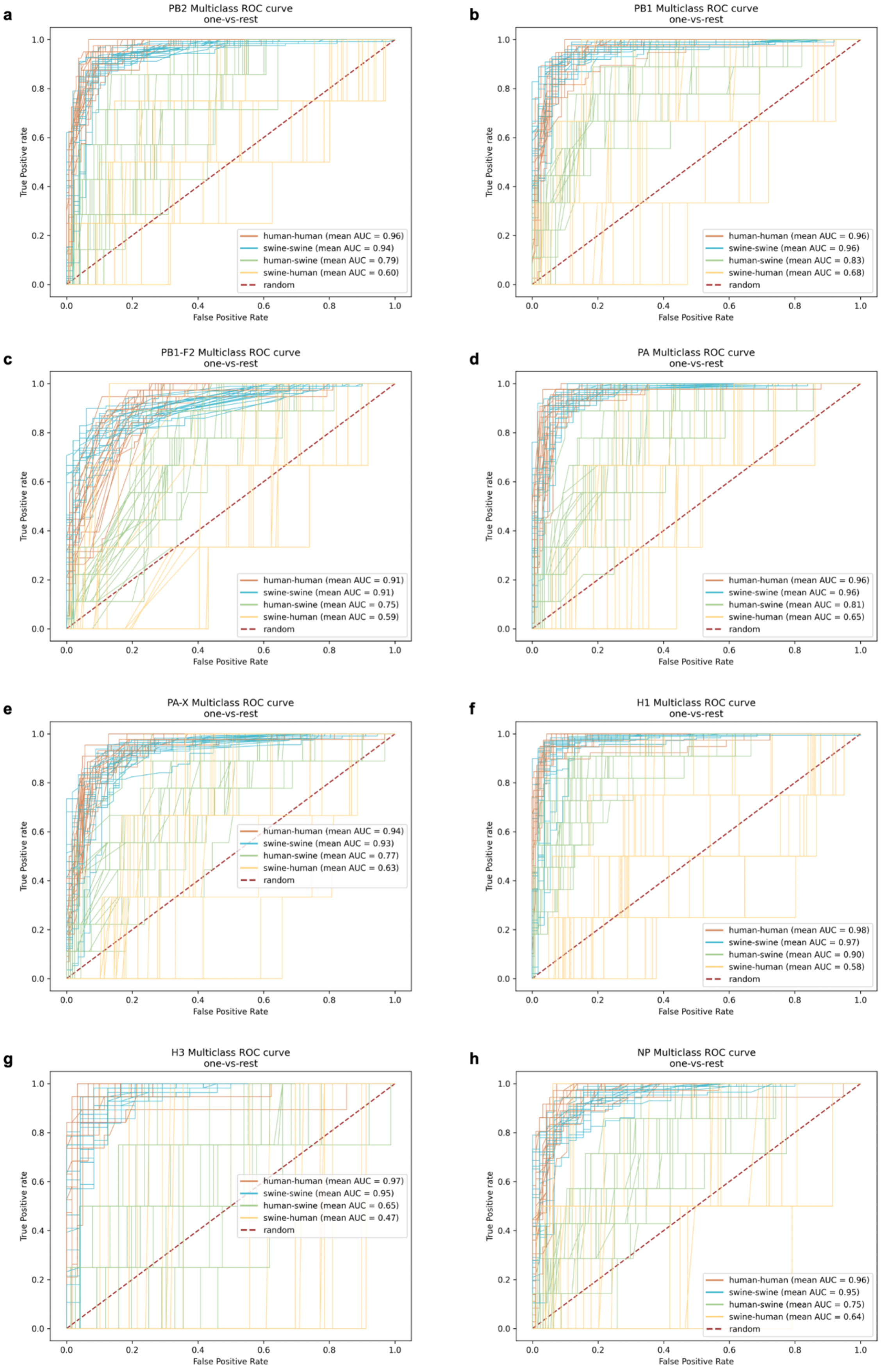
a
PB2 Multiclass ROC curve
one-vs-rest
True Positive rate
False Positive Rate
human-human (mean AUC = 0.96)
swine-swine (mean AUC = 0.94)
human-swine (mean AUC = 0.79)
swine-human (mean AUC = 0.60)
random
b
PB1 Multiclass ROC curve
one-vs-rest
True Positive rate
False Positive Rate
human-human (mean AUC = 0.96)
swine-swine (mean AUC = 0.96)
human-swine (mean AUC = 0.83)
swine-human (mean AUC = 0.68)
random
c
PB1-F2 Multiclass ROC curve
one-vs-rest
True Positive rate
False Positive Rate
human-human (mean AUC = 0.91)
swine-swine (mean AUC = 0.91)
human-swine (mean AUC = 0.75)
swine-human (mean AUC = 0.59)
random
d
PA Multiclass ROC curve
one-vs-rest
True Positive rate
False Positive Rate
human-human (mean AUC = 0.96)
swine-swine (mean AUC = 0.96)
human-swine (mean AUC = 0.81)
swine-human (mean AUC = 0.65)
random
e
PA-X Multiclass ROC curve
one-vs-rest
True Positive rate
False Positive Rate
human-human (mean AUC = 0.94)
swine-swine (mean AUC = 0.93)
human-swine (mean AUC = 0.77)
swine-human (mean AUC = 0.63)
random
f
H1 Multiclass ROC curve
one-vs-rest
True Positive rate
False Positive Rate
human-human (mean AUC = 0.98)
swine-swine (mean AUC = 0.97)
human-swine (mean AUC = 0.90)
swine-human (mean AUC = 0.58)
random
g
H3 Multiclass ROC curve
one-vs-rest
True Positive rate
False Positive Rate
human-human (mean AUC = 0.97)
swine-swine (mean AUC = 0.95)
human-swine (mean AUC = 0.65)
swine-human (mean AUC = 0.47)
random
h
NP Multiclass ROC curve
one-vs-rest
True Positive rate
False Positive Rate
human-human (mean AUC = 0.96)
swine-swine (mean AUC = 0.95)
human-swine (mean AUC = 0.75)
swine-human (mean AUC = 0.64)
random

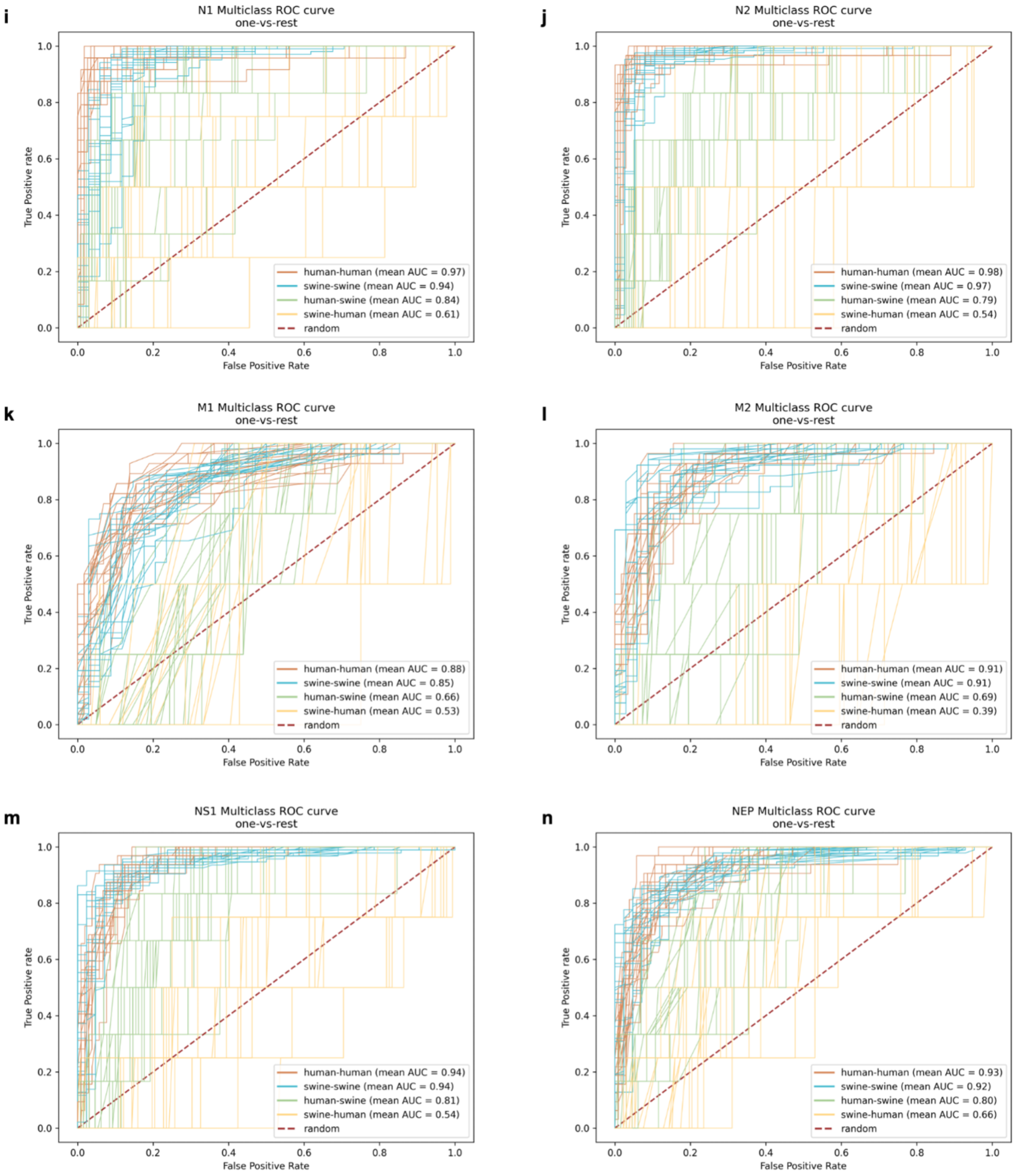

i
N1 Multiclass ROC curve
one-vs-rest
human-human (mean AUC = 0.97)
swine-swine (mean AUC = 0.94)
human-swine (mean AUC = 0.84)
swine-human (mean AUC = 0.61)
random
j
N2 Multiclass ROC curve
one-vs-rest
human-human (mean AUC = 0.98)
swine-swine (mean AUC = 0.97)
human-swine (mean AUC = 0.79)
swine-human (mean AUC = 0.54)
random
k
M1 Multiclass ROC curve
one-vs-rest
human-human (mean AUC = 0.88)
swine-swine (mean AUC = 0.85)
human-swine (mean AUC = 0.66)
swine-human (mean AUC = 0.53)
random
l
M2 Multiclass ROC curve
one-vs-rest
human-human (mean AUC = 0.91)
swine-swine (mean AUC = 0.91)
human-swine (mean AUC = 0.69)
swine-human (mean AUC = 0.39)
random
m
NS1 Multiclass ROC curve
one-vs-rest
human-human (mean AUC = 0.94)
swine-swine (mean AUC = 0.94)
human-swine (mean AUC = 0.81)
swine-human (mean AUC = 0.54)
random
n
NEP Multiclass ROC curve
one-vs-rest
human-human (mean AUC = 0.93)
swine-swine (mean AUC = 0.92)
human-swine (mean AUC = 0.80)
swine-human (mean AUC = 0.66)
random
True Positive rate
False Positive Rate

**Figure S6:** SHAP summary plots for the averages of 20 independent model trainings on amino acid data for each IAV protein as indicated by the plot headings in subfigures **a-n**. On the left, SHAP feature importance plots show the mean absolute importance scores of the top 20 features that influence the models' predictions across all classes. Class-specific impacts are denoted by color-coded bars for each feature in descending order from the most important (top bar) and the x-axis indicates the magnitude of the feature's effect on model output. Orange = human-human, blue = swine-swine, green = human-swine, yellow = swine-human. On the right, SHAP violin plots for each class separately reveal the distribution and variability of SHAP values for each feature for the given class. The directional impact on model predictions is indicated on the x-axis with negative SHAP values indicating a negative impact and positive values indicating a positive impact on mode predictions. The color represents the value of the feature from low (blue) to high (red) – in our case only two values are present, 0 (low) = feature not present, 1 (high) = feature present. Purple shades indicate overlapping points, meaning similar measured Shapley values for specific instances both in presence and absence of the feature. The violins plots are stacked by importance (sum of the absolute SHAP values for the feature) and the top 20 most important features for each class are shown.

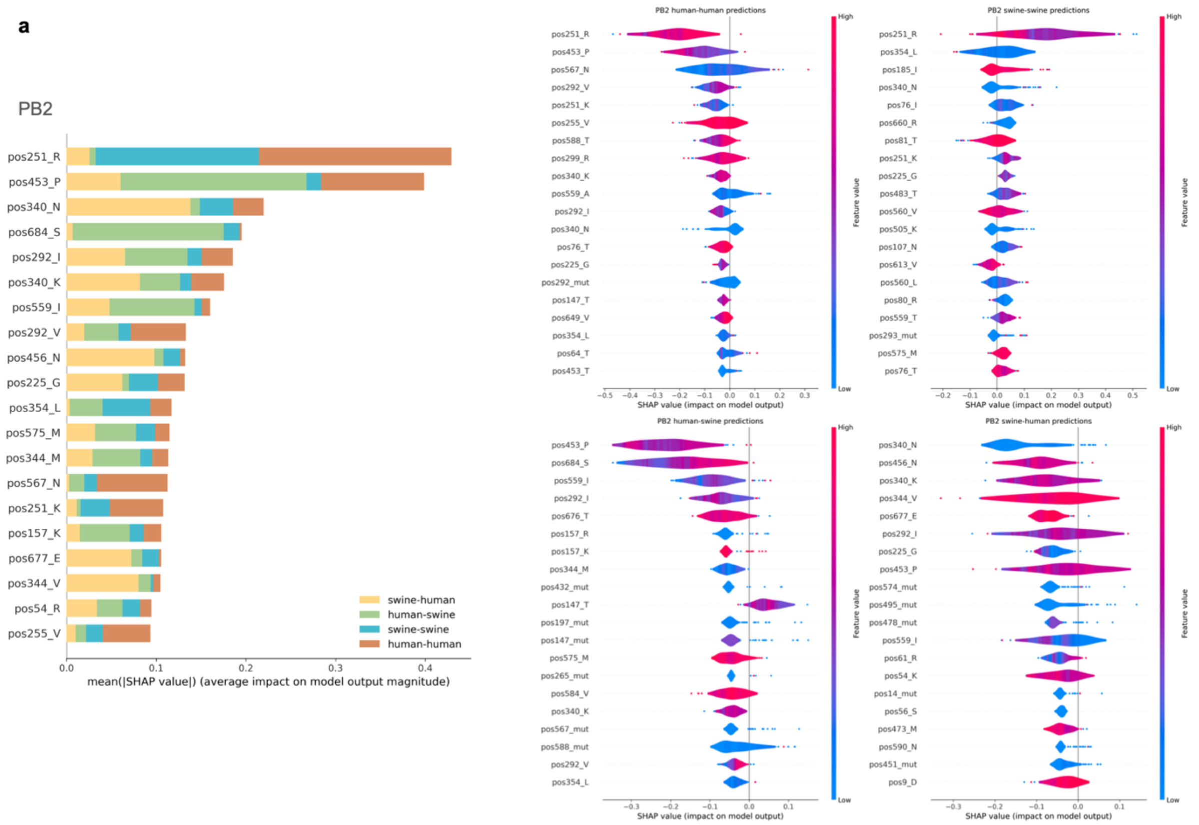
a
PB2
swine-human
human-swine
swine-swine
human-human
mean(|SHAP value|) (average impact on model output magnitude)
PB2 human-human predictions
PB2 swine-swine predictions
PB2 human-swine predictions
PB2 swine-human predictions
SHAP value (impact on model output)
Feature value
High
Low

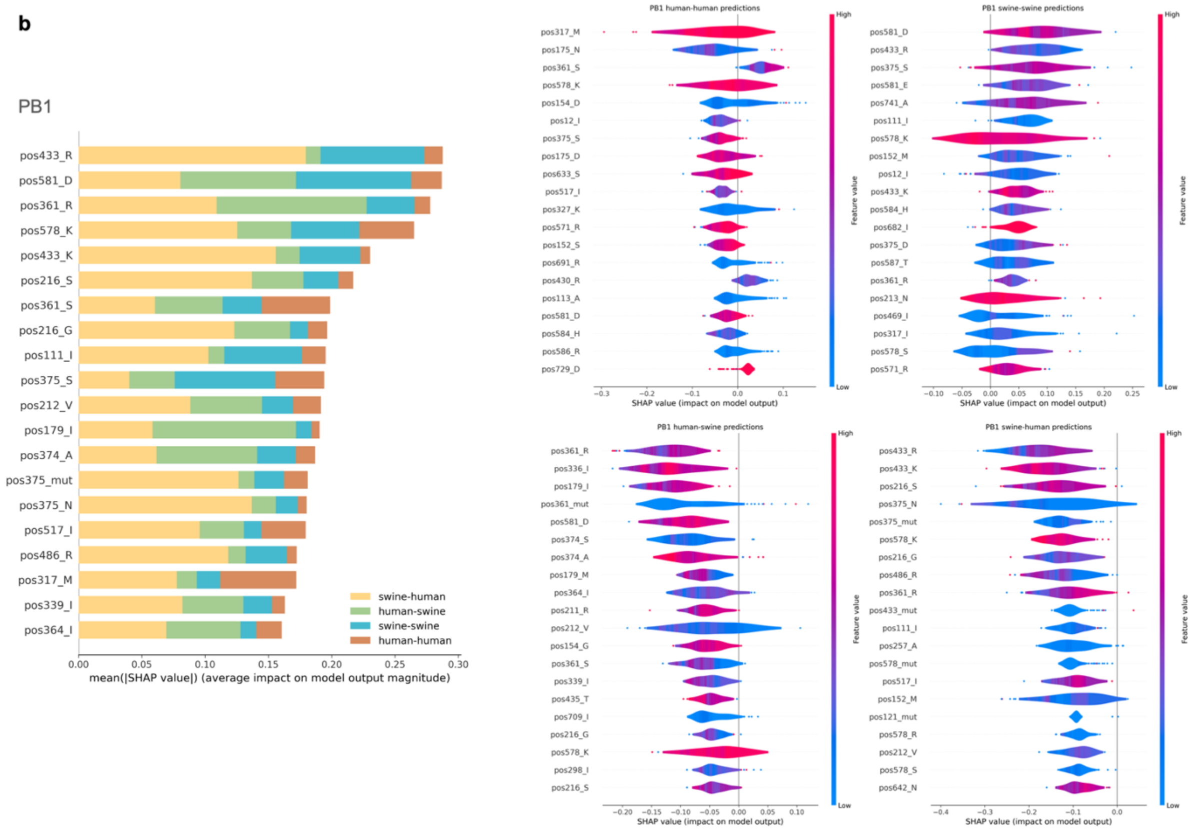
b
PB1
swine-human
human-swine
swine-swine
human-human
mean(|SHAP value|) (average impact on model output magnitude)
PB1 human-human predictions
PB1 swine-swine predictions
PB1 human-swine predictions
PB1 swine-human predictions
SHAP value (impact on model output)
Feature value
High
Low

**c**

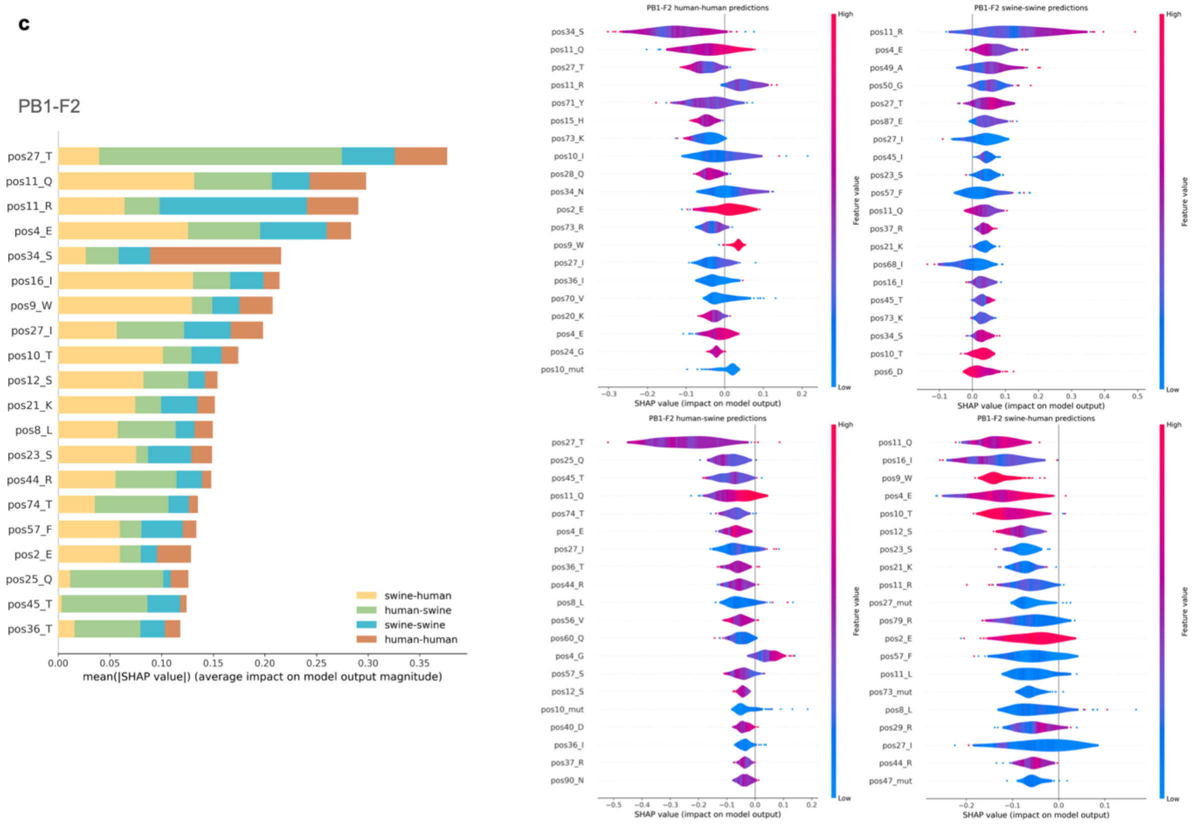
PB1-F2
pos27_T
pos11_Q
pos11_R
pos4_E
pos34_S
pos16_I
pos9_W
pos27_I
pos10_T
pos12_S
pos21_K
pos8_L
pos23_S
pos44_R
pos74_T
pos57_F
pos2_E
pos25_Q
pos45_T
pos36_T
swine-human
human-swine
swine-swine
human-human
mean(|SHAP value|) (average impact on model output magnitude)
PB1-F2 human-human predictions
PB1-F2 swine-swine predictions
PB1-F2 human-swine predictions
PB1-F2 swine-human predictions
SHAP value (impact on model output)
Feature value
High
Low

**d**

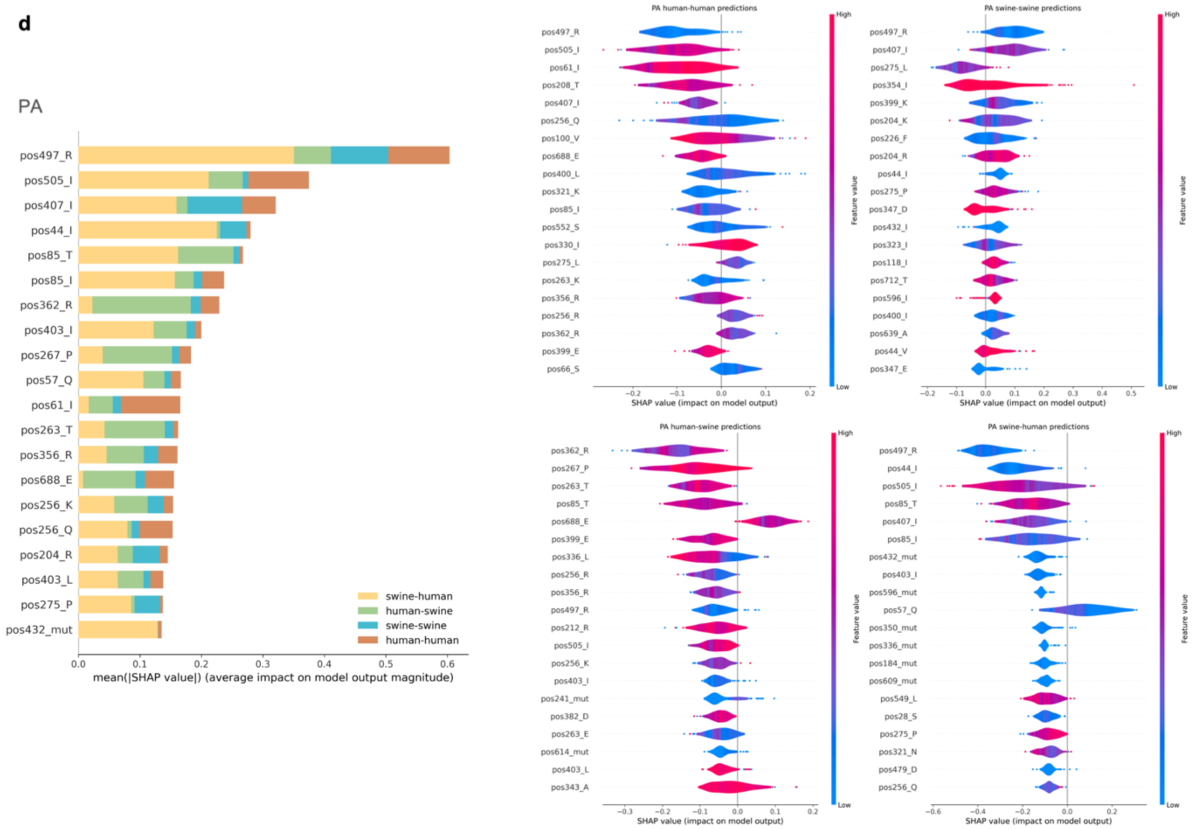
PA
pos497_R
pos505_I
pos407_I
pos44_I
pos85_T
pos85_I
pos362_R
pos403_I
pos267_P
pos57_Q
pos61_I
pos263_T
pos356_R
pos688_E
pos256_K
pos256_Q
pos204_R
pos403_L
pos275_P
pos432_mut
swine-human
human-swine
swine-swine
human-human
mean(|SHAP value|) (average impact on model output magnitude)
PA human-human predictions
PA swine-swine predictions
PA human-swine predictions
PA swine-human predictions
SHAP value (impact on model output)
Feature value
High
Low

e

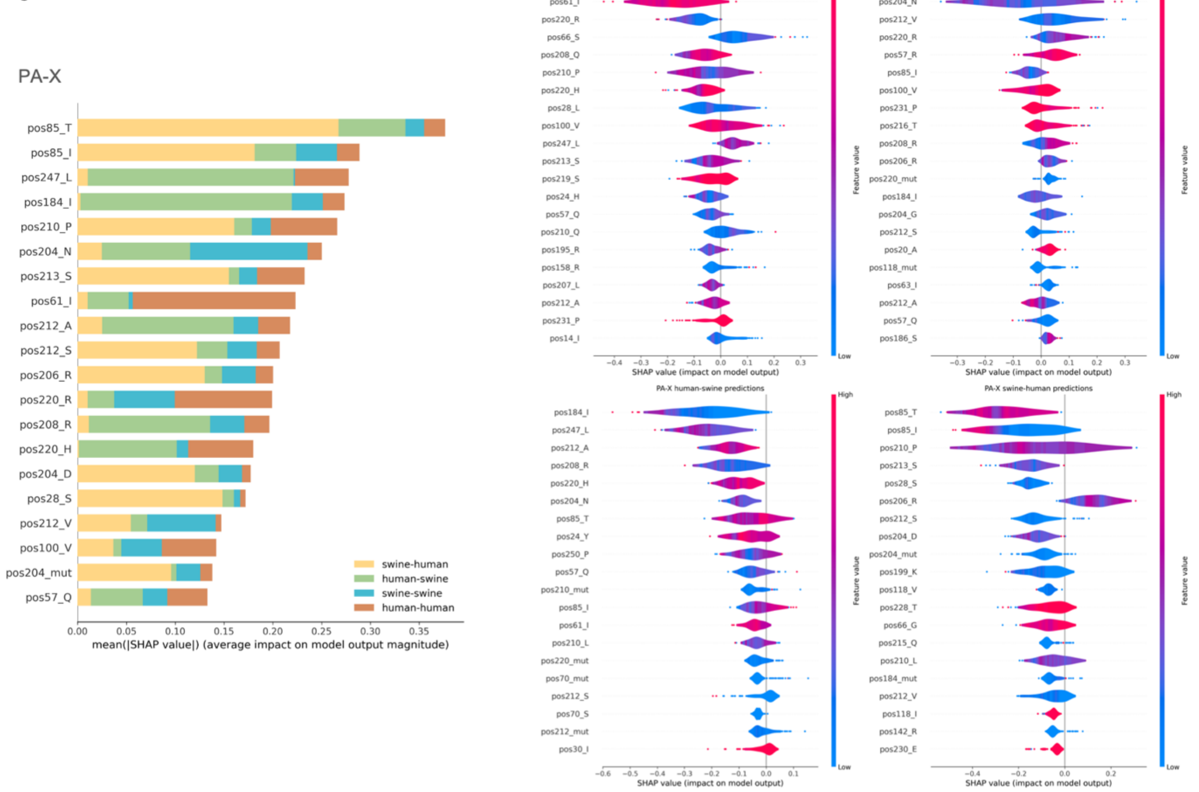
PA-X
pos85_T
pos85_I
pos247_L
pos184_I
pos210_P
pos204_N
pos213_S
pos61_I
pos212_A
pos212_S
pos206_R
pos220_R
pos208_R
pos220_H
pos204_D
pos28_S
pos212_V
pos100_V
pos204_mut
pos57_Q
0.00
0.05
0.10
0.15
0.20
0.25
0.30
0.35
mean(|SHAP value|) (average impact on model output magnitude)
swine-human
human-swine
swine-swine
human-human
PA-X human-human predictions
PA-X swine-swine predictions
PA-X human-swine predictions
PA-X swine-human predictions
SHAP value (impact on model output)
Feature value
High
Low

f

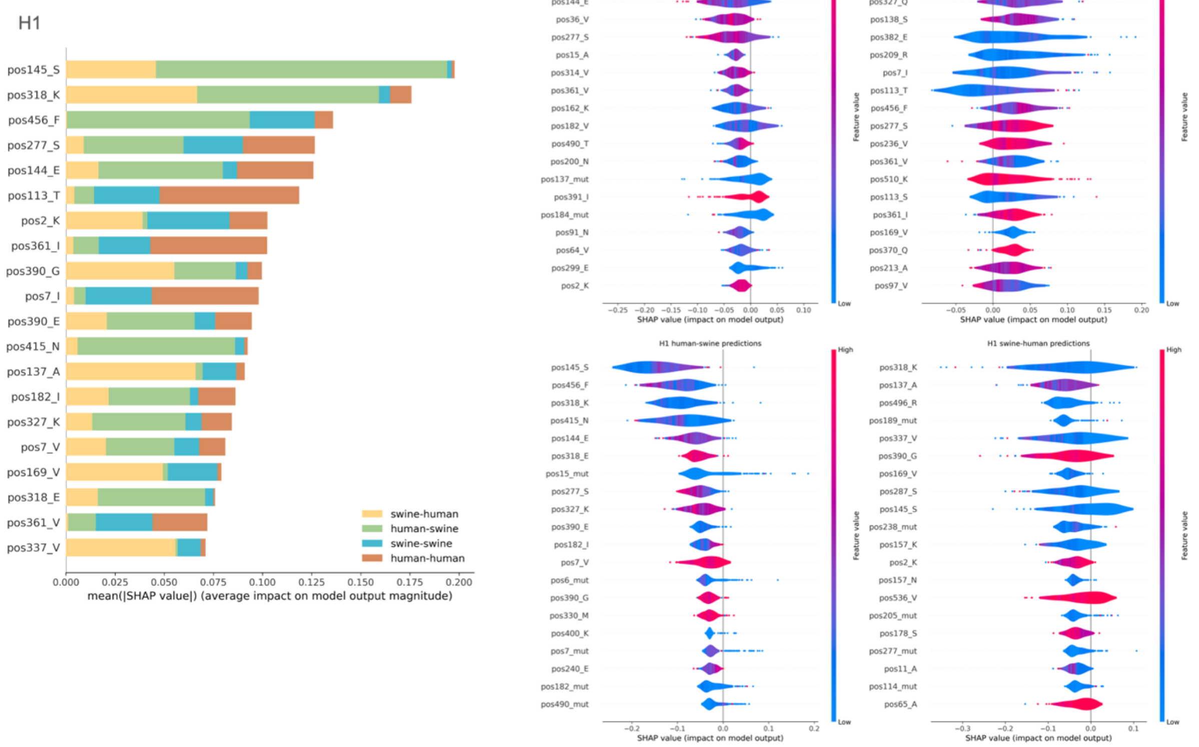
H1
pos145_S
pos318_K
pos456_F
pos277_S
pos144_E
pos113_T
pos2_K
pos361_I
pos390_G
pos7_I
pos390_E
pos415_N
pos137_A
pos182_I
pos327_K
pos7_V
pos169_V
pos318_E
pos361_V
pos337_V
0.000
0.025
0.050
0.075
0.100
0.125
0.150
0.175
0.200
mean(|SHAP value|) (average impact on model output magnitude)
swine-human
human-swine
swine-swine
human-human
H1 human-human predictions
H1 swine-swine predictions
H1 human-swine predictions
H1 swine-human predictions
SHAP value (impact on model output)
Feature value
High
Low

**g**

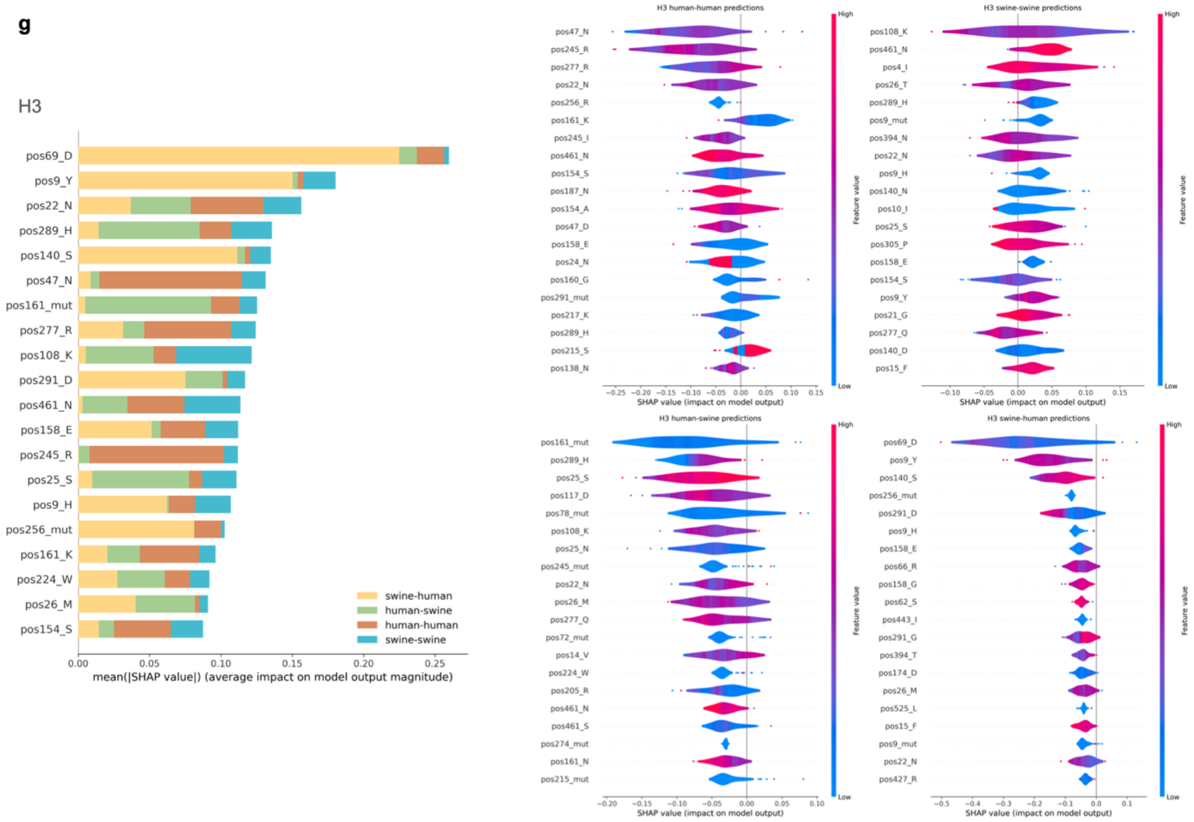
H3
pos69_D
pos9_Y
pos22_N
pos289_H
pos140_S
pos47_N
pos161_mut
pos277_R
pos108_K
pos291_D
pos461_N
pos158_E
pos245_R
pos25_S
pos9_H
pos256_mut
pos161_K
pos224_W
pos26_M
pos154_S
swine-human
human-swine
human-human
swine-swine
0.00
0.05
0.10
0.15
0.20
0.25
mean(|SHAP value|) (average impact on model output magnitude)
H3 human-human predictions
H3 swine-swine predictions
H3 human-swine predictions
H3 swine-human predictions
SHAP value (impact on model output)
Feature value
High
Low

**h**

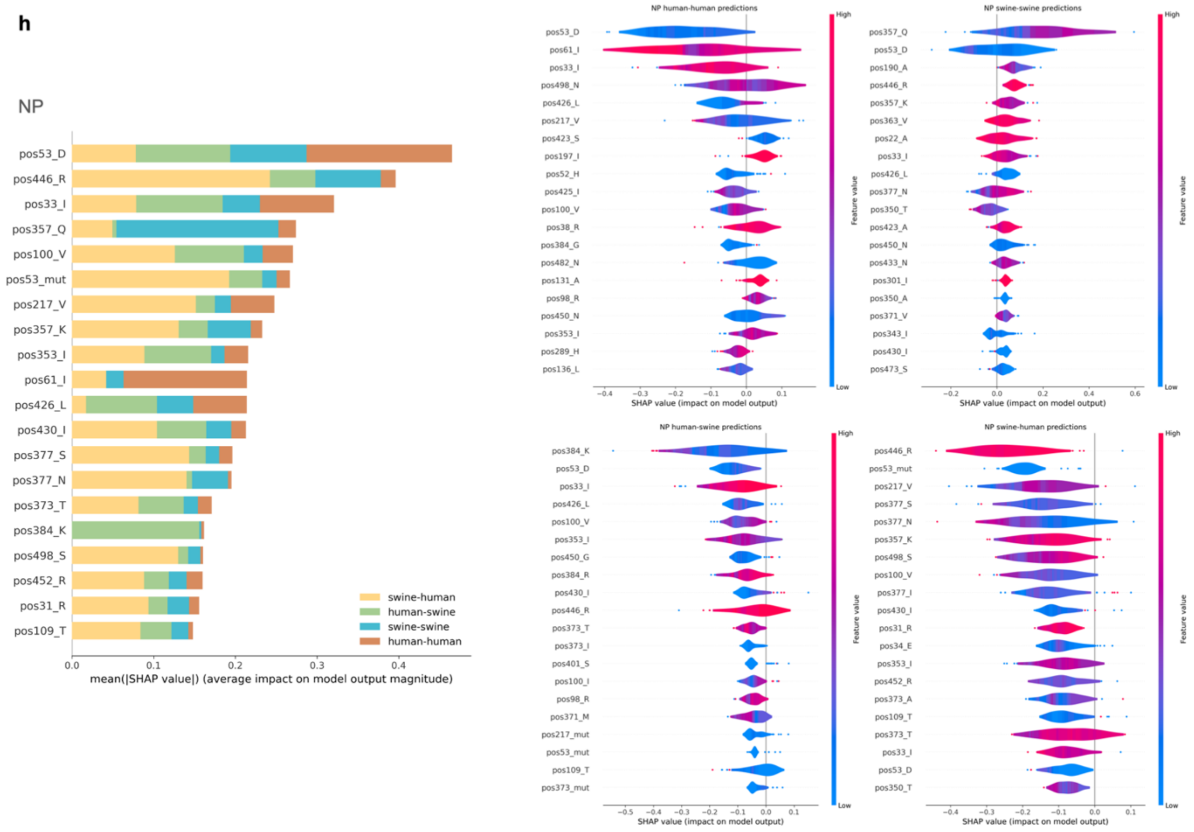
NP
pos53_D
pos446_R
pos33_I
pos357_Q
pos100_V
pos53_mut
pos217_V
pos357_K
pos353_I
pos61_I
pos426_L
pos430_I
pos377_S
pos377_N
pos373_T
pos384_K
pos498_S
pos452_R
pos31_R
pos109_T
swine-human
human-swine
swine-swine
human-human
0.0
0.1
0.2
0.3
0.4
mean(|SHAP value|) (average impact on model output magnitude)
NP human-human predictions
NP swine-swine predictions
NP human-swine predictions
NP swine-human predictions
SHAP value (impact on model output)
Feature value
High
Low

**i**

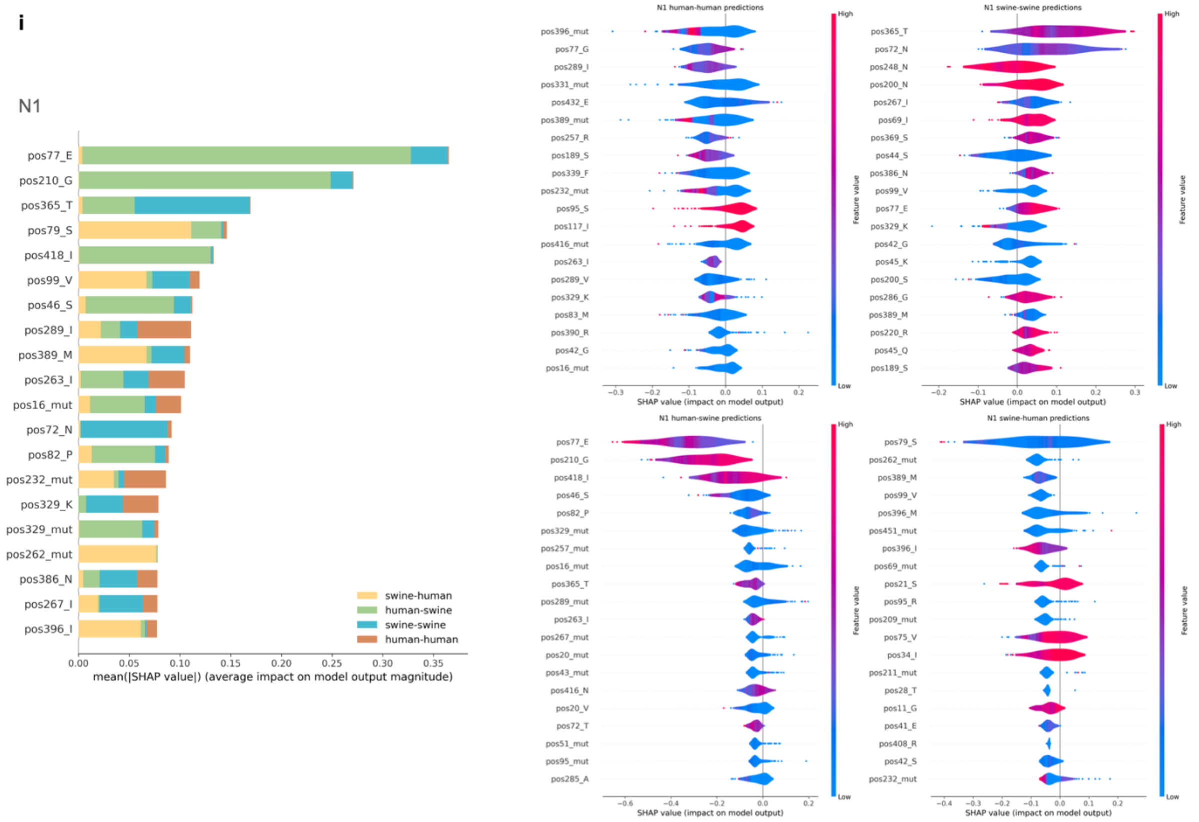
N1
pos77_E
pos210_G
pos365_T
pos79_S
pos418_I
pos99_V
pos46_S
pos289_I
pos389_M
pos263_I
pos16_mut
pos72_N
pos82_P
pos232_mut
pos329_K
pos329_mut
pos262_mut
pos386_N
pos267_I
pos396_I
swine-human
human-swine
swine-swine
human-human
mean(|SHAP value|) (average impact on model output magnitude)
N1 human-human predictions
N1 swine-swine predictions
N1 human-swine predictions
N1 swine-human predictions
SHAP value (impact on model output)
Feature value
High
Low

**j**

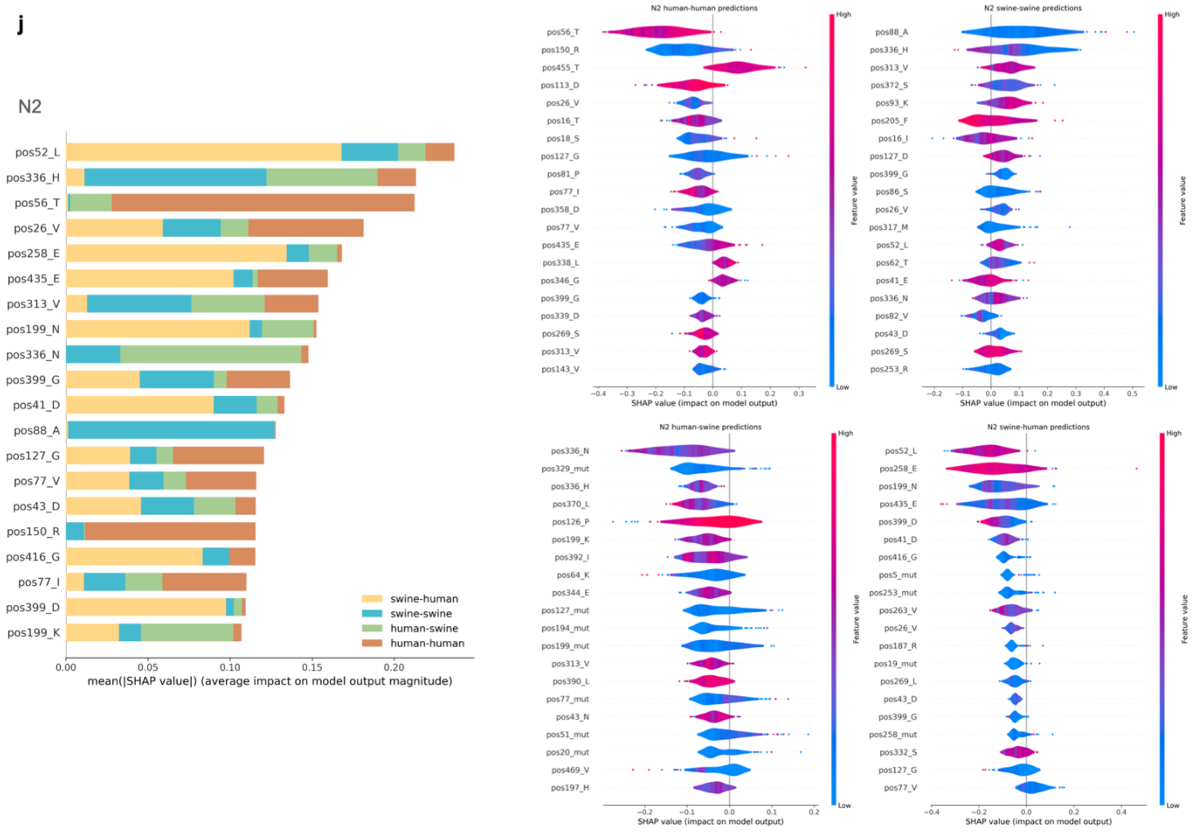
N2
pos52_L
pos336_H
pos56_T
pos26_V
pos258_E
pos435_E
pos313_V
pos199_N
pos336_N
pos399_G
pos41_D
pos88_A
pos127_G
pos77_V
pos43_D
pos150_R
pos416_G
pos77_I
pos399_D
pos199_K
swine-human
swine-swine
human-swine
human-human
mean(|SHAP value|) (average impact on model output magnitude)
N2 human-human predictions
N2 swine-swine predictions
N2 human-swine predictions
N2 swine-human predictions
SHAP value (impact on model output)
Feature value
High
Low

**k**

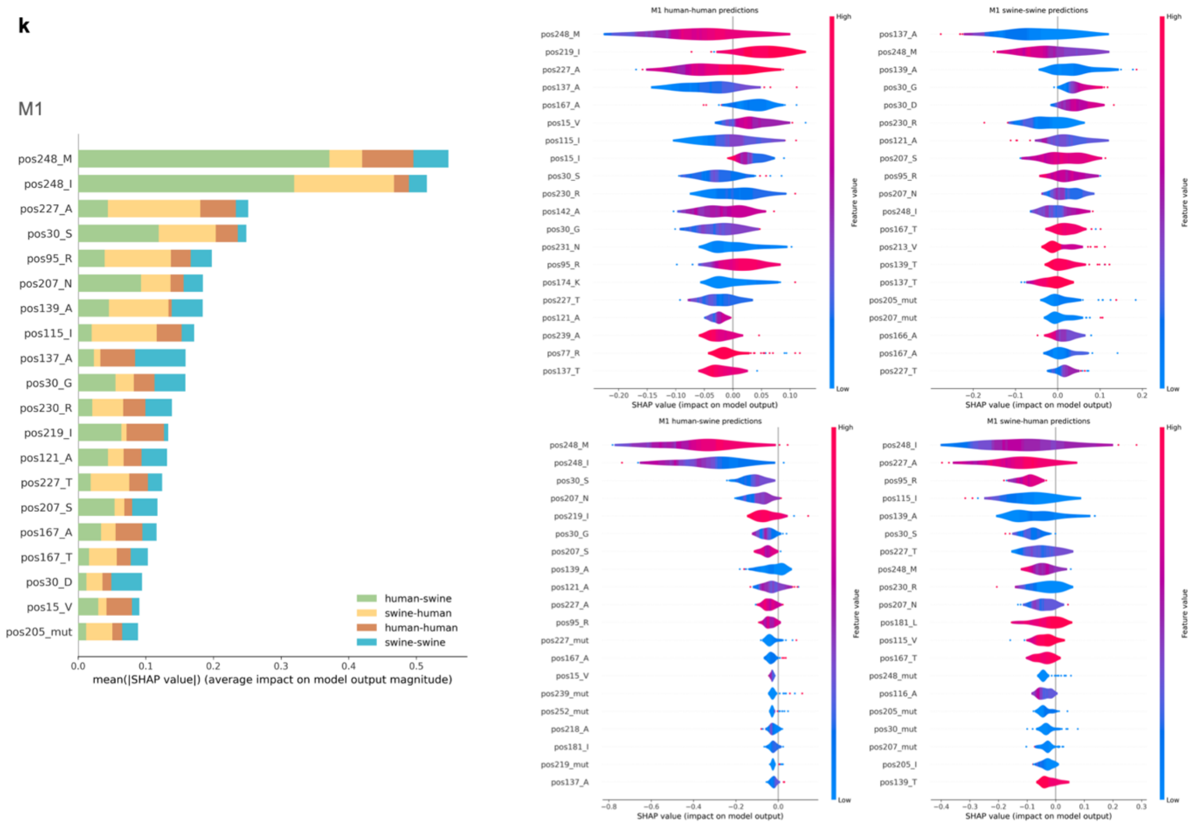
M1
pos248_M
pos248_I
pos227_A
pos30_S
pos95_R
pos207_N
pos139_A
pos115_I
pos137_A
pos30_G
pos230_R
pos219_I
pos121_A
pos227_T
pos207_S
pos167_A
pos167_T
pos30_D
pos15_V
pos205_mut
human-swine
swine-human
human-human
swine-swine
mean(|SHAP value|) (average impact on model output magnitude)
M1 human-human predictions
M1 swine-swine predictions
M1 human-swine predictions
M1 swine-human predictions
SHAP value (impact on model output)
Feature value
High
Low

**l**

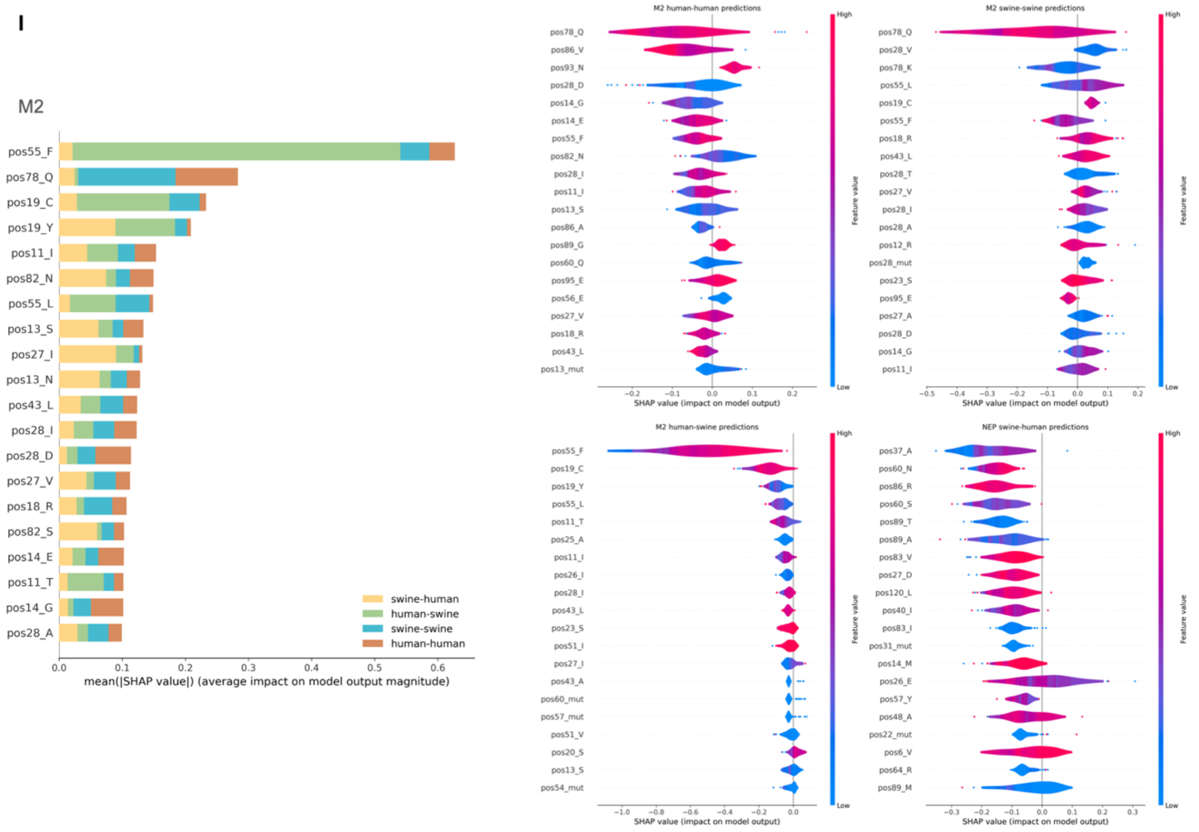
M2
pos55_F
pos78_Q
pos19_C
pos19_Y
pos11_I
pos82_N
pos55_L
pos13_S
pos27_I
pos13_N
pos43_L
pos28_I
pos28_D
pos27_V
pos18_R
pos82_S
pos14_E
pos11_T
pos14_G
pos28_A
swine-human
human-swine
swine-swine
human-human
mean(|SHAP value|) (average impact on model output magnitude)
M2 human-human predictions
M2 swine-swine predictions
M2 human-swine predictions
NEP swine-human predictions
SHAP value (impact on model output)
Feature value
High
Low

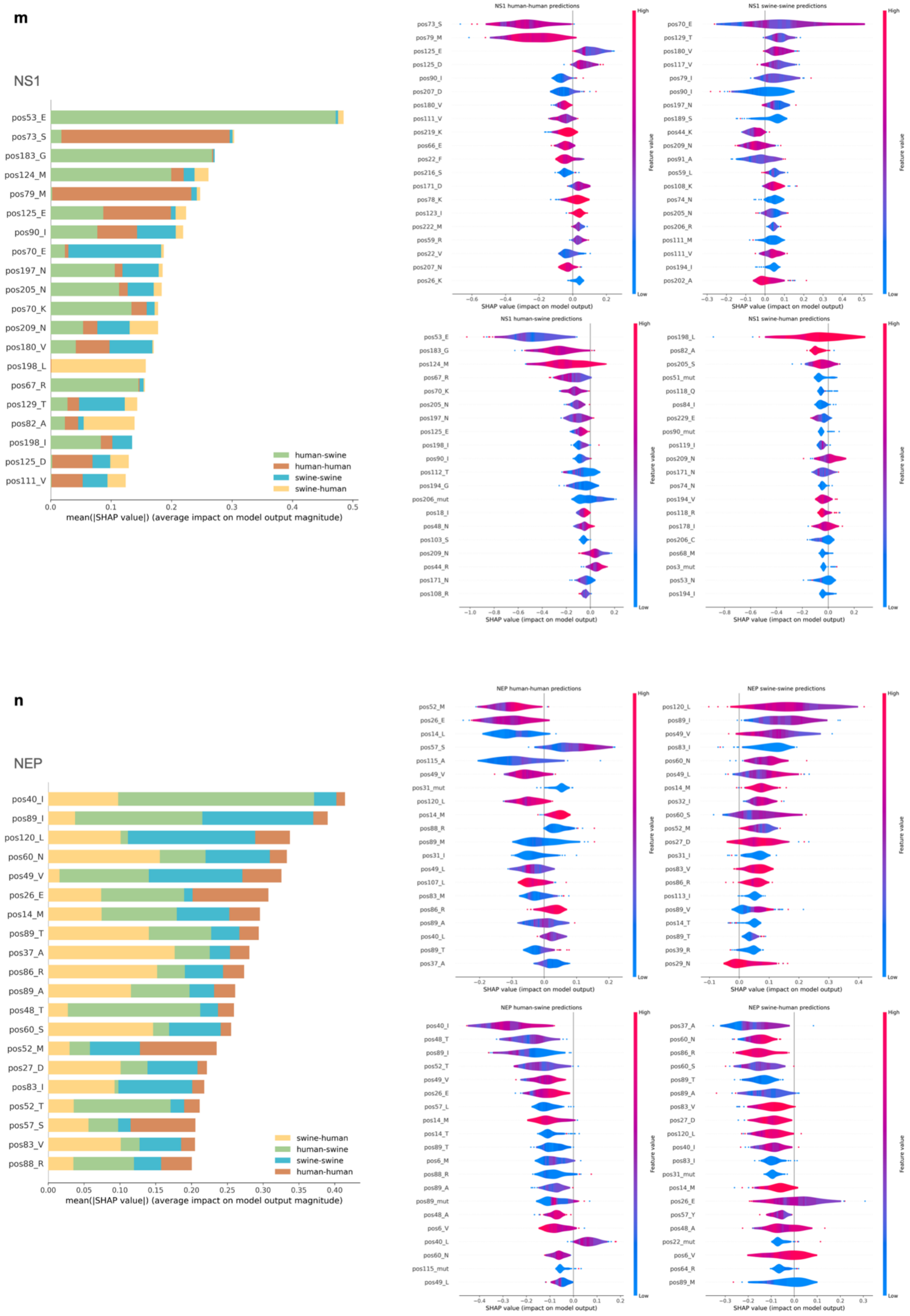
m
NS1
pos53_E
pos73_S
pos183_G
pos124_M
pos79_M
pos125_E
pos90_I
pos70_E
pos197_N
pos205_N
pos70_K
pos209_N
pos180_V
pos198_L
pos67_R
pos129_T
pos82_A
pos198_I
pos125_D
pos111_V
human-swine
human-human
swine-swine
swine-human
mean(|SHAP value|) (average impact on model output magnitude)
NS1 human-human predictions
NS1 swine-swine predictions
NS1 human-swine predictions
NS1 swine-human predictions
SHAP value (impact on model output)
Feature value
High
Low
n
NEP
pos40_I
pos89_I
pos120_L
pos60_N
pos49_V
pos26_E
pos14_M
pos89_T
pos37_A
pos86_R
pos89_A
pos48_T
pos60_S
pos52_M
pos27_D
pos83_I
pos52_T
pos57_S
pos83_V
pos88_R
swine-human
human-swine
swine-swine
human-human
mean(|SHAP value|) (average impact on model output magnitude)
NEP human-human predictions
NEP swine-swine predictions
NEP human-swine predictions
NEP swine-human predictions
SHAP value (impact on model output)

**Table S4**: Amino acid summary for positions included in the top 5 most important features from the XGBoost models based on the SHAP summary statistics. For each feature, the distribution of amino acids at the specific position is listed for all observed sequences from either humans or swine (external nodes in phylogenetic trees) as well as for each transmission type in our final datasets along with the frequency of mutations observed. NA indicates that there were no mutations at the specific position for branches belonging to the specific transmission category.

| Segment | Position | Human samples Amino Acids (%) | Swine samples Amino Acids (%) | Transmission | Amino Acids (%) | Mutations (%) |
|---|---|---|---|---|---|---|
| PB2 | 185 | I(98.1%), V(1.9%) | I(95.2%), V(4.7%), L(0.1%) | human-human | I(98%), V(2%) | NA |
| | | | | human-swine | I(98.6%), V(1.4%) | I-V(1.4%) |
| | | | | swine-human | I(100%) | NA |
| | | | | swine-swine | I(95.8%), V(4.1%), L(0.1%) | I-V(1.1%), I-L(0.1%) |
| | 251 | R(93.9%), K(6.1%) | R(61.5%), K(38%), E(0.1%), G(0.1%), I(0.1%) | human-human | R(98.5%), K(1.5%) | R-K(0.5%) |
| | | | | human-swine | R(94.6%), K(5.4%) | R-K(5.4%) |
| | | | | swine-human | R(76.9%), K(23.1%) | K-R(2.6%) |
| | | | | swine-swine | R(58.5%), K(41.2%), E(0.1%), G(0.1%), I(0.1%) | R-K(1.8%), K-R(0.5%), K-E(0.1%), R-G(0.1%), R-I(0.1%) |
| | 340 | K(62.9%), R(36.6%), N(0.5%) | K(55.7%), R(39.9%), N(3.3%), D(0.4%), E(0.1%), G(0.1%), M(0.1%), S(0.1%), T(0.1%) | human-human | K(68.2%), R(31.8%) | K-R(2%), R-K(0.2%) |
| | | | | human-swine | K(78.4%), R(21.6%) | K-R(2.7%), R-K(1.4%) |
| | | | | swine-human | K(53.8%), R(43.6%), N(2.6%) | K-R(2.6%), N-K(2.6%) |
| | | | | swine-swine | K(52%), R(43.9%), N(3.3%), D(0.4%), E(0.1%), G(0.1%), M(0.1%), S(0.1%), T(0.1%) | R-K(1.6%), K-R(0.8%), K-N(0.3%), K-E(0.1%), K-T(0.1%), N-D(0.1%), N-K(0.1%), N-S(0.1%), R-G(0.1%), R-M(0.1%) |
| | 344 | V(78.4%), M(21.1%), L(0.5%) | V(87.9%), M(10.8%), L(0.7%), I(0.6%) | human-human | V(74.2%), M(25.5%), L(0.2%) | V-M(1%), M-V(0.5%), V-L(0.2%) |
| | | | | human-swine | V(64.9%), M(35.1%) | V-M(2.7%) |
| | | | | swine-human | V(97.4%), M(2.6%) | M-V(2.6%) |
| | | | | swine-swine | V(92.8%), M(6.5%), L(0.4%), I(0.3%) | V-M(1.6%), V-I(0.3%), V-L(0.3%), M-L(0.1%) |
| | 432 | H(100%) | H(99%), Y(0.9%), P(0.1%) | human-human | H(100%) | NA |
| | | | | human-swine | H(95.9%), Y(2.7%), P(1.4%) | H-Y(2.7%), H-P(1.4%) |
| | | | | swine-human | H(100%) | NA |
| | | | | swine-swine | H(99.3%), Y(0.7%) | NA |
| | 453 | S(39.9%), P(33.8%), H(19.2%), T(7%) | P(67.2%), S(27.5%), H(2.7%), T(1.3%), F(0.9%), Y(0.3%), N(0.1%) | human-human | S(47%), P(22.8%), H(22.5%), T(7.8%) | P-H(0.5%), P-S(0.5%), S-T(0.5%), H-P(0.2%), P-T(0.2%), S-P(0.2%) |
| | | | | human-swine | S(59.5%), P(21.6%), H(10.8%), T(5.4%), N(1.4%), Y(1.4%) | S-P(6.8%), H-N(1.4%), S-Y(1.4%), T-P(1.4%) |
| | | | | swine-human | P(79.5%), S(20.5%) | P-S(2.6%) |
| | | | | swine-swine | P(73.6%), S(23.5%), H(1.5%), F(0.8%), T(0.5%), Y(0.1%) | S-P(1.4%), P-S(0.9%), P-T(0.3%), S-F(0.2%), P-H(0.1%), S-Y(0.1%) |
| | 456 | N(67.6%), S(30.5%), D(1.4%), T(0.5%) | N(65.7%), S(24.4%), D(9.7%), A(0.3%) | human-human | N(71%), S(28.7%), D(0.2%) | N-S(0.5%), S-N(0.5%) |
| | | | | human-swine | N(73%), S(25.7%), D(1.4%) | N-S(2.7%), S-N(2.7%) |
| | | | | swine-human | N(48.7%), S(41%), D(7.7%), T(2.6%) | N-D(2.6%), N-T(2.6%) |
| | | | | swine-swine | N(66.1%), S(23.8%), D(9.9%), A(0.2%) | N-D(1.6%), N-S(0.6%), S-N(0.2%), N-A(0.2%), D-N(0.1%) |

| | | | | | | |
|---|---|---|---|---|---|---|
| | 505 | R(97.2%), Q(1.9%), K(0.9%) | R(95.3%), K(4.4%), Q(0.1%), W(0.1%) | human-human | R(97.5%), Q(2%), K(0.5%) | NA |
| | | | | human-swine | R(98.6%), K(1.4%) | R-K(1.4%) |
| | | | | swine-human | R(97.4%), K(2.6%) | K-R(2.6%) |
| | | | | swine-swine | R(95.3%), K(4.4%), Q(0.2%), W(0.1%) | R-K(0.6%), K-R(0.1%), Q-K(0.1%), R-Q(0.1%), R-W(0.1%) |
| | 660 | K(93%), R(7%) | K(93.9%), R(6.1%) | human-human | K(92.5%), R(7.5%) | K-R(0.5%) |
| | | | | human-swine | K(83.8%), R(16.2%) | K-R(5.4%), R-K(1.4%) |
| | | | | swine-human | K(94.9%), R(5.1%) | K-R(2.6%) |
| | | | | swine-swine | K(95.5%), R(4.5%) | K-R(1%) |
| | 684 | S(62.4%), A(32.4%), G(2.3%), T(1.9%), X(0.9%) | A(61.3%), S(32.1%), T(3.8%), G(2.1%), N(0.6%), P(0.1%) | human-human | S(72.8%), A(23.8%), T(2%), G(1%), X(0.5%) | A-S(0.5%), S-X(0.5%) |
| | | | | human-swine | S(78.4%), A(20.3%), T(1.4%) | A-T(1.4%) |
| | | | | swine-human | A(71.8%), S(20.5%), G(7.7%) | T-A(2.6%) |
| | | | | swine-swine | A(69.1%), S(25.1%), T(2.8%), G(2.4%), N(0.5%), P(0.1%) | A-T(1.5%), A-S(0.4%), A-G(0.2%), A-P(0.1%), G-S(0.1%), N-G(0.1%) |
| **PB1** | 152 | S(85%), L(8.3%), M(4.7%), T(1%), K(0.5%), V(0.5%) | S(51.1%), M(25.8%), L(19.2%), K(1.9%), T(1.3%), A(0.2%), E(0.2%), F(0.2%), V(0.2%) | human-human | S(88.7%), L(8.9%), M(2.4%) | NA |
| | | | | human-swine | S(88%), L(9.8%), M(2.2%) | S-L(2.2%), S-M(1.1%) |
| | | | | swine-human | S(68.8%), M(12.5%), L(6.2%), T(6.2%), K(3.1%), V(3.1%) | M-V(3.1%) |
| | | | | swine-swine | S(43.2%), M(30.7%), L(22.1%), K(2%), T(1.5%), A(0.1%), E(0.1%), F(0.1%), V(0.1%) | L-S(0.4%), S-L(0.4%), M-T(0.3%), L-M(0.2%), K-M(0.1%), L-F(0.1%), M-V(0.1%), S-A(0.1%), S-T(0.1%), T-E(0.1%) |
| | 168 | K(99.5%), R(0.5%) | K(98.8%), R(1%), E(0.2%) | human-human | K(100%) | NA |
| | | | | human-swine | K(98.9%), R(1.1%) | K-R(1.1%) |
| | | | | swine-human | K(96.9%), R(3.1%) | NA |
| | | | | swine-swine | K(98.8%), R(1.1%), E(0.1%) | K-E(0.1%), K-R(0.1%) |
| | 175 | D(64.8%), N(34.7%), E(0.5%) | D(82.6%), N(16.9%), S(0.3%), E(0.2%) | human-human | D(58.7%), N(41.1%), E(0.3%) | D-N(1.6%), D-E(0.3%), N-D(0.3%) |
| | | | | human-swine | D(52.2%), N(46.7%), E(1.1%) | N-D(3.3%), D-E(1.1%) |
| | | | | swine-human | D(90.6%), N(9.4%) | D-N(3.1%) |
| | | | | swine-swine | D(89.3%), N(10.5%), S(0.2%) | D-N(0.3%), N-D(0.3%), N-S(0.2%) |
| | 179 | I(53.4%), M(40.9%), V(5.7%) | M(54.5%), I(43.8%), V(1.7%) | human-human | I(58.9%), M(35.5%), V(5.5%) | I-M(0.8%), M-I(0.5%), M-V(0.3%) |
| | | | | human-swine | I(79.3%), M(19.6%), V(1.1%) | I-M(4.3%) |
| | | | | swine-human | I(59.4%), M(40.6%) | I-M(3.1%) |
| | | | | swine-swine | M(61.7%), I(36.8%), V(1.4%) | I-M(2%), M-I(1%), M-V(0.5%), V-M(0.2%) |
| | 216 | S(49.7%), G(46.1%), N(2.6%), C(1%), A(0.5%) | S(56.5%), G(26.1%), N(14.5%), D(1%), C(0.7%), I(0.3%), V(0.3%), A(0.2%), R(0.2%), | human-human | G(56.3%), S(41.3%), C(1.1%), N(1.1%), A(0.3%) | G-S(1.8%), G-A(0.3%), G-C(0.3%), S-G(0.3%), S-N(0.3%) |
| | | | | human-swine | G(51.1%), S(42.4%), N(3.3%), D(2.2%), C(1.1%) | G-S(7.6%), G-D(2.2%), C-G(1.1%), G-C(1.1%), G-N(1.1%), S-G(1.1%), S-N(1.1%) |
| | | | | swine-human | S(71.9%), G(21.9%), N(6.2%) | NA |
| | | | | swine-swine | S(61.5%), G(21.4%), N(15.6%), D(0.5%), C(0.3%), I(0.2%), V(0.2%), A(0.1%), R(0.1%), T(0.1%) | G-S(1.6%), S-N(1.6%), S-G(0.5%), G-D(0.4%), N-S(0.3%), S-C(0.3%), D-N(0.1%), G-A(0.1%), G-V(0.1%), N-I(0.1%), S-I(0.1%), S-R(0.1%), S-T(0.1%), S-V(0.1%) |
| | 317 | M(91.7%), I(4.7%), V(3.6%) | | human-human | M(96.1%), I(2.1%), V(1.8%) | NA |

| | | | | | |
|---|---|---|---|---|---|
| | | M(72.5%), I(14%), V(12.5%), T(0.8%), D(0.2%) | human-swine | M(90.2%), V(6.5%), I(3.3%) | M-V(5.4%), M-I(3.3%) |
| | | | swine-human | M(78.1%), I(15.6%), V(6.2%) | M-V(6.2%) |
| | | | swine-swine | M(76%), I(13.4%), V(10%), T(0.5%), D(0.1%) | M-V(3.5%), M-I(3%), I-V(0.7%), M-T(0.4%), V-M(0.3%), I-T(0.1%), M-D(0.1%) |
| 336 | I(71%), V(29%) | V(50.9%), I(48.9%), T(0.2%) | human-human | I(76.6%), V(23.4%) | I-V(0.8%), V-I(0.3%) |
| | | | human-swine | I(87%), V(13%) | V-I(1.1%) |
| | | | swine-human | I(65.6%), V(34.4%) | NA |
| | | | swine-swine | V(59.4%), I(40.5%), T(0.1%) | I-V(0.6%), V-I(0.6%), I-T(0.1%) |
| 361 | R(56%), S(25.9%), K(11.9%), N(5.7%), I(0.5%) | S(43.2%), R(36.8%), K(10.6%), N(6.1%), I(2.4%), G(0.8%), T(0.2%) | human-human | R(65%), S(19.5%), K(8.2%), N(7.1%), I(0.3%) | R-K(1.6%), R-S(0.5%), R-I(0.3%) |
| | | | human-swine | R(69.6%), K(16.3%), N(6.5%), S(4.3%), I(2.2%), G(1.1%) | R-K(10.9%), K-R(1.1%), R-G(1.1%), S-R(1.1%) |
| | | | swine-human | R(53.1%), S(21.9%), K(15.6%), N(6.2%), I(3.1%) | S-R(3.1%) |
| | | | swine-swine | S(52.4%), R(30.5%), K(8.4%), N(5.8%), I(2.2%), G(0.5%), T(0.1%) | R-K(1.1%), S-N(0.3%), R-G(0.2%), R-I(0.2%), S-R(0.2%), N-T(0.1%), R-N(0.1%), S-G(0.1%), S-I(0.1%) |
| 374 | A(88.1%), S(9.3%), V(1.6%), E(0.5%), T(0.5%) | A(84.1%), S(11.5%), T(2.5%), V(1%), E(0.7%), I(0.2%) | human-human | A(86.3%), S(12.1%), V(1.6%) | A-S(0.5%) |
| | | | human-swine | A(70.7%), S(23.9%), E(2.2%), I(1.1%), T(1.1%), V(1.1%) | A-E(2.2%), A-S(2.2%), A-T(1.1%), V-I(1.1%) |
| | | | swine-human | A(81.2%), S(9.4%), E(3.1%), T(3.1%), V(3.1%) | A-E(3.1%), A-T(3.1%), A-V(3.1%) |
| | | | swine-swine | A(88%), S(9.3%), T(1.7%), V(0.7%), E(0.3%) | A-T(1.3%), A-V(0.3%), A-S(0.2%), A-E(0.1%) |
| 375 | S(79.8%), N(10.9%), D(6.2%), G(2.6%), E(0.5%) | S(50.8%), D(21.6%), N(17.9%), Y(5.1%), G(3.2%), T(0.7%), V(0.3%), C(0.2%), H(0.2%) | human-human | S(84.5%), N(10%), G(2.9%), D(2.6%) | S-N(1.1%), D-N(0.3%), N-D(0.3%) |
| | | | human-swine | S(87%), N(9.8%), D(2.2%), G(1.1%) | S-N(5.4%), D-N(1.1%), S-D(1.1%) |
| | | | swine-human | S(75%), D(15.6%), E(3.1%), G(3.1%), N(3.1%) | D-E(3.1%), D-S(3.1%), G-S(3.1%) |
| | | | swine-swine | S(42.7%), D(27.9%), N(18.9%), Y(6%), G(3.5%), T(0.5%), V(0.2%), C(0.1%), H(0.1%), K(0.1%) | D-N(1.7%), S-N(0.8%), D-G(0.5%), N-S(0.5%), N-D(0.4%), D-Y(0.3%), N-T(0.3%), G-D(0.2%), G-S(0.2%), Y-N(0.2%), D-H(0.1%), D-V(0.1%), N-K(0.1%), S-G(0.1%), S-V(0.1%), Y-C(0.1%), Y-S(0.1%) |
| 433 | K(77.2%), R(22.3%), N(0.5%) | K(72.8%), R(26.3%), N(0.8%) | human-human | K(82.6%), R(17.4%) | K-R(0.8%), R-K(0.3%) |
| | | | human-swine | K(71.7%), R(28.3%) | K-R(1.1%) |
| | | | swine-human | R(56.2%), K(40.6%), N(3.1%) | K-R(3.1%) |
| | | | swine-swine | K(73.2%), R(25.7%), N(1%) | K-R(0.7%), R-K(0.5%), K-N(0.1%) |
| 633 | S(88.1%), N(11.4%), C(0.5%) | S(64.6%), N(34.1%), T(0.5%), G(0.3%), R(0.3%), Y(0.2%) | human-human | S(95%), N(5%) | S-N(0.3%) |
| | | | human-swine | S(90.2%), N(6.5%), G(1.1%), R(1.1%), T(1.1%) | S-N(4.3%), S-G(1.1%), S-R(1.1%), S-T(1.1%) |
| | | | swine-human | S(65.6%), N(31.2%), C(3.1%) | S-C(3.1%), S-N(3.1%) |
| | | | swine-swine | S(60.5%), N(38.8%), T(0.4%), G(0.1%), R(0.1%), Y(0.1%) | S-N(1.4%), N-S(0.3%), N-Y(0.1%), S-G(0.1%), S-R(0.1%) |
| 741 | | S(50.4%), A(46.9%), | human-human | S(73.7%), A(15.3%), T(9.7%), V(1.3%) | A-T(0.3%), A-V(0.3%), S-A(0.3%), T-A(0.3%) |

| | | | | | | |
|---|---|---|---|---|---|---|
| | | S(68.9%), A(19.2%), T(9.8%), V(2.1%) | T(1.2%), V(1.2%), Y(0.3%) | human-swine | S(88%), A(9.8%), T(1.1%), Y(1.1%) | A-S(1.1%), S-Y(1.1%) |
| | | | | swine-human | S(71.9%), A(25%), V(3.1%) | NA |
| | | | | swine-swine | A(55.7%), S(42.2%), V(1.2%), T(0.7%), Y(0.1%) | A-T(0.5%), A-S(0.4%), S-A(0.4%), A-V(0.3%), S-Y(0.1%) |
| PB1-F2 | 4 | E(82.4%), G(17.1%), X(0.5%) | E(60.7%), G(39%), V(0.3%) | human-human | E(85.8%), G(13.9%), X(0.3%) | E-G(1.1%), G-E(1.1%), E-X(0.3%) |
| | | | | human-swine | E(88%), G(12%) | E-G(4.3%) |
| | | | | swine-human | E(78.1%), G(21.9%) | NA |
| | | | | swine-swine | E(53.1%), G(46.6%), V(0.3%) | G-E(2.6%), E-G(1.1%), E-V(0.1%) |
| | 7 | T(96.4%), I(3.1%), X(0.5%) | T(94.9%), I(5.1%) | human-human | T(98.7%), I(1.1%), X(0.3%) | T-X(0.3%) |
| | | | | human-swine | T(98.9%), I(1.1%) | T-I(1.1%) |
| | | | | swine-human | T(87.5%), I(12.5%) | NA |
| | | | | swine-swine | T(94.5%), I(5.5%) | T-I(0.6%), I-T(0.1%) |
| | 9 | W(96.9%), X(3.1%) | W(95.3%), X(3.9%), L(0.5%), C(0.2%), S(0.2%) | human-human | W(98.2%), X(1.8%) | W-X(0.3%) |
| | | | | human-swine | W(94.6%), X(4.3%), L(1.1%) | W-X(2.2%), W-L(1.1%) |
| | | | | swine-human | W(90.6%), X(9.4%) | W-X(3.1%) |
| | | | | swine-swine | W(97%), X(2.5%), L(0.2%), C(0.1%), S(0.1%) | W-X(1.5%), W-L(0.2%), W-C(0.1%), W-S(0.1%) |
| | 10 | T(78.2%), I(18.7%), X(3.1%) | T(87%), I(8.8%), X(4%), S(0.2%) | human-human | T(78.4%), I(19.7%), X(1.8%) | I-T(0.3%), T-I(0.3%), T-X(0.3%) |
| | | | | human-swine | T(85.9%), I(9.8%), X(4.3%) | T-I(2.2%), T-X(2.2%), I-T(1.1%) |
| | | | | swine-human | T(84.4%), X(9.4%), I(6.2%) | T-X(3.1%) |
| | | | | swine-swine | T(89.5%), I(7.7%), X(2.6%), S(0.1%) | T-X(1.4%), T-I(1.1%), I-X(0.2%), T-S(0.1%) |
| | 11 | Q(78.8%), L(11.9%), R(5.7%), X(3.1%), P(0.5%) | Q(54.1%), R(33.4%), L(7.1%), X(4.2%), P(1.2%) | human-human | Q(81.8%), L(13.2%), R(3.2%), X(1.8%) | L-Q(0.5%), Q-X(0.3%) |
| | | | | human-swine | Q(81.5%), L(7.6%), R(5.4%), X(4.3%), P(1.1%) | Q-R(3.3%), Q-X(2.2%), Q-L(1.1%), Q-P(1.1%) |
| | | | | swine-human | Q(71.9%), R(9.4%), X(9.4%), L(6.2%), P(3.1%) | Q-P(3.1%), Q-X(3.1%) |
| | | | | swine-swine | Q(47.1%), R(42.5%), L(6.9%), X(2.7%), P(0.7%) | R-Q(3%), Q-X(0.9%), R-L(0.7%), Q-R(0.5%), R-X(0.5%), L-X(0.3%), Q-P(0.3%), Q-L(0.2%), L-P(0.1%), L-Q(0.1%), L-R(0.1%), R-P(0.1%) |
| | 16 | X(41.5%), T(34.7%), I(23.8%) | I(38.4%), X(31.9%), T(29.7%) | human-human | X(47.9%), T(32.1%), I(20%) | T-I(0.8%), I-T(0.5%), T-X(0.5%) |
| | | | | human-swine | X(62%), T(29.3%), I(8.7%) | T-I(1.1%) |
| | | | | swine-human | T(53.1%), X(28.1%), I(18.8%) | T-X(6.2%), I-T(3.1%) |
| | | | | swine-swine | I(46.9%), T(29.8%), X(23.3%) | I-T(1.8%), T-X(1.5%), I-X(1.1%), T-I(0.7%), X-T(0.1%) |
| | 27 | X(42%), T(39.4%), I(18.7%) | T(55.8%), X(33.6%), I(10.3%), A(0.3%) | human-human | X(48.2%), T(30%), I(21.8%) | I-X(0.8%), I-T(0.3%), T-I(0.3%) |
| | | | | human-swine | X(62%), I(21.7%), T(16.3%) | T-I(1.1%) |
| | | | | swine-human | T(65.6%), X(28.1%), I(6.2%) | T-X(6.2%), T-I(3.1%) |
| | | | | swine-swine | T(66.9%), X(24.3%), I(8.5%), A(0.2%) | T-X(3.1%), T-I(1%), I-X(0.7%), I-T(0.5%), T-A(0.2%), X-T(0.2%) |
| | 34 | X(42%), N(32.1%), S(25.9%) | S(58%), X(33.6%), N(8.4%) | human-human | X(48.2%), N(31.8%), S(20%) | S-X(0.5%), N-S(0.3%), N-X(0.3%), S-N(0.3%) |
| | | | | human-swine | X(62%), S(25%), N(13%) | N-S(1.1%) |
| | | | | swine-human | S(65.6%), X(28.1%), N(6.2%) | S-X(6.2%) |

| | | | | | |
|---|---|---|---|---|---|
| | | | swine-swine | S(69.8%), X(24.3%), N(5.9%) | S-X(3.8%), S-N(1.4%), X-N(0.1%), X-S(0.1%) |
| | 45 | X(41.5%), I(38.3%), T(20.2%) | T(37.1%), X(36.3%), I(26.5%), A(0.2%) | human-human | X(47.9%), I(37.1%), T(15%) | I-X(0.5%) |
| | | | | human-swine | X(64.1%), I(33.7%), T(2.2%) | I-X(1.1%), T-X(1.1%) |
| | | | | swine-human | I(46.9%), X(28.1%), T(25%) | I-X(6.2%) |
| | | | | swine-swine | T(47.1%), X(26.8%), I(26%), A(0.1%) | T-X(2.5%), I-X(1.8%), I-T(0.8%), T-I(0.8%), X-I(0.2%), T-A(0.1%) |
| | 73 | X(52.3%), R(32.6%), K(15%) | X(44.9%), K(31.5%), R(22.8%), I(0.7%), T(0.2%) | human-human | X(56.8%), R(33.2%), K(10%) | R-X(1.1%), K-X(0.5%), R-K(0.5%), K-R(0.3%) |
| | | | | human-swine | X(67.4%), R(30.4%), K(2.2%) | R-X(3.3%), K-X(1.1%) |
| | | | | swine-human | R(46.9%), X(31.2%), K(21.9%) | K-R(6.2%), R-K(3.1%), R-X(3.1%) |
| | | | | swine-swine | K(42.1%), X(34.5%), R(22.7%), I(0.5%), T(0.2%) | K-X(4.1%), R-X(2.2%), K-R(0.5%), R-K(0.3%), R-I(0.2%), K-I(0.1%), K-T(0.1%), T-X(0.1%), X-K(0.1%), X-R(0.1%) |
| | 87 | X(60.1%), G(28.5%), E(11.4%) | X(55.1%), E(23.9%), G(20.9%) | human-human | X(63.2%), G(28.4%), E(8.4%) | G-X(1.3%), E-X(0.5%), G-E(0.3%) |
| | | | | human-swine | X(77.2%), G(20.7%), E(2.2%) | G-X(3.3%), E-X(1.1%), G-E(1.1%) |
| | | | | swine-human | X(46.9%), G(40.6%), E(12.5%) | E-G(3.1%), G-X(3.1%), X-E(3.1%) |
| | | | | swine-swine | X(43.6%), E(33.9%), G(22.5%) | E-X(5.3%), G-X(2.7%), E-G(0.8%), G-E(0.7%), X-E(0.1%), X-G(0.1%) |
| **PA** | 38 | I(99.1%), X(0.9%) | I(98.1%), V(1.5%), T(0.2%), X(0.2%) | human-human | I(99.5%), X(0.5%) | I-X(0.5%) |
| | | | | human-swine | I(98.9%), V(1.1%) | I-V(1.1%) |
| | | | | swine-human | I(100%) | NA |
| | | | | swine-swine | I(98.7%), V(1.1%), T(0.1%), X(0.1%) | I-T(0.1%), I-V(0.1%), I-X(0.1%) |
| | 44 | V(95.3%), I(3.7%), X(0.9%) | V(91.2%), I(7.9%), A(0.6%), M(0.2%), X(0.2%) | human-human | V(97.7%), I(1.8%), X(0.5%) | V-X(0.5%), V-I(0.2%) |
| | | | | human-swine | V(96.6%), I(3.4%) | V-I(1.1%) |
| | | | | swine-human | V(89.3%), I(10.7%) | NA |
| | | | | swine-swine | V(92.2%), I(7.2%), A(0.4%), M(0.1%), X(0.1%) | V-I(1.4%), V-A(0.3%), I-V(0.1%), V-M(0.1%), V-X(0.1%) |
| | 61 | I(92.6%), T(6%), M(0.5%), V(0.5%), X(0.5%) | I(71.1%), T(21.6%), V(5.9%), M(0.8%), K(0.5%), R(0.2%) | human-human | I(97.7%), T(1.6%), M(0.2%), V(0.2%), X(0.2%) | I-M(0.2%), I-V(0.2%), I-X(0.2%) |
| | | | | human-swine | I(94.3%), V(3.4%), M(1.1%), T(1.1%) | I-M(1.1%), I-V(1.1%) |
| | | | | swine-human | I(67.9%), T(28.6%), V(3.6%) | NA |
| | | | | swine-swine | I(69.3%), T(25.8%), V(4.1%), K(0.4%), M(0.4%), R(0.1%) | I-V(1.4%), I-T(0.5%), T-I(0.4%), I-M(0.2%), I-R(0.1%), T-K(0.1%), V-I(0.1%) |
| | 85 | T(53%), I(41.9%), N(4.2%), S(0.5%), X(0.5%) | T(59.1%), I(30.4%), N(5.9%), A(3.4%), V(0.6%), S(0.5%), G(0.2%) | human-human | I(52.6%), T(43.3%), N(3.8%), X(0.2%) | I-T(0.2%), T-I(0.2%), T-X(0.2%) |
| | | | | human-swine | I(69.3%), T(22.7%), N(6.8%), V(1.1%) | I-T(2.3%), I-V(1.1%), T-N(1.1%) |
| | | | | swine-human | T(78.6%), I(14.3%), N(3.6%), S(3.6%) | N-S(3.6%) |
| | | | | swine-swine | T(70.1%), I(21.5%), N(5.3%), A(2.5%), S(0.3%), V(0.3%), G(0.1%) | T-A(1.3%), T-N(0.8%), T-I(0.4%), I-T(0.3%), I-V(0.3%), N-S(0.2%), I-A(0.1%), T-G(0.1%), T-S(0.1%) |
| | 208 | T(82.3%), K(13.5%), S(3.7%), R(0.5%) | T(54.2%), K(42.1%), R(1.2%), A(1.1%), S(0.8%), I(0.3%), N(0.3%) | human-human | T(88.3%), K(7.4%), S(4.3%) | T-K(0.2%), T-S(0.2%) |
| | | | | human-swine | T(81.8%), K(13.6%), A(2.3%), S(2.3%) | T-K(4.5%), T-A(2.3%) |
| | | | | swine-human | T(53.6%), K(42.9%), R(3.6%) | K-R(3.6%) |

| | | | | | | |
|---|---|---|---|---|---|---|
| | | | | swine-swine | K(49.4%), T(48.3%), R(1%), A(0.6%), S(0.4%), I(0.2%), N(0.2%) | K-R(0.5%), T-K(0.5%), K-T(0.3%), T-A(0.3%), T-I(0.2%), K-N(0.1%), R-K(0.1%), T-N(0.1%) |
| | 267 | P(93.5%), S(5.1%), L(0.9%), H(0.5%) | P(74.4%), S(21.5%), L(3.4%), H(0.3%), A(0.2%), Q(0.2%), Y(0.2%) | human-human | P(98.4%), S(1.1%), L(0.5%) | S-P(0.2%) |
| | | | | human-swine | P(100%) | NA |
| | | | | swine-human | P(64.3%), S(32.1%), H(3.6%) | P-H(3.6%) |
| | | | | swine-swine | P(68.7%), S(27.7%), L(3.1%), H(0.2%), A(0.1%), Q(0.1%), Y(0.1%) | S-P(1.7%), P-L(1%), P-S(0.8%), P-H(0.2%), L-S(0.1%), P-Q(0.1%), S-A(0.1%), S-Y(0.1%) |
| | 275 | P(57.2%), L(40.5%), R(1.9%), S(0.5%) | P(61.1%), L(37.8%), A(0.3%), H(0.3%), R(0.3%), F(0.2%) | human-human | L(51.9%), P(47%), R(0.9%), S(0.2%) | L-P(0.5%), P-S(0.2%) |
| | | | | human-swine | L(70.5%), P(28.4%), R(1.1%) | L-P(3.4%), L-R(1.1%), P-L(1.1%) |
| | | | | swine-human | P(78.6%), L(21.4%) | NA |
| | | | | swine-swine | P(69.3%), L(30.1%), A(0.2%), H(0.2%), F(0.1%), R(0.1%) | P-L(1.2%), L-P(1.1%), P-A(0.2%), P-H(0.2%), L-F(0.1%), P-R(0.1%) |
| | 354 | I(97.7%), V(1.9%), T(0.5%) | I(93.2%), T(3.1%), V(2.9%), L(0.6%), F(0.2%) | human-human | I(98.2%), V(1.6%), T(0.2%) | I-T(0.2%) |
| | | | | human-swine | I(98.9%), V(1.1%) | I-V(1.1%) |
| | | | | swine-human | I(100%) | V-I(3.6%) |
| | | | | swine-swine | I(94%), T(2.8%), V(2.2%), L(0.9%), F(0.1%) | I-V(0.8%), I-T(0.5%), I-L(0.1%), T-F(0.1%), V-I(0.1%) |
| | 362 | K(51.2%), R(48.8%) | K(55.1%), R(44.9%) | human-human | R(56.2%), K(43.8%) | R-K(0.7%) |
| | | | | human-swine | R(79.5%), K(20.5%) | R-K(4.5%) |
| | | | | swine-human | K(53.6%), R(46.4%) | R-K(3.6%) |
| | | | | swine-swine | K(61.3%), R(38.7%) | R-K(2.1%), K-R(0.3%) |
| | 407 | V(59.1%), I(40.5%), M(0.5%) | V(56.2%), I(43.8%) | human-human | V(63.4%), I(36.3%), M(0.2%) | V-I(0.5%), I-M(0.2%), I-V(0.2%) |
| | | | | human-swine | V(80.7%), I(19.3%) | V-I(6.8%) |
| | | | | swine-human | V(67.9%), I(32.1%) | I-V(3.6%), V-I(3.6%) |
| | | | | swine-swine | V(50.3%), I(49.7%) | I-V(1.7%), V-I(1.7%) |
| | 497 | K(89.3%), R(10.7%) | K(91.5%), R(8.2%), I(0.2%), T(0.2%) | human-human | K(91%), R(9%) | NA |
| | | | | human-swine | K(87.5%), R(12.5%) | NA |
| | | | | swine-human | K(82.1%), R(17.9%) | NA |
| | | | | swine-swine | K(93.1%), R(6.7%), I(0.1%), T(0.1%) | K-R(0.6%), K-I(0.1%), K-T(0.1%) |
| | 505 | I(82.3%), V(17.2%), X(0.5%) | I(61.6%), V(38.4%) | human-human | I(90.7%), V(9.3%) | I-V(0.5%), V-I(0.2%) |
| | | | | human-swine | I(92%), V(8%) | I-V(3.4%) |
| | | | | swine-human | V(50%), I(46.4%), X(3.6%) | I-X(3.6%) |
| | | | | swine-swine | I(53.3%), V(46.7%) | V-I(1.1%), I-V(0.4%) |
| | 581 | M(96.3%), L(3.7%) | M(99.4%), V(0.3%), I(0.2%), L(0.2%) | human-human | M(96.6%), L(3.4%) | M-L(0.2%) |
| | | | | human-swine | M(98.9%), L(1.1%) | NA |
| | | | | swine-human | M(100%) | NA |
| | | | | swine-swine | M(99.7%), V(0.2%), I(0.1%) | M-V(0.2%), M-I(0.1%) |
| **PA-X** | 44 | V(95.3%), I(3.7%), X(0.9%) | V(91.2%), I(7.9%), A(0.6%), M(0.2%), X(0.2%) | human-human | V(97.7%), I(1.8%), X(0.5%) | V-X(0.5%), V-I(0.2%) |
| | | | | human-swine | V(96.6%), I(3.4%) | V-I(1.1%) |
| | | | | swine-human | V(89.3%), I(10.7%) | NA |
| | | | | swine-swine | V(92.2%), I(7.2%), A(0.4%), M(0.1%), X(0.1%) | V-I(1.4%), V-A(0.3%), I-V(0.1%), V-M(0.1%), V-X(0.1%) |
| | 61 | I(92.6%), T(6%), M(0.5%), V(0.5%), X(0.5%) | I(71.1%), T(21.6%), V(5.9%), M(0.8%), K(0.5%), R(0.2%) | human-human | I(97.7%), T(1.6%), M(0.2%), V(0.2%), X(0.2%) | I-M(0.2%), I-V(0.2%), I-X(0.2%) |
| | | | | human-swine | I(94.3%), V(3.4%), M(1.1%), T(1.1%) | I-M(1.1%), I-V(1.1%) |
| | | | | swine-human | I(67.9%), T(28.6%), V(3.6%) | NA |

| | | | | | |
|---|---|---|---|---|---|
| | | | swine-swine | I(69.3%), T(25.8%), V(4.1%), K(0.4%), M(0.4%), R(0.1%) | I-V(1.4%), I-T(0.5%), T-I(0.4%), I-M(0.2%), I-R(0.1%), T-K(0.1%), V-I(0.1%) |
| 66 | G(67.9%), D(19.5%), S(6%), E(5.6%), X(0.9%) | G(79.8%), S(13.1%), D(5.6%), N(0.6%), A(0.3%), E(0.3%), I(0.2%), X(0.2%) | human-human | G(71.3%), D(19.4%), E(5.6%), S(3.2%), X(0.5%) | G-D(0.7%), D-X(0.5%), D-E(0.2%), G-S(0.2%) |
| | | | human-swine | G(84.1%), D(10.2%), S(2.3%), A(1.1%), E(1.1%), X(1.1%) | G-D(3.4%), G-A(1.1%), G-S(1.1%), G-X(1.1%) |
| | | | swine-human | G(89.3%), S(7.1%), D(3.6%) | NA |
| | | | swine-swine | G(79.5%), S(15.4%), D(4.2%), N(0.5%), A(0.2%), E(0.1%), I(0.1%) | G-S(0.8%), G-D(0.7%), S-G(0.3%), S-N(0.2%), D-E(0.1%), D-G(0.1%), S-I(0.1%) |
| 85 | T(53%), I(41.9%), N(4.2%), S(0.5%), X(0.5%) | T(59.1%), I(30.4%), N(5.9%), A(3.4%), V(0.6%), S(0.5%), G(0.2%) | human-human | I(52.6%), T(43.3%), N(3.8%), X(0.2%) | I-T(0.2%), T-I(0.2%), T-X(0.2%) |
| | | | human-swine | I(69.3%), T(22.7%), N(6.8%), V(1.1%) | I-T(2.3%), I-V(1.1%), T-N(1.1%) |
| | | | swine-human | T(78.6%), I(14.3%), N(3.6%), S(3.6%) | N-S(3.6%) |
| | | | swine-swine | T(70.1%), I(21.5%), N(5.3%), A(2.5%), S(0.3%), V(0.3%), G(0.1%) | T-A(1.3%), T-N(0.8%), T-I(0.4%), I-T(0.3%), I-V(0.3%), N-S(0.2%), I-A(0.1%), T-G(0.1%), T-S(0.1%) |
| 184 | S(74%), N(14%), I(8.4%), G(2.3%), T(1.4%) | S(59.7%), I(32.6%), N(3.4%), T(2.6%), G(0.6%), L(0.6%), V(0.3%), A(0.2%) | human-human | S(80.4%), N(14.7%), I(2.9%), G(1.6%), T(0.5%) | N-S(0.2%), S-N(0.2%) |
| | | | human-swine | S(88.6%), N(6.8%), G(3.4%), I(1.1%) | S-N(2.3%), S-G(1.1%) |
| | | | swine-human | S(53.6%), I(35.7%), T(7.1%), G(3.6%) | S-G(3.6%) |
| | | | swine-swine | S(52.7%), I(41.3%), T(2.8%), N(2.4%), L(0.4%), V(0.3%), A(0.1%), G(0.1%) | I-T(0.6%), I-L(0.4%), S-N(0.4%), I-S(0.3%), T-I(0.3%), S-I(0.2%), I-V(0.1%), N-S(0.1%), S-G(0.1%), T-A(0.1%) |
| 206 | K(59.5%), R(38.6%), T(1.9%) | K(64%), R(34.6%), T(0.9%), N(0.3%), Q(0.2%) | human-human | R(49.7%), K(49%), T(1.4%) | R-K(0.7%) |
| | | | human-swine | R(65.9%), K(34.1%) | R-K(6.8%) |
| | | | swine-human | K(75%), R(25%) | K-R(3.6%), R-K(3.6%) |
| | | | swine-swine | K(72%), R(26.7%), T(1.1%), N(0.2%), Q(0.1%) | K-R(1.2%), R-K(1.1%), K-N(0.2%), K-Q(0.1%), K-T(0.1%) |
| 208 | Q(82.3%), R(8.8%), K(5.1%), P(2.3%), L(0.9%), E(0.5%) | Q(52.6%), R(31.9%), K(12.2%), G(1.2%), P(0.8%), X(0.8%), M(0.3%), L(0.2%) | human-human | Q(90.5%), K(4.1%), R(3.4%), P(1.6%), L(0.5%) | Q-K(0.2%) |
| | | | human-swine | Q(81.8%), K(11.4%), R(3.4%), P(2.3%), X(1.1%) | Q-K(4.5%), K-R(1.1%), Q-R(1.1%), Q-X(1.1%) |
| | | | swine-human | Q(46.4%), R(35.7%), K(10.7%), E(3.6%), P(3.6%) | R-E(3.6%) |
| | | | swine-swine | Q(46.8%), R(40.9%), K(10.2%), G(1%), P(0.4%), X(0.4%), M(0.2%), L(0.1%) | R-K(1.8%), Q-R(0.5%), R-G(0.5%), Q-K(0.4%), Q-X(0.3%), R-Q(0.3%), R-M(0.2%), G-R(0.1%), Q-L(0.1%), Q-P(0.1%) |
| 212 | A(80.9%), I(8.8%), E(5.1%), V(3.3%), S(1.4%), L(0.5%) | A(51.1%), I(22.2%), V(12.7%), E(4.9%), S(2.8%), T(2.8%), L(1.9%), X(1.2%), G(0.3%) | human-human | A(89.8%), E(4.5%), I(3.4%), V(1.8%), S(0.5%) | A-V(0.5%), A-E(0.2%) |
| | | | human-swine | A(85.2%), E(8%), I(3.4%), S(1.1%), V(1.1%), X(1.1%) | A-E(2.3%), A-I(2.3%), A-S(1.1%), A-V(1.1%), A-X(1.1%) |
| | | | swine-human | A(46.4%), I(35.7%), S(7.1%), E(3.6%), L(3.6%), V(3.6%) | NA |
| | | | swine-swine | A(44.4%), I(26.5%), V(16.3%), E(4.3%), S(2.9%), T(2.2%), | A-V(0.8%), I-T(0.6%), V-I(0.6%), A-E(0.4%), V-A(0.4%), A-X(0.3%), I- |

| | | | | | | |
|---|---|---|---|---|---|---|
| | | | | | L(2%), X(0.8%), G(0.3%), R(0.2%) | V(0.3%), A-T(0.2%), E-X(0.1%), I-A(0.1%), I-L(0.1%), I-R(0.1%), S-I(0.1%), S-X(0.1%), V-G(0.1%), V-L(0.1%), V-T(0.1%) |
| | 220 | H(73%), R(27%) | H(67.6%), R(28.2%), L(2%), X(1.2%), P(0.8%), Y(0.2%) | human-human | H(76.3%), R(23.5%), P(0.2%) | R-H(0.5%), H-P(0.2%), H-R(0.2%) |
| | | | | human-swine | H(86.4%), R(6.8%), L(3.4%), P(2.3%), X(1.1%) | H-L(3.4%), H-R(3.4%), R-H(2.3%), H-X(1.1%) |
| | | | | swine-human | H(64.3%), R(35.7%) | R-H(3.6%) |
| | | | | swine-swine | H(61.2%), R(36.6%), L(1%), X(0.8%), P(0.4%), Y(0.1%) | R-H(3%), R-L(0.8%), H-R(0.4%), H-X(0.4%), H-P(0.1%), H-Y(0.1%), R-P(0.1%), R-X(0.1%) |
| | 221 | R(80.5%), Q(19.5%) | R(75.6%), Q(19%), L(3.5%), X(1.2%), P(0.6%) | human-human | R(82.2%), Q(17.8%) | R-Q(1.1%), Q-R(0.5%) |
| | | | | human-swine | R(80.7%), Q(18.2%), X(1.1%) | R-Q(2.3%), R-X(1.1%) |
| | | | | swine-human | R(82.1%), Q(17.9%) | NA |
| | | | | swine-swine | R(80.5%), Q(15.4%), L(3%), X(0.8%), P(0.4%) | R-Q(3.7%), R-L(1.1%), R-X(0.3%), Q-X(0.2%), R-P(0.2%), L-P(0.1%), L-R(0.1%), Q-P(0.1%) |
| | 231 | P(100%) | P(93.5%), H(3.9%), X(1.2%), L(0.9%), S(0.5%) | human-human | P(100%) | NA |
| | | | | human-swine | P(96.6%), H(2.3%), X(1.1%) | P-H(2.3%), P-X(1.1%) |
| | | | | swine-human | P(100%) | NA |
| | | | | swine-swine | P(95.2%), H(3.1%), X(0.8%), L(0.6%), S(0.3%) | P-H(0.7%), P-L(0.4%), P-X(0.4%), P-S(0.3%) |
| | 247 | X(56.3%), L(40.9%), P(2.3%), H(0.5%) | X(53.7%), L(40.3%), P(4.3%), H(1.5%), F(0.2%) | human-human | X(63%), L(34.5%), P(2.5%) | X-L(0.9%), L-X(0.2%) |
| | | | | human-swine | X(83%), L(8%), P(8%), F(1.1%) | X-L(2.3%), X-P(2.3%), L-F(1.1%) |
| | | | | swine-human | X(50%), L(42.9%), H(3.6%), P(3.6%) | NA |
| | | | | swine-swine | L(48.3%), X(46.8%), P(3.4%), H(1.5%) | X-L(1.2%), L-P(0.6%), L-X(0.4%), L-H(0.4%), X-P(0.4%), P-X(0.2%) |
| | 250 | X(56.3%), P(37.7%), Q(3.7%), L(2.3%) | X(54%), P(38.4%), Q(5.1%), L(2.5%) | human-human | X(63%), P(32.1%), L(2.5%), Q(2.5%) | X-Q(0.7%), P-Q(0.2%), P-X(0.2%), X-P(0.2%) |
| | | | | human-swine | X(84.1%), P(6.8%), L(5.7%), Q(3.4%) | X-Q(3.4%) |
| | | | | swine-human | X(50%), P(46.4%), L(3.6%) | NA |
| | | | | swine-swine | X(46.9%), P(46.7%), Q(4%), L(2.4%) | X-Q(1.4%), P-Q(0.4%), P-L(0.4%), P-X(0.4%), L-X(0.2%), Q-P(0.2%), Q-X(0.1%), X-P(0.1%) |
| **H1** | 7 | V(87.8%), I(8.5%), T(2.7%), A(0.5%), F(0.5%) | V(80.2%), I(17.4%), A(1%), F(0.5%), T(0.5%), L(0.4%) | human-human | V(93.8%), I(4.4%), T(1.8%) | V-I(1.3%) |
| | | | | human-swine | V(94.8%), I(3.4%), A(0.9%), L(0.9%) | V-I(1.7%), V-A(0.9%), V-L(0.9%) |
| | | | | swine-human | V(79.5%), I(15.4%), A(2.6%), F(2.6%) | V-A(2.6%) |
| | | | | swine-swine | V(80.1%), I(17.9%), A(0.9%), F(0.5%), T(0.4%), L(0.2%) | V-I(1.2%), V-A(0.2%), I-T(0.1%), I-V(0.1%), V-F(0.1%), V-L(0.1%), V-T(0.1%) |
| | 113 | I(46.8%), A(40.4%), T(5.9%), E(4.8%), N(1.1%), S(1.1%) | A(41.4%), I(29.4%), T(15.5%), S(9.1%), E(1.9%), V(1%), D(0.9%), N(0.6%), F(0.2%), | human-human | I(57.6%), A(34.9%), E(4.7%), T(2.1%), N(0.5%), S(0.3%) | A-S(0.3%), I-A(0.3%) |
| | | | | human-swine | I(74.1%), A(14.7%), T(6%), E(4.3%), V(0.9%) | I-T(2.6%), A-T(1.7%), I-V(0.9%) |
| | | | | swine-human | I(35.9%), A(33.3%), T(20.5%), E(5.1%), N(2.6%), S(2.6%) | T-E(2.6%), T-N(2.6%) |
| | | | | swine-swine | A(47.5%), I(24.6%), T(14.6%), S(9.7%), E(1.4%), V(0.8%), | A-T(2.2%), A-S(0.4%), I-T(0.4%), A-D(0.3%), A-V(0.2%), T-A(0.2%), I- |

| | | | | | |
|---|---|---|---|---|---|
| | | | | D(0.7%), N(0.3%), F(0.1%), H(0.1%), X(0.1%) | N(0.1%), I-V(0.1%), T-S(0.1%), X-A(0.1%), A-E(0.1%), A-H(0.1%), A-N(0.1%), A-X(0.1%), I-A(0.1%), I-F(0.1%), I-S(0.1%), S-F(0.1%), T-I(0.1%), T-N(0.1%) |
| 137 | T(38.3%), E(36.7%), A(17.6%), K(4.8%), S(1.1%), D(0.5%), V(0.5%) | A(41.6%), E(27.3%), T(23%), S(2.1%), K(1.9%), V(1.1%), G(0.8%), R(0.8%), X(0.7%), | human-human | T(47.8%), E(32.6%), A(15%), K(4.7%) | A-T(0.3%), T-A(0.3%), T-E(0.3%) |
| | | | human-swine | T(62.9%), A(15.5%), E(14.7%), K(3.4%), V(1.7%), D(0.9%), S(0.9%) | T-A(3.4%), A-V(1.7%), A-D(0.9%), A-S(0.9%), K-E(0.9%) |
| | | | swine-human | E(28.2%), T(28.2%), A(25.6%), K(5.1%), S(5.1%), D(2.6%), V(2.6%), X(2.6%) | A-E(2.6%), A-T(2.6%), E-K(2.6%) |
| | | | swine-swine | A(48%), E(27.8%), T(17.4%), S(1.9%), K(1.5%), R(0.9%), X(0.7%), G(0.7%), V(0.7%), I(0.2%), D(0.2%), L(0.1%) | A-T(1.7%), A-E(1.2%), T-A(0.6%), A-S(0.4%), A-V(0.3%), E-G(0.2%), T-I(0.2%), E-K(0.2%), K-E(0.2%), E-R(0.1%), G-E(0.1%), X-E(0.1%), A-D(0.1%), A-K(0.1%), A-L(0.1%), E-D(0.1%), E-X(0.1%), R-G(0.1%), T-K(0.1%), T-V(0.1%), T-X(0.1%) |
| 145 | T(35.6%), V(27.7%), S(25.5%), A(5.3%), X(2.7%), I(1.6%), L(1.1%) | T(59.7%), V(16.2%), S(11.4%), A(4.7%), I(3.2%), X(2.4%), P(1%), L(0.9%), K(0.2%), | human-human | S(40.1%), T(26.6%), V(25.8%), A(4.1%), I(1.8%), X(0.8%), L(0.5%), P(0.3%) | A-T(0.3%), I-T(0.3%), S-P(0.3%), T-I(0.3%), V-A(0.3%) |
| | | | human-swine | S(58.6%), T(19%), V(15.5%), P(3.4%), A(1.7%), I(0.9%), W(0.9%) | S-P(2.6%), S-V(0.9%), S-W(0.9%), T-A(0.9%), T-P(0.9%), V-I(0.9%), X-T(0.9%) |
| | | | swine-human | T(56.4%), V(23.1%), X(10.3%), S(5.1%), A(2.6%), L(2.6%) | S-A(2.6%) |
| | | | swine-swine | T(65.8%), V(16.9%), S(5.8%), A(4.3%), I(3%), X(2.5%), L(0.9%), P(0.5%), K(0.2%), M(0.1%) | T-A(0.6%), V-I(0.5%), V-A(0.3%), A-T(0.2%), T-X(0.2%), S-P(0.2%), T-I(0.1%), A-V(0.1%), A-X(0.1%), I-M(0.1%), I-T(0.1%), I-X(0.1%), S-T(0.1%), T-K(0.1%), T-P(0.1%), T-S(0.1%), T-V(0.1%), V-L(0.1%), V-X(0.1%), X-A(0.1%), X-V(0.1%) |
| 157 | A(48.9%), K(23.4%), T(8%), E(7.4%), N(6.4%), V(2.1%), S(1.6%), | A(52.6%), K(13.6%), T(11.7%), V(6.1%), E(6%), N(4.5%), S(2.1%), I(0.9%), R(0.9%), | human-human | A(58.1%), K(24.5%), N(5.4%), E(4.9%), T(3.9%), V(1.6%), S(0.8%), I(0.5%), R(0.3%) | K-E(1.8%), A-K(0.3%), A-T(0.3%), K-N(0.3%), K-R(0.3%), N-S(0.3%) |
| | | | human-swine | A(66.4%), E(8.6%), K(8.6%), T(6%), N(3.4%), V(3.4%), G(0.9%), I(0.9%), R(0.9%), S(0.9%) | A-T(4.3%), A-E(3.4%), A-V(3.4%), N-K(1.7%), A-G(0.9%), A-I(0.9%), A-K(0.9%), A-S(0.9%), K-E(0.9%), K-R(0.9%) |
| | | | swine-human | A(43.6%), T(20.5%), N(10.3%), E(7.7%), K(7.7%), I(2.6%), Q(2.6%), S(2.6%), V(2.6%) | A-S(2.6%), K-E(2.6%) |
| | | | swine-swine | A(56.4%), K(14.7%), T(10.9%), E(5.1%), V(4.9%), N(3.9%), S(1.3%), I(0.9%), R(0.7%), Q(0.6%), D(0.3%), G(0.1%), M(0.1%), X(0.1%) | A-T(2.2%), A-V(1.7%), K-E(1.1%), A-S(0.9%), E-K(0.8%), K-N(0.4%), A-E(0.3%), T-A(0.3%), A-N(0.2%), K-R(0.2%), A-D(0.2%), A-K(0.1%), K-G(0.1%), N-K(0.1%), T-K(0.1%), V-I(0.1%), X- |

| | | | | | |
|---|---|---|---|---|---|
| | | | | | E(0.1%), A-X(0.1%), D-N(0.1%), E-A(0.1%), E-N(0.1%), E-Q(0.1%), I-V(0.1%), K-Q(0.1%), K-X(0.1%), M-K(0.1%), N-D(0.1%), T-E(0.1%), T-I(0.1%), T-S(0.1%), V-M(0.1%) |
| 209 | Q(60.1%), H(34%), K(2.7%), R(2.7%), S(0.5%) | Q(72%), H(15.4%), R(8.4%), K(2.1%), N(0.8%), E(0.6%), S(0.2%), D(0.1%), G(0.1%), L | human-human | Q(65.4%), H(31.3%), R(2.3%), K(0.8%), S(0.3%) | H-S(0.3%), Q-H(0.3%), Q-K(0.3%), Q-R(0.3%), R-H(0.3%) |
| | | | human-swine | Q(81%), H(16.4%), R(1.7%), K(0.9%) | H-R(0.9%), Q-K(0.9%), Q-R(0.9%) |
| | | | swine-human | Q(66.7%), H(25.6%), K(7.7%) | Q-K(2.6%) |
| | | | swine-swine | Q(72%), H(15.8%), R(8.8%), K(2.1%), N(0.5%), E(0.4%), S(0.1%), D(0.1%), G(0.1%), L(0.1%), P(0.1%), T(0.1%), X(0.1%), Y(0.1%) | H-R(0.6%), H-N(0.4%), Q-H(0.4%), Q-K(0.4%), H-Q(0.3%), R-H(0.2%), Q-E(0.1%), Q-R(0.1%), X-H(0.1%), H-D(0.1%), H-P(0.1%), H-X(0.1%), H-Y(0.1%), K-E(0.1%), K-R(0.1%), K-T(0.1%), N-K(0.1%), Q-G(0.1%), Q-L(0.1%) |
| 213 | A(84%), T(14.4%), S(1.6%) | A(53.6%), T(45.1%), S(1%), P(0.1%), V(0.1%), X(0.1%) | human-human | A(90.4%), T(9%), S(0.5%) | A-T(0.3%) |
| | | | human-swine | A(92.2%), T(7.8%) | A-T(2.6%) |
| | | | swine-human | A(69.2%), T(25.6%), S(5.1%) | NA |
| | | | swine-swine | A(49.3%), T(49.3%), S(1.2%), X(0.1%), P(0.1%), V(0.1%) | A-T(1.1%), T-A(0.2%), A-S(0.1%), A-X(0.1%), X-A(0.1%), A-V(0.1%), T-P(0.1%) |
| 318 | E(67%), K(29.8%), N(1.6%), Q(1.6%) | E(82.6%), K(15.9%), Q(0.8%), N(0.3%), G(0.1%), R(0.1%), V(0.1%) | human-human | E(54.5%), K(43.9%), N(1%), Q(0.5%) | E-K(0.5%) |
| | | | human-swine | K(62.9%), E(35.3%), N(1.7%) | K-E(1.7%), K-N(1.7%) |
| | | | swine-human | E(87.2%), K(7.7%), Q(5.1%) | K-Q(2.6%) |
| | | | swine-swine | E(90%), K(8.8%), Q(0.9%), N(0.1%), G(0.1%), R(0.1%), V(0.1%), X(0.1%) | E-K(1.4%), K-E(0.1%), X-E(0.1%), E-G(0.1%), E-V(0.1%), E-X(0.1%), K-R(0.1%) |
| 337 | I(62.8%), V(26.1%), T(10.1%), E(1.1%) | I(69.6%), V(18.9%), T(9.8%), N(0.8%), D(0.6%), E(0.1%), K(0.1%), X(0.1%) | human-human | I(52.2%), V(40.1%), T(7.2%), E(0.5%) | I-T(0.3%), V-T(0.3%) |
| | | | human-swine | V(54.3%), I(35.3%), T(10.3%) | V-I(6%), I-T(1.7%), V-T(0.9%) |
| | | | swine-human | I(71.8%), T(20.5%), V(5.1%), E(2.6%) | D-E(2.6%), I-T(2.6%) |
| | | | swine-swine | I(75.5%), V(14.7%), T(8.3%), D(0.7%), N(0.6%), X(0.1%), E(0.1%), K(0.1%) | I-T(1.4%), I-V(1.2%), V-I(1%), D-N(0.1%), I-X(0.1%), X-I(0.1%), D-E(0.1%), I-K(0.1%), I-N(0.1%), T-I(0.1%) |
| 361 | V(60.6%), I(38.8%), M(0.5%) | I(69.8%), V(29.7%), L(0.4%), M(0.1%) | human-human | V(70.3%), I(29.2%), M(0.5%) | I-M(0.3%), I-V(0.3%), M-V(0.3%), V-I(0.3%) |
| | | | human-swine | V(75%), I(25%) | V-I(6.9%) |
| | | | swine-human | I(64.1%), V(35.9%) | I-V(5.1%) |
| | | | swine-swine | I(77.3%), V(22.4%), L(0.2%), M(0.1%), X(0.1%) | I-V(1.8%), V-I(0.4%), I-L(0.2%), X-V(0.1%), I-M(0.1%), V-L(0.1%), V-X(0.1%) |
| 372 | E(96.8%), D(2.1%), G(1.1%) | E(93.8%), G(3.8%), D(2.2%), A(0.1%), K(0.1%) | human-human | E(99%), D(1%) | E-D(0.3%), G-E(0.3%) |
| | | | human-swine | E(96.6%), G(1.7%), D(0.9%), K(0.9%) | E-G(1.7%), D-E(0.9%), E-D(0.9%), E-K(0.9%) |
| | | | swine-human | E(87.2%), G(7.7%), D(5.1%) | E-G(2.6%), G-E(2.6%) |
| | | | swine-swine | E(93.8%), G(4.2%), D(1.9%), A(0.1%), X(0.1%) | E-D(0.6%), E-G(0.6%), G-E(0.4%), X-E(0.1%), D-E(0.1%), E-A(0.1%), E-X(0.1%) |

| | | | | | | |
|---|---|---|---|---|---|---|
| | 390 | G(72.9%), E(13.3%), K(11.7%), Q(1.6%), R(0.5%) | G(85.7%), E(5.5%), K(4.4%), R(3.7%), S(0.5%), M(0.1%), T(0.1%) | human-human | G(61%), E(22%), K(16%), Q(1%) | E-G(1%), E-K(0.5%), G-E(0.5%), K-Q(0.3%) |
| | | | | human-swine | G(46.6%), E(32.8%), K(18.1%), R(1.7%), T(0.9%) | E-G(4.3%), K-E(2.6%), E-K(1.7%), G-E(0.9%), G-R(0.9%), K-G(0.9%), K-R(0.9%), K-T(0.9%) |
| | | | | swine-human | G(92.3%), E(2.6%), K(2.6%), R(2.6%) | NA |
| | | | | swine-swine | G(91.1%), R(3.8%), K(2.5%), E(2.1%), S(0.4%), M(0.1%), X(0.1%) | G-R(0.5%), R-K(0.2%), G-E(0.1%), G-S(0.1%), R-G(0.1%), X-G(0.1%), E-K(0.1%), G-K(0.1%), G-M(0.1%), G-X(0.1%), K-E(0.1%) |
| | 415 | H(41%), K(35.1%), N(10.1%), S(8%), Q(2.7%), D(1.1%), Y(1.1%), | H(30.6%), N(25.8%), K(21.7%), D(11.6%), S(4.1%), Q(3.7%), R(1.2%), E(0.9%), G(0.3% | human-human | H(50.9%), K(32.6%), S(7.2%), N(4.9%), Q(2.8%), D(0.8%), Y(0.5%), R(0.3%) | N-K(0.5%), K-N(0.3%), K-R(0.3%) |
| | | | | human-swine | H(69.8%), K(16.4%), Q(5.2%), S(5.2%), R(1.7%), D(0.9%), N(0.9%) | H-K(0.9%), H-R(0.9%), K-R(0.9%) |
| | | | | swine-human | H(33.3%), N(28.2%), K(23.1%), S(5.1%), D(2.6%), E(2.6%), Q(2.6%), Y(2.6%) | H-Y(2.6%), N-S(2.6%) |
| | | | | swine-swine | N(27.3%), H(26.4%), K(22.5%), D(15.2%), S(3.7%), Q(3%), E(0.9%), R(0.7%), G(0.2%), I(0.1%), X(0.1%) | D-N(1.6%), K-R(0.4%), H-Q(0.3%), N-S(0.3%), N-D(0.2%), D-E(0.2%), D-G(0.2%), D-S(0.1%), K-N(0.1%), Q-H(0.1%), S-N(0.1%), X-K(0.1%), K-I(0.1%), K-X(0.1%), N-H(0.1%) |
| | 456 | S(85.1%), F(7.4%), L(7.4%) | S(60.1%), F(36.3%), L(3.2%), A(0.2%), T(0.1%), Y(0.1%) | human-human | S(90.4%), L(7%), F(2.6%) | L-F(0.3%), S-L(0.3%) |
| | | | | human-swine | S(94.8%), L(4.3%), F(0.9%) | NA |
| | | | | swine-human | S(74.4%), F(23.1%), L(2.6%) | NA |
| | | | | swine-swine | S(55.6%), F(41.4%), L(2.8%), A(0.1%), T(0.1%), X(0.1%), Y(0.1%) | F-L(0.3%), F-S(0.3%), S-F(0.2%), L-F(0.1%), S-A(0.1%), X-S(0.1%), F-Y(0.1%), S-T(0.1%), S-X(0.1%) |
| **H3** | 9 | Y(70.2%), C(10.6%), H(7.4%), N(5.3%), Q(5.3%), X(1.1%) | Y(56.1%), C(38.7%), H(2.4%), Q(1.5%), S(0.6%), G(0.3%), N(0.3%) | human-human | Y(82.5%), C(4.8%), N(4.8%), H(4.2%), Q(3.2%), X(0.5%) | C-X(0.5%), Y-N(0.5%) |
| | | | | human-swine | Y(85.7%), C(8.6%), H(5.7%) | Y-H(5.7%), Y-C(2.9%) |
| | | | | swine-human | C(42.9%), Y(28.6%), H(21.4%), Q(7.1%) | Y-H(14.3%), C-Y(7.1%) |
| | | | | swine-swine | Y(52.8%), C(42.5%), H(2.2%), Q(1.6%), S(0.5%), G(0.2%), N(0.2%) | Y-C(1.6%), C-Y(1.3%), Y-H(0.4%), C-G(0.2%), C-S(0.2%), H-Q(0.2%), Y-N(0.2%) |
| | 21 | G(97.9%), E(1.1%), R(1.1%) | G(78.4%), E(10.7%), R(8.8%), K(1.5%), S(0.6%) | human-human | G(98.4%), R(1.1%), E(0.5%) | G-E(0.5%), G-R(0.5%) |
| | | | | human-swine | G(82.9%), R(8.6%), E(5.7%), K(2.9%) | G-E(5.7%), G-R(5.7%), G-K(2.9%) |
| | | | | swine-human | G(100%) | NA |
| | | | | swine-swine | G(82.7%), E(9%), R(7.2%), K(0.7%), S(0.4%) | G-E(3.4%), G-R(2.5%), G-S(0.4%), R-K(0.4%), E-K(0.2%), E-R(0.2%), G-K(0.2%), R-G(0.2%) |
| | 25 | S(85.1%), N(14.9%) | S(66.5%), N(26.5%), T(3%), G(1.5%), R(1.2%), I(0.9%), C(0.3%) | human-human | S(91.5%), N(8.5%) | NA |
| | | | | human-swine | S(91.4%), N(5.7%), R(2.9%) | S-N(5.7%), S-R(2.9%) |
| | | | | swine-human | S(64.3%), N(35.7%) | NA |
| | | | | swine-swine | S(66.5%), N(27.2%), T(3.2%), G(1.3%), | S-N(3.6%), N-S(0.5%), S-G(0.5%), S-R(0.4%), |

| | | | | | |
|---|---|---|---|---|---|
| | | | | I(0.9%), R(0.7%), C(0.2%) | N-T(0.2%), S-C(0.2%), T-I(0.2%) |
| 47 | N(72.3%), D(26.6%), S(1.1%) | D(66.5%), N(29.6%), S(2.4%), G(0.9%), T(0.6%) | human-human | N(85.7%), D(13.8%), S(0.5%) | D-N(0.5%), N-S(0.5%) |
| | | | human-swine | N(71.4%), D(22.9%), S(5.7%) | N-D(11.4%), N-S(5.7%) |
| | | | swine-human | D(78.6%), N(21.4%) | NA |
| | | | swine-swine | D(71.7%), N(25.8%), S(1.4%), G(0.7%), T(0.4%) | N-D(1.3%), N-S(0.7%), D-G(0.4%), D-N(0.4%) |
| 69 | D(55.3%), N(37.2%), S(3.2%), G(2.1%), K(1.1%), X(1.1%) | N(60.4%), D(22.3%), S(8.2%), K(7%), G(1.5%), T(0.6%) | human-human | D(68.3%), N(29.1%), G(1.6%), S(0.5%), X(0.5%) | D-N(1.6%), D-G(0.5%), N-D(0.5%), N-X(0.5%) |
| | | | human-swine | D(60%), N(37.1%), S(2.9%) | D-N(5.7%) |
| | | | swine-human | N(71.4%), S(21.4%), K(7.1%) | N-S(7.1%) |
| | | | swine-swine | N(63.6%), D(18.2%), S(8.8%), K(7.6%), G(1.3%), T(0.5%) | D-N(1.1%), N-S(0.9%), D-G(0.5%), K-N(0.5%), N-D(0.4%), N-K(0.4%), N-T(0.2%), S-N(0.2%) |
| 108 | K(73.4%), T(14.9%), R(10.6%), N(1.1%) | T(48.5%), K(36%), R(6.7%), E(4%), N(2.7%), G(1.2%), M(0.3%), Q(0.3%), S(0.3%) | human-human | K(84.7%), R(7.9%), T(6.9%), E(0.5%) | K-E(0.5%), K-R(0.5%), K-T(0.5%) |
| | | | human-swine | K(80%), T(14.3%), E(2.9%), R(2.9%) | K-E(2.9%), K-R(2.9%), K-T(2.9%) |
| | | | swine-human | T(57.1%), K(28.6%), N(7.1%), R(7.1%) | NA |
| | | | swine-swine | T(53.3%), K(32.3%), R(6.1%), E(3.4%), N(3.1%), G(1.3%), M(0.2%), Q(0.2%), S(0.2%) | K-R(1.1%), R-K(0.5%), K-N(0.4%), T-K(0.4%), E-G(0.2%), E-Q(0.2%), K-E(0.2%), N-T(0.2%), R-M(0.2%), T-S(0.2%) |
| 117 | D(85.1%), Y(14.9%) | D(53.7%), Y(43.3%), F(1.5%), E(0.9%), H(0.6%) | human-human | D(94.7%), Y(5.3%) | NA |
| | | | human-swine | D(94.3%), Y(5.7%) | NA |
| | | | swine-human | Y(64.3%), D(35.7%) | NA |
| | | | swine-swine | D(49%), Y(49%), F(1.1%), E(0.5%), H(0.4%) | Y-F(0.7%), D-E(0.5%), D-Y(0.5%), Y-H(0.4%) |
| 140 | S(67%), G(22.3%), D(4.3%), N(4.3%), E(2.1%) | S(53.4%), D(15.9%), N(13.1%), G(11.6%), I(3%), R(1.2%), E(0.9%), V(0.6%), A(0.3%) | human-human | S(60.3%), G(29.6%), D(6.3%), E(2.1%), N(1.6%) | G-D(1.1%), D-E(0.5%), D-G(0.5%), G-S(0.5%), S-G(0.5%) |
| | | | human-swine | S(42.9%), G(40%), D(14.3%), N(2.9%) | G-D(5.7%), S-N(2.9%) |
| | | | swine-human | S(85.7%), N(14.3%) | NA |
| | | | swine-swine | S(58.9%), D(14.8%), G(10.3%), N(10.3%), I(3.2%), R(1.3%), E(0.7%), V(0.4%), A(0.2%) | S-N(3.4%), G-D(1.6%), D-N(0.9%), G-S(0.7%), S-G(0.5%), S-I(0.4%), S-R(0.4%), D-E(0.2%), G-A(0.2%), G-E(0.2%), N-D(0.2%), R-V(0.2%), S-D(0.2%), S-V(0.2%) |
| 158 | G(48.9%), R(40.4%), K(6.4%), E(4.3%) | G(59.1%), E(13.7%), K(12.2%), R(11.6%), N(2.7%), A(0.3%), Q(0.3%) | human-human | G(54%), R(42.3%), K(3.2%), E(0.5%) | R-G(1.1%), R-K(1.1%), G-R(0.5%) |
| | | | human-swine | G(71.4%), R(20%), E(5.7%), K(2.9%) | G-E(2.9%), G-R(2.9%), R-G(2.9%) |
| | | | swine-human | G(42.9%), E(28.6%), K(14.3%), R(14.3%) | E-G(7.1%) |
| | | | swine-swine | G(56.2%), E(17.5%), K(13.5%), R(10.5%), N(2%), A(0.2%), Q(0.2%) | E-G(2.2%), E-K(1.3%), G-E(0.9%), K-N(0.9%), G-K(0.5%), G-R(0.5%), R-G(0.5%), K-E(0.4%), K-G(0.4%), E-A(0.2%), E-N(0.2%), G-N(0.2%), K-Q(0.2%), K-R(0.2%), R-K(0.2%) |
| 161 | N(53.2%), S(24.5%), K(22.3%) | N(62.2%), K(26.5%), S(9.1%), I(0.9%), M(0.3%), R(0.3%), X(0.3%), Y(0.3%) | human-human | N(49.7%), K(26.5%), S(23.8%) | N-K(1.6%), N-S(1.1%), K-N(0.5%), S-N(0.5%) |
| | | | human-swine | N(40%), K(31.4%), S(20%), I(5.7%), R(2.9%) | K-N(5.7%), N-K(5.7%), N-S(5.7%), S-I(5.7%), S-N(2.9%), S-R(2.9%) |

| | | | | | | |
|---|---|---|---|---|---|---|
| | | | | swine-human | N(64.3%), S(21.4%), K(14.3%) | N-S(7.1%) |
| | | | | swine-swine | N(67.9%), K(23.8%), S(7.6%), I(0.2%), M(0.2%), X(0.2%), Y(0.2%) | N-K(5%), K-N(2%), N-S(0.5%), S-N(0.4%), K-I(0.2%), K-M(0.2%), K-S(0.2%), N-X(0.2%), N-Y(0.2%), S-K(0.2%) |
| | 187 | N(89.4%), K(7.4%), S(2.1%), D(1.1%) | N(81.7%), S(12.8%), D(3.4%), K(1.5%), G(0.3%), I(0.3%) | human-human | N(93.1%), K(6.9%) | K-N(0.5%) |
| | | | | human-swine | N(91.4%), D(5.7%), S(2.9%) | N-D(5.7%), N-S(2.9%) |
| | | | | swine-human | N(78.6%), S(14.3%), D(7.1%) | N-D(7.1%) |
| | | | | swine-swine | N(81.6%), S(14.8%), D(2%), K(1.3%), G(0.2%), I(0.2%) | N-D(1.3%), S-N(0.7%), N-S(0.4%), K-N(0.2%), N-I(0.2%), S-G(0.2%) |
| | 210 | L(100%) | L(97.3%), V(1.5%), I(1.2%) | human-human | L(100%) | NA |
| | | | | human-swine | L(100%) | NA |
| | | | | swine-human | L(100%) | NA |
| | | | | swine-swine | L(97.7%), V(1.4%), I(0.9%) | L-I(0.5%), L-V(0.2%) |
| | 245 | R(85.1%), I(14.9%) | R(50.3%), I(48.8%), T(0.6%), V(0.3%) | human-human | R(94.7%), I(5.3%) | NA |
| | | | | human-swine | R(85.7%), I(14.3%) | R-I(8.6%) |
| | | | | swine-human | I(64.3%), R(35.7%) | NA |
| | | | | swine-swine | I(54.1%), R(45.4%), T(0.4%), V(0.2%) | I-R(0.4%), I-T(0.4%), R-I(0.4%), I-V(0.2%) |
| | 246 | I(95.7%), V(3.2%), M(1.1%) | I(94.5%), V(3.4%), M(2.1%) | human-human | I(96.8%), V(2.6%), M(0.5%) | I-V(1.1%), I-M(0.5%) |
| | | | | human-swine | I(91.4%), V(5.7%), M(2.9%) | I-M(2.9%) |
| | | | | swine-human | I(100%) | NA |
| | | | | swine-swine | I(95.5%), V(3.1%), M(1.4%) | I-M(0.7%), V-I(0.4%), M-V(0.2%) |
| | 277 | R(77.7%), Q(19.1%), H(3.2%) | Q(60.7%), R(33.8%), H(3.7%), K(0.9%), E(0.3%), L(0.3%), Y(0.3%) | human-human | R(88.9%), Q(9%), H(2.1%) | R-Q(1.1%), R-H(0.5%) |
| | | | | | R(82.9%), Q(11.4%), H(5.7%) | R-H(5.7%) |
| | | | | | Q(71.4%), R(21.4%), H(7.1%) | NA |
| | | | | | Q(66.1%), R(28.3%), H(4.3%), K(0.7%), E(0.2%), L(0.2%), Y(0.2%) | R-Q(0.9%), H-Q(0.4%), Q-K(0.4%), Q-R(0.4%), H-Y(0.2%), Q-E(0.2%), Q-H(0.2%), Q-L(0.2%), R-H(0.2%) |
| **NP** | 22 | A(91.9%), T(8.1%) | A(77.7%), T(20.9%), V(1.3%) | human-human | A(95.9%), T(4.1%) | A-T(0.8%) |
| | | | | human-swine | A(97%), T(1.5%), V(1.5%) | A-T(1.5%), A-V(1.5%) |
| | | | | swine-human | A(68.4%), T(31.6%) | NA |
| | | | | swine-swine | A(75.8%), T(23.3%), V(0.9%) | A-T(0.6%), A-V(0.5%), T-A(0.1%) |
| | 33 | I(91.4%), V(8.6%) | I(65.6%), V(34.4%) | human-human | I(96.4%), V(3.6%) | I-V(1.1%), V-I(0.3%) |
| | | | | human-swine | I(97%), V(3%) | NA |
| | | | | swine-human | I(52.6%), V(47.4%) | I-V(5.3%) |
| | | | | swine-swine | I(60.1%), V(39.9%) | I-V(1.6%), V-I(0.9%) |
| | 53 | E(80.6%), D(19.4%) | E(94.4%), D(5.6%) | human-human | E(78.2%), D(21.8%) | NA |
| | | | | human-swine | E(77.3%), D(22.7%) | D-E(3%) |
| | | | | swine-human | E(94.7%), D(5.3%) | E-D(5.3%) |
| | | | | swine-swine | E(97.2%), D(2.8%) | D-E(0.1%) |
| | 61 | I(54.3%), L(40.9%), M(2.7%), V(1.6%), T(0.5%) | I(93.3%), L(5.8%), M(1%) | human-human | I(51.9%), L(44.8%), M(1.7%), V(1.4%), T(0.3%) | I-L(0.6%), I-T(0.3%), I-V(0.3%), L-I(0.3%) |
| | | | | human-swine | I(71.2%), L(28.8%) | NA |
| | | | | swine-human | I(100%) | NA |
| | | | | swine-swine | I(97.3%), L(2.1%), M(0.6%) | I-M(0.6%) |

| | | | | | |
|---|---|---|---|---|---|
| 100 | V(62.4%), I(28%), R(9.7%) | R(37.2%), V(35.3%), I(24%), L(1.5%), M(0.8%), K(0.6%), A(0.4%), T(0.2%) | human-human | V(69.3%), I(26.2%), R(4.4%) | I-V(0.6%), V-I(0.6%) |
| | | | human-swine | V(66.7%), I(30.3%), R(3%) | V-I(4.5%), I-V(1.5%) |
| | | | swine-human | R(52.6%), I(26.3%), V(21.1%) | NA |
| | | | swine-swine | R(44%), V(29.5%), I(24%), L(1.3%), M(0.5%), K(0.3%), A(0.2%), T(0.1%) | V-I(1%), I-L(0.6%), I-V(0.6%), I-M(0.5%), R-K(0.3%), V-A(0.2%), I-R(0.1%), I-T(0.1%) |
| 190 | V(50.5%), A(45.7%), T(3.2%), I(0.5%) | A(48.8%), V(45.1%), T(4.8%), I(1%), N(0.2%), S(0.2%) | human-human | A(49.2%), V(48.9%), T(1.9%) | A-T(0.3%), A-V(0.3%) |
| | | | human-swine | A(63.6%), V(33.3%), N(1.5%), T(1.5%) | A-N(1.5%), A-T(1.5%), A-V(1.5%) |
| | | | swine-human | A(47.4%), V(47.4%), I(5.3%) | V-I(5.3%) |
| | | | swine-swine | V(48.6%), A(46.9%), T(3.8%), I(0.6%), S(0.1%) | A-T(1.6%), V-A(1%), A-V(0.7%), T-I(0.2%), V-I(0.2%), A-I(0.1%), A-S(0.1%) |
| 217 | I(41.4%), V(29%), S(21.5%), G(7%), A(1.1%) | I(63%), V(30.7%), S(4%), A(1%), T(0.8%), L(0.4%), G(0.2%) | human-human | I(42%), V(26.8%), S(23.5%), G(6.9%), A(0.8%) | I-V(0.6%), S-G(0.6%), G-S(0.3%), I-S(0.3%), V-A(0.3%), V-I(0.3%) |
| | | | human-swine | I(47%), V(30.3%), S(21.2%), G(1.5%) | I-V(6.1%) |
| | | | swine-human | I(63.2%), V(36.8%) | NA |
| | | | swine-swine | I(66.2%), V(31.2%), S(1.4%), A(0.6%), T(0.5%), L(0.2%) | V-I(2.1%), I-V(1.3%), T-A(0.2%), I-L(0.1%), V-A(0.1%), V-L(0.1%) |
| 343 | V(79.6%), L(20.4%) | V(89.8%), I(6.1%), L(3.8%), P(0.2%) | human-human | V(77.1%), L(22.7%), I(0.3%) | V-L(0.6%), L-V(0.3%), V-I(0.3%) |
| | | | human-swine | V(74.2%), L(19.7%), I(6.1%) | V-I(3%) |
| | | | swine-human | V(100%) | NA |
| | | | swine-swine | V(93.7%), I(4.8%), L(1.4%), P(0.1%) | V-I(1.5%), V-P(0.1%) |
| 357 | K(90.9%), Q(5.4%), R(3.8%) | K(67.4%), Q(31.3%), R(1.3%) | human-human | K(93.4%), Q(3.3%), R(3.3%) | K-R(0.8%) |
| | | | human-swine | K(93.9%), Q(3%), R(3%) | K-R(1.5%) |
| | | | swine-human | K(78.9%), Q(21.1%) | NA |
| | | | swine-swine | K(61.5%), Q(37.8%), R(0.7%) | Q-K(0.9%), K-R(0.2%), Q-R(0.2%) |
| 377 | S(52.2%), N(37.6%), I(8.6%), V(1.1%), R(0.5%) | N(46.3%), I(29.9%), S(16.9%), V(5.6%), M(1.3%) | human-human | S(56.9%), N(38.4%), I(3.9%), V(0.6%), R(0.3%) | S-N(0.3%), S-R(0.3%) |
| | | | human-swine | N(53%), S(43.9%), I(3%) | N-S(1.5%) |
| | | | swine-human | I(47.4%), N(36.8%), S(10.5%), V(5.3%) | NA |
| | | | swine-swine | N(43.3%), I(36%), S(13.4%), V(5.8%), M(1.4%) | I-V(0.9%), S-N(0.7%), N-S(0.5%), I-M(0.2%), N-I(0.1%), S-I(0.1%) |
| 384 | R(75.3%), G(13.4%), K(11.3%) | R(60.1%), K(38%), G(1.9%) | human-human | R(79.6%), G(14.4%), K(6.1%) | R-G(0.6%), R-K(0.3%) |
| | | | human-swine | R(86.4%), G(10.6%), K(3%) | G-R(3%) |
| | | | swine-human | K(52.6%), R(47.4%) | NA |
| | | | swine-swine | R(55.3%), K(44.1%), G(0.6%) | R-K(0.5%), R-G(0.1%) |
| 426 | M(91.9%), L(7.5%), I(0.5%) | M(91.7%), L(7.5%), I(0.8%) | human-human | M(89.8%), L(10.2%) | M-L(0.6%) |
| | | | human-swine | M(80.3%), L(19.7%) | NA |
| | | | swine-human | M(94.7%), I(5.3%) | M-I(5.3%) |
| | | | swine-swine | M(95%), L(4.5%), I(0.5%) | M-L(0.9%), M-I(0.5%) |
| 446 | R(89.2%), K(10.2%), X(0.5%) | R(88.5%), K(11.5%) | human-human | R(91.2%), K(8.6%), X(0.3%) | R-K(0.6%), K-R(0.3%), K-X(0.3%) |

| | | | | | |
|---|---|---|---|---|---|
| | | | human-swine | R(93.9%), K(6.1%) | R-K(1.5%) |
| | | | swine-human | R(78.9%), K(21.1%) | R-K(5.3%) |
| | | | swine-swine | R(88.9%), K(11.1%) | R-K(1.5%), K-R(0.1%) |
| N1 | 45 | Q(51.7%), H(38.5%), Y(5.6%), K(3.5%), N(0.7%) | Q(69.7%), H(23.1%), Y(3.5%), K(1.8%), N(0.7%), R(0.7%), L(0.3%), F(0.2%), P(0.2%) | human-human | Q(54.2%), H(37%), Y(4.6%), K(3.8%), N(0.4%) | H-N(0.4%), Q-H(0.4%), Q-K(0.4%) |
| | | | | human-swine | Q(53.3%), H(38.3%), K(3.3%), Y(3.3%), N(1.7%) | H-N(1.7%), H-Y(1.7%), Q-H(1.7%) |
| | | | | swine-human | Q(47.4%), H(44.7%), Y(5.3%), K(2.6%) | H-Y(5.3%), Q-K(2.6%) |
| | | | | swine-swine | Q(71.7%), H(22.7%), Y(3.1%), K(1.2%), N(0.5%), R(0.4%), L(0.2%), F(0.1%), P(0.1%) | Q-H(0.6%), Q-K(0.5%), H-Y(0.3%), Q-R(0.3%), H-Q(0.2%), Q-L(0.2%), H-N(0.1%), H-P(0.1%), H-R(0.1%), Q-Y(0.1%), Y-F(0.1%) |
| | 72 | T(82.5%), I(11.2%), N(5.6%), X(0.7%) | T(49.4%), N(34.2%), I(12.8%), S(1.7%), A(1.2%), V(0.3%), Y(0.3%), K(0.2%) | human-human | T(86.6%), I(11.3%), N(1.7%), X(0.4%) | T-I(0.8%), T-X(0.4%) |
| | | | | human-swine | T(81.7%), I(16.7%), A(1.7%) | T-I(3.3%), T-A(1.7%) |
| | | | | swine-human | T(73.7%), I(13.2%), N(13.2%) | NA |
| | | | | swine-swine | T(47.5%), N(38.3%), I(10.5%), S(1.8%), A(1.1%), V(0.3%), Y(0.3%), K(0.1%) | T-I(1.7%), N-S(0.3%), S-N(0.3%), T-N(0.3%), A-T(0.1%), N-I(0.1%), N-K(0.1%), N-Y(0.1%), T-V(0.1%) |
| | 77 | G(58.7%), E(22.4%), R(8.4%), I(7%), V(2.8%), K(0.7%) | E(54.9%), G(25.4%), I(6.1%), V(5.8%), K(3.5%), R(3.3%), T(0.3%), A(0.2%), D(0.2%), | human-human | G(65.5%), R(13%), E(12.2%), I(8.4%), K(0.4%), V(0.4%) | G-E(0.8%), G-R(0.4%), R-K(0.4%) |
| | | | | human-swine | G(61.7%), I(15%), R(13.3%), E(6.7%), K(1.7%), V(1.7%) | E-K(1.7%), G-E(1.7%), G-R(1.7%) |
| | | | | swine-human | E(42.1%), G(39.5%), V(10.5%), I(7.9%) | NA |
| | | | | swine-swine | E(62.1%), G(21.7%), V(6.8%), I(5%), K(2.3%), R(1.5%), T(0.2%), A(0.1%), D(0.1%), M(0.1%), Q(0.1%) | E-K(1.3%), E-G(1.1%), G-E(1.1%), G-R(0.5%), G-K(0.3%), E-R(0.2%), E-D(0.1%), E-T(0.1%), G-V(0.1%), I-T(0.1%), K-Q(0.1%), V-A(0.1%), V-I(0.1%), V-M(0.1%) |
| | 79 | D(33.6%), S(33.6%), A(12.6%), G(9.8%), V(4.2%), E(2.1%), P(2.1%) | A(27.2%), T(23.4%), S(16.3%), D(15.9%), G(8.1%), V(3.6%), E(2.8%), I(0.5%), N(0.5% | human-human | S(46.6%), D(29.4%), G(11.3%), A(7.1%), V(2.5%), P(1.3%), E(0.8%), T(0.8%) | S-P(0.4%) |
| | | | | human-swine | S(51.7%), D(30%), G(11.7%), A(6.7%) | NA |
| | | | | swine-human | D(44.7%), A(23.7%), V(7.9%), E(5.3%), G(5.3%), S(5.3%), T(5.3%), P(2.6%) | S-P(2.6%) |
| | | | | swine-swine | A(32%), T(25.2%), D(15.8%), S(10.5%), G(8.2%), V(3.8%), E(2.6%), N(0.4%), P(0.4%), I(0.3%), R(0.3%), K(0.2%), L(0.1%), M(0.1%), Y(0.1%) | A-T(2%), T-A(0.5%), A-E(0.4%), A-V(0.4%), G-S(0.4%), D-G(0.3%), G-D(0.3%), A-S(0.2%), D-E(0.2%), S-P(0.2%), T-I(0.2%), A-K(0.1%), D-A(0.1%), D-N(0.1%), D-Y(0.1%), G-R(0.1%), S-L(0.1%), S-M(0.1%), T-K(0.1%), T-N(0.1%), T-S(0.1%), V-I(0.1%) |
| | 210 | G(91.6%), D(5.6%), S(2.1%), N(0.7%) | G(62%), D(31.5%), N(4%), S(2%), Y(0.3%), E(0.2%) | human-human | G(97.5%), D(1.7%), S(0.8%) | NA |
| | | | | human-swine | G(100%) | NA |

| | | | | | |
|---|---|---|---|---|---|
| | | | swine-human | G(78.9%), D(13.2%), S(5.3%), N(2.6%) | N-D(2.6%) |
| | | | swine-swine | G(57.1%), D(37.3%), N(3.3%), S(2.1%), Y(0.2%), E(0.1%) | D-G(1.1%), D-N(1.1%), G-D(0.4%), D-Y(0.2%), G-S(0.2%), D-E(0.1%), D-S(0.1%), S-G(0.1%), S-N(0.1%) |
| 232 | V(52.4%), A(44.8%), T(2.8%) | A(55.9%), V(40%), T(3.6%), I(0.3%), D(0.2%) | human-human | A(51.3%), V(46.2%), T(2.5%) | NA |
| | | | human-swine | A(50%), V(46.7%), I(1.7%), T(1.7%) | A-V(5%), A-T(1.7%), V-I(1.7%) |
| | | | swine-human | V(63.2%), A(34.2%), T(2.6%) | A-V(5.3%), A-T(2.6%), V-A(2.6%) |
| | | | swine-swine | A(60.1%), V(37%), T(2.8%), D(0.1%), I(0.1%) | A-V(3.2%), A-T(1.1%), V-A(0.6%), A-I(0.1%), V-D(0.1%), V-T(0.1%) |
| 257 | K(65%), R(35%) | K(85.7%), R(14.1%), Q(0.2%) | human-human | K(52.9%), R(47.1%) | NA |
| | | | human-swine | K(61.7%), R(38.3%) | R-K(13.3%) |
| | | | swine-human | K(92.1%), R(7.9%) | NA |
| | | | swine-swine | K(89.7%), R(10.2%), Q(0.1%) | R-K(0.6%), K-Q(0.1%), K-R(0.1%) |
| 262 | K(97.9%), R(2.1%) | K(97%), R(2.8%), M(0.2%) | human-human | K(99.6%), R(0.4%) | NA |
| | | | human-swine | K(96.7%), R(3.3%) | K-R(1.7%) |
| | | | swine-human | K(92.1%), R(7.9%) | K-R(2.6%), R-K(2.6%) |
| | | | swine-swine | K(97.5%), R(2.4%), M(0.1%) | K-R(0.7%), K-M(0.1%) |
| 331 | G(45.5%), K(42.7%), R(11.2%), N(0.7%) | G(35.2%), R(34.8%), K(25.4%), N(3.2%), E(0.7%), M(0.5%), Q(0.2%), V(0.2%) | human-human | K(51.3%), G(40.8%), R(8%) | NA |
| | | | human-swine | G(41.7%), K(41.7%), R(13.3%), E(1.7%), N(1.7%) | K-R(8.3%), K-E(1.7%), K-N(1.7%) |
| | | | swine-human | G(57.9%), K(21.1%), R(18.4%), N(2.6%) | K-R(2.6%), R-G(2.6%) |
| | | | swine-swine | R(40.2%), G(32.8%), K(23.5%), N(2.7%), E(0.4%), M(0.3%), Q(0.1%), V(0.1%) | R-G(2.5%), K-G(0.8%), K-N(0.8%), K-R(0.7%), R-K(0.6%), K-E(0.3%), K-M(0.3%), E-G(0.1%), G-K(0.1%), G-R(0.1%), K-Q(0.1%), N-K(0.1%), R-V(0.1%) |
| 365 | I(55.2%), N(23.1%), T(21%), S(0.7%) | T(54.7%), I(41.6%), N(3%), A(0.5%), P(0.2%) | human-human | I(66.8%), N(19.3%), T(13.9%) | I-T(0.4%), N-T(0.4%) |
| | | | human-swine | I(80%), N(11.7%), T(8.3%) | I-T(1.7%), T-I(1.7%) |
| | | | swine-human | T(36.8%), I(34.2%), N(26.3%), S(2.6%) | T-S(2.6%) |
| | | | swine-swine | T(61.3%), I(35.3%), N(3%), A(0.3%), P(0.1%) | T-I(2.1%), I-T(1.2%), T-A(0.3%), N-T(0.1%), T-P(0.1%) |
| 389 | V(39.2%), I(37.8%), M(18.9%), K(2.8%), T(1.4%) | V(49.6%), I(22.7%), M(17.1%), A(3.5%), L(2.8%), K(1.8%), T(1.7%), E(0.2%), G(0.2%) | human-human | I(48.7%), V(30.3%), M(18.5%), K(2.5%) | I-K(1.3%), M-I(0.4%) |
| | | | human-swine | I(48.3%), M(21.7%), V(21.7%), L(3.3%), K(1.7%), Q(1.7%), T(1.7%) | M-L(3.3%), I-T(1.7%), I-V(1.7%), M-Q(1.7%), M-V(1.7%), V-I(1.7%) |
| | | | swine-human | V(52.6%), M(31.6%), I(10.5%), T(5.3%) | A-T(2.6%), I-M(2.6%), V-I(2.6%), V-T(2.6%) |
| | | | swine-swine | V(58.6%), M(17.6%), I(15.8%), A(3.1%), L(2%), K(1.2%), T(1.2%), E(0.1%), G(0.1%), S(0.1%), X(0.1%) | V-I(2.9%), V-A(1.1%), V-L(0.9%), V-M(0.6%), M-I(0.5%), M-K(0.5%), I-M(0.3%), M-T(0.2%), A-T(0.1%), I-K(0.1%), I-T(0.1%), I-V(0.1%), L-I(0.1%), L-S(0.1%), M-L(0.1%), M-V(0.1%), M-X(0.1%), V-E(0.1%), V-G(0.1%) |

| | | | | | | |
|---|---|---|---|---|---|---|
| | 416 | D(67.8%), N(30.1%), G(1.4%), K(0.7%) | D(55.4%), N(37.3%), S(4.8%), G(1.3%), K(1%), T(0.2%) | human-human | D(71%), N(28.6%), G(0.4%) | D-N(0.8%), D-G(0.4%) |
| | | | | human-swine | D(61.7%), N(36.7%), S(1.7%) | D-N(3.3%), D-S(1.7%), N-D(1.7%) |
| | | | | swine-human | D(57.9%), N(36.8%), G(2.6%), K(2.6%) | D-N(5.3%) |
| | | | | swine-swine | D(56.5%), N(36.6%), S(4.3%), G(1.5%), K(1%), T(0.1%) | D-N(1.8%), D-S(0.9%), N-D(0.9%), N-S(0.3%), S-N(0.3%), D-G(0.2%), G-D(0.1%), N-K(0.1%), N-T(0.1%), S-K(0.1%) |
| | 418 | I(69.9%), M(29.4%), V(0.7%) | I(63.2%), M(36%), T(0.5%), V(0.3%) | human-human | I(82.8%), M(16.8%), V(0.4%) | I-V(0.4%) |
| | | | | human-swine | I(93.3%), M(6.7%) | M-I(1.7%) |
| | | | | swine-human | M(60.5%), I(39.5%) | NA |
| | | | | swine-swine | I(58%), M(41.4%), T(0.4%), V(0.2%) | M-I(1.1%), I-M(0.2%), M-V(0.2%), I-T(0.1%), M-T(0.1%) |
| **N2** | 18 | A(52%), S(46.6%), T(1.4%) | A(88.1%), S(6.5%), T(4.9%), G(0.5%) | human-human | A(56.8%), S(43.2%) | A-S(0.3%) |
| | | | | human-swine | A(84.2%), S(15.8%) | NA |
| | | | | swine-human | A(81.8%), S(9.1%), T(9.1%) | NA |
| | | | | swine-swine | A(89.7%), S(5.2%), T(4.8%), G(0.4%) | A-T(0.9%), A-G(0.2%), A-S(0.1%), T-A(0.1%) |
| | 26 | I(81.1%), V(12.2%), T(6.1%), S(0.7%) | I(64.8%), V(25.2%), M(5.7%), T(3.7%), A(0.4%), L(0.3%) | human-human | I(83.8%), V(10.1%), T(5.7%), S(0.3%) | I-S(0.3%), I-T(0.3%) |
| | | | | human-swine | I(78.9%), V(17.5%), T(3.5%) | V-I(1.8%) |
| | | | | swine-human | I(68.2%), V(31.8%) | V-I(4.5%) |
| | | | | swine-swine | I(64.2%), V(26.2%), M(6.5%), T(2.7%), A(0.2%), L(0.2%) | I-V(1.3%), V-I(1.2%), I-T(0.6%), I-M(0.2%), V-A(0.2%), I-L(0.2%), M-I(0.1%), M-T(0.1%), M-V(0.1%) |
| | 52 | L(76.4%), P(16.9%), F(4.1%), Q(2%), X(0.7%) | L(48.6%), P(29.7%), T(7.5%), F(5.3%), Q(3.3%), S(1.9%), R(1.6%), M(0.7%), I(0.5%), | human-human | L(76.7%), P(18.9%), F(4.1%), X(0.3%) | L-P(1%), L-F(0.3%), L-X(0.3%), P-L(0.3%) |
| | | | | human-swine | L(61.4%), P(28.1%), F(7%), I(1.8%), S(1.8%) | L-P(3.5%), F-S(1.8%), L-I(1.8%) |
| | | | | swine-human | L(68.2%), P(13.6%), Q(13.6%), F(4.5%) | L-Q(4.5%), S-L(4.5%) |
| | | | | swine-swine | L(48.1%), P(31%), T(8.2%), F(4.9%), Q(3%), R(1.7%), S(1.6%), M(0.5%), I(0.2%), K(0.2%), X(0.2%), A(0.1%), V(0.1%), W(0.1%) | L-P(0.5%), P-S(0.5%), L-R(0.4%), P-T(0.4%), L-M(0.3%), P-Q(0.2%), L-F(0.2%), L-I(0.2%), L-Q(0.2%), R-Q(0.2%), F-I(0.1%), L-S(0.1%), L-V(0.1%), P-L(0.1%), P-X(0.1%), R-W(0.1%), T-A(0.1%), T-K(0.1%) |
| | 56 | T(83.1%), I(16.2%), X(0.7%) | T(58.9%), I(33.6%), K(2.3%), M(2.2%), L(2%), V(0.5%), A(0.3%), X(0.1%) | human-human | T(83.1%), I(16.6%), X(0.3%) | I-T(0.7%), T-I(0.3%), T-X(0.3%) |
| | | | | human-swine | T(66.7%), I(28.1%), K(5.3%) | T-K(5.3%), T-I(3.5%) |
| | | | | swine-human | T(72.7%), I(27.3%) | T-I(4.5%) |
| | | | | swine-swine | T(58.2%), I(36.1%), L(2%), M(1.8%), K(1.2%), V(0.3%), A(0.2%), X(0.1%) | I-M(0.7%), T-I(0.7%), T-K(0.7%), I-T(0.4%), I-V(0.3%), I-K(0.2%), T-A(0.2%), I-L(0.1%), I-X(0.1%), T-M(0.1%) |
| | 86 | N(94.6%), I(3.4%), D(1.4%), K(0.7%) | N(80.5%), S(11.2%), D(5.7%), K(0.8%), T(0.8%), H(0.3%), I(0.3%), R(0.3%), E(0.1%) | human-human | N(96.3%), I(2.7%), D(0.7%), K(0.3%) | N-K(0.3%) |
| | | | | human-swine | N(98.2%), S(1.8%) | N-S(1.8%) |
| | | | | swine-human | N(90.9%), D(4.5%), I(4.5%) | N-I(4.5%) |
| | | | | swine-swine | N(83.1%), S(10.1%), D(5.2%), K(0.5%), | N-S(2%), N-D(1.5%), N-T(0.5%), S-N(0.4%), N- |

| | | | | |
|---|---|---|---|---|
| | | | T(0.5%), H(0.2%), I(0.2%), R(0.2%), E(0.1%) | K(0.3%), N-H(0.2%), N-I(0.2%), D-E(0.1%), D-N(0.1%), K-R(0.1%), S-R(0.1%) |
| 88 | S(98.6%), T(1.4%) | S(83.1%), A(14.8%), L(1.4%), E(0.3%), T(0.3%), K(0.1%), V(0.1%) | human-human | S(99.3%), T(0.7%) | NA |
| | | | human-swine | S(98.2%), L(1.8%) | S-L(1.8%) |
| | | | swine-human | S(95.5%), T(4.5%) | A-T(4.5%) |
| | | | swine-swine | S(82.2%), A(16.4%), L(0.9%), E(0.2%), T(0.2%), K(0.1%), V(0.1%) | S-L(0.4%), S-A(0.2%), A-E(0.2%), A-T(0.2%), A-K(0.1%), A-V(0.1%), L-S(0.1%) |
| 150 | H(62.2%), R(37.8%) | H(96.6%), R(2.7%), N(0.4%), X(0.1%), Y(0.1%) | human-human | H(64.9%), R(35.1%) | H-R(0.7%), R-H(0.3%) |
| | | | human-swine | H(89.5%), R(10.5%) | NA |
| | | | swine-human | H(95.5%), R(4.5%) | NA |
| | | | swine-swine | H(97.6%), R(2%), N(0.2%), X(0.1%), Y(0.1%) | H-N(0.2%), H-R(0.2%), R-H(0.2%), H-X(0.1%), H-Y(0.1%) |
| 258 | E(85.1%), K(9.5%), V(4.1%), M(0.7%), R(0.7%) | E(78.9%), K(13.6%), V(6.4%), R(0.7%), N(0.3%), Q(0.3%) | human-human | E(92.6%), K(5.4%), V(2%) | E-K(0.3%) |
| | | | human-swine | E(89.5%), K(5.3%), V(5.3%) | E-K(3.5%), E-V(1.8%) |
| | | | swine-human | E(50%), K(22.7%), V(18.2%), M(4.5%), R(4.5%) | E-K(9.1%), K-R(4.5%), V-M(4.5%) |
| | | | swine-swine | E(81%), K(11.6%), V(6.6%), R(0.5%), N(0.2%), Q(0.2%) | E-K(4.1%), E-V(0.2%), E-N(0.2%), E-Q(0.2%), E-R(0.2%), K-R(0.2%), K-E(0.1%), V-E(0.1%) |
| 336 | H(63.5%), N(20.3%), Y(12.8%), R(2%), D(0.7%), P(0.7%) | N(47%), H(27%), Y(20.6%), D(2.7%), R(2.4%), F(0.3%) | human-human | H(66.2%), Y(17.9%), N(15.2%), P(0.3%), R(0.3%) | H-N(0.7%), Y-H(0.7%), H-P(0.3%), H-R(0.3%), N-Y(0.3%) |
| | | | human-swine | H(59.6%), Y(24.6%), N(12.3%), D(1.8%), R(1.8%) | H-N(3.5%), H-R(1.8%), N-D(1.8%) |
| | | | swine-human | H(40.9%), N(36.4%), R(9.1%), Y(9.1%), D(4.5%) | H-N(4.5%) |
| | | | swine-swine | N(49.3%), H(26.2%), Y(20.2%), D(2%), R(2%), F(0.2%) | H-N(1.7%), H-Y(1.4%), N-D(0.7%), H-R(0.5%), Y-N(0.5%), Y-D(0.2%), N-Y(0.1%), Y-F(0.1%), Y-H(0.1%), Y-R(0.1%) |
| 370 | S(56.1%), L(39.9%), F(4.1%) | L(53.5%), S(39.3%), F(3.5%), V(2.2%), A(0.4%), I(0.4%), T(0.3%), P(0.1%), Q(0.1%), | human-human | S(52.4%), L(43.2%), F(4.4%) | L-S(1.4%), S-L(0.7%), L-F(0.3%) |
| | | | human-swine | S(50.9%), L(38.6%), F(8.8%), V(1.8%) | L-S(7%), L-F(5.3%), F-S(1.8%), L-V(1.8%) |
| | | | swine-human | L(50%), S(45.5%), F(4.5%) | L-F(4.5%) |
| | | | swine-swine | L(58%), S(36.7%), F(2.6%), V(1.9%), A(0.2%), I(0.2%), T(0.2%), P(0.1%), Q(0.1%), X(0.1%) | L-S(2.8%), L-F(0.7%), S-L(0.5%), L-V(0.4%), F-L(0.2%), L-I(0.2%), S-A(0.2%), S-T(0.2%), L-Q(0.1%), S-P(0.1%), S-V(0.1%), S-X(0.1%), V-A(0.1%), V-I(0.1%) |
| 399 | D(69.6%), E(20.3%), G(8.8%), K(0.7%), Y(0.7%) | D(58.7%), E(27.5%), G(11.2%), K(1.2%), N(0.5%), V(0.3%), A(0.1%), S(0.1%), X(0.1%) | human-human | D(74.7%), E(16.2%), G(8.8%), Y(0.3%) | D-E(0.7%), D-Y(0.3%) |
| | | | human-swine | D(64.9%), E(22.8%), G(12.3%) | G-D(3.5%), D-E(1.8%) |
| | | | swine-human | E(45.5%), D(36.4%), G(13.6%), K(4.5%) | E-K(4.5%) |
| | | | swine-swine | D(59.6%), E(28.3%), G(10.4%), K(1%), N(0.3%), V(0.2%), A(0.1%), S(0.1%), X(0.1%), Y(0.1%) | D-G(0.8%), D-E(0.5%), G-D(0.5%), E-G(0.5%), E-K(0.4%), D-N(0.3%), E-D(0.2%), D-V(0.2%), D-A(0.1%), D-Y(0.1%), E-X(0.1%), G-S(0.1%) |

| | | | | | | |
|---|---|---|---|---|---|---|
| | 435 | E(54.7%), R(21.6%), K(15.5%), A(8.1%) | R(42.1%), K(27.5%), E(24.5%), A(4.7%), D(0.4%), G(0.1%), I(0.1%), N(0.1%), Q(0.1%) | human-human | E(59.5%), R(21.3%), K(13.2%), A(6.1%) | E-R(0.7%), E-K(0.3%), K-E(0.3%), R-K(0.3%) |
| | | | | human-swine | E(38.6%), K(29.8%), R(28.1%), A(3.5%) | E-K(7%) |
| | | | | swine-human | K(36.4%), R(36.4%), A(13.6%), E(13.6%) | NA |
| | | | | swine-swine | R(45%), K(26.7%), E(22.7%), A(4.8%), D(0.3%), G(0.1%), I(0.1%), N(0.1%), Q(0.1%), T(0.1%) | R-K(0.8%), K-E(0.3%), K-R(0.3%), E-A(0.2%), A-T(0.1%), E-G(0.1%), K-N(0.1%), K-Q(0.1%), R-I(0.1%) |
| **M1** | 30 | D(55.9%), S(29.7%), G(10.3%), N(3.4%), Y(0.7%) | D(41.6%), G(35%), S(22.4%), K(0.3%), N(0.3%), X(0.3%) | human-human | D(59.6%), S(31.3%), G(6.5%), N(2.2%), Y(0.4%) | D-N(1.5%), G-S(0.7%), D-S(0.4%), D-Y(0.4%), S-D(0.4%), S-N(0.4%) |
| | | | | human-swine | D(56.4%), S(35.9%), G(5.1%), N(2.6%) | NA |
| | | | | swine-human | G(44.4%), D(27.8%), S(27.8%) | G-S(5.6%) |
| | | | | swine-swine | D(40.8%), G(40.5%), S(18.3%), K(0.2%), X(0.2%) | G-S(0.8%), D-G(0.4%), D-S(0.2%), S-K(0.2%), S-X(0.2%) |
| | 95 | R(62.8%), K(37.2%) | R(64.4%), K(35.6%) | human-human | R(62.2%), K(37.8%) | R-K(1.5%), K-R(0.7%) |
| | | | | human-swine | R(53.8%), K(46.2%) | R-K(2.6%) |
| | | | | swine-human | R(83.3%), K(16.7%) | NA |
| | | | | swine-swine | R(67.7%), K(32.3%) | R-K(2.1%), K-R(0.6%) |
| | 96 | A(100%) | A(97.4%), G(2%), S(0.7%) | human-human | A(100%) | NA |
| | | | | human-swine | A(100%) | NA |
| | | | | swine-human | A(100%) | NA |
| | | | | swine-swine | A(97.5%), G(1.9%), S(0.6%) | A-G(0.4%), A-S(0.2%) |
| | 115 | V(56.6%), I(42.8%), M(0.7%) | V(87.5%), I(12.5%) | human-human | V(53.8%), I(45.8%), M(0.4%) | V-I(0.7%), I-M(0.4%), I-V(0.4%) |
| | | | | human-swine | V(69.2%), I(30.8%) | NA |
| | | | | swine-human | V(94.4%), I(5.6%) | NA |
| | | | | swine-swine | V(91.6%), I(8.4%) | V-I(0.8%) |
| | 137 | T(55.9%), A(44.1%) | T(90.1%), A(9.6%), K(0.3%) | human-human | T(53.8%), A(46.2%) | T-A(0.7%), A-T(0.4%) |
| | | | | human-swine | T(69.2%), A(30.8%) | NA |
| | | | | swine-human | T(88.9%), A(11.1%) | T-A(5.6%) |
| | | | | swine-swine | T(94.7%), A(5.2%), K(0.2%) | T-A(0.2%), T-K(0.2%) |
| | 139 | T(99.3%), A(0.7%) | T(93.1%), A(6.9%) | human-human | T(100%) | NA |
| | | | | human-swine | T(100%) | NA |
| | | | | swine-human | T(94.4%), A(5.6%) | NA |
| | | | | swine-swine | T(91.8%), A(8.2%) | A-T(0.4%), T-A(0.2%) |
| | 208 | Q(93.8%), K(4.8%), R(1.4%) | Q(99.7%), K(0.3%) | human-human | Q(94.5%), K(4.4%), R(1.1%) | Q-K(1.1%), K-Q(0.4%), Q-R(0.4%) |
| | | | | human-swine | Q(100%) | NA |
| | | | | swine-human | Q(100%) | NA |
| | | | | swine-swine | Q(99.8%), K(0.2%) | Q-K(0.2%) |
| | 213 | V(99.3%), I(0.7%) | V(95.4%), I(4.6%) | human-human | V(99.6%), I(0.4%) | NA |
| | | | | human-swine | V(100%) | NA |
| | | | | swine-human | V(100%) | NA |
| | | | | swine-swine | V(95.6%), I(4.4%) | V-I(1.1%) |
| | 227 | A(66.9%), T(33.1%) | A(72.6%), T(25.7%), N(1%), S(0.7%) | human-human | A(67.3%), T(32.7%) | A-T(2.2%) |
| | | | | human-swine | A(59%), T(35.9%), N(5.1%) | A-T(5.1%), T-N(5.1%) |
| | | | | swine-human | A(88.9%), T(11.1%) | NA |
| | | | | swine-swine | A(78.1%), T(21.2%), S(0.6%), N(0.2%) | A-T(2.3%), A-N(0.2%), T-S(0.2%) |
| | 248 | | | human-human | M(92%), I(8%) | NA |

| | | | | | | |
|---|---|---|---|---|---|---|
| | | M(85.5%), I(14.5%) | M(57.8%), I(41.9%), T(0.3%) | human-swine | M(97.4%), I(2.6%) | NA |
| | | | | swine-human | I(61.1%), M(38.9%) | I-M(5.6%) |
| | | | | swine-swine | I(50.2%), M(49.6%), T(0.2%) | I-M(1%), M-I(0.6%), M-T(0.2%) |
| | 252 | K(100%) | K(97.7%), R(2.3%) | human-human | K(100%) | NA |
| | | | | human-swine | K(97.4%), R(2.6%) | K-R(2.6%) |
| | | | | swine-human | K(100%) | NA |
| | | | | swine-swine | K(98.5%), R(1.5%) | K-R(0.8%) |
| **M2** | 11 | I(60.7%), T(39.3%) | T(61.7%), I(38%), F(0.3%) | human-human | I(62.2%), T(37.8%) | T-I(1.8%), I-T(0.4%) |
| | | | | human-swine | I(64.1%), T(33.3%), F(2.6%) | I-F(2.6%), I-T(2.6%) |
| | | | | swine-human | T(66.7%), I(33.3%) | T-I(5.6%) |
| | | | | swine-swine | T(67.7%), I(32.3%) | T-I(1.9%), I-T(0.8%) |
| | 13 | N(69.7%), S(29%), K(1.4%) | N(77.6%), S(19.8%), K(1.3%), R(0.7%), D(0.3%), T(0.3%) | human-human | N(66.9%), S(32.7%), K(0.4%) | S-N(2.2%), N-S(1.1%), N-K(0.4%) |
| | | | | human-swine | N(71.8%), S(25.6%), R(2.6%) | S-N(2.6%), S-R(2.6%) |
| | | | | swine-human | N(77.8%), S(16.7%), K(5.6%) | NA |
| | | | | swine-swine | N(81.1%), S(17.4%), K(1%), D(0.2%), R(0.2%), T(0.2%) | N-S(2.3%), S-N(1.5%), N-K(0.8%), N-D(0.2%), N-T(0.2%), S-R(0.2%) |
| | 19 | C(85.5%), Y(14.5%) | C(57.8%), Y(41.3%), S(0.7%), R(0.3%) | human-human | C(92%), Y(8%) | NA |
| | | | | human-swine | C(97.4%), Y(2.6%) | NA |
| | | | | swine-human | Y(61.1%), C(38.9%) | Y-C(5.6%) |
| | | | | swine-swine | Y(49.8%), C(49.6%), S(0.4%), R(0.2%) | Y-C(1%), C-Y(0.6%), Y-S(0.4%), C-R(0.2%) |
| | 27 | V(79.3%), A(10.3%), I(6.9%), T(3.4%) | V(53.1%), A(27.7%), I(15.2%), T(3.3%), S(0.7%) | human-human | V(83.3%), I(7.3%), A(6.9%), T(2.5%) | V-I(1.5%), V-A(0.4%), V-T(0.4%) |
| | | | | human-swine | V(61.5%), I(20.5%), A(17.9%) | I-V(2.6%), V-A(2.6%) |
| | | | | swine-human | V(50%), A(38.9%), I(5.6%), T(5.6%) | V-I(5.6%) |
| | | | | swine-swine | V(56.7%), A(26.7%), I(13.5%), T(2.7%), S(0.4%) | V-A(2.1%), V-I(1.1%), I-T(0.8%), A-T(0.4%), A-V(0.4%), I-V(0.4%), A-S(0.2%), I-S(0.2%), V-T(0.2%) |
| | 28 | I(49.7%), V(35.9%), A(9%), N(2.1%), T(2.1%), D(0.7%), F(0.7%) | I(44.2%), T(16.5%), A(14.2%), V(11.2%), D(8.3%), N(3.3%), F(0.7%), M(0.7%), E(0.3% | human-human | I(52%), V(37.8%), A(9.1%), T(0.7%), F(0.4%) | I-V(0.7%), V-I(0.7%), I-F(0.4%) |
| | | | | human-swine | I(46.2%), V(28.2%), A(23.1%), D(2.6%) | A-D(2.6%) |
| | | | | swine-human | I(50%), N(16.7%), A(11.1%), T(11.1%), D(5.6%), V(5.6%) | T-I(5.6%), T-N(5.6%) |
| | | | | swine-swine | I(46.6%), T(17.4%), A(13.7%), D(10.1%), V(7.1%), N(3.6%), F(0.6%), M(0.4%), E(0.2%), L(0.2%), Y(0.2%) | I-T(1%), V-I(0.8%), A-T(0.4%), D-V(0.4%), I-M(0.4%), I-V(0.4%), A-I(0.2%), A-V(0.2%), D-E(0.2%), D-I(0.2%), D-N(0.2%), D-Y(0.2%), I-F(0.2%), I-L(0.2%), N-V(0.2%), T-I(0.2%), T-N(0.2%) |
| | 43 | L(65.5%), T(25.5%), I(6.2%), F(1.4%), P(0.7%), V(0.7%) | L(78.5%), T(16.2%), A(1.3%), F(1.3%), V(1.3%), I(0.7%), S(0.7%) | human-human | L(62.5%), T(29.5%), I(5.1%), V(1.1%), F(0.7%), P(0.7%), A(0.4%) | L-I(0.7%), P-L(0.4%), T-A(0.4%), T-I(0.4%), T-P(0.4%), V-T(0.4%) |
| | | | | human-swine | L(56.4%), T(33.3%), V(5.1%), A(2.6%), I(2.6%) | NA |
| | | | | swine-human | L(77.8%), T(11.1%), F(5.6%), V(5.6%) | L-F(5.6%), L-V(5.6%) |

| | | | | | | |
|---|---|---|---|---|---|---|
| | | | | swine-swine | L(84.5%), T(12.4%), A(0.8%), F(0.8%), V(0.8%), S(0.6%), I(0.2%) | L-F(0.8%), T-A(0.4%), L-I(0.2%), L-T(0.2%), T-S(0.2%) |
| | 55 | F(82.8%), L(15.2%), I(2.1%) | F(51.8%), L(45.5%), I(1.7%), S(0.3%), V(0.3%), Y(0.3%) | human-human | F(89.8%), L(9.1%), I(1.1%) | L-F(0.7%), F-I(0.4%), F-L(0.4%) |
| | | | | human-swine | F(97.4%), L(2.6%) | NA |
| | | | | swine-human | L(55.6%), F(38.9%), I(5.6%) | L-F(5.6%), L-I(5.6%) |
| | | | | swine-swine | L(55%), F(43.1%), I(1.3%), S(0.2%), V(0.2%), Y(0.2%) | L-F(1.3%), F-L(1%), L-I(0.6%), F-I(0.4%), F-S(0.2%), F-V(0.2%), F-Y(0.2%) |
| | 56 | K(88.3%), R(6.9%), E(4.8%) | K(92.7%), E(5.3%), R(1.7%), N(0.3%) | human-human | K(89.5%), R(5.8%), E(4.7%) | K-E(0.4%), K-R(0.4%) |
| | | | | human-swine | K(89.7%), E(7.7%), N(2.6%) | K-E(2.6%), K-N(2.6%) |
| | | | | swine-human | K(100%) | NA |
| | | | | swine-swine | K(94.3%), E(3.8%), R(1.9%) | K-E(0.6%), K-R(0.6%), E-K(0.2%), R-K(0.2%) |
| | 78 | Q(53.8%), K(40%), E(6.2%) | Q(91.7%), K(7.3%), E(1%) | human-human | Q(51.3%), K(42.5%), E(6.2%) | Q-K(0.7%), K-E(0.4%) |
| | | | | human-swine | Q(69.2%), K(28.2%), E(2.6%) | NA |
| | | | | swine-human | Q(94.4%), K(5.6%) | NA |
| | | | | swine-swine | Q(95.8%), K(3.8%), E(0.4%) | K-Q(0.2%), Q-K(0.2%) |
| | 86 | V(56.6%), A(42.1%), T(1.4%) | V(90.8%), A(8.9%), S(0.3%) | human-human | V(54.2%), A(44.7%), T(1.1%) | V-A(0.7%), A-T(0.4%), A-V(0.4%) |
| | | | | human-swine | V(71.8%), A(28.2%) | NA |
| | | | | swine-human | V(94.4%), A(5.6%) | NA |
| | | | | swine-swine | V(95.4%), A(4.4%), S(0.2%) | V-A(0.4%), V-S(0.2%) |
| | 93 | N(63.4%), S(36.6%) | N(91.4%), S(8.6%) | human-human | N(61.1%), S(38.9%) | N-S(1.5%), S-N(1.5%) |
| | | | | human-swine | N(69.2%), S(30.8%) | N-S(2.6%) |
| | | | | swine-human | N(94.4%), S(5.6%) | NA |
| | | | | swine-swine | N(96%), S(4%) | NA |
| **NS1** | 22 | F(61.3%), V(37.1%), L(1.1%), C(0.5%) | F(94.1%), V(4.3%), L(1.1%), I(0.5%) | human-human | F(59.3%), V(39.8%), L(0.9%) | F-V(0.6%), F-L(0.3%) |
| | | | | human-swine | F(79.7%), V(20.3%) | NA |
| | | | | swine-human | F(85.7%), V(11.4%), C(2.9%) | F-C(2.9%) |
| | | | | swine-swine | F(97.1%), V(1.8%), L(0.6%), I(0.4%) | F-L(0.6%), F-V(0.4%), F-I(0.1%) |
| | 53 | D(71%), N(17.7%), E(11.3%) | D(44.5%), E(44.5%), N(8.8%), G(1.1%), K(0.5%), A(0.4%), Y(0.4%) | human-human | D(78.6%), N(17.1%), E(4.3%) | D-N(0.9%), D-E(0.3%), N-D(0.3%) |
| | | | | human-swine | D(89.8%), N(10.2%) | D-N(5.1%) |
| | | | | swine-human | D(54.3%), E(34.3%), N(11.4%) | D-N(2.9%), N-D(2.9%) |
| | | | | swine-swine | E(52.8%), D(37.8%), N(7.8%), G(0.8%), K(0.3%), A(0.2%), Y(0.2%) | D-N(1.5%), E-D(0.4%), E-G(0.4%), D-E(0.3%), E-K(0.3%), D-Y(0.2%), D-A(0.1%), E-A(0.1%), N-D(0.1%) |
| | 70 | K(84.9%), E(10.2%), R(3.2%), G(1.1%), T(0.5%) | K(48.2%), E(45.2%), G(3.2%), R(3%), N(0.4%) | human-human | K(92.2%), E(4.3%), R(3.1%), T(0.3%) | K-E(0.3%), K-T(0.3%) |
| | | | | human-swine | K(91.5%), R(8.5%) | NA |
| | | | | swine-human | K(57.1%), E(28.6%), R(8.6%), G(5.7%) | E-G(2.9%) |
| | | | | swine-swine | E(54.8%), K(40.7%), G(2.4%), R(1.9%), N(0.2%) | E-G(1.3%), K-R(0.6%), E-K(0.5%), K-E(0.2%), E-N(0.1%), G-R(0.1%), K-G(0.1%), K-N(0.1%) |

| | | | | | |
|---|---|---|---|---|---|
| 73 | S(89.2%), Y(8.6%), H(2.2%) | S(51.6%), Y(33.4%), H(10.7%), C(2.1%), F(1.1%), P(0.5%), N(0.2%), Q(0.2%), R(0.2%) | human-human | S(96%), Y(3.4%), H(0.6%) | NA |
| | | | human-swine | S(96.6%), F(1.7%), P(1.7%) | S-F(1.7%), S-P(1.7%) |
| | | | swine-human | S(65.7%), Y(25.7%), H(8.6%) | NA |
| | | | swine-swine | S(44.4%), Y(41%), H(11.4%), C(1.9%), F(0.7%), P(0.2%), N(0.1%), Q(0.1%), R(0.1%) | Y-H(1.2%), Y-C(0.6%), S-P(0.2%), Y-S(0.2%), S-F(0.1%), S-Y(0.1%), Y-N(0.1%), Y-Q(0.1%), Y-R(0.1%) |
| 79 | M(94.1%), I(3.2%), L(1.6%), T(1.1%) | M(59.3%), I(33.6%), L(4.8%), T(1.2%), V(0.7%), K(0.2%), R(0.2%) | human-human | M(98.4%), I(0.6%), L(0.6%), T(0.3%) | M-T(0.3%) |
| | | | human-swine | M(94.9%), I(1.7%), R(1.7%), T(1.7%) | M-I(1.7%), M-R(1.7%), M-T(1.7%) |
| | | | swine-human | M(77.1%), I(14.3%), L(5.7%), T(2.9%) | I-T(2.9%) |
| | | | swine-swine | M(52.5%), I(41.2%), L(5.1%), T(0.6%), V(0.4%), K(0.1%) | I-M(1.4%), I-T(0.6%), I-L(0.2%), I-V(0.2%), M-I(0.2%), M-V(0.2%), L-I(0.1%), M-K(0.1%), M-L(0.1%) |
| 82 | A(87.6%), V(12.4%) | A(98.2%), V(1.8%) | human-human | A(86.6%), V(13.4%) | A-V(0.6%), V-A(0.6%) |
| | | | human-swine | A(93.2%), V(6.8%) | NA |
| | | | swine-human | A(94.3%), V(5.7%) | NA |
| | | | swine-swine | A(99.2%), V(0.8%) | A-V(0.1%) |
| 124 | M(83.3%), I(7.5%), L(5.9%), T(3.2%) | M(79.6%), I(11.8%), L(5.2%), T(1.6%), V(1.2%), A(0.2%), R(0.2%), S(0.2%) | human-human | M(87.9%), I(5.6%), L(3.7%), T(2.8%) | M-I(0.9%), M-T(0.6%), I-T(0.3%), M-L(0.3%) |
| | | | human-swine | M(89.8%), I(6.8%), L(1.7%), T(1.7%) | M-I(5.1%), M-T(1.7%) |
| | | | swine-human | M(74.3%), I(14.3%), L(11.4%) | NA |
| | | | swine-swine | M(81%), I(11.4%), L(4.9%), T(1.4%), V(1%), A(0.1%), R(0.1%), S(0.1%) | M-I(0.5%), M-V(0.5%), I-T(0.2%), M-T(0.2%), I-S(0.1%), M-R(0.1%), T-A(0.1%) |
| 125 | E(64%), D(31.2%), G(2.7%), N(2.2%) | E(51.2%), D(41.6%), N(5.5%), G(1.2%), A(0.2%), V(0.2%) | human-human | E(68.9%), D(28.9%), G(2.2%) | E-D(0.9%), D-E(0.6%), E-G(0.6%) |
| | | | human-swine | E(91.5%), D(8.5%) | E-D(1.7%) |
| | | | swine-human | E(62.9%), D(25.7%), N(11.4%) | D-N(2.9%) |
| | | | swine-swine | D(50.2%), E(43.6%), N(5.2%), G(0.7%), A(0.1%), V(0.1%) | D-N(1.9%), D-E(0.5%), D-G(0.5%), E-G(0.2%), D-A(0.1%), D-V(0.1%), E-D(0.1%), N-D(0.1%) |
| 129 | I(43%), V(24.7%), M(21.5%), T(10.8%) | T(46.2%), I(40.7%), V(8.6%), M(3.8%), A(0.4%), L(0.2%), R(0.2%) | human-human | I(41.6%), V(30.1%), M(23.9%), T(4.3%) | I-M(0.3%), M-I(0.3%), M-T(0.3%), M-V(0.3%), V-I(0.3%), V-M(0.3%) |
| | | | human-swine | I(45.8%), V(35.6%), M(16.9%), T(1.7%) | I-T(1.7%), V-I(1.7%) |
| | | | swine-human | I(57.1%), T(31.4%), M(8.6%), V(2.9%) | I-M(2.9%) |
| | | | swine-swine | T(54.2%), I(38.9%), V(4.8%), M(1.7%), A(0.2%), L(0.1%), R(0.1%) | T-I(1.1%), I-T(0.8%), I-M(0.2%), I-V(0.2%), T-A(0.2%), V-I(0.2%), I-L(0.1%), T-R(0.1%) |
| 171 | N(28%), Y(25.3%), D(21%), I(20.4%), A(2.7%), G(1.1%), E(0.5%), | D(51.4%), N(30.4%), Y(8.6%), I(3.2%), G(2.5%), H(1.2%), E(1.1%), C(0.7%), K(0.4%), | human-human | Y(31.4%), I(23.3%), N(22.7%), D(18.6%), A(2.8%), H(0.6%), F(0.3%), G(0.3%) | I-Y(0.6%), D-A(0.3%), D-G(0.3%), D-N(0.3%), I-F(0.3%), Y-H(0.3%), Y-I(0.3%) |
| | | | human-swine | Y(33.9%), N(25.4%), I(16.9%), D(11.9%), H(6.8%), A(1.7%), C(1.7%), E(1.7%) | Y-H(5.1%), D-E(1.7%), Y-C(1.7%) |
| | | | swine-human | N(51.4%), D(34.3%), I(5.7%), E(2.9%), G(2.9%), Y(2.9%) | D-N(2.9%) |

| | | | | | | |
|---|---|---|---|---|---|---|
| | | | | swine-swine | D(62.1%), N(27.6%), Y(4.8%), G(2%), I(1.2%), E(1%), C(0.5%), H(0.3%), K(0.3%), T(0.2%) | D-N(1.9%), D-G(1%), D-Y(0.3%), Y-H(0.3%), D-E(0.2%), N-T(0.2%), N-G(0.1%), N-I(0.1%), N-K(0.1%), Y-C(0.1%) |
| | 180 | V(87.6%), I(11.8%), F(0.5%) | V(54.6%), I(42%), T(1.6%), A(0.7%), F(0.7%), L(0.4%) | human-human | V(93.5%), I(6.2%), F(0.3%) | V-I(0.6%), V-F(0.3%) |
| | | | | human-swine | V(96.6%), I(3.4%) | V-I(3.4%) |
| | | | | swine-human | V(68.6%), I(31.4%) | NA |
| | | | | swine-swine | V(50.4%), I(47.1%), T(1.2%), A(0.5%), F(0.5%), L(0.2%) | V-I(2.3%), I-T(0.7%), I-V(0.5%), V-A(0.3%), I-F(0.2%), I-L(0.1%), V-F(0.1%), V-L(0.1%) |
| | 183 | G(89.2%), K(10.8%) | G(52.7%), K(45.9%), R(0.9%), E(0.4%), N(0.2%) | human-human | G(96%), K(4%) | NA |
| | | | | human-swine | G(100%) | NA |
| | | | | swine-human | G(65.7%), K(34.3%) | NA |
| | | | | swine-swine | K(53.8%), G(45%), R(0.8%), E(0.2%), N(0.1%) | G-R(0.3%), K-E(0.2%), K-G(0.1%), K-N(0.1%), K-R(0.1%), R-K(0.1%) |
| | 198 | L(73.7%), I(26.3%) | L(89.5%), I(10.2%), V(0.4%) | human-human | L(67.4%), I(32.6%) | L-I(1.2%), I-L(0.3%) |
| | | | | human-swine | L(55.9%), I(44.1%) | L-I(3.4%), I-L(1.7%) |
| | | | | swine-human | L(100%) | NA |
| | | | | swine-swine | L(95.1%), I(4.7%), V(0.2%) | L-I(0.7%), I-L(0.2%), L-V(0.2%) |
| | 206 | S(36.6%), C(26.3%), R(23.7%), I(5.4%), T(5.4%), H(2.2%), L(0.5%) | R(32.1%), T(25.7%), I(19.3%), C(11.8%), S(6.2%), H(1.8%), A(1.1%), L(0.7%), X(0.5% | human-human | S(39.4%), C(32.3%), R(21.7%), I(2.8%), H(2.2%), T(1.2%), L(0.3%) | S-C(0.9%), C-S(0.3%), R-H(0.3%), R-L(0.3%) |
| | | | | human-swine | C(42.4%), R(35.6%), S(20.3%), N(1.7%) | C-S(3.4%), R-C(3.4%), C-R(1.7%), S-C(1.7%), S-N(1.7%) |
| | | | | swine-human | R(45.7%), T(20%), I(14.3%), S(14.3%), C(2.9%), H(2.9%) | R-S(2.9%) |
| | | | | swine-swine | R(32.1%), T(30.9%), I(22.8%), C(6.5%), S(3.8%), H(1.7%), A(0.8%), L(0.7%), X(0.3%), K(0.1%), P(0.1%), Y(0.1%) | T-I(1.4%), R-C(1.1%), I-T(0.5%), R-H(0.4%), R-S(0.4%), T-A(0.4%), R-L(0.3%), I-S(0.2%), T-S(0.2%), T-X(0.2%), C-S(0.1%), H-Y(0.1%), I-K(0.1%), I-X(0.1%), L-I(0.1%), S-I(0.1%), T-P(0.1%) |
| | 227 | X(57%), R(34.4%), G(7.5%), E(0.5%), K(0.5%) | X(58.2%), G(29.3%), R(12.1%), E(0.4%) | human-human | X(58.1%), R(37.6%), G(3.7%), K(0.6%) | R-X(0.9%), R-G(0.3%), R-K(0.3%) |
| | | | | human-swine | X(79.7%), R(18.6%), G(1.7%) | R-G(1.7%) |
| | | | | swine-human | X(65.7%), G(17.1%), R(14.3%), E(2.9%) | G-R(2.9%), R-E(2.9%), R-X(2.9%) |
| | | | | swine-swine | X(52.6%), G(35.1%), R(11.4%), E(0.8%) | G-R(0.8%), G-X(0.6%), R-X(0.6%), X-G(0.2%), E-G(0.1%), E-X(0.1%), R-G(0.1%) |
| **NEP** | 14 | M(73.1%), L(23.7%), T(1.6%), V(1.1%), K(0.5%) | M(88%), L(3.8%), V(3.2%), T(2.3%), K(1.4%), I(1.2%) | human-human | M(71.1%), L(26.4%), T(2.2%), V(0.3%) | L-M(0.6%), M-L(0.6%), M-T(0.3%), M-V(0.3%) |
| | | | | human-swine | M(71.2%), L(18.6%), T(6.8%), K(1.7%), V(1.7%) | M-K(1.7%), M-V(1.7%), T-M(1.7%) |
| | | | | swine-human | M(85.7%), L(5.7%), K(2.9%), T(2.9%), V(2.9%) | NA |
| | | | | swine-swine | M(92.1%), V(2.5%), T(1.8%), L(1.4%), K(1.3%), I(1%) | M-V(1.2%), M-I(0.4%), M-K(0.3%), M-L(0.3%), M-T(0.2%), T-M(0.1%) |
| | 26 | E(80.6%), K(10.8%), G(8.6%) | | human-human | E(88.5%), G(7.5%), K(4%) | E-G(0.6%), G-E(0.3%) |

| | | | | | |
|---|---|---|---|---|---|
| | | E(52%), K(45.5%), G(1.6%), R(0.7%), T(0.2%) | human-swine | E(94.9%), G(5.1%) | E-G(3.4%) |
| | | | swine-human | E(57.1%), K(34.3%), G(8.6%) | E-G(2.9%) |
| | | | swine-swine | K(53.6%), E(45%), G(0.8%), R(0.4%), T(0.1%) | E-G(0.6%), K-R(0.4%), K-E(0.2%), E-K(0.1%), K-T(0.1%) |
| 37 | S(82.8%), A(17.2%) | S(77.5%), A(22.5%) | human-human | S(86.3%), A(13.7%) | NA |
| | | | human-swine | S(76.3%), A(23.7%) | NA |
| | | | swine-human | S(62.9%), A(37.1%) | NA |
| | | | swine-swine | S(79.3%), A(20.7%) | A-S(0.2%), S-A(0.1%) |
| 40 | I(67.7%), L(31.2%), V(1.1%) | L(49.3%), I(46.1%), V(4.6%) | human-human | I(72.4%), L(26.4%), V(1.2%) | L-I(0.3%), V-I(0.3%) |
| | | | human-swine | I(91.5%), L(8.5%) | NA |
| | | | swine-human | I(57.1%), L(40%), V(2.9%) | NA |
| | | | swine-swine | L(57.1%), I(37.7%), V(5.2%) | L-I(0.6%), I-V(0.2%), I-L(0.1%), V-I(0.1%), V-L(0.1%) |
| 48 | A(57.5%), T(40.3%), S(1.6%), N(0.5%) | A(58.8%), T(35%), S(2.7%), N(2%), V(0.7%), X(0.4%), E(0.2%), P(0.2%), Y(0.2%) | human-human | A(54%), T(43.8%), S(1.9%), N(0.3%) | A-T(0.6%), T-A(0.3%), T-N(0.3%) |
| | | | human-swine | T(67.8%), A(30.5%), S(1.7%) | A-S(1.7%), T-A(1.7%) |
| | | | swine-human | A(57.1%), T(40%), S(2.9%) | A-T(2.9%), N-S(2.9%) |
| | | | swine-swine | A(66.2%), T(28.9%), S(2.1%), N(1.8%), V(0.4%), X(0.2%), E(0.1%), P(0.1%), Y(0.1%) | T-A(1%), A-T(0.8%), A-S(0.4%), A-V(0.4%), T-N(0.4%), A-X(0.2%), A-E(0.1%), S-Y(0.1%), T-P(0.1%), T-S(0.1%) |
| 49 | V(86.6%), L(11.3%), I(2.2%) | V(49.1%), L(47.9%), I(2.1%), A(0.5%), K(0.2%), X(0.2%) | human-human | V(93.5%), L(4.3%), I(2.2%) | V-I(0.3%), V-L(0.3%) |
| | | | human-swine | V(98.3%), I(1.7%) | V-I(1.7%) |
| | | | swine-human | V(62.9%), L(34.3%), I(2.9%) | NA |
| | | | swine-swine | L(56%), V(41.5%), I(1.8%), A(0.4%), K(0.1%), X(0.1%) | V-I(0.4%), V-L(0.4%), L-V(0.2%), V-A(0.2%), L-K(0.1%), L-X(0.1%) |
| 52 | M(83.3%), T(12.4%), L(4.3%) | T(48.8%), M(47.9%), K(1.4%), V(0.9%), L(0.4%), A(0.2%), I(0.2%), N(0.2%), R(0.2%) | human-human | M(89.8%), T(5.9%), L(4.3%) | M-L(0.6%), L-M(0.3%) |
| | | | human-swine | M(93.2%), L(1.7%), R(1.7%), T(1.7%), V(1.7%) | M-R(1.7%), M-T(1.7%), M-V(1.7%) |
| | | | swine-human | M(62.9%), T(37.1%) | NA |
| | | | swine-swine | T(57.7%), M(40.4%), K(1%), V(0.5%), A(0.1%), I(0.1%), L(0.1%), N(0.1%) | T-M(1%), T-K(0.7%), M-T(0.4%), M-V(0.3%), M-I(0.1%), M-L(0.1%), T-A(0.1%), T-N(0.1%) |
| 57 | Y(55.4%), S(26.9%), L(17.2%), F(0.5%) | Y(55.2%), S(33.8%), F(5.4%), L(3.2%), H(1.8%), T(0.5%), A(0.2%) | human-human | Y(57.5%), S(23.3%), L(19.3%) | L-Y(0.3%), S-L(0.3%) |
| | | | human-swine | Y(78%), L(16.9%), S(3.4%), H(1.7%) | Y-H(1.7%) |
| | | | swine-human | Y(62.9%), S(28.6%), L(5.7%), F(2.9%) | NA |
| | | | swine-swine | Y(50.5%), S(40.7%), F(5.5%), H(1.5%), L(1.2%), T(0.5%), A(0.1%) | S-F(1.1%), Y-H(0.5%), F-S(0.3%), S-Y(0.3%), F-L(0.1%), S-A(0.1%), S-T(0.1%), Y-S(0.1%) |
| 60 | N(50.5%), S(40.3%), H(7%), D(1.6%), T(0.5%) | N(57.7%), S(35.9%), D(4.8%), T(0.5%), H(0.4%), R(0.4%), C(0.2%), I(0.2%) | human-human | N(46.9%), S(45.3%), H(7.1%), D(0.6%) | N-S(0.6%) |
| | | | human-swine | S(55.9%), N(39%), C(1.7%), H(1.7%), R(1.7%) | S-N(3.4%), S-C(1.7%), S-R(1.7%) |
| | | | swine-human | N(57.1%), S(31.4%), D(5.7%), H(2.9%), T(2.9%) | N-H(2.9%), N-S(2.9%), N-T(2.9%), S-N(2.9%) |

| | | | | | | |
|---|---|---|---|---|---|---|
| | | | | swine-swine | N(58.6%), S(35%), D(5.7%), T(0.4%), H(0.1%), I(0.1%), R(0.1%) | S-N(2.1%), N-S(1.2%), D-N(0.5%), N-D(0.2%), N-T(0.2%), D-S(0.1%), N-H(0.1%), S-I(0.1%), S-R(0.1%) |
| | 89 | A(36%), T(32.8%), I(17.2%), M(8.1%), L(2.2%), V(2.2%), E(1.1%), | I(37.3%), A(23.8%), T(12.3%), L(9.5%), M(9.3%), V(6.2%), E(0.9%), K(0.4%), S(0.4%) | human-human | A(42.5%), T(33.2%), I(12.4%), M(8.7%), E(1.2%), V(0.9%), L(0.6%), S(0.3%) | T-A(0.9%), A-V(0.6%), A-E(0.3%), A-T(0.3%), I-M(0.3%), M-S(0.3%), T-I(0.3%) |
| | | | | human-swine | A(45.8%), T(27.1%), M(13.6%), V(6.8%), E(3.4%), I(1.7%), S(1.7%) | A-T(6.8%), A-V(5.1%), A-E(1.7%), A-M(1.7%), A-S(1.7%), M-A(1.7%) |
| | | | | swine-human | I(28.6%), T(25.7%), A(20%), M(11.4%), L(8.6%), V(5.7%) | A-T(2.9%), I-T(2.9%) |
| | | | | swine-swine | I(45.1%), A(20.7%), L(10.8%), M(9.5%), T(8.9%), V(4.3%), E(0.3%), K(0.2%), S(0.1%) | I-T(1.1%), A-V(1%), M-I(0.5%), A-T(0.4%), I-M(0.4%), A-E(0.3%), I-L(0.3%), I-V(0.3%), T-M(0.3%), L-I(0.2%), M-T(0.2%), M-V(0.2%), T-K(0.2%), A-M(0.1%), L-S(0.1%), T-I(0.1%) |
| | 120 | L(94.1%), F(5.9%) | L(64.1%), F(35.7%), I(0.2%) | human-human | L(98.4%), F(1.6%) | L-F(0.3%) |
| | | | | human-swine | L(96.6%), F(3.4%) | L-F(3.4%) |
| | | | | swine-human | L(80%), F(20%) | F-L(2.9%) |
| | | | | swine-swine | L(58.9%), F(41%), I(0.1%) | F-L(0.8%), L-F(0.4%), F-I(0.1%) |
